# Outline and divergence time of subkingdom Mucoromyceta: two new phyla, five new orders, six new families and seventy-three new species

**DOI:** 10.1101/2022.07.05.498902

**Authors:** Heng Zhao, Yu-Cheng Dai, Xiao-Yong Liu

## Abstract

Zygomycetes are phylogenetically early diverged, ecologically diverse, industrially valuable, agriculturally beneficial, and clinically pathogenic fungi. Although new phyla and subphyla have been constantly established to accommodate specific members and a subkingdom, Mucoromyceta, was erected to unite core zygomycetous fungi, their phylogenetic relationships have not been well resolved. Taking account of the information of monophyly and divergence time estimated from ITS and LSU rDNA sequences, the present study updates the classification framework of the subkingdom Mucoromyceta from the phylum down to the generic rank: six phyla (including two new phyla Endogonomycota and Umbelopsidomycota), eight classes, 15 orders (including five new orders Claroideoglomerales, Cunninghamellales, Lentamycetales, Phycomycetales and Syncephalastrales), 41 families (including six new families Circinellaceae, Gongronellaceae, Protomycocladaceae, Rhizomucoraceae, Syzygitaceae and Thermomucoraceae), and 121 genera. The taxonomic hierarchy was calibrated with estimated divergence times: phyla 810–639 Mya, classes 651–585 Mya, orders 570–400 Mya, and families 488–107 Mya. Along with this outline, 71 genera are annotated and 73 new species are described. In addition, three new combinations are proposed. In this paper, we update the taxonomic backbone of the subkingdom Mucoromyceta and reinforce its phylogeny. We also contribute numerous new taxa and enrich the diversity of Mucoromyceta.

## Introduction

As an early-diverging group of fungi, zygomycetes are considered an evolutionary transition from ancestral aquatic zoosporic chytrids to terrestrial nonflagellated higher fungi (Spatafora *et al*. 2016, Voigt *et al*. 2021), playing a critical role at the dawn of terrestrial life (Berbee *et al*. 2017). They are ecologically diverse, including pathogens of animals and plants, parasites on other fungi, mycorrhizal symbionts, endophytes, and decomposers of organic matter (Domsch *et al*. 1980, Yao *et al*. 1996, Kamei 2000, Zheng *et al*. 2009, Benny *et al*. 2014, Rashmi *et al* 2019, Wagner *et al*. 2020, Voigt *et al*. 2021). It is well-known that some zygomycetous fungi are used not only to synthesize industrially valuable products such as glucoamylase, laccase, lipase, polygalacturonase, pectinase, proteases, β-carotene, carotenoids, chitosan, fatty acids, biodiesel, progesterone and testosterone, but also to ferment traditional foods like Asian soybean-based staples and European cheese (Davoust & Persson 1992, Papp *et al*. 2009, Liu & Voigt 2010, Hermet *et al*. 2012, Hong *et al*. 2012, Kristanti *et al*. 2016, Kosa *et al*. 2018, Zoghi *et al*. 2019, de Carvalho Tavares *et al*. 2020, Luo *et al*. 2020, Jia *et al*. 2021, Sandmann 2021, Zhao *et al*. 2021a). Many species are agriculturally useful, being able to solubilize phosphate, degrade metalaxyl, and promote crop growth (Doilom *et al*. 2020, Li *et al*. 2020, Martins *et al*. 2020, Ozimek & Hanaka 2021). A few species are widely applied in pharmaceutical industries (Abrashev *et al*. 2021). Several species are also model organisms for dimorphism, drug metabolism/biotransformation, phototropism/light-growth response, and forcible ejection (Page 1964, Schulz *et al*. 1974, Galland & Lipson 1985, Khale & Deshpande 1992, Sepuri & Vidyavathi 2008). However, some zygomycetes are still notorious for infecting humans and causing deadly diseases, such as the mucormycosis during the COVID-19 pandemic (Kwon-Chung 2012, Cheng *et al*. 2017, Morin-Sardin *et al*. 2017, Hallur *et al*. 2021, Mahalaxmi *et al*. 2021).

Zygomycetous fungi are characterized by 1) fast growing mycelia, 2) coenocytic aseptate filamentous hyphae except for irregular septa with age or at the point of branching, 3) asexual intercalary or terminal chlamydospores as well as sporangiospores in a sack such as multi-spored sporangia, one to multi-spored sporangiola, and one to few-spored merosporangiola, and 4) sexual zygospores or azygospores in a zygosporangium or sporocarp (Berbee *et al*. 2017). Zygomycetes were once classified in the phylum Zygomycota (Moreau 1952), though it is invalid and illegal because of a lack of Latin diagnosis or description (Liu & Voigt 2010, Turland *et al*. 2018). Half a century later, the phylum Zygomycota was rejected by Hibbett *et al*. (2007) and its members were assigned to the phylum Glomeromycota and four subphyla *incertae sedis*, viz., Entomophthoromycotina, Kickellomycotina, Mucoromycotina and Zoopagomycotina. Due to a paraphyletic grade relationship to the Dikarya in a genome-scale phylogeny, they were reclassified in two monophyletic phyla Mucoromycota and Zoopagomycota (Spatafora *et al*. 2016). The former phylum accommodated the core members of traditional zygomycetous fungi, comprising three subphyla Glomeromycotina, Mortierellomycotina and Mucoromycotina (Spatafora *et al*. 2016). Recently, it was further raised to the subkingdom rank as Mucoromyceta accommodating Calcarisporiellomycota, Glomeromycota, Mortierellomycota and Mucoromycota (Tedersoo *et al*. 2018). However, the subkingdom was still a paraphyletic grade to the subkingdom Dikarya rather than a monophyletic clade, and the placement of the Glomeromycota and Mortierellomycota was incongruent among several studies (James *et al*. 2006, Oehl *et al*. 2011a, b, c, Spatafora *et al*. 2016, Tedersoo *et al*. 2018, Wijayawardene *et al*. 2020).

Nowadays, it is more and more popular to incorporate divergence time in ranking fungal taxa, especially in Basidiomycota (Hyde *et al*. 2017, Zhao *et al*. 2017, He *et al*. 2019, Wang *et al*. 2021a). Zygomycetous fossil evidence has been accumulated recently, making it possible to date the molecular evolution within zygomycetes (Batra *et al*. 1964, Baxter 1975, Taylor & Alexander 2005, Krings *et al*. 2011, 2018, Taylor *et al*. 2014, Gan *et al*. 2021). The present study adopts molecular clock hypothesis for further refining the Mucoromyceta with a hierarchical outline from class to genus.

Since the 1780’s, a total of 792 species has been proposed for the subkingdom Mucoromyceta, though much less than those in the subkingdom Dikarya (https://nmdc.cn/fungarium/fungi/worldfungi, accessed on November 12, 2021, Fig. 1 and Table S1). Voigt *et al*. (2021) supposed that species in Mucoromyceta could be significantly higher than what was known to exist. Inspiringly, new species have been described in the subkingdom (Hurdeal *et al*. 2021, Nguyen *et al*. 2021, Zhao *et al*. 2021b, Zong *et al*. 2021) at an accelerated pace after the inflection point around the year of 2000 on the trend curve of species number (Fig. 1), which is due to advances in molecular phylogeny (Dai *et al*. 2015, Voigt *et al*. 2021). In this paper we describe new species in the subkingdom Mucoromyceta, along with the proposal of an updated higher-rank taxonomic framework. The internal transcribed spacer (ITS) region of the ribosomal RNA gene (rDNA) has been recommended as a universal fungal DNA barcode for identification at the species rank (Schoch *et al*. 2012, Nilsson *et al*. 2014). In the present study, all new species are identified with a combination of morphological traits and ITS rDNA sequences.

**Fig. 1.**
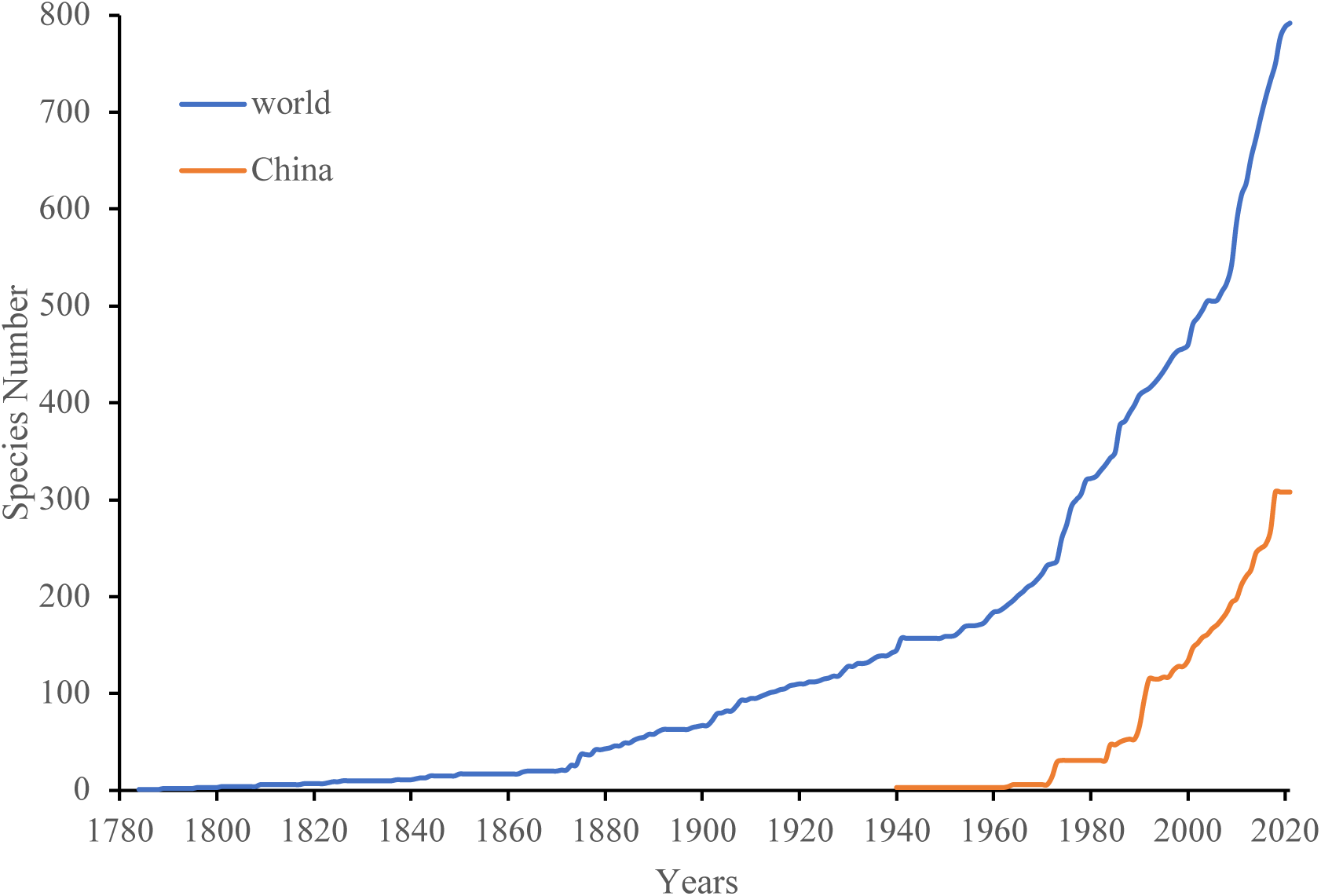
Trends in accumulative number of species of Mucoromyceta recorded in the world and China.

## Materials and Methods

### Strains and culture conditions

Samples including soil, plant debris, leaves, rotten wood, flowers, macrofungi, paper and animal dung were collected, and strains were isolated according to methods described previously (Zhao *et al*. 2021b, Zong *et al*. 2021). In brief, approximately 1 g of sample was suspended and shaken with sterilized water. About 200 μL of the suspension was then incubated at room temperature with potato dextrose agar (PDA: 20 g / L glucose, 200 g / L potato, 20 g / L agar, pH7), and examined daily with a stereo microscope (SMZ1500, Nikon Corporation, Japan). Selective antibiotics (100 mg / mL streptomycin sulfate, and 100 mg / mL ampicillin) were added to the PDA medium to inhibit bacterial growth. Living and dried cultures for nomenclatural types were deposited in the China General Microbiological Culture Collection Center, Beijing, China (CGMCC) and the Herbarium Mycologicum Academiae Sinicae, Beijing, China (HMAS), respectively. Living cultures numbered with initial acronyms “XY” were preserved in both Shandong Normal University and Beijing Forestry University.

### Morphology and growth temperature tests

Pure cultures were established with PDA or synthetic mucor agar (SMA: 20 g / L dextrose, 2 g / L asparagine, 0.5 g / L KH_2_PO_4_, 0.25 g / L MgSO_4_·H_2_O, 0.5 mg / L thiamin chloride, 20 g / L agar, pH7) or malt-extract agar (MEA: 20 g / L malt-extract, 20 g / L agar, pH7) for morphological observation. Strains were incubated inversely at 20 °C, 27 °C or 30 °C for two weeks, and examined and photographed daily under a light microscope (Axio Imager A2, Carl Zeiss, Oberkochen, Germany). Colonies reaching 90 mm in diameter within three days were referred to as ‘fast growing’, and if more than seven days as ‘slow growing’. Colony colors were taken from Ridgway (1912). The sizes of microscopic features were measured with a software platform (Leica Application Suite v4.5, Leica Microsystems Inc., Buffalo Grove, Illinois, USA). In order to determine the maximum growth temperature, strains were initially cultivated at 25 °C for two days, and then the temperature was gradually increased by a gradient of 1 °C until the colony stopped growing.

### DNA extraction, amplification and sequencing

Cultures were grown at 27 °C for one week on PDA plates. For Sanger sequencing and whole genomic sequencing (WGS), cell total DNAs were extracted from the fungal thalli following the instruction of the GO-GPLF-400 kit (GeneOnBio Corporation, Changchun, China). The internal transcribed spacer of the ribosomal DNA (ITS rDNA) was amplified with the primer pairs LR5M (5′-GCT ATC CTG AGG GAA ACT TCG-3′) and NS5M (5′-GGC TTA ATT TGA CTC AAC ACG G-3′; Zhao *et al*. 2021b). The polymerase chain reaction (PCR) program was performed as follows: an initial temperature at 95 °C for 5 min; then 30 cycles of denaturation at 95 °C for 1 min, annealing at 55 °C for 45 s and extension at 72 °C for 1 min; and a final extra extension at 72 °C for 10 min. The PCR products were sequenced with the primers ITS4 and ITS5 (White *et al*. 1990). Sanger sequencing of the PCR products was conducted by BGI Tech Solutions Beijing Liuhe Co., Limited (https://www.bgi.com/, Beijing, China). For WGS, total DNAs were applied to the HiSeq 4000 platform (https://www.illumina.com, Illumina Inc., California, USA) with Novogene Co., Ltd. (https://novogene.com/, Beijing, China).

### Phylogenomic and phylogenetic analyses

De novo assembly, manual proofreading and target extraction for ITS rDNA sequences were carried out with Geneious 9.0.2 (http://www.geneious.com, Kearse *et al*. 2012). Assembled sequences were then submitted to GenBank under accession numbers in Table 1. The ITS rDNA dataset containing sequences retrieved from GenBank was realigned locally with AliView 3.0 (Larsson 2014) and MAFFT 7 (Katoh *et al*. 2019), and then was manually proofread. The annotated genome sequences were then submitted to GenBank under accession numbers in Table 2. The whole proteome was obtained from the annotated genome with Gffread v0.12.4 (Pertea & Pertea 2020), and 192 clusters of orthologous proteins (COPs) were retrieved with HMMER 3.3.1 (http://hmmer.org/download.html) and Trimal 1.4.4 (Sanchez *et al*. 2011) according to the method by Spatafora *et al*. (2016). Phylogenetically informative protein markers followed James *et al*. (2013). Phylogenomic and phylogenetic analyses were carried out following the methods by Nie *et al*. (2020a, b), using the PROTGAMMALGX model with 100 bootstrap replications. Phylogenetic analyses were carried out following the methods by Nie *et al*. (2020a, b), including Maximum Likelihood (ML) and Bayesian Inference (BI) implemented in RAxML version 8 (Stamatakis 2014) and MrBayes 3.2.7a (Ronquist *et al*. 2012), respectively. Maximum Likelihood analysis was performed using the GTRGAMMA model with 1,000 bootstrap replications. For BI analysis, eight cold Markov chains ran simultaneously for two million generations with the GTR + I + G model, sampling every 1,000 generations and removing the first 25% sampled trees as burn-in. In the BI analyses, Markov Chain Monte Carlo (MCMC) chains ran until the convergence met and the standard deviation fell below 0.01. The phylogram was viewed and modified with FigTree version 1.4.4 (http://tree.bio.ed.ac.uk/software/figtree/).

**Table 1.** Information of new species and Chinese new record species reported in this study. * New species proposed herein are in bold, new combinations herein are underlined. ** T represents ex-holotype.

| Species* | Strains** | Substrate | Locality | GenBank<br>accession Nos.<br>of ITS rDNA |
| --- | --- | --- | --- | --- |
| *Absidia alpina* | CGMCC 3.16104<sup>T</sup> | soil | Yunnan, China | OL678133 |
| *A. ampullacea* | CGMCC 3.16054<sup>T</sup> | soil | Beijing, China | MZ354138 |
| *A. biappendiculata* | NRRL 3033 | leaves | Wyoming, USA | MZ354153 |
| *A. brunnea* | CGMCC 3.16055<sup>T</sup> | soil | Qinghai, China | MZ354139 |
| *A. caerulea* | XY00608 | flower | Guangdong, China | OL620081 |
|  | XY00729 | flower | Guangdong, China | OL620082 |
| *A. chinensis* | CGMCC 3.16056<sup>T</sup> | soil | Yunnan, China | MZ354140 |
|  | CGMCC 3.16057 | soil | Sichuan, China | MZ354141 |
| *A. cinerea* | CGMCC 3.16062<sup>T</sup> | soil | Beijing, China | MZ354146 |
| *A. digitula* | CGMCC 3.16058<sup>T</sup> | soil | Xinjiang, China | MZ354142 |
| *A. jiangxiensis* | CGMCC 3.16105<sup>T</sup> | soil | Jiangxi, China | OL678134 |
| *A. koreana* | XY00816 | soil | Hubei, China | OL620083 |
|  | XY00596 | seed | Hubei, China | OL620084 |
| *A. nigra* | NRRL 3060<sup>T</sup> | soil | Wisconsin, USA | MZ354143 |
|  | CGMCC 3.16059 | soil | Jilin, China | MZ354144 |
|  | CGMCC 3.16060 | soil | Inner Mongolia, China | MZ354152 |
| *A. oblongispora* | CGMCC 3.16061<sup>T</sup> | soil | Yunnan, China | MZ354145 |
| *A. pararepens* | XY00631 | soil | Beijing, China | OL620085 |
|  | XY00615 | soil | Beijing, China | OL620086 |
|  | XY05899 | soil | Tibet, China | OL620087 |
| *A. purpurea* | CGMCC 3.16106<sup>T</sup> | soil | Yunnan, China | OL678135 |
| *A. sympodialis* | CGMCC 3.16064<sup>T</sup> | soil | Shaanxi, China | MZ354148 |
|  | CGMCC 3.16063<sup>T</sup> | soil | Beijing, China | MZ354147 |
| *A. xinjiangensis* | CGMCC 3.16107<sup>T</sup> | soil | Xinjiang, China | OL678136 |
| *A. varians* | CGMCC 3.16065<sup>T</sup> | soil | Yunnan, China | MZ354149 |
| *A. virescens* | CGMCC 3.16067<sup>T</sup> | soil | Yunnan, China | MZ354151 |
|  | CGMCC 3.16066<sup>T</sup> | soil | Yunnan, China | MZ354150 |
| *Actinomortierella* | XY06923 | soil | Yunnan, China | OL620101 |
| *ambigua* |  |  |  |  |
| *Backusella* | CGMCC 3.16108<sup>T</sup> | soil | Yunnan, China | OL678137 |
| *dichotoma* |  |  |  |  |
|  | XY07504 | soil | Yunnan, China | OL678138 |
| *B. dispersa* | XY08806 | soil | Xinjiang, China | OL620088 |
| *B. locustae* | XY07940 | soil | Hunan, China | OL620089 |
| *B. moniliformis* | CGMCC 3.16109<sup>T</sup> | soil | Shanxi, China | OL678139 |
| *B. ovalispora* | CGMCC 3.16110<sup>T</sup> | soil | Guangxi, China | OL678140 |
|  | XY07481 | soil | Guangxi, China | OL678141 |
| *B. oblongielliptica* | XY08152 | soil | Yunnan, China | OL620090 |
|  | XY08767 | soil | Zhejiang, China | OL620091 |
|  | XY08768 | soil | Zhejiang, China | OL620092 |
| *B. tuberculispora* | XY07548 | soil | Yunnan, China | OL620093 |
|  | XY7557 | soil | Yunnan, China | OL620094 |
| *Circinella* | CGMCC 3.16026<sup>T</sup> | soil | Hainan, China | MW580599 |
| *homothallica* |  |  |  |  |
| *Cunninghamella* | XY07003 | soil | Anhui, China | OL620095 |
| *antarctica* |  |  |  |  |
|  | XY07050 | soil | Gansu, China | OL620096 |
| *C. arrhiza* | CGMCC 3.16111<sup>T</sup> | soil | Hunan, China | OL678142 |
|  | XY08047 | soil | Beijing, China | OL678143 |
| *C. guttata* | CGMCC 3.16112<sup>T</sup> | soil | Beijing, China | OL678144 |
| *C. irregularis* | CGMCC 3.16113<sup>T</sup> | soil | Inner Mongolia, China | OL678144 |
|  | XY07683 | soil | Inner Mongolia, China | OL678146 |
|  | XY07657 | soil | Inner Mongolia, China | OL678147 |
| *C. nodosa* | AS. 3.5628<sup>T</sup> | flower | Beijing, China | AF346407 |
| *C. regularis* | CGMCC 3.16114<sup>T</sup> | soil | Yunnan, China | OL678148 |
|  | XY07510 | soil | Yunnan, China | OL678149 |
|  | XY07512 | soil | Yunnan, China | OL678150 |
|  | XY07519 | soil | Yunnan, China | OL678151 |
| *C. subclavata* | CGMCC 3.16115<sup>T</sup> | soil | Yunnan, China | OL678152 |
|  | XY07766 | soil | Yunnan, China | OL678153 |
| *C. varians* | CGMCC 3.16116<sup>T</sup> | soil | Jiangsu, China | OL678154 |
|  | XY06999 | soil | Jiangsu, China | OL678155 |
| *Dissophora* | XY08870 | soil | Xinjiang, China | OL620097 |
| *globulifera* |  |  |  |  |
| *Gamsiella* | CGMCC 3.16117<sup>T</sup> | soil | Tibet, China | OL678156 |
| *globistyluspora* |  |  |  |  |
| *Gongronella* | CGMCC 3.16118<sup>T</sup> | soil | Jiangxi, China | OL678157 |
| *chlamydospora* |  |  |  |  |
| *G. multispora* | CGMCC 3.16119<sup>T</sup> | soil | Beijing, China | OL678158 |
| *G. namwonensis* | XY08131 | soil | Zhejiang, China | OL620098 |
| *Isomucor trufemiae* | XY07543 | soil | Yunnan, China | OL620099 |
|  | XY07544 | soil | Yunnan, China | OL620100 |
| *Lichtheimia alba* | CGMCC 3.16120<sup>T</sup> | soil | Beijing, China | OL678159 |
| *L. globospora* | CGMCC 3.16025<sup>T</sup> | soil | Hainan, China | MW580598 |
| *Modicella abundans* | CGMCC 3.16121<sup>T</sup> | soil | Anhui, China | OL678160 |
| *Mortierella* | CGMCC 3.16122<sup>T</sup> | soil | Yunnan, China | OL678161 |
| *amphispora* |  |  |  |  |
|  | XY08653 | soil | Yunnan, China | OL678162 |
|  | XY08796 | soil | Yunnan, China | OL678292 |
| *M. annulata* | CGMCC 3.16123<sup>T</sup> | soil | Yunnan, China | OL678163 |
| *M. bainieri* | XY08661 | soil | Yunnan, China | OL620102 |
| *M. beljakovae* | XY03433 | soil | Heilongjiang, China | OL620103 |
| *M. chlamydospora* | XY07448 | soil | Zhejiang, China | OL620104 |
|  | XY08109 | soil | Guangxi, China | OL620105 |
| *M. cylindrispora* | CGMCC 3.16124<sup>T</sup> | soil | Jiangsu, China | OL678164 |
| *M. cystojenkinii* | XY03898 | soil | Inner Mongolia, China | OL620106 |
|  | XY03876 | plant debris | Heilongjiang, China | OL620107 |
|  | XY03882 | plant debris | Heilongjiang, China | OL620108 |
| *M. epicladia* | XY06924 | soil | Yunnan, China | OL620109 |
|  | XY06927 | soil | Yunnan, China | OL620110 |
| *M. floccosa* | CGMCC 3.16125<sup>T</sup> | soil | Jiangsu, China | OL678165 |
| *M. fluviae* | XY04370 | soil | Heilongjiang, China | OL620111 |
| *M. fusiformispora* | CGMCC 3.16029<sup>T</sup> | soil | Fields Peninsula, Antarctica | MW580602 |
|  | CGMCC 3.16030 | soil | Fields Peninsula, Antarctica | MW580603 |
| *M. lignicola* | XY08224 | soil | Yunnan, China | OL620112 |
| *M. lobata* | CGMCC 3.16126<sup>T</sup> | soil | Beijing, China | OL678166 |
| *M. longigemmata* | XY07567 | soil | Yunnan, China | OL620113 |
|  | XY08599 | soil | Yunnan, China | OL620114 |
|  | XY08601 | soil | Yunnan, China | OL620115 |
| *M. macrochlamydospora* | CGMCC 3.16127<sup>T</sup> | plant | Inner Mongolia, China | OL678167 |
|  |  | debris |  |  |
|  | XY04275 | plant | Inner Mongolia, China | OL678168 |
|  |  | debris |  |  |
| *M. microchlamydospora* | CGMCC 3.16128<sup>T</sup> | soil | Yunnan, China | OL678169 |
|  | XY07258 | soil | Beijing, China | OL678170 |
|  | XY07283 | soil | Beijing, China | OL678171 |
| *M. mongolica* | CGMCC 3.16129<sup>T</sup> | soil | Inner Mongolia, China | OL678172 |
|  | XY03534 | soil | Inner Mongolia, China | OL678173 |
| *M. nantahalensis* | XY07486 | soil | Guangxi, China | OL620116 |
|  | XY06017 | soil | Hainan, China | OL620117 |
| *M. rishiksha* | XY07315 | soil | Beijing, China | OL620118 |
| *M. schmuckeri* | XY07412 | soil | Beijing, China | OL620119 |
|  | XY07419 | soil | Beijing, China | OL620120 |
| *M. sclerotiella* | XY03528 | soil | Inner Mongolia, China | OL620121 |
| *M. seriatoinflata* | CGMCC 3.16130<sup>T</sup> | plant | Beijing, China | OL678174 |
|  |  | debris |  |  |
|  | XY06985 | soil | Anhui, China | OL678175 |
|  | CBS 314.52 | soil | Germany | MH868591 |
| *M. sparsa* | CGMCC 3.16131<sup>T</sup> | leaves | Heilongjiang, China | OL678176 |
|  | XY03860 | leaves | Heilongjiang, China | OL678291 |
|  | XY04406 | leaves | Inner Mongolia, China | OL678177 |
| *M. turficola* | XY03346 | plant debris | Heilongjiang, China | OL620122 |
| *M. varians* | CGMCC 3.16132<sup>T</sup> | soil | Tibet, China | OL678178 |
|  | XY05920 | soil | Tibet, China | OL678179 |
| *Mucor* | CGMCC 3.16133<sup>T</sup> | soil | Guangxi, China | OL678180 |
| *abortisporangium* |  |  |  |  |
| *M. aligarensis* | XY07583 | soil | Yunnan, China | OL620123 |
| *M. amphisorus* | CGMCC 3.16134<sup>T</sup> | soil | Inner Mongolia, China | OL678181 |
|  | XY07649 | soil | Inner Mongolia, China | OL678182 |
| *M. atramentarius* | XY06751 | soil | Tibet, China | OL620124 |
| *M. bainieri* | XY08747 | soil | Yunnan, China | OL620125 |
| *M. breviphorus* | CGMCC 3.16135<sup>T</sup> | soil | Yunnan, China | OL678183 |
| *M. brunneolus* | CGMCC 3.16136<sup>T</sup> | soil | Guangxi, China | OL678184 |
| *M. changshaensis* | CGMCC 3.16137<sup>T</sup> | soil | Hunan, China | OL678185 |
|  | XY07965 | soil | Hunan, China | OL678186 |
| *M. chlamydosporus* | CGMCC 3.16138<sup>T</sup> | soil | Hebei, China | OL678187 |
|  | XY08211 | soil | Yunnan, China | OL678188 |
|  | XY08225 | soil | Yunnan, China | OL678189 |
| *M. donglingensis* | CGMCC 3.16139<sup>T</sup> | plant | Beijing, China | OL678190 |
|  |  | debris |  |  |
|  | XY08501 | soil | Yunnan, China | OL678191 |
| *M. floccosus* | CGMCC 3.16140<sup>T</sup> | soil | Hunan, China | OL678192 |
|  | XY07967 | soil | Hunan, China | OL678193 |
| *M. fluvii* | XY07446 | soil | Zhejiang, China | OL620126 |
|  | XY07447 | soil | Zhejiang, China | OL620127 |
| *M. fuscus* | XY06718 | soil | Tibet, China | OL620128 |
|  | XY06719 | soil | Tibet, China | OL620129 |
| *M. fusiformisporus* | CGMCC 3.16141<sup>T</sup> | soil | Jiangsu, China | OL678194 |
|  | XY08153 | soil | Yunnan, China | OL678195 |
|  | XY08154 | soil | Yunnan, China | OL678196 |
|  | XY08117 | soil | Zhejiang, China | OL678197 |
| *M. griseocyanus* | XY06685 | soil | Tibet, China | OL620130 |
|  | XY06697 | soil | Tibet, China | OL620131 |
| *M. heilongjiangensis* | CGMCC 3.16142<sup>T</sup> | soil | Heilongjiang, China | OL678198 |
|  | XY05057 | soil | Heilongjiang, China | OL678199 |
| *M. hemisphaericum* | CGMCC 3.16143<sup>T</sup> | soil | Tibet, China | OL678200 |
| *M. homothallicus* | CGMCC 3.16144<sup>T</sup> | soil | Jiangsu, China | OL678201 |
|  | XY06967 | soil | Jiangsu, China | OL678202 |
| *M. hyalinosporus* | CGMCC 3.16145<sup>T</sup> | soil | Beijing, China | OL678203 |
| *M. lobatus* | CGMCC 3.16146<sup>T</sup> | soil | Jiangsu, China | OL678204 |
|  | XY07029 | soil | Hubei, China | OL678205 |
| *M. lusitanicus* | XY06042 | soil | Shandong, China | OL620132 |
|  | XY06074 | soil | Shandong, China | OL620133 |
|  | XY07298 | dung | Beijing, China | OL620134 |
|  | XY07299 | dung | Beijing, China | OL620135 |
| *M. minutus* | XY06939 | soil | Yunnan, China | OL620136 |
|  | XY06940 | soil | Yunnan, China | OL620137 |
|  | XY07460 | soil | Hubei, China | OL620138 |
| *M. moniliformis* | CGMCC 3.16147<sup>T</sup> | soil | Zhejiang, China | OL678206 |
|  | XY07458 | soil | Hubei, China | OL678207 |
| *M. mucedo* | XY07301 | soil | Beijing, China | OL620139 |
|  | XY07302 | soil | Beijing, China | OL620140 |
| *M. nanus* | XY07037 | soil | Hubei, China | OL620141 |
|  | XY07221 | soil | Beijing, China | OL620142 |
| *M. nidicola* | XY08161 | soil | Yunnan, China | OL620143 |
| *M. orientalis* | CGMCC 3.16148<sup>T</sup> | soil | Anhui, China | OL678208 |
| *M. pseudocircinelloides* | XY07713 | soil | Shanxi, China | OL620144 |
|  | XY07714 | soil | Shanxi, China | OL620145 |
| *M. radiatus* | CGMCC 3.16149<sup>T</sup> | soil | Yunnan, China | OL678209 |
|  | XY08167 | soil | Yunnan, China | OL678210 |
| *M. rhizosporus* | CGMCC 3.16150<sup>T</sup> | soil | Guangxi, China | OL678211 |
| *M. robustus* | CGMCC 3.16151<sup>T</sup> | soil | Xinjiang, China | OL678212 |
|  | **XY08976** | **soil** | **Xinjiang, China** | **OL678213** |
|  | **XY09024** | **soil** | **Xinjiang, China** | **OL678214** |
| *M. silvaticus* | XY04328 | plant debris | Heilongjiang, China | OL620146 |
|  | XY04329 | plant debris | Heilongjiang, China | OL620147 |
| *M. sino-saturninus* | CGMCC 3.16152<sup>T</sup> | soil | Yunnan, China | OL678215 |
|  | XY08556 | soil | Yunnan, China | OL678216 |
| *M. stercorearius* | XY07007 | soil | Anhui, China | OL620148 |
| *M. variicolumellatus* | XY07369 | rotten wood | Beijing, China | OL620149 |
|  | XY07370 | rotten wood | Beijing, China | OL620150 |
| *Syncephalastrum* | **CGMCC 3.16153<sup>T</sup>** | **paper** | **Fujian, China** | **OL678217** |
| *breviphorum* |  |  |  |  |
|  | **XY06838** | **paper** | **Fujian, China** | **OL678218** |
| *S. elongatum* | **CGMCC 3.16154<sup>T</sup>** | **paper** | **Fujian, China** | **OL678219** |
| *S. simplex* | **CGMCC 3.16155<sup>T</sup>** | **soil** | **Hubei, China** | **OL678220** |
| *S. sympodiale* | **CGMCC 3.16156<sup>T</sup>** | **soil** | **Beijing, China** | **OL678221** |
|  | **XY06803** | **soil** | **Beijing, China** | **OL678222** |
|  | **XY06811** | **mushroom** | **Beijing, China** | **OL678223** |
| *Umbelopsis brunnea* | **CGMCC 3.16023<sup>T</sup>** | **soil** | **Heilongjiang, China** | **MV580596** |
|  | **CGMCC 3.16024** | **soil** | **Inner Mongolia, China** | **MV580597** |
| *U. chlamydospora* | **CGMCC 3.16157<sup>T</sup>** | **soil** | **Beijing, China** | **OL678224** |
| *U. crustacea* | **CGMCC 3.16158<sup>T</sup>** | **soil** | **Inner Mongolia, China** | **OL678225** |
|  | **XY03542** | **soil** | **Inner Mongolia, China** | **OL678226** |
|  | **XY03944** | **soil** | **Heilongjiang, China** | **OL678227** |
| *U. gibberispora* | XY06667 | soil | Zhejiang, China | OL620151 |
| *U. globospora* | **CGMCC 3.16159<sup>T</sup>** | **soil** | **Zhejiang, China** | **OL678228** |
| *U. gutianensis* | **CGMCC 3.16160<sup>T</sup>** | **soil** | **Zhejiang, China** | **OL678229** |
|  | **XY06604** | **soil** | **Zhejiang, China** | **OL678230** |
|  | **XY06605** | **soil** | **Zhejiang, China** | **OL678231** |
| *U. oblongispora* | **CGMCC 3.16161<sup>T</sup>** | **soil** | **Zhejiang, China** | **OL678232** |
| *U. ovata* | XY03175 | soil | Heilongjiang, China | OL620152 |
|  | XY03176 | soil | Heilongjiang, China | OL620153 |
| ***U. septata*** | **CGMCC 3.16162<sup>T</sup>** | **soil** | **Zhejiang, China** | **OL678233** |
| *U. sinsidoensis* | XY06194 | soil | Zhejiang, China | OL620154 |
| ***U. tibetica*** | **CGMCC 3.16163<sup>T</sup>** | **soil** | **Tibet, China** | **OL678234** |
|  | **XY07881** | **soil** | **Tibet, China** | **OL678235** |
|  | **XY07882** | **soil** | **Tibet, China** | **OL678236** |

**Table 2.** The genomic information used for molecular dating in this study. * SRA NCBI stands for Sequence Read Archive, National Center for Biotechnology Information, US National Library of Medicine.

| Species | Strains | BioSample | SRA NCBI* | Reference |
| --- | --- | --- | --- | --- |
| *Absidia glauca* | CBS 101.48 | SAMEA3923633 | ERS1110767 | (Ellenberger *et al.* 2016) |
| *Acaulospora morrowiae* | CL551 | SAMEA8911292 | ERS6594176 |  |
| *Ambispora gerdemannii* | MT106 | SAMEA8911308 | ERS6594192 |  |
| *A. leptoticha* | FL130A | SAMEA8911307 | ERS6594191 |  |
| *Apophysomyces elegans* | B7760 | SAMN02351510 | SRS478092 |  |
| *A. trapeziformis* | B9324 | SAMN02351504 | SRS478090 |  |
| *Backusella circina* | CGMCC3.15908 | SAMN28403603 |  | This study |
| *B. lamprospora* | CGMCC3.15922 | SAMN28403604 |  | This study |
| *Cetraspora pellucida* | FL966 | SAMEA8911298 | ERS6594182 |  |
| *Choanephora cucurbitarum* | KUS-F282377 | SAMN04532838 |  |  |
| *Claroideoglossus candidum* | CCK_pot_B_6-9 | SAMEA8911304 | ERS6594188 |  |
| *Cokeromyces recurvatus* | B5483 | SAMN02370952 | SRS489494 |  |
| *Cunninghamella elegans* | B9769 | SAMN02351511 | SRS478097 |  |
| *Dentiscutata erythropus* | MA453B | SAMEA8911297 | ERS6594181 |  |
| *Dissophora decumbens* | CBS 301.87 | SAMN23435287 |  | This study |
| *D. globulifera* | CBS 858.70 | SAMN23461011 |  | This study |
| *Diversispora eburnea* | AZ414A | SAMEA8911293 | ERS6594177 |  |
| *Endogone sp.* | FLAS-F59071 | SAMN09071421 | SRS5707057 | (Chang *et al.* 2019) |
| *Entomortierella lignicola* | CBS 207.37 | SAMN23441792 |  | This study |
| *E. parvispora* | CBS 304.52 | SAMN23437757 |  | This study |
| *Funneliformis caledonium* | UK204 | SAMEA8911289 | ERS6594173 |  |
| *F. mosseae* | 87-<br>6_pot_B_2015 | SAMEA8911288 | ERS6594172 |  |
| *Gamsiella multivaricata* | CBS 227.78 | SAMN23437755 |  | This study |
| *Geosiphon pyriformis* | V1.0 |  |  | (Malar *et al.* 2021,<br>2022) |
| *Gigaspora margarita* | BEG34 | SAMEA6235760 | ERS4035453 |  |
| *G. rosea* | DAOM194757 |  |  | (Morin *et al.* 2019) |
| *Hesseltinella vesiculosa* | NRRL 3301 | SAMN02745228 | SRS710783 | (Mondo *et al.* 2017) |
| *Jimgerdemannia*<br>*flammicorona* | AD002 | SAMN09071422 | SRS5707054 | (Chang *et al.* 2019) |
| *J. flammicorona* | GMNB39 | SAMN05444478 |  | (Chang *et al.* 2019) |
| *J. lactiflua* | OSC166217 | SAMN10169340 |  | (Chang *et al.* 2019) |
| *Lichtheimia corymbifera* | FSU 9682 | SAMEA2189700 | ERS339162 | (Schwartz *et al.* 2014) |
| *L. ramosa* | FSU 6197 | SAMN05179542 | SRS1468292 | (Linde *et al.* 2014) |
| *Linnemannia amoeboides* | CBS 889.72 | SAMN19911466 | SRS9293840 | (Yang *et al.* 2022) |
| *Mortierella indohii* | CBS 720.71 | SAMN23441793 |  | This study |
| *M. microzygospora* | CBS 880.97 | SAMN23441791 |  | This study |
| *M. parazychnae* | CBS 868.71 | SAMN23441790 |  | This study |
| *M. rishikeshae* | CBS 652.68 | SAMN23437756 |  | This study |
| *Mucor circinelloides* | 1006PhL | SAMN00103456 | SRS074119 | (Lee *et al.* 2014) |
| *M. lusitanicus* | CBS 277.49 | SAMN00120579 |  | (Corrochano *et al.* 2016) |
| *M. saturninus* |  | SAMN16393843 |  |  |
| *Paraglomus brasilianum* | BR232B | SAMEA8911306 | ERS6594190 |  |
| *P. occultum* | IA702 | SAMEA8911305 | ERS6594189 |  |
| *Parasitella parasitica* | CBS 412.66 | SAMEA278055 | ERS550304 | (Ellenberger *et al.* 2016) |
| *Podila clonocystis* | CBS 357.76 | SAMN23461012 |  | This study |
| *Racocetra persica* | MA461A | SAMEA8911300 | ERS6594184 |  |
| *Rhizophagus cerebriforme* | DAOM 227022 |  |  | (Morin *et al.* 2019) |
| *R. clarus* | RCL21 | SAMD00190756 |  |  |
| *R. irregularis* | DAOM 197198 |  |  | (Morin *et al.* 2019) |
| *Rhizopus arrhizus* | GL8 | SAMN14162349 |  | (Nguyen *et al.* 2020) |
| *R. arrhizus* | GL4 | SAMN14162309 |  | (Nguyen *et al.* 2020) |
| *R. microsporus* | ATCC 52813 | SAMN06821222 | SRS2140116 | (Horn *et al.* 2015) |
| *Phycomyces blakesleeana* | NRRL 1555 | SAMN00189023 |  | (Corrochano *et al.* 2016) |
| *Smittium culicis* | GSMNP | SAMN04489870 |  | (Wang *et al.* 2016) |
| *Syncephalastrum monosporum* | B8922 | SAMN02370995 | SRS489641 |  |
| *Thermomucor indicae-seudaticae* | HACC 243 | SAMN03070115 | SRS702596 |  |
| *Umbelopsis isabellina* | WA0000067209 | SAMN16393839 |  | (Takeda *et al.* 2014) |
| *U. vinacea* | WA0000051536 | SAMN16393840 | SRS7563300 |  |

### Molecular dating

A set of ITS and LSU rDNA sequences retrieved from GenBank database, was used to infer the divergence time within the Mucoromyceta (Table 2). The divergence time was estimated with BEAST 2.6.5 (Bouckaert *et al*. 2014). A BEAST XML input file was generated with BEAUti v2 implemented in the BEAST 2.6.5. The estimation of rates of evolutionary changes at amino acid sites was performed by ModelTest 3.7 (Posada & Crandall 1998) using the GTR substitution model. The relaxed clock model with a log-normal distribution was used. A Yule model was selected as prior, assuming a constant speciation rate with no extinction (Barraclough & Nee 2001) and an offset age with a gamma distributed prior (scale = 10, shape = 1.0). The divergence time analysis was calibrated using two fossil fungi and three molecular divergence estimates. The two fossil fungi were 1) *Protoascon missouriensis* L.R. Batra *et al*. in the Middle Pennsylvanian sub-epoch (approximately 315‒307 Mya) which is remarkably similar to *Absidia spinosa* Lendn. or *A. spinosa* var. *azygospora* Boedijn in azygosporangia and appendages (Boedijn 1958, Batra *et al*. 1964, Baxter 1975, Taylor & Alexander 2005, Taylor *et al*. 2014), and 2) *Jimwhitea circumtecta* M. Krings & T.N. Taylor which is the most persuasive fossil representative of the *Endogone* and was discovered from the Middle Triassic epoch (roughly 247‒237 Mya; Krings *et al*. 2011, Taylor *et al*. 2014). The three molecular divergence estimates were as follows: 1) Although some molecular studies suggested that zygomycetous fungi appeared at the Precambrian super eon (1.4‒1.2 Gya), the conservative estimate of 800 Mya or so during the Proterozoic eon was adopted herein (Berbee & Taylor 2001, Heckman *et al*. 2001, Blair 2009); 2) The crown age of Mucoromycota was set to 560 Mya with 95 % HPD of 183–735 Mya (Tedersoo *et al*. 2018). 3) Glomeromycota was set to differentiate 642 Mya with 95 % HPD of 597–720 Mya (Tedersoo *et al*. 2018). The analysis ran for 200 million generations, logging states every 1,000 generations. Log files were mixed with Tracer v1.7.2 (Rambaut & Drummond 2013) and checked for convergence. Finally, a maximum clade credibility (MCC) tree was summarized with TreeAnnotator 1.8.2 (within BEAST), removing the first 20 % of states as burn-in, and annotating clades with more than 0.8 posterior probability.

## Results

### Phylogenomics, phylogenetics and molecular dating in Mucoromyceta

We sequenced the genomes of 12 strains of 12 species in 6 genera for phylogenomic analyses (Table 2). We also downloaded the published genomes of 44 strains of 42 species in 29 genera including an outgroup from *Smittium culicis* Tuzet & Manier ex Kobayasi in the phylum Zoopagomycota, and all of these retrieved genomes are freely available for public use in accordance with the usage policy in the GenBank database and the Joint Genome Institute (JGI) portal MycoCosm database (Table 2). Unfortunately, genomes on Calcarisporiellomycota are unavailable. Phylogenetic analyses and molecular dating were conducted based on ITS and LSU rDNA sequences of 170 strains of 165 species in 98 genera (accounting for 81 %, 98 / 121) in the Mucoromyceta and that of an outgroup in *Neurospora crassa* Shear & B.O. Dodge. The most optimal model of BI was GTR + I + R, and the average standard deviation of split frequencies was 0.009026. Topology of ML tree was selected to represent the phylogenetic relationship of Mucoromyceta (Fig. 3). The phylogenomic (Fig. 2) and phylogenetic analyses (Fig. 3) suggest an identical taxonomic backbone of the subkingdom Mucoromyceta: four accepted phyla (Calcarisporiellomycota, Glomeromycota, Mortierellomycota and Mucoromycota) and two new phyla (Endogonomycota *phyl. nov*. and Umbelopsidomycota *phyl. nov*.). These results also provide a strong phylogenetic resolution for the phylum, class, order and family in the subkingdom Mucoromyceta, proposing five new orders (Claroideoglomerales *ord. nov.*, Cunninghamellales *ord. nov.*, Lentamycetales *ord. nov.*, Phycomycetales *ord. nov.* and Syncephalastrales *ord. nov.*) and six new families (Circinellaceae *fam. nov.*, Gongronellaceae *fam. nov.*, Protomycocladaceae *fam. nov.*, Rhizomucoraceae *fam. nov.*, Syzygitaceae *fam. nov.* and Thermomucoraceae *fam. nov.*).

**Fig. 2.**
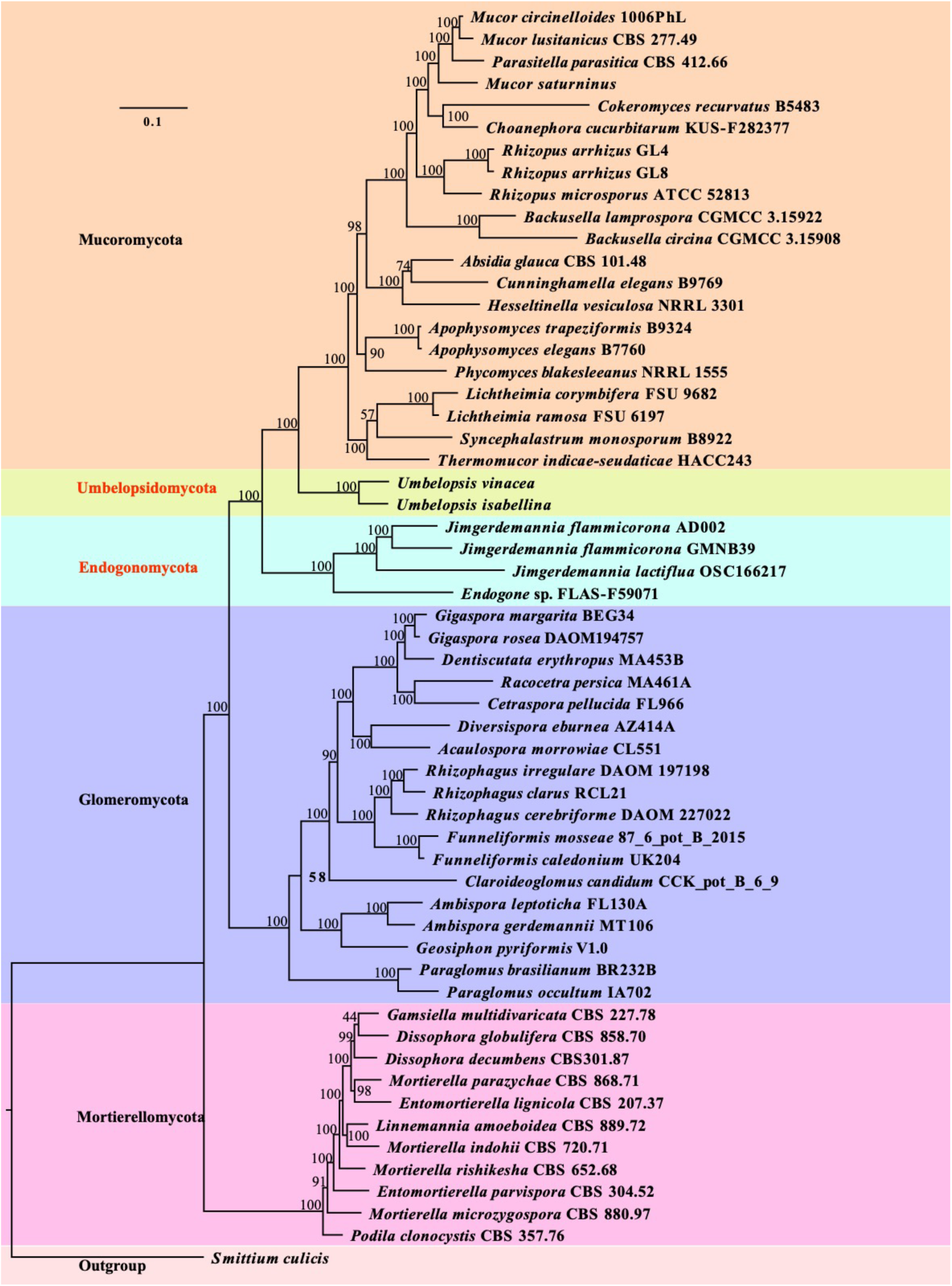
A Maximum Likelihood phylogenomic tree illustrating relationships within subkingdom Mucoromyceta based on 192 clusters of orthologous proteins. New taxa proposed in the present study are in red. Maximum Likelihood bootstrap values (≥50%) are indicated along branches. Phyla are highlighted in different colors. A scale bar in the upper left indicates substitutions per site.

**Fig. 3.**
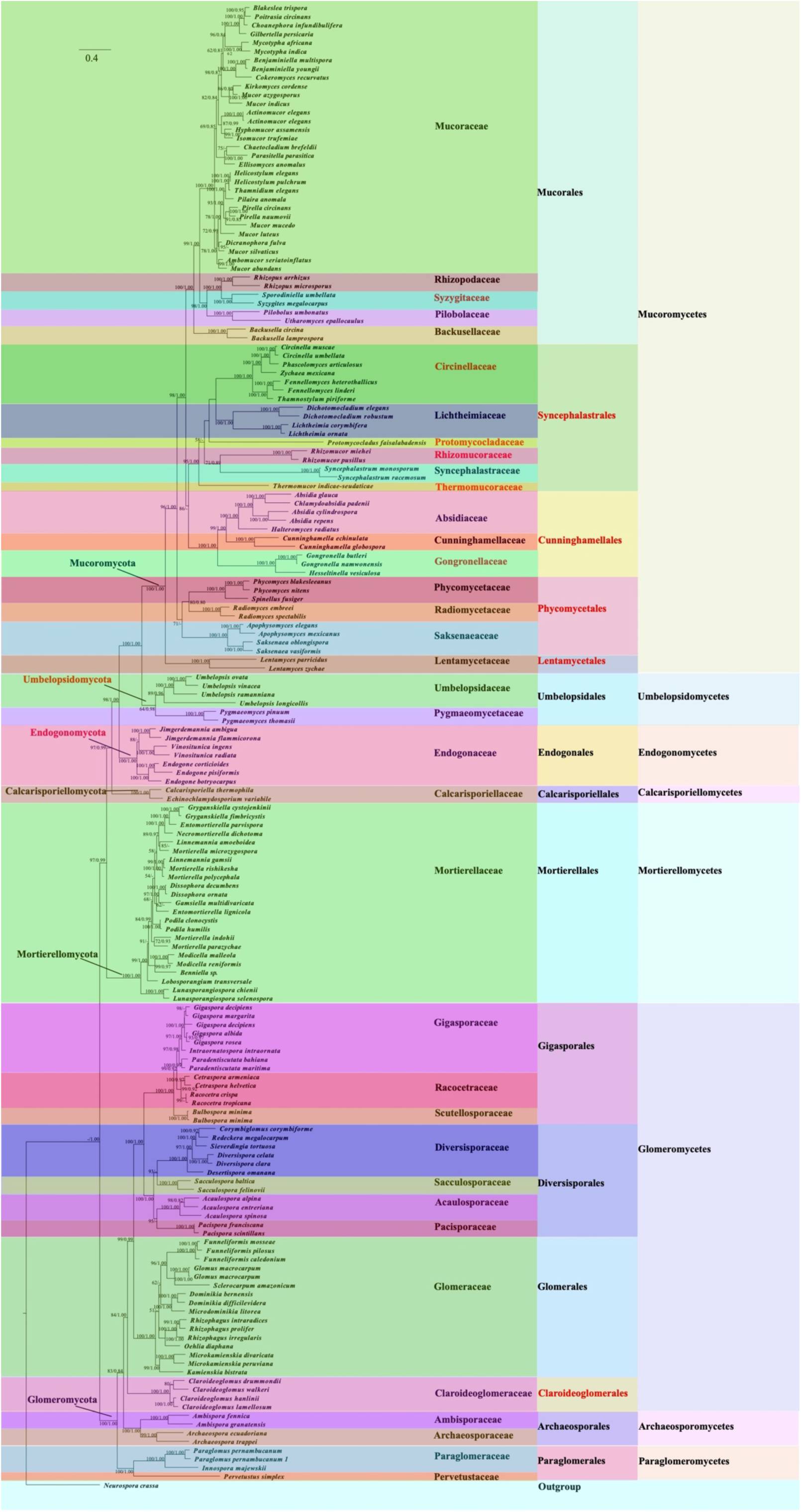
A Maximum Likelihood phylogenetic tree illustrating relationships within subkingdom Mucoromyceta based on ITS and LSU rDNA sequences, with *Neurospora crassa* as outgroup. New taxa proposed in the present study are in red. Maximum Likelihood bootstrap values (≥50%) / Bayesian Inference Posterior Probabilities (≥0.80) are indicated along branches. Taxa are highlighted in different colors. A scale bar in the upper left indicates substitutions per site.

The taxonomic hierarchy above is dated with estimated divergence times: phyla 810–639 Mya, classes 651–585 Mya, orders 570–400 Mya, and families 488–107 Mya (Fig. 4 and Table 4). The six phylum clades are supported with high posterior probabilities (PP): Glomeromycota (810 Mya / PP = 1.0), Mortierellomycota (777 Mya / PP = 0.9), Calcarisporiellomycota (748 Mya / PP = 1.0), Endogonomycota *phyl. nov*. (708 Mya / PP = 1.0), Mucoromycota (639 Mya / PP = 1.0), and Umbelopsidomycota *phyl. nov*. (639 Mya / PP = 1.0).

**Fig. 4.**
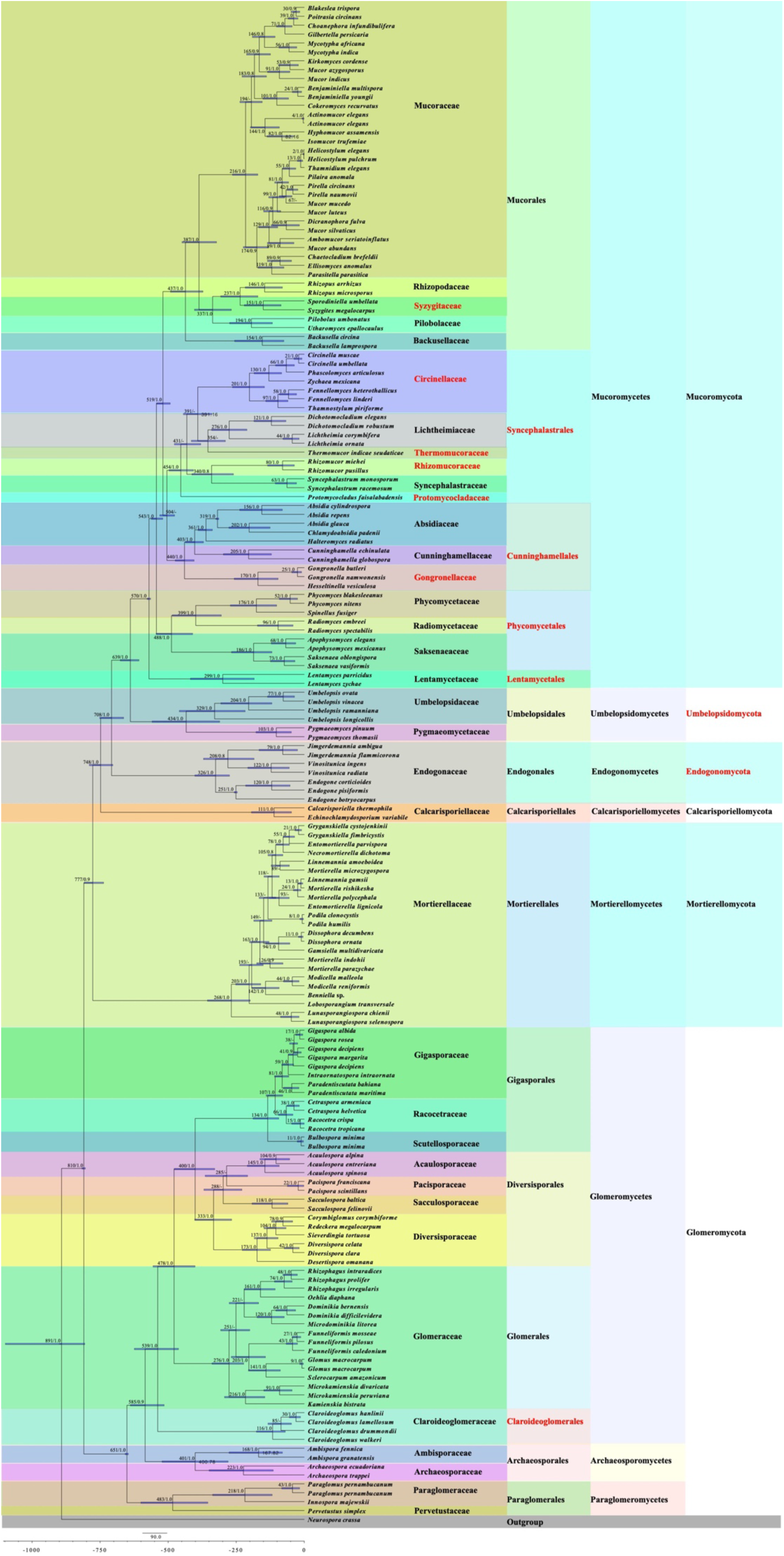
Time-scaled Bayesian maximum clade credibility phylogenomic tree inferred from ITS and LSU rDNA sequences for the subkingdom Mucoromyceta, with *Neurospora crassa* as outgroup Estimated mean divergence time (Mya) and posterior probabilities (PP) > 0.8 are annotated at the internodes. The 95 % highest posterior density (HPD) interval of divergence time estimates are marked by horizontal blue bars. New taxa proposed in the present study are in red.

**Table 3.**
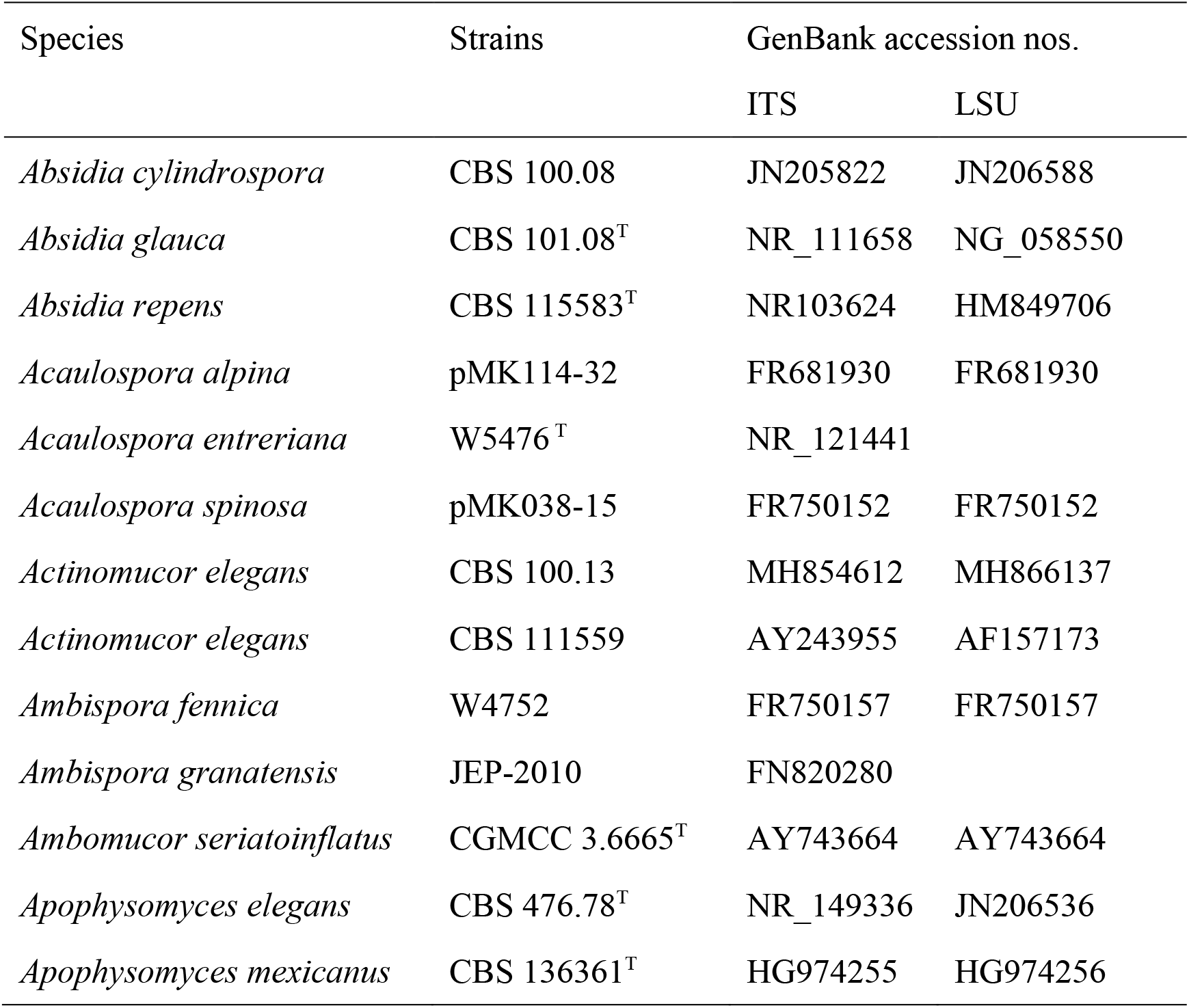
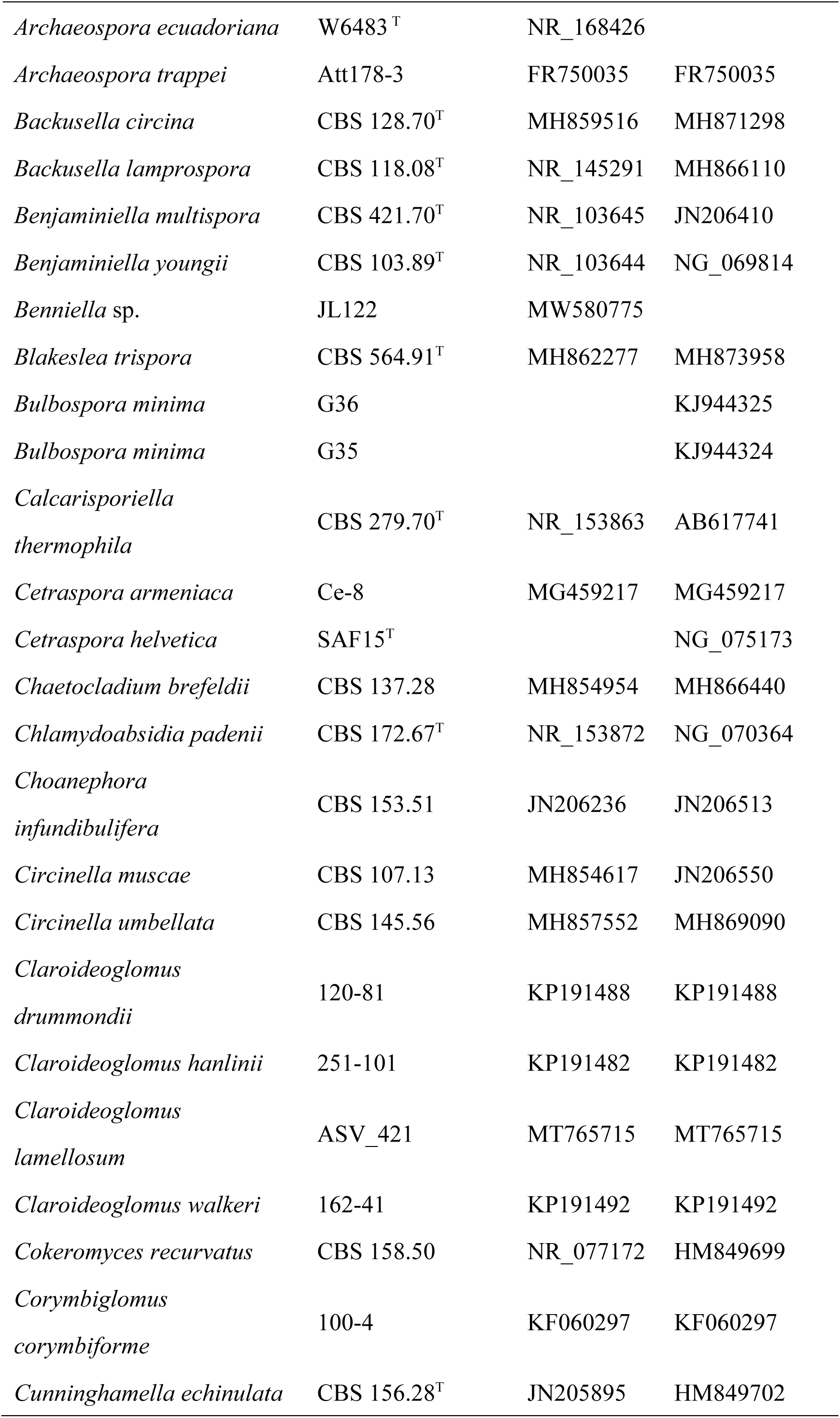
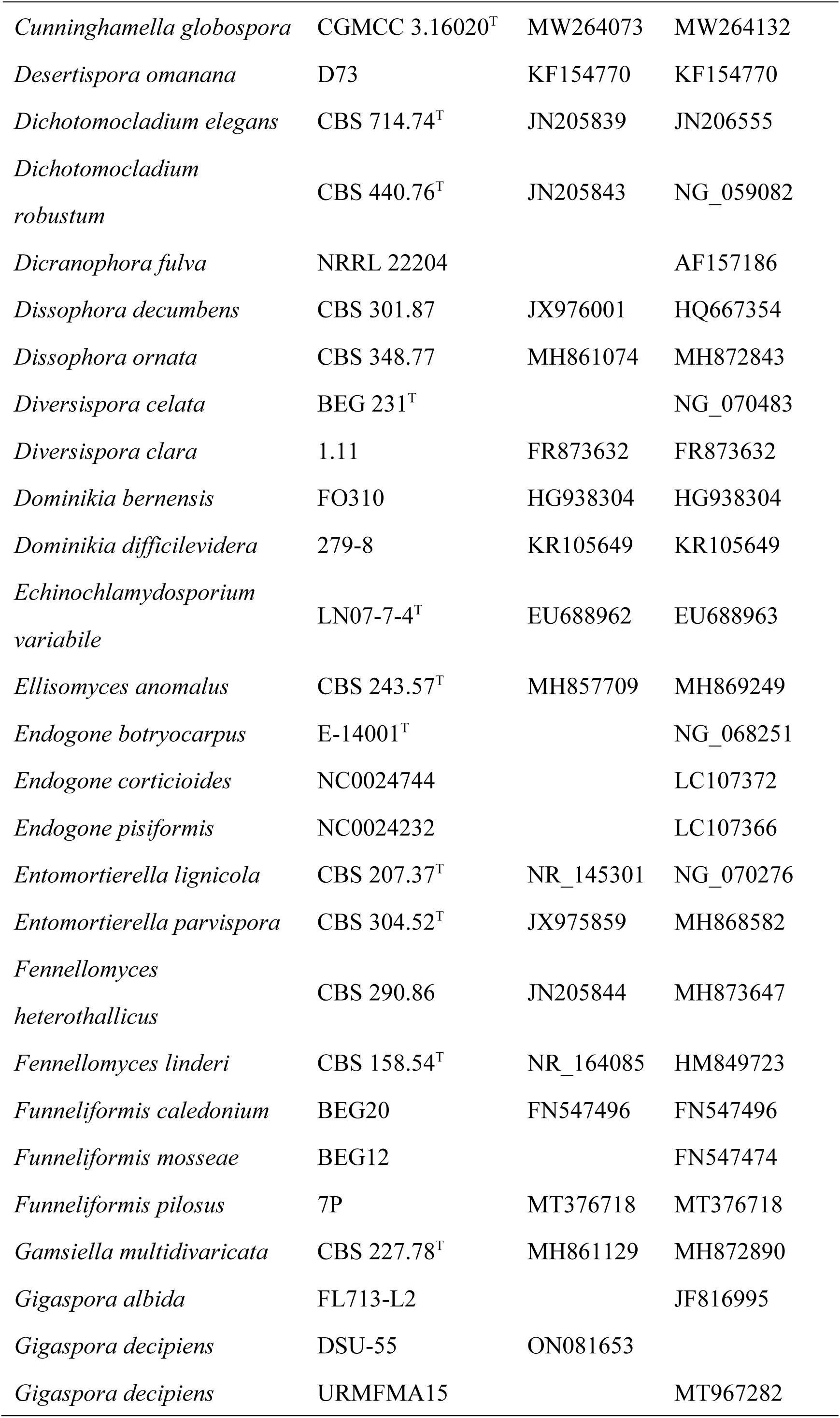
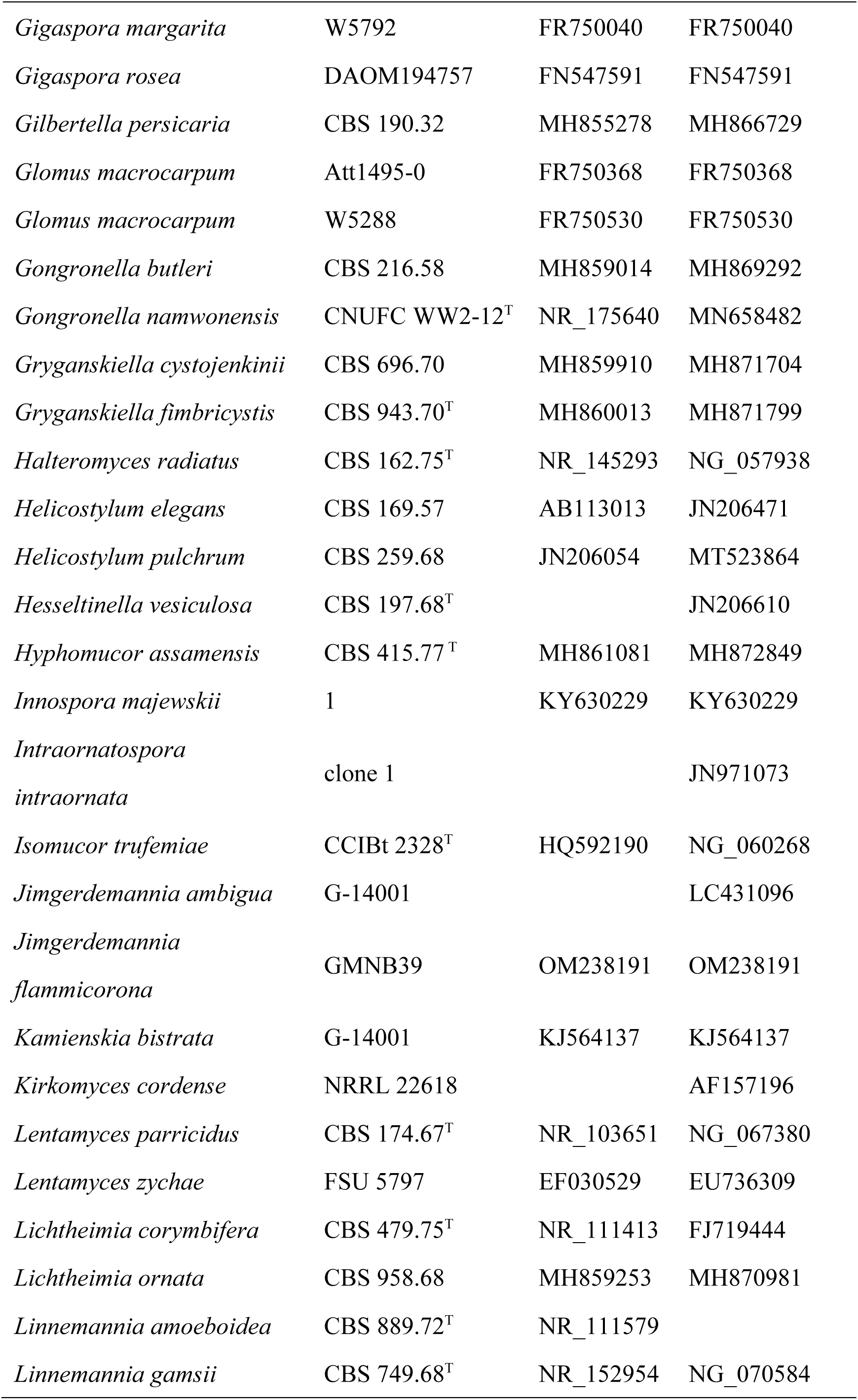
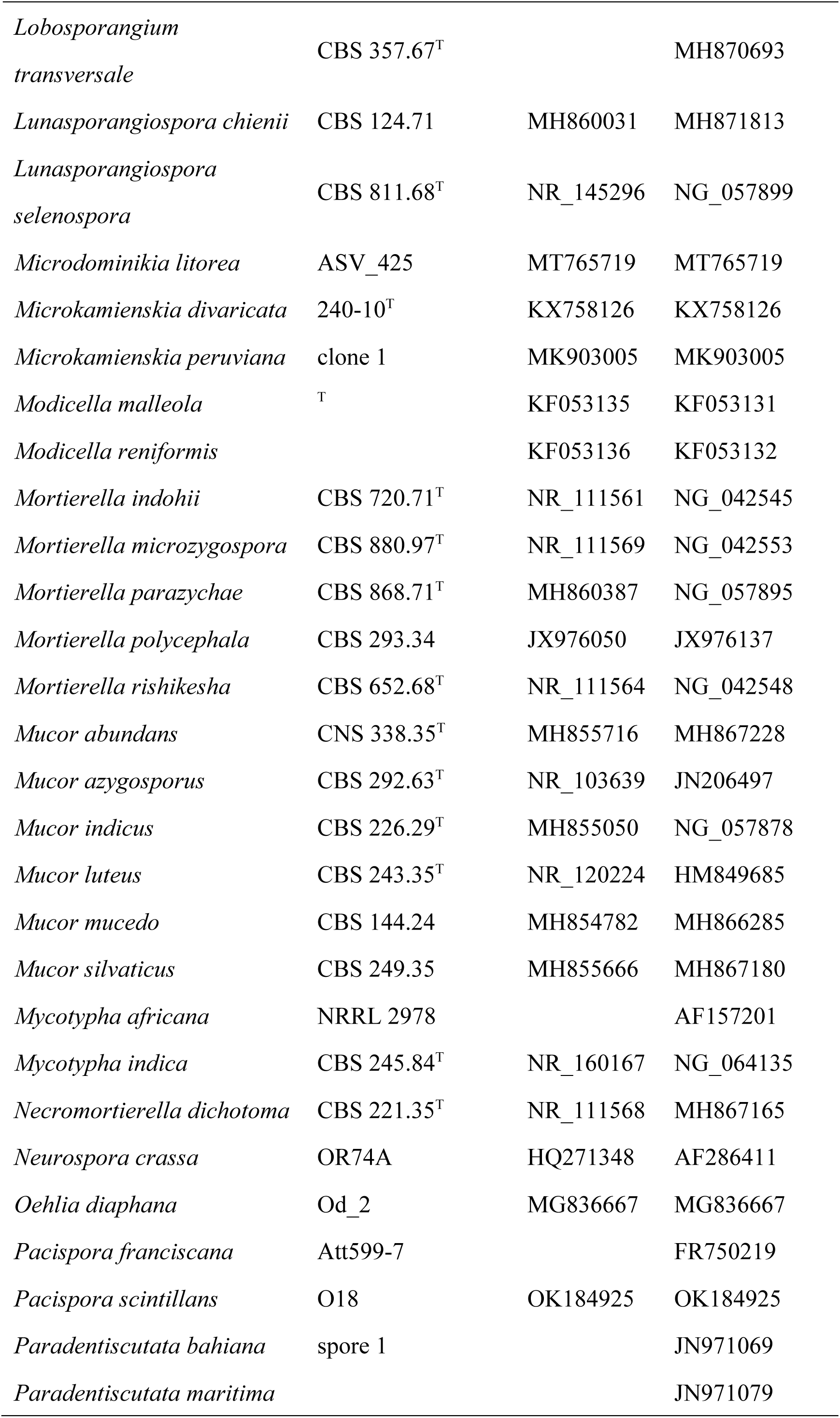
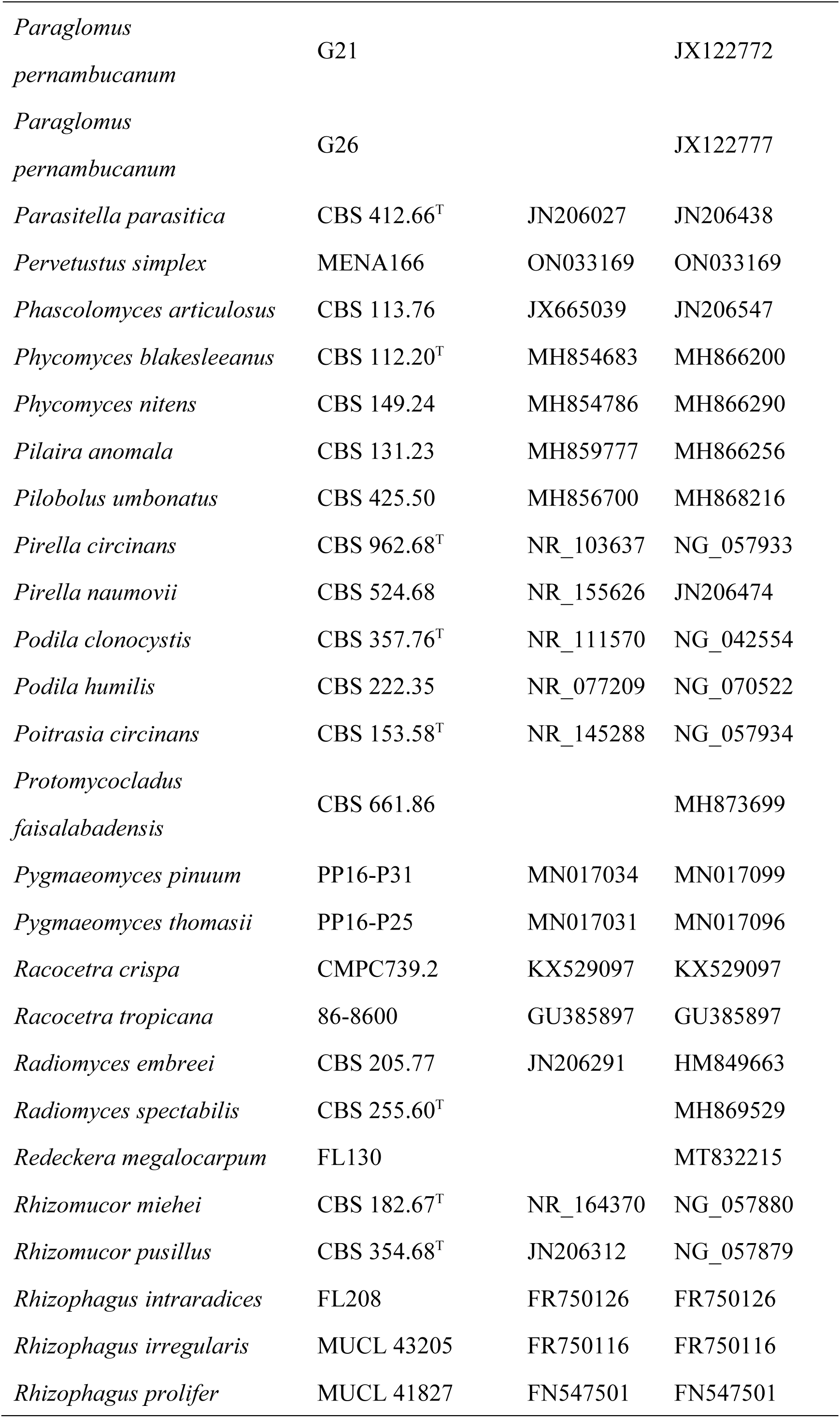
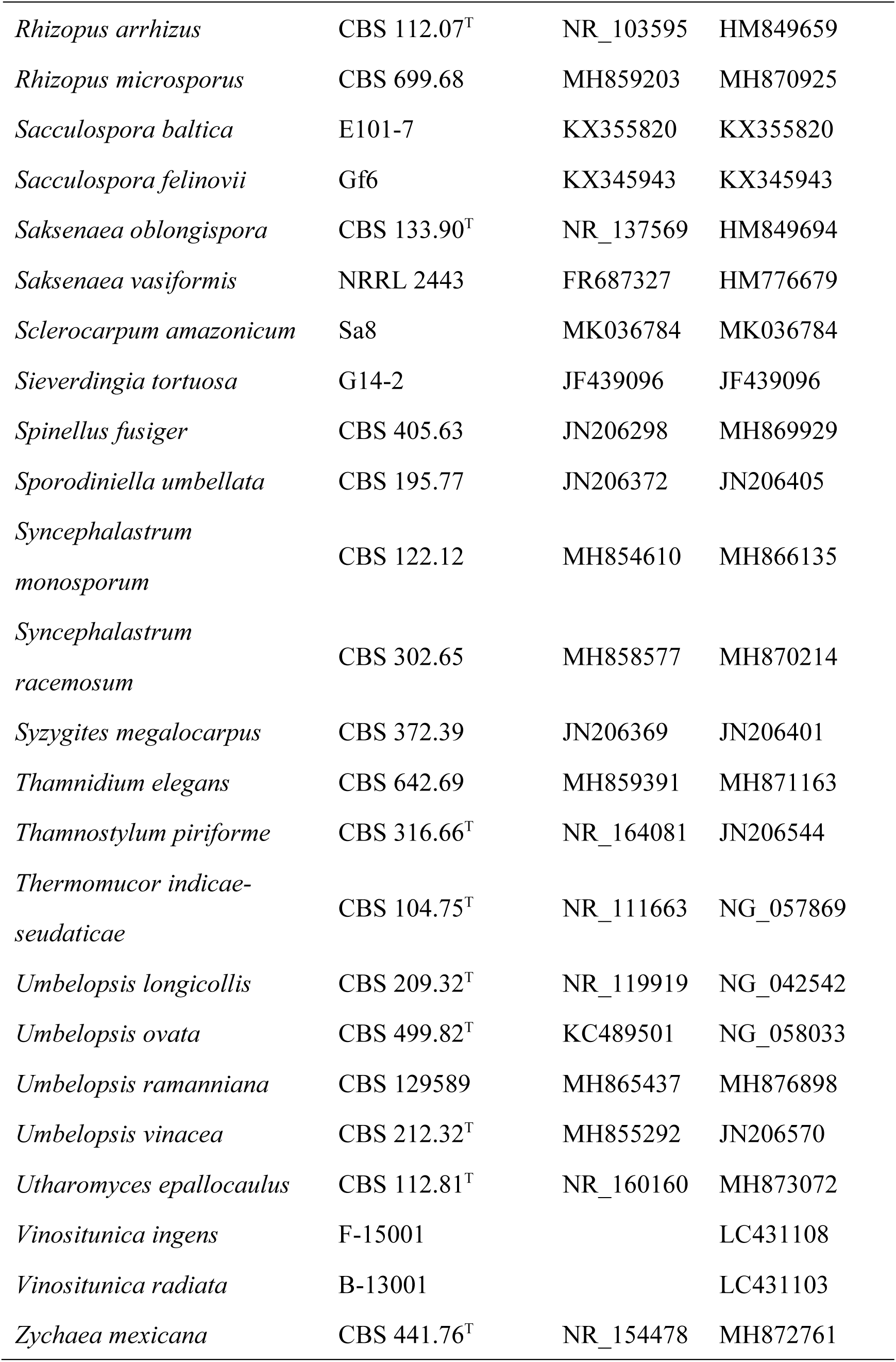
GenBank accession numbers of sequences used in this study. “T” represents ex-holotype.

| Species | Strains | GenBank accession nos. |  |
| --- | --- | --- | --- |
|  |  | ITS | LSU |
| *Absidia cylindrospora* | CBS 100.08 | JN205822 | JN206588 |
| *Absidia glauca* | CBS 101.08<sup>T</sup> | NR_111658 | NG_058550 |
| *Absidia repens* | CBS 115583<sup>T</sup> | NR103624 | HM849706 |
| *Acaulospora alpina* | pMK114-32 | FR681930 | FR681930 |
| *Acaulospora entreeriana* | W5476<sup>T</sup> | NR_121441 |  |
| *Acaulospora spinosa* | pMK038-15 | FR750152 | FR750152 |
| *Actinomucor elegans* | CBS 100.13 | MH854612 | MH866137 |
| *Actinomucor elegans* | CBS 111559 | AY243955 | AF157173 |
| *Ambispora fennica* | W4752 | FR750157 | FR750157 |
| *Ambispora granatensis* | JEP-2010 | FN820280 |  |
| *Ambomucor seriatoinflatus* | CGMCC 3.6665<sup>T</sup> | AY743664 | AY743664 |
| *Apophysomyces elegans* | CBS 476.78<sup>T</sup> | NR_149336 | JN206536 |
| *Apophysomyces mexicanus* | CBS 136361<sup>T</sup> | HG974255 | HG974256 |
| *Archaeospora ecuadoriana* | W6483<sup>T</sup> | NR_168426 |  |
| *Archaeospora trappei* | Att178-3 | FR750035 | FR750035 |
| *Backusella circina* | CBS 128.70<sup>T</sup> | MH859516 | MH871298 |
| *Backusella lamprospora* | CBS 118.08<sup>T</sup> | NR_145291 | MH866110 |
| *Benjaminiella multispora* | CBS 421.70<sup>T</sup> | NR_103645 | JN206410 |
| *Benjaminiella youngii* | CBS 103.89<sup>T</sup> | NR_103644 | NG_069814 |
| *Benniella* sp. | JL122 | MW580775 |  |
| *Blakeslea trispora* | CBS 564.91<sup>T</sup> | MH862277 | MH873958 |
| *Bulbospora minima* | G36 |  | KJ944325 |
| *Bulbospora minima* | G35 |  | KJ944324 |
| *Calcarisporiella thermophila* | CBS 279.70<sup>T</sup> | NR_153863 | AB617741 |
| *Cetraspora armeniaca* | Ce-8 | MG459217 | MG459217 |
| *Cetraspora helvetica* | SAF15<sup>T</sup> |  | NG_075173 |
| *Chaetocladium brefeldii* | CBS 137.28 | MH854954 | MH866440 |
| *Chlamydoabsidia padenii* | CBS 172.67<sup>T</sup> | NR_153872 | NG_070364 |
| *Choanephora infundibulifera* | CBS 153.51 | JN206236 | JN206513 |
| *Circinella muscae* | CBS 107.13 | MH854617 | JN206550 |
| *Circinella umbellata* | CBS 145.56 | MH857552 | MH869090 |
| *Claroideoglomerus drummondii* | 120-81 | KP191488 | KP191488 |
| *Claroideoglomerus hanlinii* | 251-101 | KP191482 | KP191482 |
| *Claroideoglomerus lamellosum* | ASV_421 | MT765715 | MT765715 |
| *Claroideoglomerus walkeri* | 162-41 | KP191492 | KP191492 |
| *Cokeromyces recurvatus* | CBS 158.50 | NR_077172 | HM849699 |
| *Corymbiglomerus corymbiforme* | 100-4 | KF060297 | KF060297 |
| *Cunninghamella echinulata* | CBS 156.28<sup>T</sup> | JN205895 | HM849702 |
| *Cunninghamella globospora* | CGMCC 3.16020<sup>T</sup> | MW264073 | MW264132 |
| *Desertispora omanana* | D73 | KF154770 | KF154770 |
| *Dichotomocladium elegans* | CBS 714.74<sup>T</sup> | JN205839 | JN206555 |
| *Dichotomocladium robustum* | CBS 440.76<sup>T</sup> | JN205843 | NG_059082 |
| *Dicranophora fulva* | NRRL 22204 |  | AF157186 |
| *Dissophora decumbens* | CBS 301.87 | JX976001 | HQ667354 |
| *Dissophora ornata* | CBS 348.77 | MH861074 | MH872843 |
| *Diversispora celata* | BEG 231<sup>T</sup> |  | NG_070483 |
| *Diversispora clara* | 1.11 | FR873632 | FR873632 |
| *Dominikia bernensis* | FO310 | HG938304 | HG938304 |
| *Dominikia difficilevidera* | 279-8 | KR105649 | KR105649 |
| *Echinochlamydosporium variabile* | LN07-7-4<sup>T</sup> | EU688962 | EU688963 |
| *Ellisomyces anomalus* | CBS 243.57<sup>T</sup> | MH857709 | MH869249 |
| *Endogone botryocarpus* | E-14001<sup>T</sup> |  | NG_068251 |
| *Endogone corticioides* | NC0024744 |  | LC107372 |
| *Endogone pisiformis* | NC0024232 |  | LC107366 |
| *Entomortierella lignicola* | CBS 207.37<sup>T</sup> | NR_145301 | NG_070276 |
| *Entomortierella parvispora* | CBS 304.52<sup>T</sup> | JX975859 | MH868582 |
| *Fennellomyces heterothallicus* | CBS 290.86 | JN205844 | MH873647 |
| *Fennellomyces linderi* | CBS 158.54<sup>T</sup> | NR_164085 | HM849723 |
| *Funneliformis caledonium* | BEG20 | FN547496 | FN547496 |
| *Funneliformis mosseae* | BEG12 |  | FN547474 |
| *Funneliformis pilosus* | 7P | MT376718 | MT376718 |
| *Gamsiella multivaricata* | CBS 227.78<sup>T</sup> | MH861129 | MH872890 |
| *Gigaspora albida* | FL713-L2 |  | JF816995 |
| *Gigaspora decipiens* | DSU-55 | ON081653 |  |
| *Gigaspora decipiens* | URMFMA15 |  | MT967282 |
| *Gigaspora margarita* | W5792 | FR750040 | FR750040 |
| *Gigaspora rosea* | DAOM194757 | FN547591 | FN547591 |
| *Gilbertella persicaria* | CBS 190.32 | MH855278 | MH866729 |
| *Glomus macrocarpum* | Att1495-0 | FR750368 | FR750368 |
| *Glomus macrocarpum* | W5288 | FR750530 | FR750530 |
| *Gongronella butleri* | CBS 216.58 | MH859014 | MH869292 |
| *Gongronella namwonensis* | CNUFC WW2-12<sup>T</sup> | NR_175640 | MN658482 |
| *Gryganskiella cystojenkinii* | CBS 696.70 | MH859910 | MH871704 |
| *Gryganskiella fimbricystis* | CBS 943.70<sup>T</sup> | MH860013 | MH871799 |
| *Halteromyces radiatus* | CBS 162.75<sup>T</sup> | NR_145293 | NG_057938 |
| *Helicostylum elegans* | CBS 169.57 | AB113013 | JN206471 |
| *Helicostylum pulchrum* | CBS 259.68 | JN206054 | MT523864 |
| *Hesseltinella vesiculosa* | CBS 197.68<sup>T</sup> |  | JN206610 |
| *Hyphomucor assamensis* | CBS 415.77<sup>T</sup> | MH861081 | MH872849 |
| *Innospora majewskii* | 1 | KY630229 | KY630229 |
| *Intraornatospora* |  |  |  |
| *intraornata* | clone 1 |  | JN971073 |
| *Isomucor trufemiae* | CCIBt 2328<sup>T</sup> | HQ592190 | NG_060268 |
| *Jimgerdemannia ambigua* | G-14001 |  | LC431096 |
| *Jimgerdemannia* |  |  |  |
| *flammicorona* | GMNB39 | OM238191 | OM238191 |
| *Kamienskia bistrata* | G-14001 | KJ564137 | KJ564137 |
| *Kirkomyces cordense* | NRRL 22618 |  | AF157196 |
| *Lentamyces parricidus* | CBS 174.67<sup>T</sup> | NR_103651 | NG_067380 |
| *Lentamyces zychae* | FSU 5797 | EF030529 | EU736309 |
| *Lichtheimia corymbifera* | CBS 479.75<sup>T</sup> | NR_111413 | FJ719444 |
| *Lichtheimia ornata* | CBS 958.68 | MH859253 | MH870981 |
| *Linnemannia amoeboides* | CBS 889.72<sup>T</sup> | NR_111579 |  |
| *Linnemannia gamsii* | CBS 749.68<sup>T</sup> | NR_152954 | NG_070584 |
| *Lobosporangium transversale* | CBS 357.67<sup>T</sup> |  | MH870693 |
| *Lunasporangiospora chienii* | CBS 124.71 | MH860031 | MH871813 |
| *Lunasporangiospora selenospora* | CBS 811.68<sup>T</sup> | NR_145296 | NG_057899 |
| *Microdominikia litorea* | ASV_425 | MT765719 | MT765719 |
| *Microkamiensia divaricata* | 240-10<sup>T</sup> | KX758126 | KX758126 |
| *Microkamiensia peruviana* | clone 1 | MK903005 | MK903005 |
| *Modicella malleola* | <sup>T</sup> | KF053135 | KF053131 |
| *Modicella reniformis* |  | KF053136 | KF053132 |
| *Mortierella indohii* | CBS 720.71<sup>T</sup> | NR_111561 | NG_042545 |
| *Mortierella microzygospora* | CBS 880.97<sup>T</sup> | NR_111569 | NG_042553 |
| *Mortierella parazychae* | CBS 868.71<sup>T</sup> | MH860387 | NG_057895 |
| *Mortierella polycephala* | CBS 293.34 | JX976050 | JX976137 |
| *Mortierella rishiksha* | CBS 652.68<sup>T</sup> | NR_111564 | NG_042548 |
| *Mucor abundans* | CNS 338.35<sup>T</sup> | MH855716 | MH867228 |
| *Mucor azygosporus* | CBS 292.63<sup>T</sup> | NR_103639 | JN206497 |
| *Mucor indicus* | CBS 226.29<sup>T</sup> | MH855050 | NG_057878 |
| *Mucor luteus* | CBS 243.35<sup>T</sup> | NR_120224 | HM849685 |
| *Mucor mucedo* | CBS 144.24 | MH854782 | MH866285 |
| *Mucor silvaticus* | CBS 249.35 | MH855666 | MH867180 |
| *Mycotypha africana* | NRRL 2978 |  | AF157201 |
| *Mycotypha indica* | CBS 245.84<sup>T</sup> | NR_160167 | NG_064135 |
| *Necromortierella dichotoma* | CBS 221.35<sup>T</sup> | NR_111568 | MH867165 |
| *Neurospora crassa* | OR74A | HQ271348 | AF286411 |
| *Oehlia diaphana* | Od_2 | MG836667 | MG836667 |
| *Pacispora franciscana* | Att599-7 |  | FR750219 |
| *Pacispora scintillans* | O18 | OK184925 | OK184925 |
| *Paradentiscutata bahiana* | spore 1 |  | JN971069 |
| *Paradentiscutata maritima* |  |  | JN971079 |
| *Paraglomus* | G21 |  | JX122772 |
| *pernambucanum* |  |  |  |
| *Paraglomus* | G26 |  | JX122777 |
| *pernambucanum* |  |  |  |
| *Parasitella parasitica* | CBS 412.66<sup>T</sup> | JN206027 | JN206438 |
| *Pervetustus simplex* | MENA166 | ON033169 | ON033169 |
| *Phascolomyces articulatus* | CBS 113.76 | JX665039 | JN206547 |
| *Phycomyces blakesleeana* | CBS 112.20<sup>T</sup> | MH854683 | MH866200 |
| *Phycomyces nitens* | CBS 149.24 | MH854786 | MH866290 |
| *Pilaira anomala* | CBS 131.23 | MH859777 | MH866256 |
| *Pilobolus umbonatus* | CBS 425.50 | MH856700 | MH868216 |
| *Pirella circinans* | CBS 962.68<sup>T</sup> | NR_103637 | NG_057933 |
| *Pirella naumovii* | CBS 524.68 | NR_155626 | JN206474 |
| *Podila clonocystis* | CBS 357.76<sup>T</sup> | NR_111570 | NG_042554 |
| *Podila humilis* | CBS 222.35 | NR_077209 | NG_070522 |
| *Poitrasia circinans* | CBS 153.58<sup>T</sup> | NR_145288 | NG_057934 |
| *Protomyocladus* |  |  |  |
| *faisalabadensis* | CBS 661.86 |  | MH873699 |
| *Pygmaeomyces pinuum* | PP16-P31 | MN017034 | MN017099 |
| *Pygmaeomyces thomasi* | PP16-P25 | MN017031 | MN017096 |
| *Racocetra crispa* | CMPC739.2 | KX529097 | KX529097 |
| *Racocetra tropicana* | 86-8600 | GU385897 | GU385897 |
| *Radiomyces embreei* | CBS 205.77 | JN206291 | HM849663 |
| *Radiomyces spectabilis* | CBS 255.60<sup>T</sup> |  | MH869529 |
| *Redeckera megalocarpum* | FL130 |  | MT832215 |
| *Rhizomucor miehei* | CBS 182.67<sup>T</sup> | NR_164370 | NG_057880 |
| *Rhizomucor pusillus* | CBS 354.68<sup>T</sup> | JN206312 | NG_057879 |
| *Rhizophagus intraradices* | FL208 | FR750126 | FR750126 |
| *Rhizophagus irregularis* | MUCL 43205 | FR750116 | FR750116 |
| *Rhizophagus prolifer* | MUCL 41827 | FN547501 | FN547501 |
| *Rhizopus arrhizus* | CBS 112.07<sup>T</sup> | NR_103595 | HM849659 |
| *Rhizopus microsporus* | CBS 699.68 | MH859203 | MH870925 |
| *Sacculospora baltica* | E101-7 | KX355820 | KX355820 |
| *Sacculospora felinovii* | Gf6 | KX345943 | KX345943 |
| *Saksenaea oblongispora* | CBS 133.90<sup>T</sup> | NR_137569 | HM849694 |
| *Saksenaea vasiformis* | NRRL 2443 | FR687327 | HM776679 |
| *Sclerocarpum amazonicum* | Sa8 | MK036784 | MK036784 |
| *Sieverdingia tortuosa* | G14-2 | JF439096 | JF439096 |
| *Spinellus fusiger* | CBS 405.63 | JN206298 | MH869929 |
| *Sporodiniella umbellata* | CBS 195.77 | JN206372 | JN206405 |
| *Syncephalastrum monosporum* | CBS 122.12 | MH854610 | MH866135 |
| *Syncephalastrum racemosum* | CBS 302.65 | MH858577 | MH870214 |
| *Syzygites megalocarpus* | CBS 372.39 | JN206369 | JN206401 |
| *Thamnidium elegans* | CBS 642.69 | MH859391 | MH871163 |
| *Thamnostylum piriforme* | CBS 316.66<sup>T</sup> | NR_164081 | JN206544 |
| *Thermomucor indicae-seudaticae* | CBS 104.75<sup>T</sup> | NR_111663 | NG_057869 |
| *Umbelopsis longicollis* | CBS 209.32<sup>T</sup> | NR_119919 | NG_042542 |
| *Umbelopsis ovata* | CBS 499.82<sup>T</sup> | KC489501 | NG_058033 |
| *Umbelopsis ramanniana* | CBS 129589 | MH865437 | MH876898 |
| *Umbelopsis vinacea* | CBS 212.32<sup>T</sup> | MH855292 | JN206570 |
| *Utharomyces epallocaulus* | CBS 112.81<sup>T</sup> | NR_160160 | MH873072 |
| *Vinositunica ingens* | F-15001 |  | LC431108 |
| *Vinositunica radiata* | B-13001 |  | LC431103 |
| *Zychaea mexicana* | CBS 441.76<sup>T</sup> | NR_154478 | MH872761 |

**Table 4.**
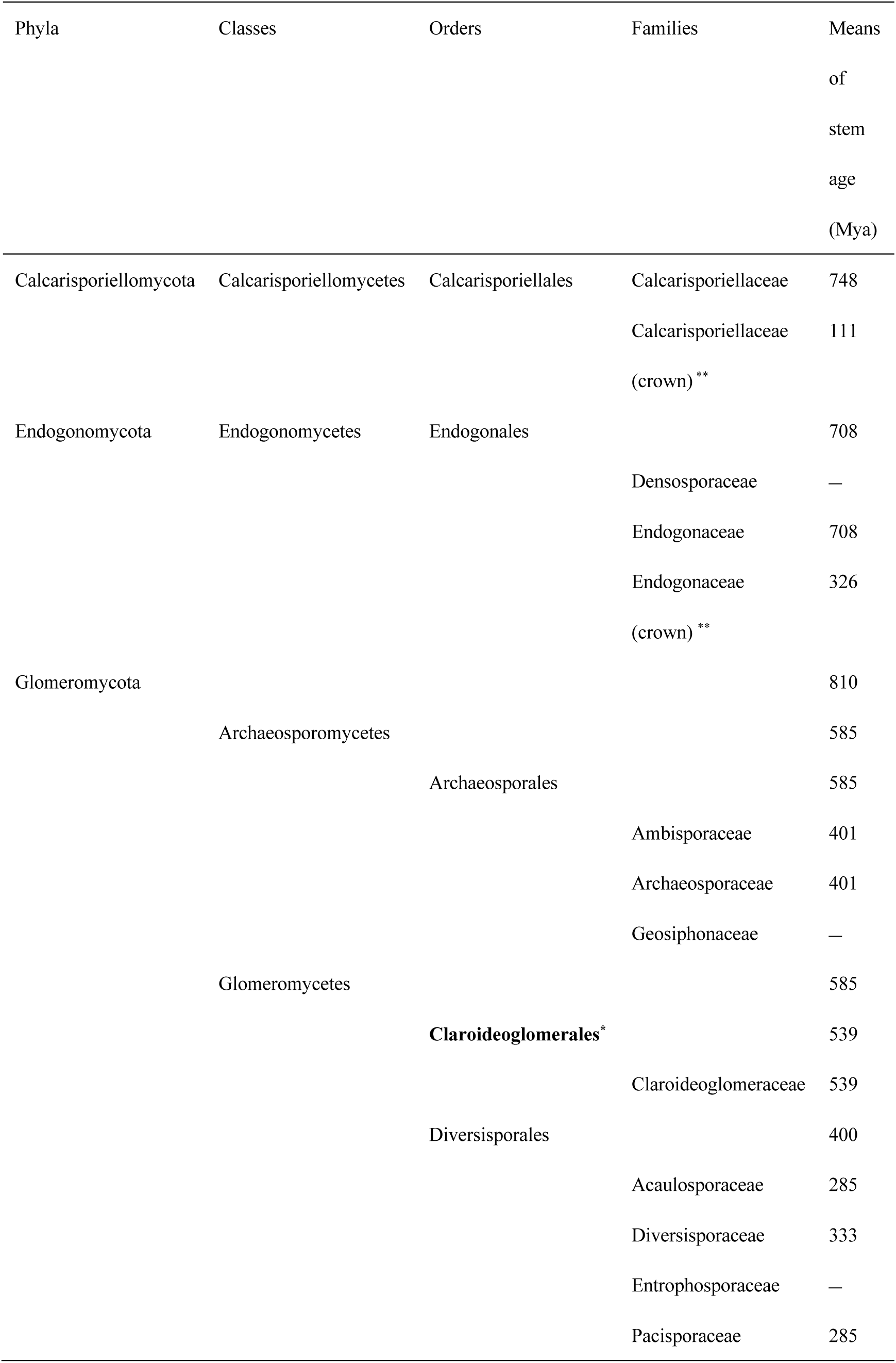
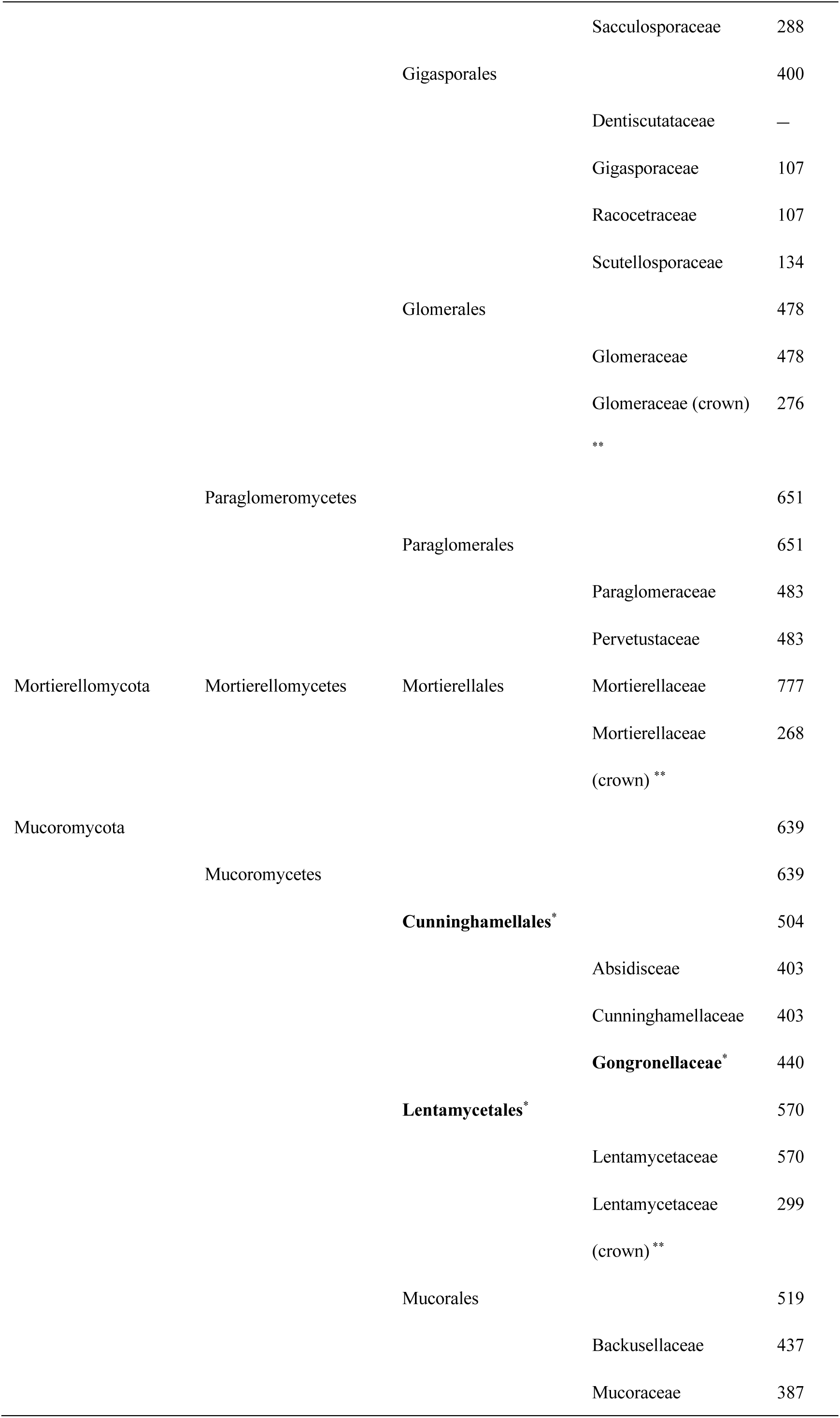
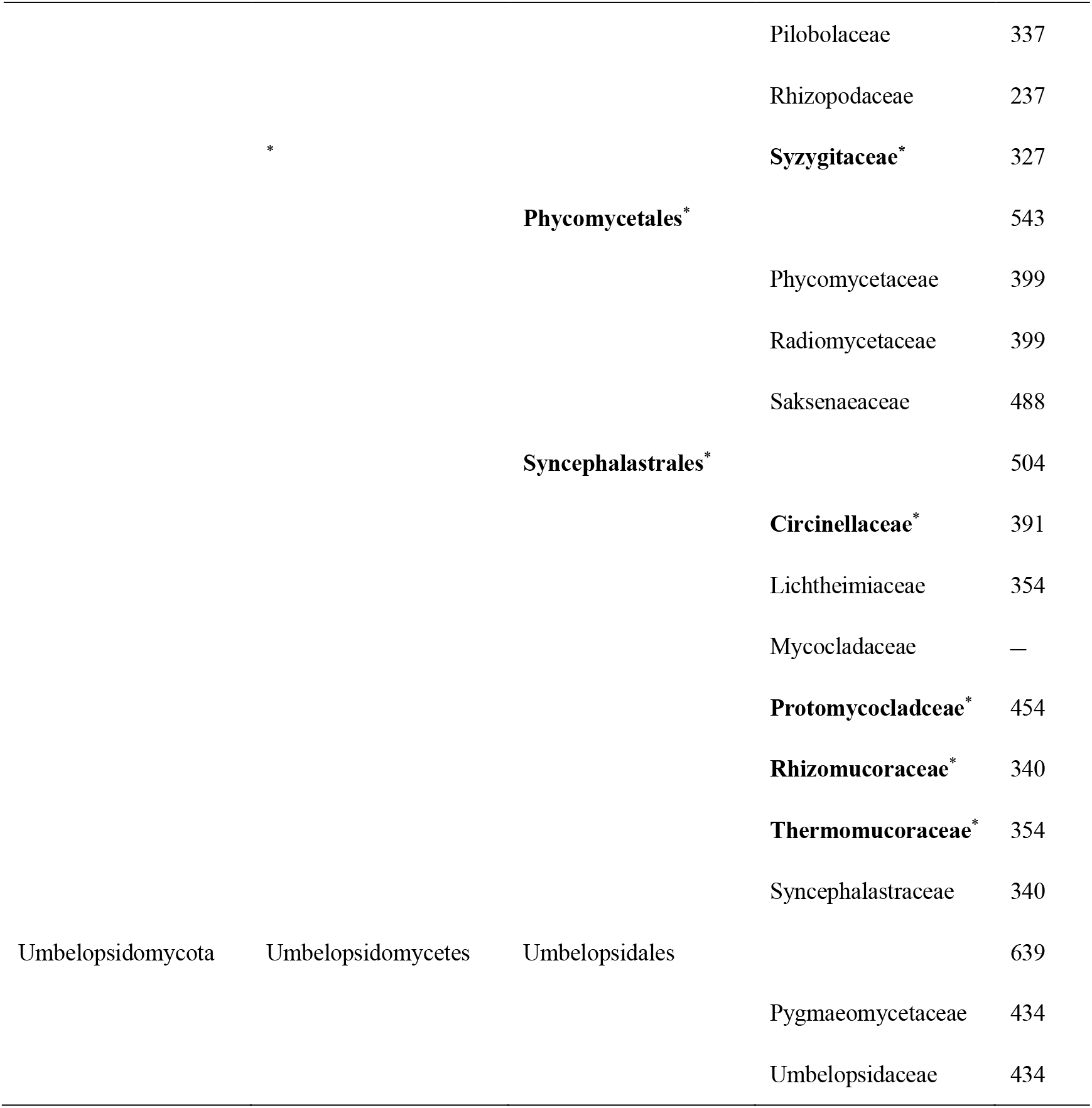
Inferred divergence time of higher taxa in subkingdom Mucoromyceta. * New taxa proposed in the present study are in bold. ** Means of mean crown age (Mya). “–” represents the absence of divergence time in this study.

We accepted the phylum Glomeromycota, and three classes Archaeosporomycetes (585 Mya / PP = 0.9), Glomeromycetes (585 Mya / PP = 0.9) and Paraglomeromycetes (651 Mya / PP = 1.0) were described (Schüßler et al. 2001, Oehl et al. 2011a, b, c). The Archaeosporomycetes contained one class Archaeosporales (585 Mya / PP = 0.9) and three families (Ambisporaceae 401 Mya / PP = 1.0 and Archaeosporaceae 401 Mya / PP = 1.0, while Geosiphonaceae absent). The Glomeromycetes held four classes (Claroideoglomerales 539 Mya / PP = 1.0, Diversisporales 400 Mya / PP = 1.0, Gigasporales 400 Mya / PP = 1.0 and Glomerales 478 Mya / PP = 1.0). The Claroideoglomerales comprised one family Claroideoglomeraceae with 539 Mya / PP = 1.0. The Diversisporales consisted of five families (Acaulosporaceae 285 Mya, Diversisporaceae 333 Mya / PP = 1.0, Pacisporaceae 285 and Sacculosporaceae 288, while Entrophosporaceae absent). The Gigasporales comprised four families (Gigasporaceae 107 Mya / PP = 1.0, Racocetraceae 107 Mya / PP = 1.0 and Scutellosporaceae 134 Mya / PP = 1.0, while Dentiscutataceae absent). The Glomerales was one family described (Glomeraceae 478 Mya / PP = 1.0 and mean crown age of Glomeraceae 276 Mya / PP = 1.0). The Paraglomeromycetes included one class (Paraglomerales 651 Mya / PP = 1.0) and two families (Paraglomeraceae 483 Mya / PP = 1.0 and Pervetustaceae 483 Mya / PP = 1.0).

The Mortierellomycota comprised one class (Mortierellomycetes), one order (Mortierellales) and one family Mortierellaceae (mean stem age 777 Mya / PP = 0.9, mean crown age 268 Mya / PP = 1.0).

The Calcarisporiellomycota accommodated one class, one order and one family Calcarisporiellaceae (mean stem age 728 Mya / PP = 1.0, mean crown age 111 Mya /PP = 1.0).

The Endogonomycota contained one class, one order and two families Endogonaceae (mean stem age 708 Mya / PP = 1.0, mean crown age 326 Mya / PP = 1.0) and Densosporaceae (data unavailable).

The Mucoromycota held one class (Mucoromycetes 639 Mya with a 1.0 PP support). The Mucoromycetes evolved into five orders (Cunninghamellales 504 Mya, Lentamycetales 570 Mya / PP =1.0, Mucorales 519 Mya / PP =1.0, Phycomycetales 543 Mya / PP =1.0 and Syncephalastrales 504 Mya). The Cunninghamellales consisted of three families (Absidiaceae 403 Mya / PP = 1.0, Cunninghamellaceae 403 Mya / PP = 1.0 and Gongronaceae 440 Mya / PP =1.0). The Lentamycetales was monotypic with 570 Mya / PP = 0.8 and mean crown ages 299 Mya / PP =1.0. The monotypic Mucorales diverged into five families (Backusellaceae 437 Mya / PP =1.0, Mucoraceae 387 Mya / PP =1.0, Pilobolaceae 337 Mya / PP =1.0, Rhizopodaceae 237 Mya / PP =1.0, and Syzygitaceae 237 Mya / PP =1.0). The Phycomycetales contained three families (Phycomycetaceae 399 Mya / PP = 1.0, Radiomycetaceae 399 Mya / PP = 1.0, and Saksenaeaceae 488 Mya / PP = 1.0). Syncephalastromycetales held seven families (Circinellaceae 391 Mya, Lichtheimiaceae 354 Mya, Protomycocladaceae 454 Mya / PP = 1.0, Rhizomucoraceae 340 Mya / PP = 0.8, Syncephalastraceae 340 Mya / PP = 0.8, Thermomucoraceae 354 Mya, and Mycocladaceae data unavailable).

The Umbelopsidomycota consisted of one class (Umbelopsidomycetes), one order (Umbelopsidales) and two families (Pygmaeomycetaceae 434 Mya / PP =1.0 and Umbelopsidaceae 434 Mya / PP =1.0).

### New taxa at high ranks

Comprehensively considering morphological features, divergence dating and monophyletic concept, a total of thirteen new taxa, including two phyla, five orders and six families, are proposed herein.

### New phyla

**Endogonomycota** H. Zhao, Y.C. Dai, B.K. Cui, F. Wu & X.Y. Liu, *phyl. nov*.

*Fungal Names*: FN570870.

*Type*: *Endogone* Link, Mag. Gesell. naturf. Freunde, Berlin 3(1–2): 33, 1809.

*Diagnosis*: Mycelium well-known to be ectomycorrhizal with plants or sometimes saprobic. Sporangiospores absent. Chlamydospores present or absent. Zygospores always formed in sporocarps.

**Umbelopsidomycota** H. Zhao, Y.C. Dai, B.K. Cui, F. Wu & X.Y. Liu, *phyl. nov*.

*Fungal Names*: FN570871.

*Type*: *Umbelopsis* Amos & H.L. Barnett, Mycologia 58(5): 807, 1966.

*Diagnosis*: Colonies frequently sectoring due to a possible genetic instability. Aerial hyphae so degenerated that the colonies are a thin layer of sporangiophores. Sporangiophores arising mainly from substrate hyphae and rarely from aerial hyphae. Sporangia one- or multi-spored, subglobose to globose, typically pigmented, resulting in reddish or ochraceous colonies. Collars and columellae obvious or degenerated. Sporangiospores globose, subglobose, teardrop-shaped, angular or irregular. Chlamydospores present in substrate hyphae. Zygospores unknown.

### New orders

**Claroideoglomerales** H. Zhao, Y.C. Dai, B.K. Cui, F. Wu & X.Y. Liu, *ord. nov*.

*Fungal Names*: FN571005.

*Type*: *Claroideoglomus* C. Walker & A. Schüßler, 2010.

*Diagnosis*: Spores single, commonly in soil or rarely in plant roots. Subtending hyphae hyaline to white, and rarely subhyaline, conspicuously bill-shaped. Spores with one to four layers, with a pore closure at the base which allowing spore to germinate from the structural layer, or from an adherent innermost (semi-)flexible layer, or from both layers.

**Cunninghamellales** H. Zhao, Y.C. Dai, B.K. Cui, F. Wu & X.Y. Liu, *ord. nov*.

*Fungal Names*: FN570873.

*Type*: *Cunninghamella* Matr., Annls Mycol. 1(1): 46, 1903.

*Diagnosis*: Sporangiophores arising from substrate and aerial hyphae, or from stolons. Rhizoids always present, finger-like or root-like. Stolons present. Apophyses present. Asexual reproduction by either sporangia or sporangiola. Sporangia pyriform to globose, multi-spored. Sporangiola borne on pedicels on vesicles, with spines all over the surface, containing one smooth sporangiospore. Chlamydospores unknown. Zygospores if present, always with appendages.

**Lentamycetales** H. Zhao, Y.C. Dai, B.K. Cui, F. Wu & X.Y. Liu, *ord. nov*.

*Fungal Names*: FN570874.

*Type*: *Lentamyces* Kerst. Hoffm. & K. Voigt, Pl. Biol. 11(4): 550, 2009.

*Diagnosis*: Colonies slow growing. Maximum growth temperatures < 30 °C. Sporangiophores erect, unbranched, without whorls. Sporangia spherical to subpyriform, multi-spored. Columellae spherical to hemispherical, without terminal appendages. Sporangiospores ovoid to cylindrical. Rhizoids present. Stolons present. Apophyses present. Projections absent. Zygospores without appendages.

**Phycomycetales** H. Zhao, Y.C. Dai, B.K. Cui, F. Wu & X.Y. Liu, *ord. nov*.

*Fungal Names*: FN570875.

*Type*: *Phycomyces* Kunze, Mykologische Hefte (Leipzig) 2: 113, 1823.

*Diagnosis*: Asexual reproduction by either multi-spored sporangia or uni-spored sporangiola, with walls either deliquescent or persistent. Apophyses typically present. Zygospores if present, with suspensors opposed.

**Syncephalastrales** H. Zhao, Y.C. Dai, B.K. Cui, F. Wu & X.Y. Liu, *ord. nov*.

*Fungal Names*: FN570876.

*Type*: *Syncephalastrum* J. Schröt., Krypt.-Fl. Schlesien (Breslau) 3.1(9–16): 217, 1886.

*Diagnosis*: Sporangiophores arising from aerial or substrate hyphae, unbranched, simple branched or sympodially branched. Rhizoids present or absent. Stolons present or absent. Sporangia multi-spored, columellate, either apophysate or nonapophysate. Chlamydospores absent or scarcely produced. Zygospores if present, with suspensors opposed.

### New families

**Circinellaceae** H. Zhao, Y.C. Dai, B.K. Cui, F. Wu & X.Y. Liu, *fam. nov*.

*Fungal Names*: FN570878.

*Type*: *Circinella* Tiegh. & G. Le Monn., Annls Sci. Nat., Bot., sér. 5 17: 298, 1873.

*Diagnosis*: Most recently originated in the Syncephalastrales clade in phylogenetic divergence time analyses. Sporangiophores arising from substrate and aerial hyphae, simple or branched, main stem straight and lateral branches circinate, curved or twisted. Terminal sporangia always subglobose to globose, multi-spored. Lateral sporangia subglobose to globose, uni-to multi-spored. Apophyses present. Sporangiospores often ovoid to ellipsoid. Chlamydospores sometimes present in substrate hyphae. Zygospores if known, ornamented, pigmented, heterothallic or homothallic, with suspensors opposed.

**Gongronellaceae** H. Zhao, Y.C. Dai, B.K. Cui, F. Wu & X.Y. Liu, *fam. nov*.

*Fungal Names*: FN570879.

*Diagnosis*: *Gongronella* Ribaldi, Riv. Biol. 44: 164, 1952.

*Description*: Basal in the Cunninghamellales clade in phylogenetic and divergence time analyses. Sporangiophores arising from substrate and aerial hyphae, erect to curved, unbranched. Sporangia globose. Zygospores if known, ornamented, pigmented, heterothallic, with suspensors opposed.

**Protomycocladaceae** H. Zhao, Y.C. Dai, B.K. Cui, F. Wu & X.Y. Liu, *fam. nov*.

*Fungal Names*: FN570880.

*Type*: *Protomycocladus* Schipper & Samson, Mycotaxon 50: 487, 1994

*Diagnosis*: Basal in the order Syncephalastrales clade in phylogenetic and divergence time analyses. Sporangiophores arising from substrate hyphae, sympodially branched. Apophyses present. Sporangia apophysate, smooth, pyriform and multi-spored, walls deliquescent. Zygospores ornamented, homothallic, with suspensors opposed.

**Rhizomucoraceae** H. Zhao, Y.C. Dai, B.K. Cui, F. Wu & X.Y. Liu, *fam. nov*.

*Fungal Names*: FN570881.

*Type*: *Rhizomucor* Lucet & Costantin, Rev. gén. Bot. 12: [81], 1900.

*Diagnosis*: Next to the Protocladaceae clade and the Syncephalastraceae clade in the Syncephalastrales group in phylogenetic and divergence time analyses. Rhizoids always present. Sporangiophores arising from hyphae, branched. Apophyses absent. Sporangia columellate and multi-spored. Chlamydospores sometimes present. Zygospores ornamented, with opposed suspensors and a pigmented zygosporangial wall. Mesophilic and thermophilic.

**Syzygitaceae** H. Zhao, Y.C. Dai, B.K. Cui, F. Wu & X.Y. Liu, *fam. nov*.

*Fungal Names*: FN571006.

*Type*: *Syzygites* Ehrenb., Sylv. mycol. berol. (Berlin): 25, 1818.

*Diagnosis*: Sporangiophores arising directly from substrate hyphae, umbel or dichotomously branched. Sporangia subglobose to globose, columellate, few-spored or multi-spored, deliquescent-walled. Sporangiospores globose to ovoid, spinose-walled. Zygospores ornamented, with suspensors opposed.

**Thermomucoraceae** H. Zhao, Y.C. Dai, B.K. Cui, F. Wu & X.Y. Liu, *fam. nov*.

*Fungal Names*: FN571007.

*Type*: *Thermomucor* Subrahm., B.S. Mehrotra & Thirum., Georgia J. Sci. 35(1): 1, 1977. *Diagnosis*: Sporangiophores arising from stolons and rhizoids, branched. Sporangia globose, columellate, multi-spored, and apophysate. Sporangiospores subglobose, smooth-walled. Zygospores produced from aerial hyphae, with opposed suspensors and a smooth zygosporangial wall. Homothallic and thermophilic.

### Outline of the subkingdom Mucoromyceta

**Subkingdom Mucoromyceta** Tedersoo, Sánchez-Ramírez, Kõljalg, Bahram, Döring, Schigel, T. May, M. Ryberg & Abarenkov 2018

**Phylum Calcarisporiellomycota** Tedersoo, Sánchez-Ramírez, Kõljalg, Bahram, Döring, Schigel, T. May, M. Ryberg & Abarenkov 2018

**Class Calcarisporiellomycetes** Tedersoo, Sánchez-Ramírez, Kõljalg, Bahram, Döring, Schigel, T. May, M. Ryberg & Abarenkov 2018

**Order Calcarisporiellales** Tedersoo, Sánchez-Ramírez, Kõljalg, Bahram, Döring, Schigel, T. May, M. Ryberg & Abarenkov 2018

**Family Calcarisporiellaceae** Tedersoo, Sánchez-Ramírez, Kõljalg, Bahram, Döring, Schigel, T. May, M. Ryberg & Abarenkov 2018

*Calcarisporiella* de Hoog 1974

*Echinochlamydosporium* X.Z. Jiang, H.Y. Yu, M.C. Xiang, X.Y. Liu & Xing Z. Liu 2011

**Phylum Endogonomycota** H. Zhao, Y.C. Dai, B.K. Cui, F. Wu & X.Y. Liu, *phyl. nov*.

**Class Endogonomycetes** Doweld 2014

**Order Endogonales** Jacz. & P.A. Jacz. 1931

**Family Densosporaceae** Desirò, M.E. Sm., Bidartondo, Trappe & Bonito 2017

*Densospora* McGee 1996

**Family Endogonaceae** Paol. 1889

*Bifiguratus* Torr.-Cruz & Porras-Alfaro 2017 *Endogone* Link 1809

*Jimgerdemannia* Trappe, Desirò, M.E. Sm., Bonito & Bidartondo 2017

*Peridiospora* C.G. Wu & Suh J. Lin 1997

*Sclerogone* Warcup 1990

*Vinositunica* Koh. Yamam., Degawa & A. Yamada 2020

**Phylum Glomeromycota C. Walker & A. Schüßler 2001**

**Class Archaeosporomycetes** Sieverd., G.A. Silva, B.T. Goto & Oehl 2011

**Order Archaeosporales** C. Walker & A. Schüßler 2001

**Family Ambisporaceae** C. Walker, Vestberg & A. Schüßler 2007

*Ambispora* C. Walker, Vestberg & A. Schüßler 2007

**Family Archaeosporaceae** J.B. Morton & D. Redecker 2001

*Archaeospora* J.B. Morton & D. Redecker 2001

**Family Geosiphonaceae** Engl. & E. Gilg 1924

*Geosiphon* F. Wettst. 1915

**Class Glomeromycetes** Caval.-Sm. 1998

**Order Claroideoglomerales** H. Zhao, Y.C. Dai, B.K. Cui, F. Wu & X.Y. Liu, *ord. nov*.

**Family Claroideoglomeraceae** C. Walker & A. Schüßler 2010

*Claroideoglomus* C. Walker & A. Schüßler 2010

**Order Diversisporales** C. Walker & A. Schüßler 2004

**Family Acaulosporaceae** J.B. Morton & Benny 1990

*Acaulospora* Gerd. & Trappe 1974

*Kuklospora* Oehl & Sieverd. 2006

*Tricispora* Oehl, Sieverd., G.A. Silva & Palenz. 2011

**Family Diversisporaceae** C. Walker & A. Schüßler 2004

*Corymbiglomus* Błaszk. & Chwat 2012

*Desertispora* Błaszk., Kozłowska, Ryszka, Al-Yahya’ei & Symanczik 2018

*Diversispora* C. Walker & A. Schüßler 2004

*Otospora* Oehl, Palenz. & N. Ferrol 2008

*Redeckera* C. Walker & A. Schüßler 2010

*Sieverdingia* Błaszk., Niezgoda & B.T. Goto 2019

**Family Entrophosporaceae** Oehl & Sieverd. 2006

*Entrophospora* R.N. Ames & R.W. Schneid. 1979

**Family Pacisporaceae** C. Walker, Błaszk., A. Schüßler & Schwarzott 2004

*Pacispora* Sieverd. & Oehl 2004

**Family Sacculosporaceae** Oehl, Sieverd., G.A. Silva, B.T. Goto, Sánchez-Castro & Palenz.

*Sacculospora* Oehl, Sieverd., G.A. Silva, B.T. Goto, Sánchez-Castro & Palenz. 2011

**Order Gigasporales** S.P. Gautam & U.S. Patel 2007

**Family Dentiscutataceae** Sieverd., F.A. Souza & Oehl 2009

*Dentiscutata* Sieverd., F.A. Souza & Oehl 2009

**Family Gigasporaceae** J.B. Morton & Benny 1990

*Gigaspora* Gerd. & Trappe 1974

*Intraornatospora* B.T. Goto, Oehl & G.A. Silva 2012

*Paradentiscutata* B.T. Goto, Oehl & G.A. Silva 2012

**Family Racocetraceae** Oehl, Sieverd. & F.A. Souza 2009

*Cetraspora* Oehl, F.A. Souza & Sieverd. 2009

*Racocetra* Oehl, F.A. Souza & Sieverd. 2009

**Family Scutellosporaceae** Sieverd., F.A. Souza & Oehl 2009

*Bulbospora* Oehl & G.A. Silva 2014

*Scutellospora* C. Walker & F.E. Sanders 1986

**Order Glomerales** J.B. Morton & Benny 1990

**Family Glomeraceae** Piroz. & Dalpé 1989

*Dominikia* Błaszk., Chwat & Kovács 2014

*Epigeocarpum* Błaszk., B.T. Goto, Jobim, Niezgoda & Marguno 2021

*Funneliformis* C. Walker & A. Schüßler 2010

*Funneliglomus* Corazon-Guivin, G.A. Silva & Oehl 2019

*Glomus* Tul. & C. Tul. 1844

*Halonatospora* Błaszkowski, Niezgoda, B.T. Goto & Kozłowska 2018

*Kamienskia* Błaszk., Chwat & Kovács 2014

*Microdominikia* Oehl, Corazon-Guivin & G.A. Silva 2019

*Microkamienskia* Corazon-Guivin, G.A. Silva & Oehl 2019

*Nanoglomus* Corazon-Guivin, G.A. Silva & Oehl 2019

*Oehlia* Błaszk., Kozłowska, Niezgoda, B.T. Goto & Dalpé 2018

*Rhizophagus* P.A. Dang. 1896

*Sclerocarpum* B.T. Goto, Błaszk., Niezgoda, A. Kozłowska & Jobim 2019

*Silvaspora* Błaszk., Niezgoda, B.T. Goto, Crossay & Magurno 2021

**Class Paraglomeromycetes** Oehl, G.A. Silva, B.T. Goto & Sieverd. 2011

**Order Paraglomerales** C. Walker & A. Schüßler 2001

**Family Paraglomeraceae** J.B. Morton & D. Redecker 2001

*Innospora* Błaszk., Kovács, Chwat & Kozłowska 2017

*Paraglomus* J.B. Morton & D. Redecker 2001

**Family Pervetustaceae** Błaszk., Chwat, Kozłowska, Symanczik & Al-Yahya’ei 2017

*Pervetustus* Błaszk., Chwat, Kozłowska, Symanczik & Al-Yahya’ei 2017

**Phylum Mortierellomycota** Tedersoo, Sánchez-Ramírez, Kõljalg, Bahram, Döring, Schigel, T. May, M. Ryberg & Abarenkov 2018

**Class Mortierellomycetes** Doweld 2013

**Order Mortierellales** Caval.-Sm. 1998

**Family Mortierellaceae** Luerss. 1877

*Actinomortierella* Chalab 1968

*Aquamortierella* Embree & Indoh 1967

*Benniella* Vandepol & Bonito 2020

*Dissophora* Thaxter 1914

*Entomortierella* Vandepol & Bonito 2020

*Gamsiella* Benny & M. Blackwell 2004

*Gryganskiella* Vandepol, Stajich & Bonito 2020

*Linnemannia* Vandepol & Bonito 2020

*Lobosporangium* M. Blackwell & Benny 2004

*Lunasporangiospora* Vandepol & Bonito 2020

*Modicella* Kanouse 1936

*Mortierella* Coemans 1863

*Necromortierella* Vandepol & Bonito 2020

*Podila* Stajich, Vandepol & Bonito 2020

**Phylum Mucoromycota** Doweld 2001

**Class Mucoromycetes** Doweld 2001

**Order Cunninghamellales** H. Zhao, Y.C. Dai, B.K. Cui, F. Wu & X.Y. Liu, *ord. nov*.

**Family Absidiaceae** Arx 1982

*Absidia* Tiegh. 1878

*Chlamydoabsidia* Hesselt. & J.J. Ellis 1966

*Halteromyces* Shipton & Schipper 1975

**Family Cunninghamellaceae** Naumov ex R.K. Benj. 1959

*Cunninghamella* Matr. 1903

**Family Gongronellaceae** H. Zhao, Y.C. Dai, B.K. Cui, F. Wu & X.Y. Liu, *fam. nov*. *Gongronella* Ribaldi 1952

*Hesseltinella* H.P. Upadhyay 1970

**Order Lentamycetales** H. Zhao, Y.C. Dai, B.K. Cui, F. Wu & X.Y. Liu, *ord. nov*.

**Family Lentamycetaceae** K. Voigt & P.M. Kirk 2012

*Lentamyces* Kerst. Hoffm. & K. Voigt 2008

**Order Mucorales** Dumort 1829

**Family Backusellaceae** K. Voigt & P.M. Kirk 2012

*Backusella* Hesselt. & J.J. Ellis 1969

**Family Mucoraceae** Fr. 1821

*Actinomucor* Schostak. 1898

*Ambomucor* R.Y. Zheng & X.Y. Liu 2014

*Benjaminiella* Arx 1981

*Blakeslea* Thaxt. 1914

*Chaetocladium* Fresen. 1863

*Choanephora* Curr. 1873

*Cokeromyces* Shanor 1950

*Dicranophora* J. Schröt. 1886

*Ellisomyces* Benny & R.K. Benj. 1975

*Gilbertella* Hesselt. 1960

*Helicostylum* Corda 1842

*Hyphomucor* Schipper & Lunn 1986

*Isomucor* J.I. Souza, Pires-Zottar. & Harakava 2012

*Kirkiana* L.S. Loh, Kuthub. & Nawawi 2001

*Kirkomyces* Benny 1996

*Mucor* Fresen. 1850

*Mycotypha* Fenner 1932

*Nawawiella* L.S. Loh & Kuthub. 2001

*Parasitella* Bainier 1903

*Pilaira* Tiegh. 1875

*Pirella* Bainier 1882

*Poitrasia* P.M. Kirk 1984

*Rhizopodopsis* Boedijn 1959

*Thamnidium* Link 1809

*Tortumyces* L.S. Loh 2001

**Family Pilobolaceae** Corda 1842

*Pilobolus* Tode 1784

*Utharomyces* Boedijn ex P.M. Kirk & Benny 1980

**Family Rhizopodaceae** K. Schum. 1894

*Rhizopus* Ehrenb. 1821

**Family Syzygitaceae** H. Zhao, Y.C. Dai, B.K. Cui, F. Wu & X.Y. Liu, *fam. nov*.

*Sporodiniella* Boedijn 1959

*Syzygites* Ehrenb. 1818

**Order Phycomycetales** H. Zhao, Y.C. Dai, B.K. Cui, F. Wu & X.Y. Liu, *ord. nov*.

**Family Phycomycetaceae** Arx 1982

*Phycomyces* Kunze 1823

*Spinellus* Tiegh. 1875

**Family Radiomycetaceae** Hesselt. & J.J. Ellis 1974

*Radiomyces* Embree 1959

**Family Saksenaeaceae** Hesselt. & J.J. Ellis 1974

*Apophysomyces* P.C. Misra 1979

*Saksenaea* S.B. Saksena 1953

**Order Syncephalastrales** H. Zhao, Y.C. Dai, B.K. Cui, F. Wu & X.Y. Liu, *ord. nov*.

**Family Circinellaceae** H. Zhao, Y.C. Dai, B.K. Cui, F. Wu & X.Y. Liu, *fam. nov*.

*Circinella* Tiegh. & G. Le Monn. 1873

*Fennellomyces* Benny & R.K. Benj. 1975

*Phascolomyces* Boedijn ex Benny & R.K. Benj. 1976

*Thamnostylum* Arx & H.P. Upadhyay 1970

*Zychaea* Benny & R.K. Benj. 1975

**Family Lichtheimiaceae** Kerst. Hoffm., Walther & K. Voigt 2009

*Dichotomocladium* Benny & R.K. Benj. 1975

*Lichtheimia* Vuill. 1903

**Family Mycocladaceae** Kerst. Hoffm., Discher & K. Voigt 2007

*Mycocladus* Beauverie 1900

**Family Protomycocladaceae** H. Zhao, Y.C. Dai, B.K. Cui, F. Wu & X.Y. Liu, *fam. nov*.

*Protomycocladus* Schipper & Samson 1994

**Family Rhizomucoraceae** H. Zhao, Y.C. Dai, B.K. Cui, F. Wu & X.Y. Liu, *fam. nov*.

*Rhizomucor* Lucet & Costantin 1900

**Family Thermomucoraceae** H. Zhao, Y.C. Dai, B.K. Cui, F. Wu & X.Y. Liu, *fam. nov*.

*Thermomucor* Subrahm., B.S. Mehrotra & Thirum. 1977

**Family Syncephalastraceae** Naumov ex R.K. Benj. 1959

*Syncephalastrum* J. Schröt. 1886

Phylum Umbelopsidomycota H. Zhao, Y.C. Dai, B.K. Cui, F. Wu & X.Y. Liu, *phyl. nov*.

**Class Umbelopsidomycetes** Tedersoo, Sánchez-Ramírez, Kõljalg, Bahram, Döring, Schigel, T. May, M. Ryberg & Abarenkov 2018

**Order Umbelopsidales** Spatafora, Stajich & Bonito 2016

**Family Pygmaeomycetaceae** E. Walsh & N. Zhang 2020

*Pygmaeomyces* E. Walsh & N. Zhang 2020

**Family Umbelopsidaceae** W. Gams & W. Mey. 2003

*Umbelopsis* Amos & H.L. Barnett 1966

### Notes of genera in Mortierellomycota, Mucoromycota and Umbelopsidomycota

In this study, we describe 73 new species, propose three new combinations and present 43 Chinese new record species. These 119 species belong to Mortierellomycota, Mucoromycota and Umbelopsidomycota. These three phyla accommodate 71 genera in total, and therefore we provide herein notes for all these genera.

1. ***Absidia*** Tiegh., *Annls Sci. Nat.*, Bot., sér. 6 4(4): 350, 1878 *Absidia* is the type of the family Absidiaceae and typified by *A. reflexa* Tiegh. (van Tieghem 1876). Members of the genus always produce sporangiophores from stolons, form rhizoids between sporangiophores, and develop sporangia which are pyriform, with deliquescent walls, and apophysate (Benny *et al*. 2001). Columellae are mainly conical, subglobose and applanate, commonly with apical projections (van Tieghem 1876, Hoffmann *et al*. 2007). Some species produce zygospores within zygosporangia, and their opposite suspensory cells possess appendages (Hoffmann *et al*. 2007, Hoffmann 2010). Currently, it contains 51 species (Zhao *et al*. 2022b). This genus is distributed worldwide in soil (Rafhaella *et al*. 2020), herbivorous dung and rotten plants (van Tieghem 1876), and particularly in the mycangia of ambrosia beetles (*A. psychrophilia* Hesselt. & J.J. Ellis; Hesseltine & Ellis 1964, Kaitera *et al*. 2019) as well as the body surface of bats (*A. stercoraria* Hyang B. Lee, H.S. Lee & T.T.T. Nguyen; https://bccm.belspo.be/content/remarkable-fungal-biodiversity-northern-belgium-bats). Some species of the genus are industrially relevant. The α-galactosidase from *A. reflexa* was used to yield rubusoside derivatives (Kitahata *et al*. 1989). Laccase from *A. spinosa* Lendn. is used for the biotransformation of cresol red (Kristanti *et al*. 2016). *A. repens* Tiegh. yields chitosan (Davoust & Persson 1992). In total, 15 new taxa are described herein.
2. ***Actinomortierella*** Chalab., *Nov. sist. Niz. Rast*., 1968 (5): 129, 1968 *Actinomortierella* is the type of the family Actinomortierellaceae and typified by *A. capitata* (Marchal) Vandepol & Bonito (Vandepol *et al*. 2020). It was once treated as a section in the genus *Mortierella* by Gams (1977), but resurrected as a separate genus by Vandepol *et al*. (2020). Short branches grow on the apical inflation of sporangiophores. Terminal sporangia form on both stem and lateral sporangiophores. Chlamydospores are absent (Vandepol *et al*. 2020). Currently, it contains three species *A. ambigua* (B.S. Mehrotra) Vandepol & Bonito, *A. capitata* and *A. wolfii* (B.S. Mehrotra & Baijal) Van-depol & Bonito. Members are distributed in soil (Mehrotra *et al*. 1963). Cultures have been recorded in Fiji, India, New Zealand, North America, and the UK. *A. wolfii* is a well-known pathogen of animals (Munday *et al*. 2006, 2010, Davies *et al*. 2010), but also reported to be associated with human invasive diseases (Layios *et al*. 2014). However, another species, *A. capitata*, is useful as being able to promote crop growth (Li *et al*. 2020). This paper newly records this genus in Yunnan, China.
3. ***Actinomucor*** Schostak., *Ber. Dt. Bot. Ges*. 16: 155, 1898 *Actinomucor* is typified by *A. elegans* (Eidam) C.R. Benj. & Hesselt. (Benjamin & Hesseltine 1957). The genus is closely related to *Mucor*, but differs in sporangiophores, stolons and rhizoids (Benjamin & Hesseltine 1957). Currently, only the type species is accepted in the genus (Zheng & Liu 2005). This species is a pathogen of mucormycosis (Davel *et al*. 2001, Tully *et al*. 2009, Mahmud *et al*. 2012, Trobisch *et al*. 2020). It produces glutaminase (Lu *et al*. 1996) and is widely used to ferment soybean food (Han *et al*. 2001), which in turn benefits human health (Yin *et al*. 2021). It also secretes proteases and lipases for biodegrading polylactic acid / polybutylene adipate-coterephthalate (Jia *et al*. 2021).
4. ***Ambomucor*** R.Y. Zheng & X.Y. Liu, *Mycotaxon* 126: 99, 2014 *Ambomucor* is typified by *A. seriatoinflatus* X.Y. Liu & R.Y. Zheng and is characterized by simultaneously having two kinds of sporangia, fertile and aborted. Currently, it contains three species, *A. clavatus* X.Y. Liu & R.Y. Zheng, *A. ovalosporus* R.Y. Zheng & X.Y. Liu and *A. seriatoinflatus* (Zheng & Liu 2014, Liu & Zheng 2015). All these three species are reported from China in soil (Romero-Olivares *et al*. 2019). *A. seriatoinflatus* is found to synthesize sialidases which are widely applied in food and pharma industries (Abrashev *et al*. 2021).
5. ***Apophysomyces*** P.C. Misra, *Mycotaxon* 8(2): 377, 1979 *Apophysomyces* is typified by *A. elegans* P.C. Misra, K.J. Srivast. & Lat and is characterized by pyriform sporangia, and conspicuous funnel- or bell-shaped apophyses (Misra & Lata 1979, Khuna *et al*. 2019, Li *et al*. 2020). Currently, it contains six species (Misra & Lata 1979, Li *et al*. 2020). All species of *Apophysomyces* are found in soil, decayed vegetation and detritus. At least four species, *A. elegans*, *A. mexicanus* A. Bonifaz *et al*., *A. ossiforms* E. Álvarez *et al*. and *A. variabilis* E. Álvarez *et al*., are reported to cause mucormycosis (Chakrabarti *et al*. 2010, Guarro *et al*. 2011, Bonifaz *et al*. 2014, Khuna *et al*. 2019, Martínez-Herrera *et al*. 2020). *Apophysomyces* species are the second common agents of mucormycosis in India (Chander *et al*. 2021).
6. ***Aquamortierella*** Embree & Indoh 1967 *Aquamortierella* is typified by *A. elegans* Embree & Indoh (Misra & Lata 1979). The genus is characterized by sporangiospores discharging under water, and reniform (kidney-shaped) to allantoid (sausage-shaped) sporangiospores (Embree & Indoh 1967). Currently, it includes the type species only (www.indexfungorum.org; accessed 5 August 2021), but no living culture is available. This species is isolated from midge larvae in a freshwater stream in New Zealand (Embree & Indoh 1967).
7. ***Backusella*** Hesselt. & J.J. Ellis, *Mycologia* 61: 863, 1969 *Backusella* is the type of the family Backusellaceae and typified by *B. circina* J.J. Ellis & Hesselt. (Ellis & Hesseltine 1969). The genus is characterized by many sporangiola-bearing lateral branches on the stem of the sporangiophores which in turn produce multi-spored sporangia (Ellis & Hesseltine 1969). In recent years, the species number of *Backusella* has been dramatically increasing, especially from Australia and Korea (Nguyen *et al*. 2021, Urquhart *et al*. 2021). Currently, 29 species are recorded in Index Fungorum (www.indexfungorum.org; accessed 5 August 2021). However, only two species, *B. circina* and *B. lamprospora* (Lendn.) Benny & R.K. Benj., are recorded in China (Zheng *et al*. 2013). Species of the genus inhabit mainly in soil, litter and herbivore dung. Two species, *B. circina* and *B. lamprospora*, are recorded as agents of mucormycosis (Zheng *et al*. 2013, Panthee *et al*. 2021). *B. lamprospora* has capacity to produce beta-carotene (Papp *et al*. 2009) and is also a potential oleaginous fungus (Zhao *et al*. 2021a). In this paper, we present three new species and four new Chinese records.
8. ***Benniella*** Vandepol & Bonito, in Vandepol *et al*., *Fungal Diversity* 104: 281, 2020 *Benniella* is typified by *B. erionia* Liber & Bonito (Vandepol *et al*. 2020). It was erected in honor of Gerald Benny’s significant contributions on Mortierellaceae (Vandepol *et al*. 2020). Aerial hyphae are abundant in this genus, near the agar surface, producing slight rings with age (Vandepol *et al*. 2020). Currently, it contains the type species only (www.indexfungorum.org; accessed 5 August 2021).
9. ***Benjaminiella*** Arx, *Gen. Fungi Sporul. Cult*., Edn 3 (Vaduz): 60, 1981 *Benjaminiella* is typified by *B. poitrasii* (R.K. Benj.) Arx (basionym *Cokeromyces poitrasii* R.K. Benj.; von Arx 1970). The genus produces sporangiola on relatively long, unbranched, curved to twisted pedicels, which in turn form on the entire surface of fertile vesicles (von Arx 1970, Kirk 1989, Benny *et al*. 1985). The fertile vesicles are produced at the terminal of the sporangiophores (von Arx 1970, Kirk 1989, Benny *et al*. 1985). The pedicels and sporangiola are released as a unit when dehiscence occurs at a circumscissile zone, leaving denticles regularly scattered on the surface of the vesicles (von Arx 1970, Kirk 1989, Benny *et al*. 1985). Zygosporangia are homothallic, pigmented and ornamented (von Arx 1970, Benny *et al*. 1985, Kirk 1989). Currently, the genus includes three species which are commonly isolated from soil and rat dung, and sometimes are pathogens of humans (Gade *et al*. 2017). *B. poitrasii* is a well-known model of a dimorphic fungus and contains a large number of chitin synthase (Khale & Deshpande 1992, Chitnis *et al*. 2002).
10. ***Blakeslea*** Thaxt., *Bot. Gaz*. 58: 353, 1914 *Blakeslea* is typified by *B. trispora* Thaxt. (Thaxter 1914). The genus is characterized by two types of sporangia on separate sporangiophores (Kirk 1984, Zheng & Chen 1986). Three species are currently accepted in the genus (www.indexfungorum.org; accessed 5 August 2021). They are commonly collected from soil and fallen flowers of cucurbitaceous or malvaceous plants (Ho & Chang 2003). *B. trispora* is used industrially for producing carotenoids (Luo *et al*. 2020).
11. ***Chaetocladium*** Fresen., *Beitr. Mykol.* 3: 97, 1863 *Chaetocladium* is typified by *C. jonesiae* (Berk. & Broome) Fresen. (Benny & Benjamin 1976). In *Chaetocladium*, uni-spored sporangiola have a separable sporangial wall (Benny & Benjamin 1976, Benny *et al*. 2001). The sporangiola are borne on pedicels which grow on vesicles (Benny & Benjamin 1976, Benny *et al*. 2001). Fertile hyphae produce two or three branches from one point, and branches usually end in a sterile spine (Benny & Benjamin 1976, Benny *et al*. 2001). Zygospores are formed between opposite suspensors, which are more or less equal (Benny & Benjamin 1976, Benny *et al*. 2001). Two species, *C. brefeldii* Tiegh. & G. Le Monn. and *C. jonesiae*, are accepted (www.indexfungorum.org; accessed 5 August 2021). Species of *Chaetocladium* are facultative parasites on other mucoralean fungi (Benny & Benjamin 1976, Benny *et al*. 2001).
12. ***Chlamydoabsidia*** Hesselt. & J.J. Ellis, *Mycologia* 58(5): 761, 1966 *Chlamydoabsidia* is typified by *C. padenii* Hesselt. & J.J. Ellis (Hesseltine & Ellis 1966). The genus differs from *Absidia* by large, pigmented, fusiform, multiseptated chlamydospores formed on aerial hyphae (Hesseltine & Ellis 1966). It includes the type species only (www.indexfungorum.org; accessed 5 August 2021).
13. ***Choanephora*** Curr., *J. Linn. Soc., Bot*. 13: 578, 1873 *Choanephora* replaced the synonym *Cunninghamia* and is typified by *Choanephora infundibulifera* (Curr.) D.D. Cunn. (basionym *Cunninghamia infundibulifera* Curr.; Kirk 1984). The genus produces multi-spored sporangia and uni-spored sporangiola at separate sporangiophores (Kirk 1984). In contrast to those in the genus *Blakeslea*, sporangiola in *Choanephora* are uni-spored, forming on fertile heads and lacking suture in the sporangial wall (Kirk 1984). There are no really separable walls in the uni-spored sporangiola (Kirk 1984). Two species, *C. cucurbitarum* and *C. infundibulifera*, are accepted in *Choanephora* so far (www.indexfungorum.org; accessed 5 August 2021). They cause a blossom blight in gourd, pumpkin, pepper, okra, and other cucubits. (Agrios 2005, Saroj *et al*. 2012).
14. ***Circinella*** Tiegh. & G. Le Monn., *Annls Sci. Nat., Bot., sér.* 5 17: 298, 1873 *Circinella* is the type of the family Circinellaceae and typified by *C. umbellata* Tiegh. & G. Le Monn. (Hesseltine & Fennell 1955). The genus is characterized by sporangiophores bearing circinate branches that terminate in globose apophysate persistent-walled sporangia (Hesseltine & Fennell 1955). Nowadays, nine species are accepted in this genus (www.indexfungorum.org; accessed 5 August 2021), and two of them were described from China (Zheng *et al*. 2017). Species of the genus can be used as microbial biosensors, and in the biotransformation of progesterone and testosterone (Alpat *et al*. 2008, Zoghi *et al*. 2019). In this paper, we present one new species from China.
15. ***Cokeromyces*** Shanor, *Mycologia* 42(2): 272, 1950 *Cokeromyces* is typified by *C. recurvatus* Poitras and is characterized by pedicellate sporangiola originating from a globoid enlargement at the hyphal terminal (Shanor *et al*. 1950). Pedicles are elongate and non-deciduous (Shanor *et al*. 1950, Benny & Benjamin 1976, Kirk 1984). Sporangiola are columellate, multi-spored, globose, persistent-walled, smooth (Shanor *et al*. 1950, Benny & Benjamin 1976, Kirk 1984). Sporangiospores are variable in shape, globose, ovoid, or irregular (Shanor *et al*. 1950, Benny & Benjamin 1976, Kirk 1984). Zygospores are globose to subglobose, with dark walls and opposed suspensors, homothallic (Shanor *et al*. 1950, Benny & Benjamin 1976, Kirk 1984). The sole species, *C. recurvatus*, is accepted in the genus, and it causes mucormycosis (Tsai *et al*. 1997).
16. ***Cunninghamella*** Matr., *Annls Mycol.* 1(1): 46, 1903 *Cunninghamella* is the type of the order Cunninghamellales and the family Cunninghamellaceae and typified by *C. africana* Matr. (Zheng & Chen 2001). The genus is characterized by terminal vesicles bearing uni-spored sporangiola, which are surrounded by spines (Zheng & Chen 2001). A worldwide monographic on a combination of morphology, maximum growth temperature, mating compatibility and molecular systematics was conducted and consequently 15 species/varieties were accepted (Zheng & Chen 2001). Subsequently, four species, *C. bigelovii* Z.H. Xin, Y. Hui Zhao & Hui Wang, *C. gigacellularis* A.L. Santiago, C.L. Lima & C.A.F. de Souza, *C. guizhouensis* Zhi.Y. Zhang, Y.F. Han & Z.Q. Liang and *C. globospora* H. Zhao & X.Y. Liu, were described in the genus (Guo *et al*. 2015, Hyde *et al*. 2016, Zhang *et al*. 2020, Zhao *et al*. 2021b). Species of *Cunninghamella* are found in soil, plant material, animal material and cheese. Some members of the genus are endophytic (Zhao *et al*. 2014). Fatty acids in the genus are dominantly composed of oleic acid (Zhao *et al*. 2014). In addition to being a common contaminant, *Cunninghamella* spp. are agents of zygomycosis (Robinson *et al*. 1990). Conversely, *Cunninghamella* spp. are also a microbial model for drug metabolism/biotransformation (Sepuri & Vidyavathi 2008). In this paper, six new species are described, and one new combination is proposed within the genus.
17. ***Dichotomocladium*** Benny & R.K. Benj., *Aliso* 8(3): 338, 1975 *Dichotomocladium* is typified by *D. elegans* Benny & R.K. Benj. (Benny & Benjamin 1975). Based on molecular data, the genus was once treated as a member of Lichtheimiaceae, closely related to *Lichtheimia* (Hoffmann *et al*. 2009b). It is distinguished from *Lichtheimia* by dichotomously branched fertile hyphae, sporangiola, sterile spines and absence of giant cells (Benny & Benjamin 1975, 1993, Hoffmann *et al*. 2009b). It contains five species (Hoffmann *et al*. 2009b) which are coprophilous and rarely found (Benny & Benjamin 1975, 1993).
18. ***Dicranophora*** J. Schröt., *Jber. Schles. Ges. Vaterl. Kultur* 64: 184, 1886 *Dicranophora* is typified by *D. fulva* J. Schröt. and is characterized by two types of sporangia, tree-like sporangiospores, yellow colonies, pronounced anisogamy, zygospores with irregular creviced, homothallism (Voglmayr & Krisai-Greilhuber 1996). Currently, it contains the type species only (Voglmayr & Krisai-Greilhuber 1996) which grows on rotting boletes in Austria (Voglmayr & Krisai-Greilhuber 1996).
19. ***Dissophora*** Thaxt., *Bot. Gaz.* 58: 361, 1914 *Dissophora* is typified by *D. decumbens* Thaxt. and is characterized by fertile hyphae abruptly differentiated from slender creeping vegetative hyphae, and by sporangiophores continuously arising as buds from fertile hyphae (Thaxter 1914). The uni- or two-spored sporangiola are similar to those in *Mortierella* (Thaxter 1914). Currently, it includes four species (www.indexfungorum.org; accessed 5 August 2021). Most species of the genus are found in soil except *D. decumbens* which was originally isolated from wood mouse (*Apodemus sylvaticus*) dung (Thaxter 1914).
20. ***Ellisomyces*** Benny & R.K. Benj., *Aliso* 8(3): 330, 1975 *Ellisomyces* is typified by *E. anomalus* (Hesselt. & P. Anderson) Benny & R.K. Benj.; basionym is *Thamnidium anomalum* Hesselt. & P. Anderson (Hesseltine & Anderson 1956). Sporangiophores arise from the substrate hyphae, producing bifurcate or trifurcate apical branches with multi-spored, columellate, persistent-walled, and globose sporangiola (Benny & Benjamin 1975). Zygospores are surrounded by an undulate wall and produced between opposed, approximately equal suspensors (Benny & Benjamin 1975). Currently, it includes the type species only (Benny & Benjamin 1975).
21. ***Entomortierella*** Vandepol & Bonito, in Vandepol *et al*., *Fungal Diversity* 104: 281, 2020 *Entomortierella* is typified by *E. lignicola* (G.W. Martin) Vandepol & Bonito (basionym *Haplosporangium lignicola* G.W. Martin) and is characterized by globose, smooth or spiny sporangiospores, as well as spiny chlamydospores in some species (Vandepol *et al*. 2020). Currently, it includes five species (Vandepol *et al*. 2020). These species are usually associated with insects and widely distributed in soil, roots and rotten plants (Vandepol *et al*. 2020). They are also associated with soil bacteria (Telagathoti *et al*. 2021).
22. ***Fennellomyces*** Benny & R.K. Benj., *Aliso* 8(3): 328, 1975 *Fennellomyces* is typified by *F. linderi* (Hesselt. & Fennell) Benny & R.K. Benj. (basionym *Circinella linderi* Hesselt. & Fennell) and is characterized by swellings immediately below a large terminal sporangium (Benny & Benjamin 1975). Four species are accepted (www.indexfungorum.org; accessed 5 August 2021), and they are found in rat dung and poplin fabric (Benny & Benjamin 1975).
23. ***Gamsiella*** (R.K. Benj.) Benny & M. Blackw., *Mycologia* 96(1): 147, 2004 *Gamsiella* typified by *G. multidivaricata* (R.K. Benj.) Benny & M. Blackw. was previously treated as a subgenus in *Mortierella* (Gams 1976, 1977, Benny & Blackwell 2004). The genus is characterized by repeatedly divaricately branched sporangiophores, two-spored sporangiola on long slender pedicels, and terminal globose ornamented chlamydospores (Benjamin 1978, Benny & Blackwell 2004). Currently, it includes two species (Vandepol *et al*. 2020). Previously, strains of the genus were isolated from detritus taken from a rotten stump (Benjamin 1978). In this paper, we describe a new species of *Gamsiella*, which is found in soil.
24. ***Gilbertella*** Hesselt., *Bull. Torrey bot. Club* 87: 24, 1960 *Gilbertella* is typified by *G. persicaria* (E.D. Eddy) Hesselt. (basionym *Choanephora persicaria* E.D. Eddy) and is characterized by sporangia with a longitudinal suture on their persistent walls (Hesseltine 1960, Benny 1991). Currently, it contains the type species only. This species is a potential new source of oleic acid (Shah *et al*. 2021) and also a post-harvest soft rot pathogen of fruits and vegetables (Guo *et al*. 2012, Pinho *et al*. 2014, Cruz-Lachica *et al*. 2015). Most recently, it has been isolated from human stool (Huët *et al*. 2020).
25. ***Gongronella*** Ribaldi, *Riv. Biol*. 44: 164, 1952 *Gongronella* is the type of the family Gongronellaceae and typified by *G. urceolifera* (Ribaldi 1952, Hesseltine & Ellis 1964). The characteristic of *Gongronella* is a distinct swollen apophysis (Ribaldi 1952, Hesseltine & Ellis 1964, Doilom *et al*. 2020). A total of twelve species are accepted in the genus (Hesseltine & Ellis 1964, Doilom *et al*. 2020). *Gongronella butleri* (Lendn.) Peyronel & Dal Vesco is well-known in chitosan industry (Tan *et al*. 1996, Babu *et al*. 2015). *Gongronella* spp. were found to be among phosphate-solubilizing (Doilom *et al*. 2020) and metalaxyl degrading fungi (Martins *et al*. 2020). A species of *Gongronella* was found capable of inducing overproduction of laccase in *Coprinopsis cinerea* (Schaeff.) Redhead, Vilgalys & Moncalvo (Hu *et al*. 2019) and *Panus rudis* Fr. (Wei *et al*. 2010). In this paper, two new species are described from China.
26. ***Gryganskiella*** Vandepol & Bonito, in Vandepol *et al*., *Fungal Diversity* 104: 282, 2020 *Gryganskiella* is typified by *G. cystojenkinii* (W. Gams & Veenb.-Rijks) Vandepol & Bonito 2020 (basionym *Mortierella cystojenkinii* W. Gams & Veenb.-Rijks; Vandepol *et al*. 2020). The genus was established in honor of Andrii Gryanskyi, a Ukrainian-American mycologist who has made a great contribution to the Mucoromycota (Vandepol *et al*. 2020). Unique characteristics are not described for this genus, though some are really remarkable, including smooth, ellipsoid to cylindrical sporangiospores, and lightly pigmented, Light Brown, ochre or orange chlamydospores (Vandepol *et al*. 2020). Currently, it includes two species (Vandepol *et al*. 2020).
27. ***Halteromyces*** Shipton & Schipper, *Antonie van Leeuwenhoek* 41: 337, 1975 *Halteromyces* is typified by *H. radiatus* Shipton & Schipper and is characterized by dumbbell-shaped, apophysate, sporangia with deliquescent walls (Shipton & Schipper 1975). Currently, it contains the type species only (Shipton & Schipper 1975). Strains of the species were isolated from mud in a mangrove forest (Shipton & Schipper 1975).
28. ***Hesseltinella*** H.P. Upadhyay, *Persoonia* 6(1): 111, 1970 *Hesseltinella* is typified by *H. vesiculosa* H.P. Upadhyay and is characterized by sporangia which are multi-spored, spiny, on pedicels on versicles that originate from multi-vesiculate sporangiophores (Upadhyay 1970, Young 1987, Benny & Benjamin 1991). Currently, it contains the type species only (Upadhyay 1970). Strains of the species were isolated from soil samples (Upadhyay 1970, Young 1987).
29. ***Helicostylum*** Corda, *Icon. Fung.* (Prague) 5: 18, 1842 *Helicostylum* is typified by *H. elegans* Corda (Corda 1842). The main stem of the sporangiophores produces larger, multi-spored, sporangia with deliquescent walls, while the lateral branches bear smaller, persistent, and pedicellate sporangia, as well as verticillate or pseudo-verticillate borne spines (Corda 1842, Benny 1995b). Two species have been described in this genus (Benny 1995b). *H. pulchrum* (Preuss) Pidopl. & Milko is a psychrotolerant fungus, gowning on refrigerated meat (Spatafora *et al*. 2016).
30. ***Hyphomucor*** Schipper & Lunn, *Mycotaxon* 27: 83, 1986 *Hyphomucor* is typified by *H. assamensis* (B.S. Mehrotra & B.R. Mehrotra) Schipper & Lunn (basionym *Mucor assamensis* B.S. Mehrotra & B.R. Mehrotra; Schipper 1986a). Sporangiophores are sympodially branched. Sporangia are apophysate and subglobose (Schipper 1986a). Sporangial walls are dry and powdery (Schipper 1986a). Sporangiospores are subglobose (Schipper 1986a). Currently, it contains the type species only (Schipper 1986a).
31. ***Isomucor*** J.I. Souza, Pires-Zottar. & Harakava, *Mycologia* 104(1): 233, 2012 *Isomucor* is typified by *I. trufemiae* J.I. Souza, Pires-Zottar. & Harakava (de Souza *et al*. 2012). It was established based on molecular data and unique morphological traits (de Souza *et al*. 2012). The genus produces lateral branches bearing multi-spored sporangiola and sporangia as well as uniformly septate hyphae (de Souza *et al*. 2012). It contains two species, *I. fuscus* (Berk. & M.A. Curtis) J.I. Souza, Pires-Zottar. & Harakava, and *I. trufemiae* (de Souza *et al*. 2012). In this paper, *I. trufemia* is recorded from China for the first time.
32. ***Kirkiana*** L.S. Loh, Kuthub. & Nawawi, *Mucoraceous Fungi from Malaysia* (Kuala Lumpur): 76, 2001 *Kirkiana* is typified by *K. ramosa* L.S. Loh, Kuthub. & Nawawi (Loh *et al*. 2001). Currently, it contains the type species only (Loh *et al*. 2001). However, no cultures and sequences are available.
33. ***Kirkomyces*** Benny, *Mycologia* 87(6): 922, 1995 *Kirkomyces*, formerly named *Kirkia*, is typified by *K. cordense* (B.S. Mehrotra & B.R. Mehrotra) Benny (Benny 1995a, b). The genus is characterized by the absence of spine-like fertile branches and a higher optimal growth temperature than that of *Helicostylum* (Benny 1995a, b). Currently, it contains the type species only (Benny 1995a, b).
34. ***Lentamyces*** Kerst. Hoffm. & K. Voigt, *Pl. Biol*. 11(4): 550, 2009 *Lentamyces* is the type of the order Lentamycetales and the family Lentamycetaceae. It is typified by *L. parricidus* (Renner & Muskat ex Hesselt. & J.J. Ellis) Kerst. Hoffm. & K. Voigt (basionym *Absidia parricida* Renner & Muskat; Hoffmann & Voigt 2009). The genus is characterized by a slow growth and a maximum growth temperature < 30 °C (Hoffmann & Voigt 2009). It contains four species so far (www.indexfungorum.org; accessed 5 August 2021).
35. ***Lichtheimia*** Vuill., *Bull. Soc. Mycol. Fr*. 19: 126, 1903 *Lichtheimia* is the type of the family Lichtheimiaceae (Hoffmann & Voigt 2009, Hoffmann 2010). It is typified by *L. corymbifera* (Cohn) Vuill. (basionym *Mucor corymbifer* Cohn; Hoffmann & Voigt 2009, Hoffmann 2010). Most species of the genus formerly belonged to *Absidia* (Hoffmann & Voigt 2009, Hoffmann 2010). The genus is differentiated from *Absidia* by a maximum growth temperature of above 37 °C (Hoffmann & Voigt 2009, Hoffmann 2010). Currently, seven species are accepted (www.indexfungorum.org; accessed 5 August 2021). Interestingly, some species, such as *L. corymbifera* and *L. ramosea* (Zopf) Vuill., are well-known as pathogenic fungi causing mucormycosis, and are widely distributed in foods and the environment (Gomes *et al*. 2011, André *et al*. 2014, Schwartze & Jacobsen 2014). In this study, two new species are illustrated from Chinese soil samples.
36. ***Linnemannia*** Vandepol & Bonito, in Vandepol *et al., Fungal Diversity* 104: 282, 2020 *Linnemannia* is typified by *L. hyalina* (Harz) Vandepol & Bonito (basionym *Hydrophora hyalina* Harz; Vandepol *et al*. 2020). It was set up in honor of Germaine Linnemann for his contribution to the family Mortierellaceae (Vandepol *et al*. 2020). Most members of the genus usually produce ellipsoid, spherical to cylindrical sporangiospores (Vandepol *et al*. 2020). Chlamydospores are irregular, with various brown shades (Vandepol *et al*. 2020). Sexual reproduction is heterothallic (Vandepol *et al*. 2020). Currently, it includes eleven species (www.indexfungorum.org; accessed 5 August 2021).
37. ***Lobosporangium*** M. Blackw. & Benny, *Mycologia* 96(1): 144, 2004 *Lobosporangium* replaced the synonym *Echinosporangium* and is typified by *Lobosporangium transversale* (Malloch) M. Blackw. & Benny (basionym *Echinosporangium transversale* Malloch; Benny & Blackwell 2004). The genus is characterized by coenocytic and anastomosing mycelia, forming 3‒4 dichotomies at lateral branches, terminating with a cylindrical sporangium and several spines (Benny & Blackwell 2004). Currently, it contains the type species only (Benny & Blackwell 2004).
38. ***Lunasporangiospora*** Vandepol & Bonito, in Vandepol *et al*., *Fungal Diversity* 104: 283, 2020 *Lunasporangiospora* is typified by *L. chienii* (P.M. Kirk) Vandepol & Bonito (basionym *Mortierella chienii*) and is characterized by smooth and lunate sporangiospores (Vandepol *et al*. 2020). It accommodates two species, *L. chienii* and *L. selenospora* (W. Gams) Vandepol & Bonito (Vandepol *et al*. 2020). They were found in mushroom compost and forest soil (Vandepol *et al*. 2020).
39. ***Modicella*** Kanouse, *Mycologia* 28(1): 60, 1936 *Modicella* is typified by *M. malleola* (Harkn.) Gerd. & Trappe (basionym *Endogone malleola* Harkn.) and is characterized by truffle-like fruitbodies, namely sporocarps which consist of smooth, thin-walled and hyaline sporangia (Kanouse 1936). Phylogenetic analysis demonstrated that *Modicella* was monophyletic in Mortierellales (Smith *et al*. 2013). Apart from the type species, the other two species in the genus are *M. albostipitata* J.A. Cooper and *M. reniformis* (Bres.) Gerd. & Trappe (Cooper & Park 2020).
40. ***Mortierella*** Coem., *Bull. Acad. R. Sci. Belg., Cl. Sci., sér.* 2 15: 536, 1863 *Mortierella* is the type of the phylum Mortierellomycota, the class Mortierellomycetes, the order Mortierellales and the family Mortierellaceae (Tedersoo *et al*. 2018). It is typified by *M. polycephala* Coem. (Coemans 1863). The genus is characterized by sporangiophores enlarging at the base and shrinking sharply towards top (Coemans 1863). Some colonies are zonate and some look like rounded to slightly pointed rosette petals (Gams 1976, 1977). A few species even grow flat on the surface of the solid media (Gams 1977, Vandepol *et al*. 2020). Four types of spores are observed in this genus, including asexual sporangiospores, chlamydospores, stylospores and sexual zygospores, while one or more types of spores are absent in some members (Gams 1977, Vandepol *et al*. 2020). Currently, it encompasses more than 110 species, the largest genus in the phylum Mortierellomycota (www.indexfungorum.org; accessed 5 August 2021). Species of the genus are widely distributed in soils, plant debris, living plant roots, mosses and insect guts (Gams 1977, Domsch *et al*. 1980). *Mortierella* seems to be the most commonly isolated Mucoromyceta genus from soil and is highly associated with soil bacteria (Telagathoti *et al*. 2021). Some species of the genus accumulate large amounts of polyunsaturated fatty acids, especially arachidonic acid, attracting widespread interest (Zhao *et al*. 2021a). *Mortierella* species can be endophytic and plant growth promoting (Ozimek & Hanaka 2021).
41. ***Mucor*** Fresen., *Beitr. Mykol.* 1: 7, 1850 *Mucor* is the type of the subkingdom Mucoromyceta, the phylum Mucoromycota, the class Mucoromycetes, the order Mucorales and the family Mucoraceae (Tedersoo *et al*. 2018). It is a conserved genus with the explicitly conserved type *M. mucedo* Fresen. (Appendix III; Turland *et al*. 2018). The genus is characterized by fast growth, luxuriant aerial and substrate hyphae, usually unbranched sporangiophores, non-apophysate, zygospores with opposed suspensors, and are heterothallic or homothallic (Schipper 1978a). Species of *Mucor* are distributed all over the world, and commonly collected in soil and dung (Walther *et al*. 2013). It accommodates more than 100 species, the largest genus in the phylum Mucoromycota (www.indexfungorum.org; accessed 5 August 2021). In practice, *Mucor* seems to be the second most common genus of the Mucoromyceta isolated from soil. Some species are widely used in fermentation and biotransformation (Hong *et al*. 2012). Several species are used in the pharmaceutical industry for podophyllotoxin, kaempferol (Huang *et al*. 2014), and stearidonic acid (Khan *et al*. 2019), and some species are well-known pathogens causing mucormycosis (Chibucos *et al*. 2016, Panthee *et al*. 2021).
42. ***Mycocladus*** Beauverie, *Annls Univ. Lyon, Ser*. 2 3: 162, 1900 *Mycocladus* is the type of the family Mycocladaceae (Hoffmann *et al*. 2007). It is typified by *M. verticillatus* Beauverie and is characterized by relatively high optimum and maximum growth temperatures (37‒42 °C and 53 °C, respectively), abundantly giant cells, stolons and rhizoids, and apophysate sporangia with deliquescent walls (Hoffmann *et al*. 2007). Several species described in this genus, have been transferred to *Absidia* and *Lichtheimia* (Beauverie 1900, Hoffmann *et al*. 2007, Hoffmann *et al*. 2009b). Currently, it contains the type species only (Hoffmann *et al*. 2009b).
43. ***Mycotypha*** Fenner, *Mycologia* 24(2): 196, 1932 *Mycotypha* is typified by *M. microspora* Fenner and is characterized by cylindrical vesicles and two distinct kinds of denticles (Benny *et al*. 1985). Three species are recorded in the genus. They are distributed in Finland, India, Japan, USA and Zimbabwe (Benny & Benjamin 1976, Benny *et al*. 1985). *M. microspora* was found to induce the aggregation of firebrats (*Thermobia domestica*), a fast-moving brownish insect frequently living indoors in warm environments (Woodbury & Gries 2013). *M. africana* R.O. Novak & Backus and *M. microspora* are traditional dimorphic fungi (Schulz *et al*. 1974) and have been most recently reported as new pathogens causing mucormycosis (Trachuk *et al*. 2018).
44. ***Nawawiella*** L.S. Loh & Kuthub., *Mucoraceous Fungi from Malaysia* (Kuala Lumpur): 73, 2001 *Nawawiella* is typified by *N. apophysa* L.S. Loh & Kuthub. (Loh *et al*. 2001). It contains the type species only (Loh *et al*. 2001). However, no cultures and sequences are available for the species.
45. ***Necromortierella*** Vandepol & Bonito, in Vandepol *et al*., *Fungal Diversity* 104: 284, 2020 *Necromortierella* is typified by *N. dichotoma* (Linnem. ex W. Gams) Vandepol & Bonito (basionym *Mortierella dichotoma* Linnem. ex W. Gams) and is characterized by sporangiophores irregularly dichotomously branched and tapering towards the apex quickly, ellipsoid to cylindric sporangiospores, and elongated or irregular chlamydospores (Vandepol *et al*. 2020). Currently, it contains the type species only (Vandepol *et al*. 2020). This species has a necrotrophic and mycophilic life mode (Vandepol *et al*. 2020).
46. ***Parasitella*** Bainier, *Bull. Soc. Mycol. Fr*. 19(2): 153, 1903 *Parasitella* is typified by *P. parasitica* (Bainier) Syd. (basionym *Mucor parasiticus* Bainier) and is characterized by mycoparasitic behavior, sporangiophores from substrate hyphae *in vitro* and from aerial hyphae *in vivo*, subglobose to globose sporangia, apophysate, with a columella and moist wall, multi-spored, zygospores, and is heterothallic (Schipper 1978b). Currently, it contains the type species only (Schipper 1978b). The species is parasitic on other Mucoraceae such as *Absidia glauca* Hagem (Schipper 1978b).
47. ***Phascolomyces*** Boedijn ex Benny & R.K. Benj., *Aliso* 8(4): 417, 1976 *Phascolomyces* is typified by *P. articulosus* Boedijn ex Benny & R.K. Benj. (Benny & Benjamin 1976). The genus is allied to *Cunninghamella*, and characterized by chlamydospores which form on substrate hyphae, swollen vesicles on the side of sporangiophores, sporangiola which are pedicellate, globose to subglobose, columellate and uni-spored (Benny & Benjamin 1976). Currently, it contains the type species only (Benny & Benjamin 1976).
48. ***Phycomyces*** Kunze, *Mykologische Hefte* (Leipzig) 2: 113, 1823 *Phycomyces* is the type of the order Phycomycetales and the family Phycomycetaceae. It is typified by *P. nitens* (C. Agardh) Kunze (basionym *Ulva nitens* C. Agardh; Benjamin & Hesseltine 1959). The genus is characterized by sporangia which are non-apophysate, multi-spored and borne on exceedingly long, unbranched, bluish or metallic sporangiophores, zygospores which are *Mucor*-like and produce dichotomously branched, darkened spines and appendages (Benjamin & Hesseltine 1959, Bergman *et al*. 1969). The dried thalli in test tubes looks like a bunch of hair, well-known as a hairy fungus (Benjamin & Hesseltine 1959, Bergman *et al*. 1969). This genus accommodates three species, *P. blakesleeanus* Burgeff, *P. microspores* Tiegh. and *P. nitens* (C. Agardh) Kunze (Benjamin & Hesseltine 1959, Bergman *et al*. 1969). Among these species, *P. blakesleeanus* is well-known as a model organism for phototropism and light-growth response (Galland & Lipson 1985).
49. ***Pilobolus*** Tode, *Schr. Ges. Naturf. Freunde, Berlin* 5: 46, 1784 *Pilobolus* is the type of the family Pilobolaceae and typified by *P. crystallinus* (F.H. Wigg.) Tode (basionym *Hydrogera crystallina* F.H. Wigg.; www.indexfungorum.org; accessed 5 August 2021). The genus is characterized by trophocytes which give rise to positively phototropic sporangiophores, sporangia which are produced on a huge swelling and are violently discharged (Page 1962, Hu *et al*. 1989, Viriato 2008, Lee *et al*. 2018). It accommodates 15 species (www.indexfungorum.org; accessed 5 August 2021). Members of the genus are cosmopolitan and coprophilous and are also known as model organisms for the forcible ejection of entire ripe sporangia (Page 1964, Page & Kennedy 1964).
50. ***Pilaira*** Tiegh., *Annls Sci. Nat., Bot., sér.* 6 1: 51, 1875 *Pilaira* is typified by *P. cesatii* Milko (basionym *Ascophora cesatii* Coem., an illegitimate name) and is characterized by sporangiophores which are simple or branched, and sporangia which have cutinized walls, and are apophysate, columellate and multi-spored. It accommodates nine species (Zheng & Liu 2009, Liu *et al*. 2012, Urquhart *et al*. 2017).
51. ***Pirella*** Bainier, *Étud. Mucor.*, (Thèse, Paris) (Paris): 83, 1882 *Pirella* is typified by *P. circinans* Bainier and is characterized by sporangiophores which produce simultaneously terminal multi-spored sporangia and lateral several-spored sporangiola. Lateral branches are twisted and contorted (Benny & Schipper 1992). Zygospores are globose to compressed, with a pigmented and ornamented wall (Benny & Schipper 1992). It accommodates two species *P. circinans* and *P. naumovii* (Milko) Benny & Schipper (Benny & Schipper 1992). *P. circinans* has been isolated from China once (Liu 2004).
52. ***Podila*** Stajich, Vandepol & Bonito, in Vandepol *et al*., *Fungal Diversity* 104: 284, 2020 *Podila*, in honor of Gopi Podila, is typified by *P. minutissima* (Tiegh.) Vandepol & Bonito (basionym *Mortierella minutissima*; Vandepol *et al*. 2020). It seems there are no characteristics unique to this genus because species of the genus produce various types of sporangiospores, chlamydospores, and zygospores. Species are distributed in forest and agricultural soil, plant roots, compost, animal dung and municipal waste (Vandepol *et al*. 2020). It accommodates seven species all transferred from *Mortierella* (Vandepol *et al*. 2020).
53. ***Poitrasia*** P.M. Kirk, *Mycol. Pap*. 152: 51, 1984 *Poitrasia* is typified by *P. circinans* (H. Nagan. & N. Kawak.) P.M. Kirk (basionym *Blakeslea circinans*) and is characterized by tong-like suspensors (Kirk 1984). Currently, it contains the type species only (Kirk 1984) which was originally isolated from soil but also found to be a marine fungus with a phytotoxicity activity (Huang *et al*. 2018). It produces protease and keratinase (Agrawal *et al*. 2021).
54. ***Protomycocladus*** Schipper & Samson, *Mycotaxon* 50: 487, 1994 *Protomycocladus* is the type of the family Protomycocladaceae. It is typified by *P. faisalabadensis* (J.H. Mirza *et al*.) Schipper & Samson (basionym *Mucor faisalabadensis et al*.). The genus is characterized by the multi-spored sporangia that are apophysate, columellate and obpyriform (Mirza *et al*. 1979, Schipper & Samson 1994). Currently, it contains the type species only (Schipper & Samson 1994).
55. ***Pygmaeomyces*** E. Walsh & N. Zhang, *Mycologia* 113(1): 141, 2021 *Pygmaeomyces* is the type of the family Pygmaeomycetaceae and typified by *P. thomasii* E. Walsh & N. Zhang (Walsh *et al*. 2021). The genus is characterized by microchlamydospores which are hyaline, globose to subglobose, and distinguished from its most closely related genus *Umbelopsis* by producing colonies without pigmentation, sporangiophores and sporangiospores (Walsh *et al*. 2021). Currently, it contains two species which are associated with plant roots (Walsh *et al*. 2021).
56. ***Radiomyces*** Embree, *Am. J. Bot.* 46: 25, 1959 *Radiomyces* is the type of the family Radiomycetaceae and typified by *R. spectabilis* Embree (Embree 1959, Benny & Benjamin 1991). The genus is characterized by two levels of vesicles, the single main vesicle proliferating several secondary vesicles which in turn produce pedicles and uni- or several-spored sporangiola. Its zygospores are *Mucor*-like, except for simple or branched appendages (Embree 1959, Benny & Benjamin 1991). Three species are accepted in the genus (www.indexfungorum.org; accessed 5 August 2021), and they are coprophilous but are thermotolerant and pathogenic to mice (Kitz *et al*. 1980).
57. ***Rhizopodopsis*** Boedijn, *Sydowia* 12(1-6): 220, 1958 *Rhizopodopsis* is typified by *R. javensis* Boedijn and is characterized by sporangiophores which directly arise from a long erect hyphal branch, by sporangia which are apophysate, columellate and multi-spored, by sporangiospores which are globose to elongated, and by zygospores which are *Mucor*-like and homothallic (Boedijn 1958). Currently, it contains the type species only (Boedijn 1958).
58. ***Rhizomucor*** Lucet & Costantin, *Rev. Gén. Bot*. 12: 81, 1900 *Rhizomucor* is the type of the family Rhizomucoraceae and typified by *R. parasiticus* (replaced synonym *Mucor parasiticus* Lucet & Costantin, competing homonym non *M. parasiticus* Bainier; Schipper 1979). The genus differs from *Mucor* by rhizoids, and from *Thermomucor* by nonapohysate sporangia and rough zygospores (Subrahmanyam *et al*. 1977, Schipper 1979). It accommodates six species (Lucet and Costantin 1900, Zheng *et al*. 2009). *Rhizomucor* is a cosmopolitan genus and found in soil, decayed fruit and vegetables, fermenting and composting organic matter. All species in this genus produce lipase (Li *et al*. 2018), also cause mucormycosis in humans (Lukacs *et al*. 2004).
59. ***Rhizopus*** Ehrenb., *Nova Acta Phys.-Med. Acad. Caes. Leop.-Carol. Nat. Cur.* 10: 198, 1821 *Rhizopus* is type genus of the family Rhizopodaceae and typified by *R. nigricans* Ehrenb. which is an illegitimate name and synonymized with *R. stolonifer* (Ehrenb.) Vuill. (basionym *Mucor stolonifer* Ehrenb; Zheng *et al*. 2007). The genus is characterized by deep gray to blackish colonies, abundant rhizoids on hyphae and stolons, solitary or more often in a group, sporangia dark, round, apophysate and columellate, sporangiospores striped and of various shapes and sizes, zygospores or azygospores hyaline, globose to broadly ovoid (Zheng *et al*. 2007). It accommodates twelve species (Li *et al*. 2016, Zheng *et al*. 2007). Species of *Rhizopus* are distributed worldwide in soil and air. They play key roles in industrial, agricultural, and medical applications as fermentation agents, and some species cause diseases in plants, humans and animals (Zheng *et al*. 2007, Cheng *et al*. 2017, Yao *et al*. 2018, Ju *et al*. 2020).
60. ***Saksenaea*** S.B. Saksena, *Mycologia* 45(3): 434, 1953 *Saksenaea* is type genus of the family Saksenaeaceae (Ellis & Hesseltine 1974) and typified by *S. vasiformis* S.B. Saksena (Saksena 1953). The genus is characterized by flask-shaped or variform sporangia with a long neck (Saksena 1953). It accommodates six species which often cause mucomycosis in humans (Saksena 1953, Labuda *et al*. 2019).
61. ***Spinellus*** Tiegh., *Annls Sci. Nat.*, Bot., sér. 6 1: 66, 1875 *Spinellus* is typified by *S. fusiger* (Link) Tiegh. (basionym *Mucor fusiger* Link; Zycha *et al*. 1969) and is characterized by unbranched sporangiophores, apophysate and multi-spored, spinose zygospores, and a facultative parasitism on fungi (Zycha *et al*. 1969, Overton 1997). Six species are reported in the genus so far (www.indexfungorum.org, accessed 5 August 2021).
62. ***Sporodiniella*** Boedijn, *Sydowia* 12(1‒6): 336, 1959 [1958] *Sporodiniella* is typified by *S. umbellata* Boedijn and is characterized by umbel-branched sporangiophores, and spinose sporangiospores (Boedijn 1958). Currently, it contains only the type species which is a facultative parasite on insect larvae in the tropics (Boedijn 1958, Chien & Hwang 1997).
63. ***Syncephalastrum*** J. Schröt., *Krypt.-Fl. Schlesien* (Breslau) 3.1(9–16): 217, 1886 *Syncephalastrum* is type genus of the class Syncephalastromycetes, the order Syncephalastrales and the family Syncephalastraceae. It is typified by *S. racemosum* Cohn ex J. Schröt. (Benjamin 1959). The genus is characterized by cylindrical merosporangia, rough or dark-walled zygosporangia, and opposed, more or less equal suspensors (Benjamin 1959). It accommodates five species (www.indexfungorum.org; accessed 5 August 2021). They are a group of fungi pathogenic to humans (Pavlovic & Bulajic 2006, Rao *et al*. 2007, Irshad *et al*. 2020). In this study, four new species are described from China.
64. ***Syzygites*** Ehrenb., *Sylv. Mycol. Berol.* (Berlin): 25, 1818 *Syzygites* is typified by *S. megalocarpus* Ehrenb. and is characterized by columellate and few-spored sporangia, dichotomously branched sporangiophores, and homothallic zygospores (Kovacs & Sundberg 1999). In addition, its spinose sporangiospores are similar to those of *Sporodiniella* (Ekpo & Young 1979). Currently, it contains the type species only (Kovacs & Sundberg 1999). In nature, this species is parasitic on fleshy fungi (Kovacs & Sundberg 1999).
65. ***Thamnidium*** Link, *Mag. Gesell. Naturf. Freunde, Berlin* 3(1‒2): 31, 1809 The monotypic *Thamnidium* is typified by *T. elegans* Link and is characterized by sporangia with deliquescent walls and persistent-walled sporangiola, primary sporangiophores and lateral dichotomous branchlets (Benny 1992). As a psychrophilic fungus, *T. elegans* is isolated from soil and cold-storage meat (Brooks & Hansford 1923), and it releases proteases and collagenolytic enzymes for colonizing meat (Dashdorj *et al*. 2016).
66. ***Thamnostylum*** Arx & H.P. Upadhyay, *Gen. Fungi Sporul. Cult*. (Lehr): 247, 1970 *Thamnostylum* is typified by *T. piriforme* (Bainier) Arx & H.P. Upadhyay (basionym *Helicostylum piriforme* Bainier) and is characterized by producing spherical to pyriform apophysate sporangia and pyriform apophysate sporangiola (von Arx 1970, Upadhyay 1973, Benny & Benjamin 1975). It accommodates four species (von Arx 1970). *T. lucknowense* (J.N. Rai, J.P. Tewari & Mukerji) Arx & H.P. Upadhyay is found to be an agent of rhino-orbital mucormycosis (Xess *et al*. 2012).
67. ***Thermomucor*** Subrahm., B.S. Mehrotra & Thirum., *Georgia J. Sci*. 35(1): 1, 1977 *Thermomucor* is typified by *T. indicae-seudaticae* Subrahm., B.S. Mehrotra & Thirum. and is characterized by apophysate sporangia and unusually smooth zygospores (Subrahmanyam *et al*. 1977, Schipper 1979). Currently, it contains the type species only (Schipper 1979). This species is a human pathogen and synthesizes cellulases, glucoamylase, pectinase, and protease (Subrahmanyam & Lakshmi 1993, Kumar & Satyanarayana 2007, Merheb-Dini *et al*. 2012, Martins *et al*. 2019, de Carvalho Tavares *et al*. 2020, Pereira *et al*. 2020).
68. ***Tortumyces*** L.S. Loh, *Mucoraceous Fungi from Malaysia* (Kuala Lumpur): 80, 2001 *Tortumyces* is typified by *T. fimicola* L.S. Loh (Loh *et al*. 2001). It accommodates two species, *T. cameronensis* L.S. Loh and *T. fimicola* (Loh *et al*. 2001). However, their cultures and sequences are unavailable so far.
69. ***Umbelopsis*** Amos & H.L. Barnett, *Mycologia* 58(5): 807, 1966 *Umbelopsis* is type genus of the phylum Umbelopsidomycota, the class Umbelopsidales and the family Umbelopsidaceae. It is typified by *U. versiformis* Amos & H.L. Barnett (Amos & Barnett 1966). The genus differs from its allied genus *Mortierella* in having small but distinct columellae and ochraceous to reddish colonies (Meyer & Gams 2003, Wang *et al*. 2013, 2014, 2015). Currently, 18 species are accepted in the genus (von Arx 1982, Yip 1986, Meyer & Gams 2003, Wang *et al*. 2014, 2015, Crous *et al*. 2017, Wijayawardene *et al*. 2018, Yuan *et al*. 2020). They are distributed worldwide in soil and are also endophytic (Qin *et al*. 2018). Some species of the genus are used for lipid production (Gardeli *et al*. 2017, Zhao *et al*. 2021a).
70. ***Utharomyces*** Boedijn ex P.M. Kirk & Benny, *Trans. Br. Mycol. Soc*. 75(1): 124, 1980 *Utharomyces* is typified by *U. epallocaulus* Boedijn and is similar to *Pilobolus* in having sporangiophores that arise from a trophocyte in the substrate hyphae and with subsporangial swellings but differs in sporangia which are not violently discharged (Boedijn 1958, Kirk & Benny 1980, Hu *et al*. 1989). Currently, it contains the type species only (Kirk & Benny 1980).
71. ***Zychaea*** Benny & R.K. Benj., *Aliso* 8(3): 334, 1975 *Zychaea* is typified by *Z. mexicana* Benny & R.K. Benj. and is characterized by sporangiola which are pedicellate, and sporangiospores which are globose to subglobose, uni-spored or multi-spored, ovoid to ellipsoid (Benny & Benjamin 1975). Currently, it contains the type species only (Benny & Benjamin 1975). Strains of the species were isolated from dung (Benny & Benjamin 1975).

**Fig. 5.**
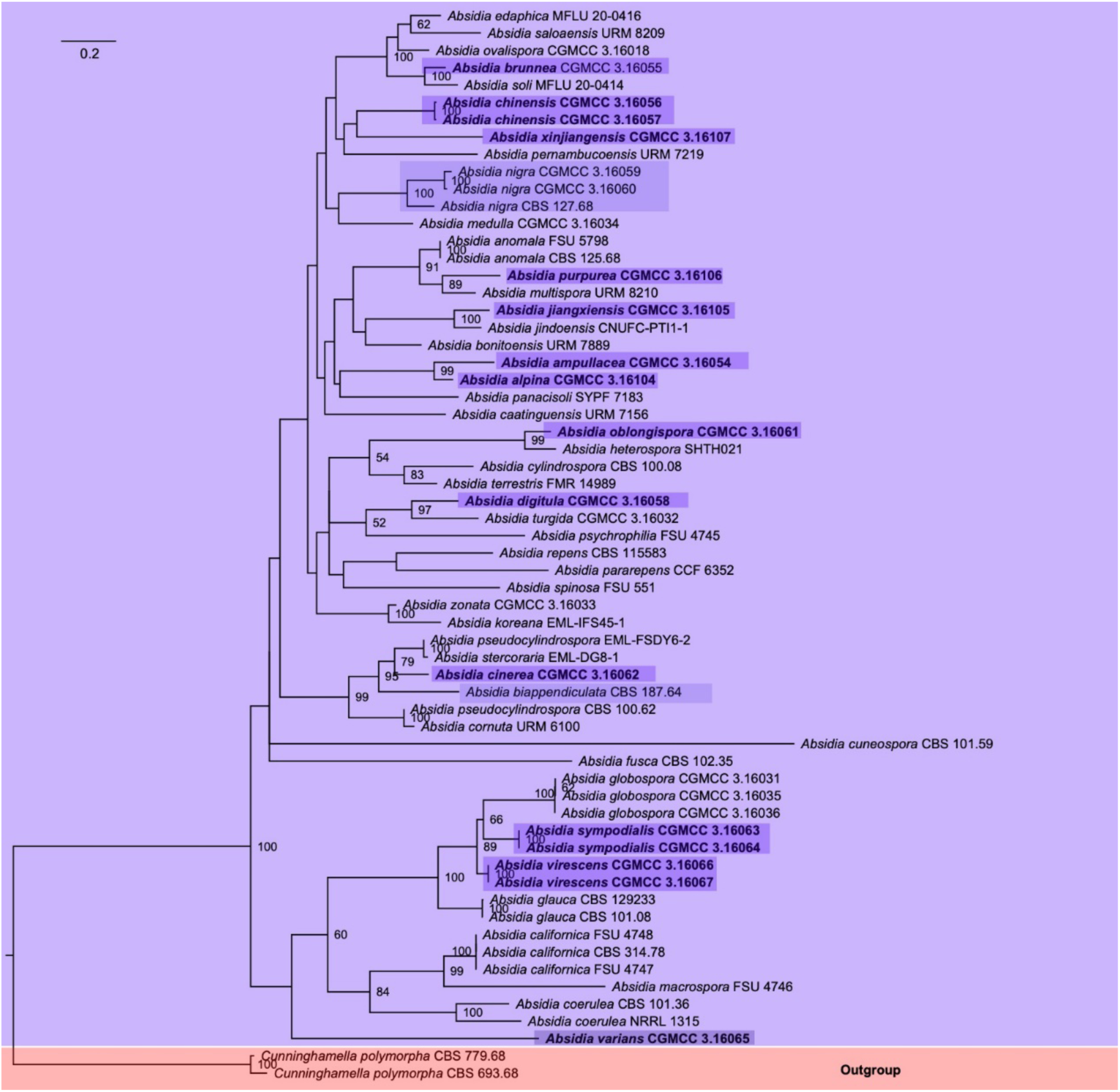
Maximum Likelihood phylogenetic tree of *Absidia* based on ITS rDNA sequences, with *Cunninghamella polymorpha* as outgroup. Thirteen new species, *A. alpina*, *A. ampullacea*, *A. brunnea*, *A. chinensis*, *A. cinerea*, *A. digitula*, *A. jiangxiensis*, *A. oblongispora*, *A. purpurea*, *A. sympodialis*, *A. xinjiangensis*, *A. varians*, *A. virescens*, and two new combinations, *A. biappendiculata* and *A. nigra*, are in shade. Maximum Likelihood (ML) bootstrap values (≥50%) of each clade are indicated at nodes. A scale bar in the upper left indicates substitutions per site.

**Fig. 6.**
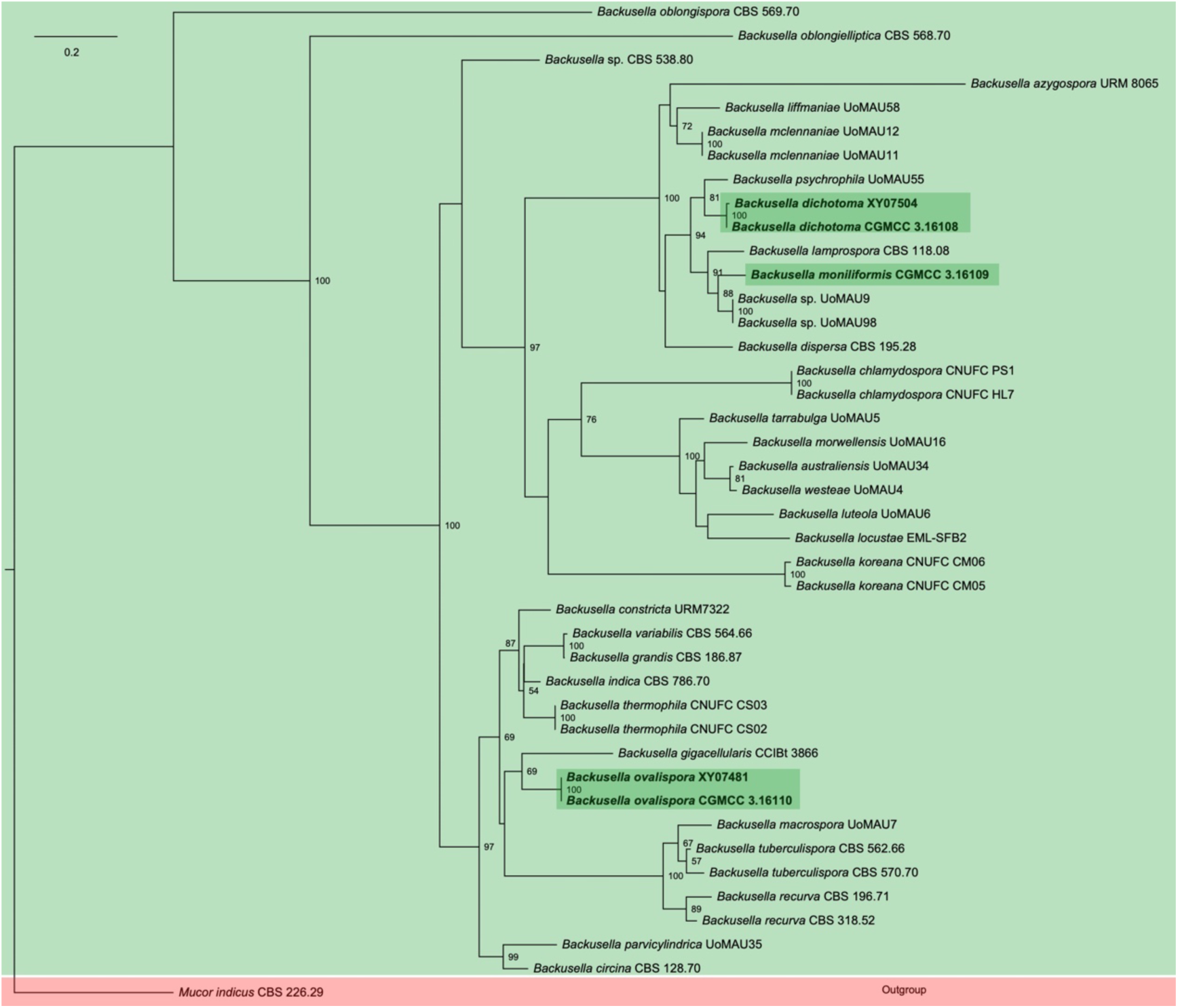
Maximum Likelihood phylogenetic tree of *Backusella* based on ITS rDNA sequences, with *Mucor indicus* as outgroup. Three new species, *B. dichotoma*, *B. moniliformis* and *B. ovalispora*, are in shade. Maximum Likelihood (ML) bootstrap values (≥50%) of each clade are indicated at nodes. A scale bar in the upper left indicates substitutions per site.

**Fig. 7.**
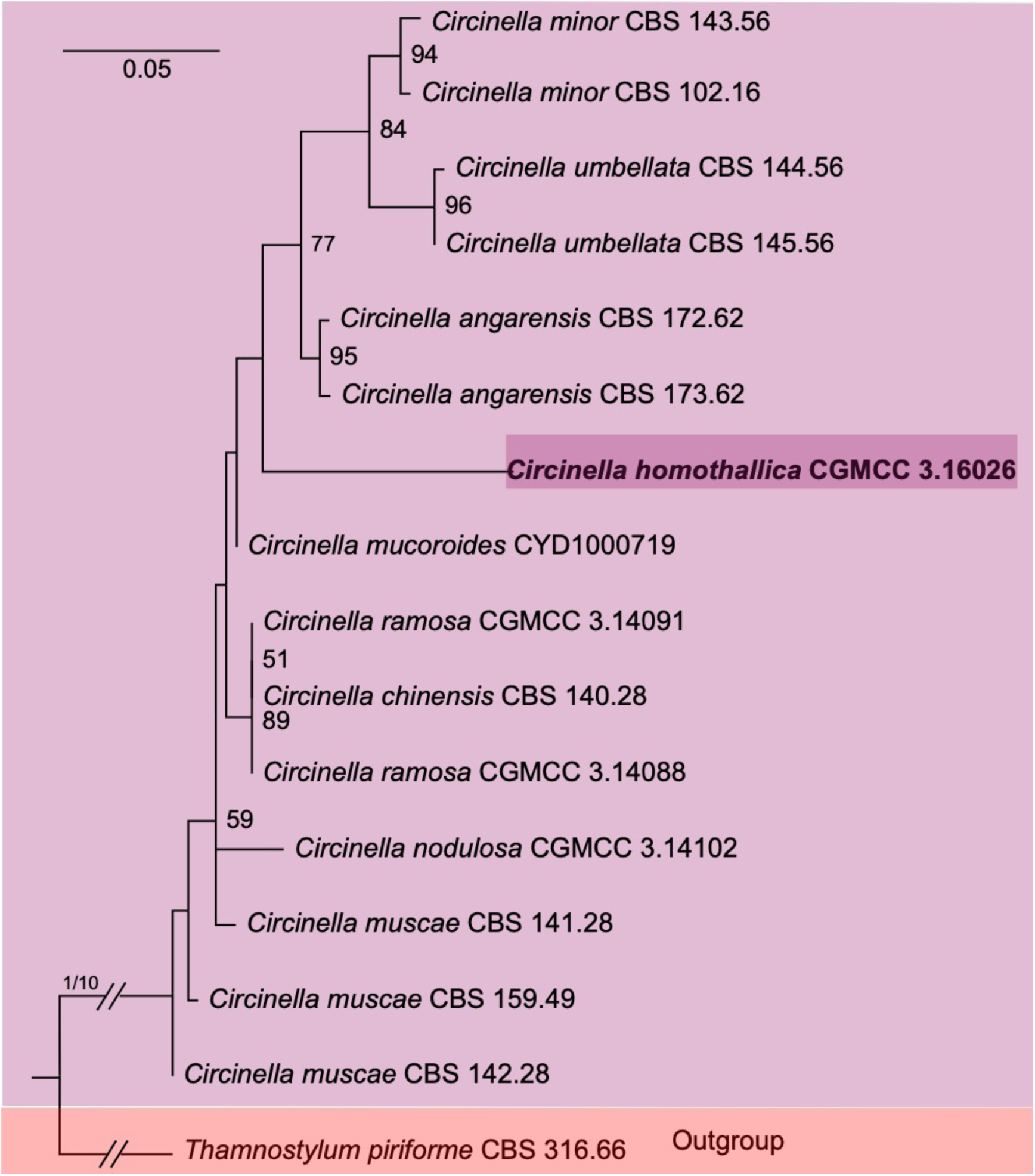
Maximum Likelihood phylogenetic tree of *Circinella* based on ITS rDNA sequences, with *Thamnostylum piniforme* as outgroup. One new species, *C. homothallica*, is in shade. Maximum Likelihood (ML) bootstrap values (≥50%) of each clade are indicated at nodes. A scale bar in the upper left indicates substitutions per site. Double slashes indicate branch lengths shortened ten-fold to facilitate visualization.

**Fig. 8.**
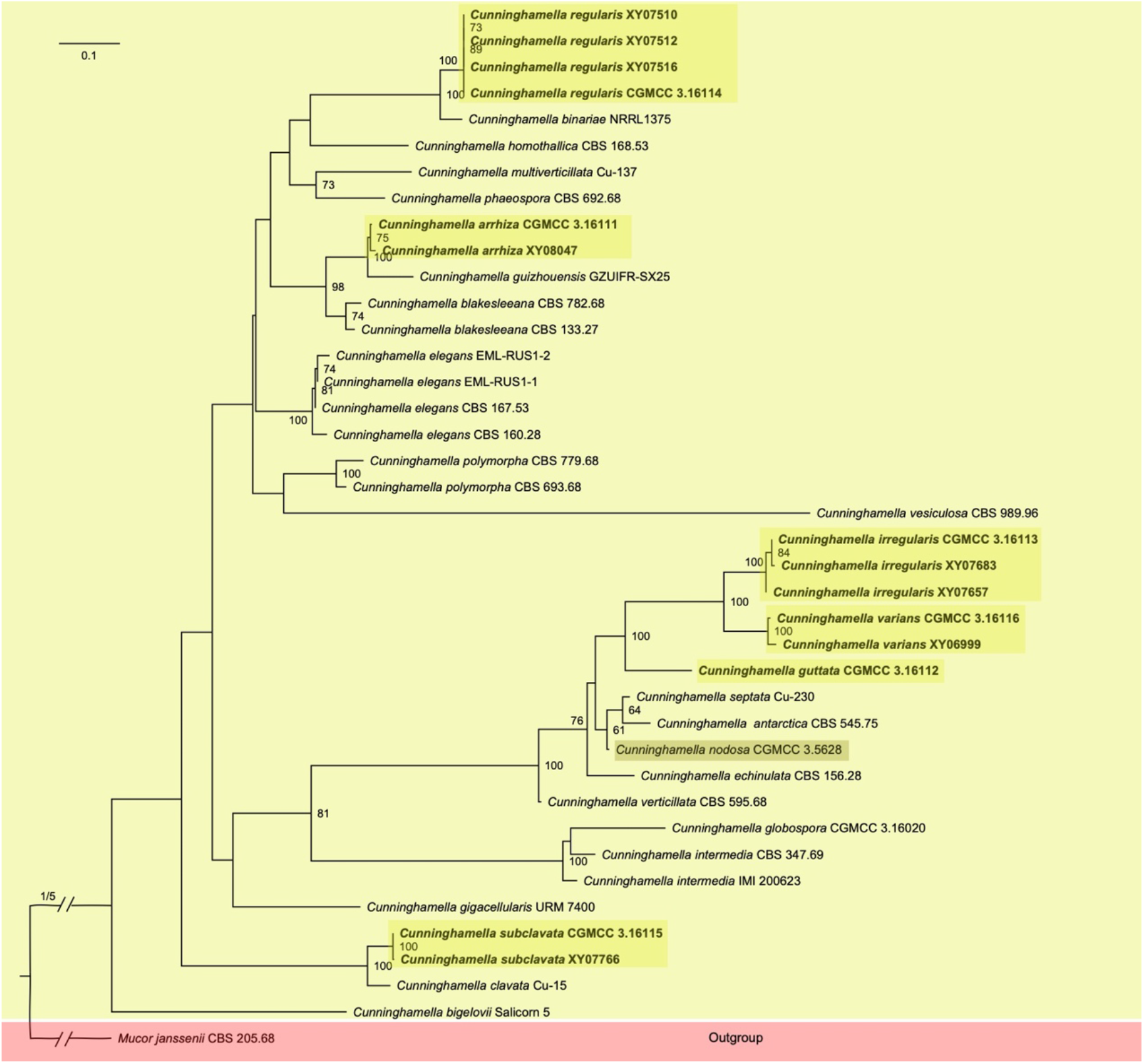
Maximum Likelihood phylogenetic tree of *Cunninghamella* based on ITS rDNA sequences, with *Mucor janssenii* as outgroup. Six new species, *C. arrhiza*, *C. guttata*, *C. irregularis*, *C. regularis*, *C. subclavata*, *C. varians*, and one combination, *C. nodosa*, are in shade. Maximum Likelihood (ML) bootstrap values (≥50%) of each clade are indicated at nodes. A scale bar in the upper left indicates substitutions per site. Double slashes indicate branch lengths shortened five-fold to facilitate visualization.

**Fig. 9.**
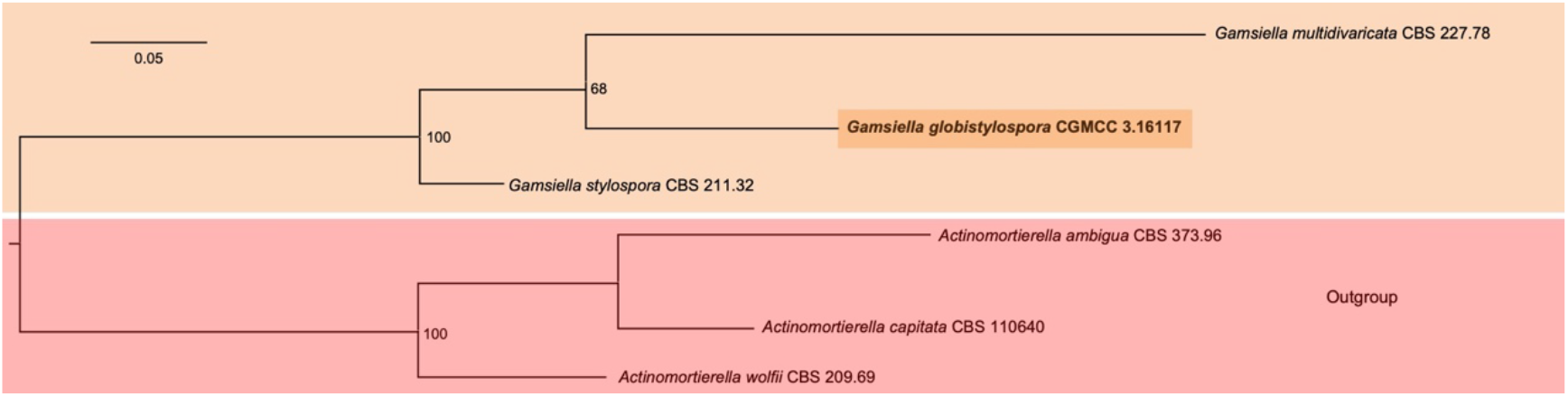
Maximum Likelihood phylogenetic tree of *Gamsiella* based on ITS rDNA sequences, with *Actinomortierella* as outgroup. One new species, *G. globistylospora*, is in shade. Maximum Likelihood (ML) bootstrap values (≥50%) of each clade are indicated at nodes. A scale bar in the upper left indicates substitutions per site.

**Fig. 10.**
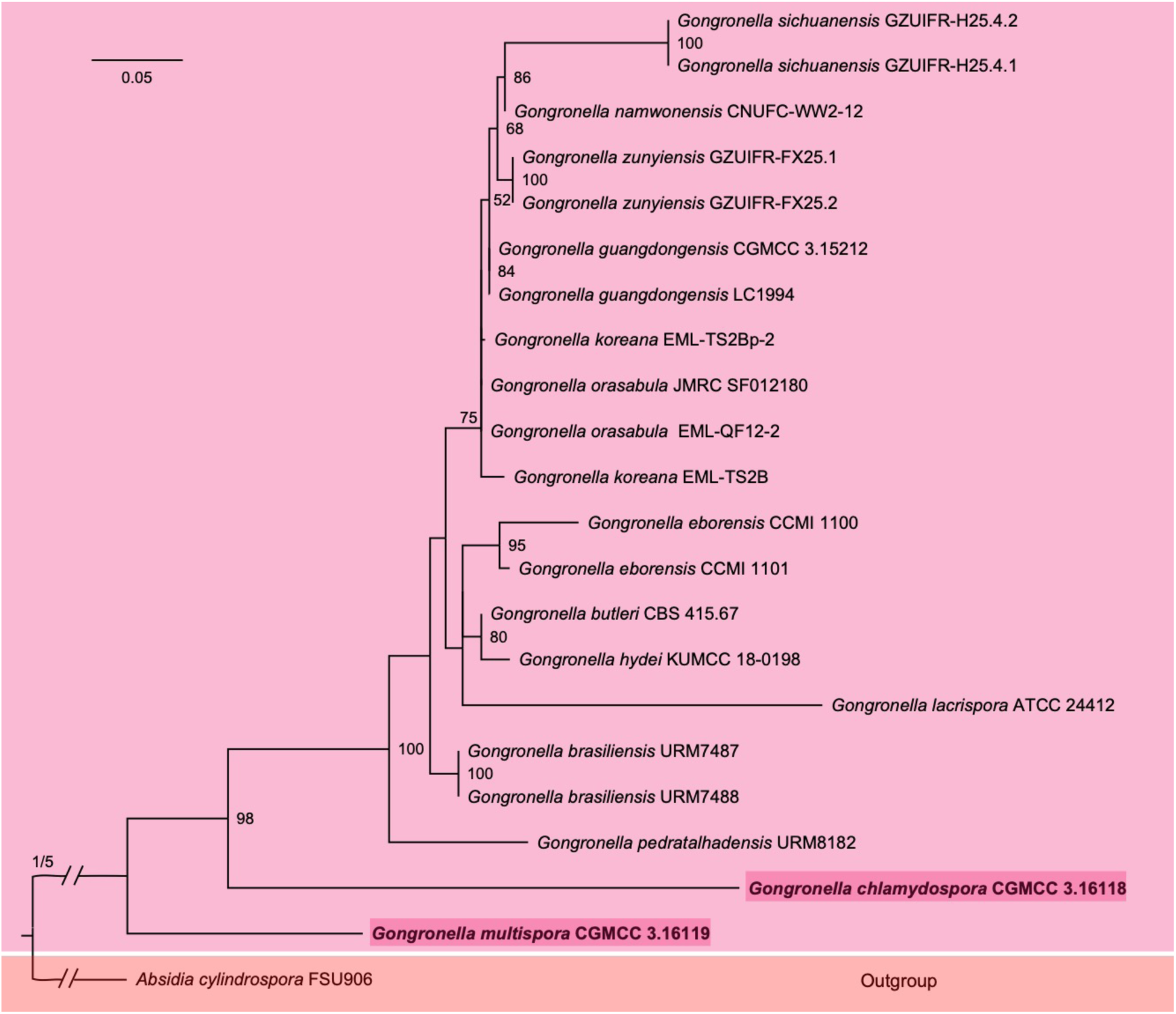
Maximum Likelihood phylogenetic tree of *Gongronella* based on ITS rDNA sequences with *Absidia cylindrospora* as outgroup. Two new species, *G. chlamydospora* and *G. multispora*, are in shade. Maximum Likelihood (ML) bootstrap values (≥50%) of each clade are indicated at nodes. A scale bar in the upper left indicates substitutions per site. Double slashes indicate branch lengths shortened five-fold to facilitate visualization.

**Fig. 11.**
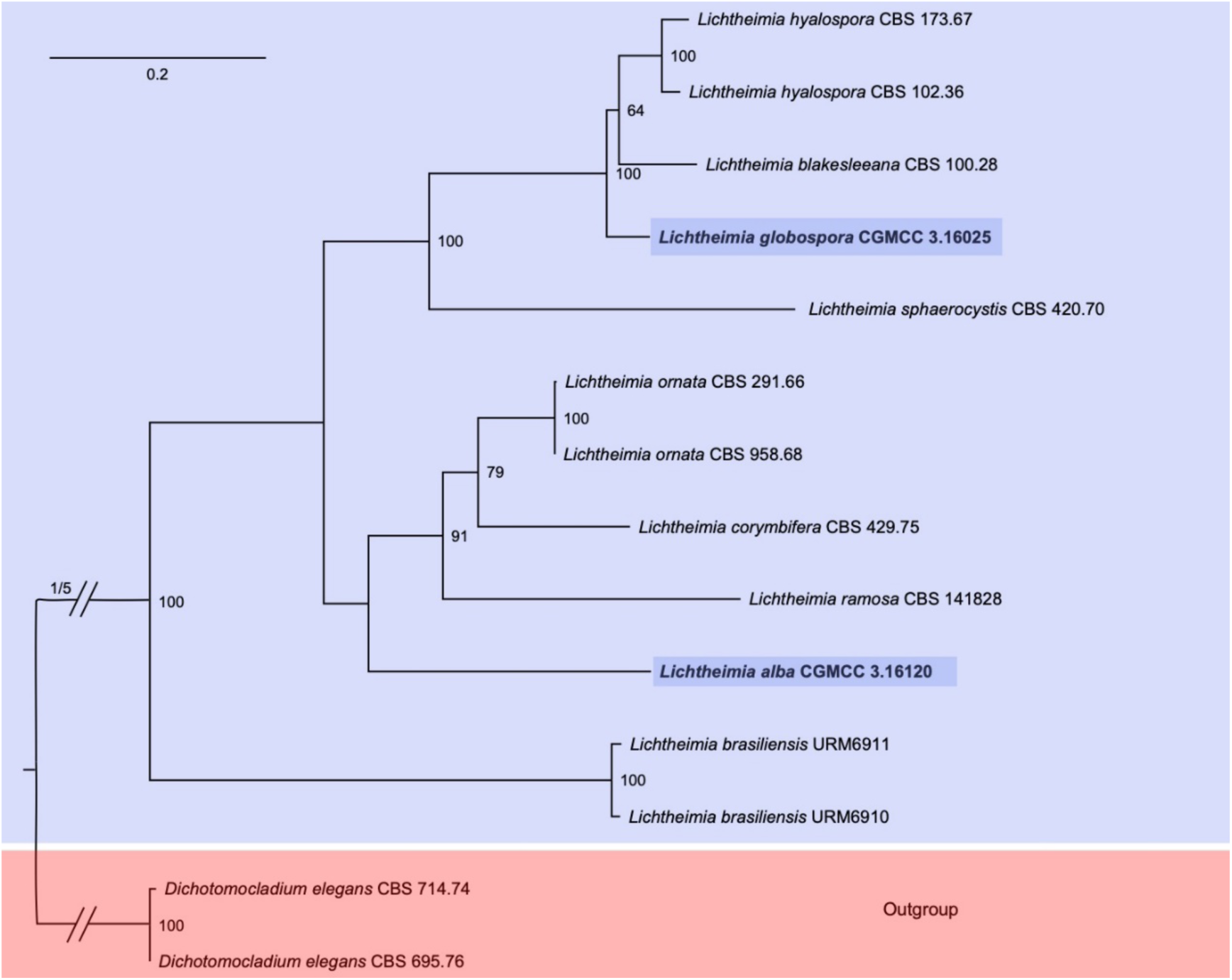
Maximum Likelihood phylogenetic tree of *Lichtheimia* based on ITS rDNA sequences, with *Dichotomocladium elegans* as outgroup. Two new species, *L. alba* and *L. globospora*, are in shade. Maximum Likelihood (ML) bootstrap values (≥50%) of each clade are indicated at nodes. A scale bar in the upper left indicates substitutions per site. Double slashes indicate branch lengths shortened five-fold to facilitate visualization.

**Fig. 12.**
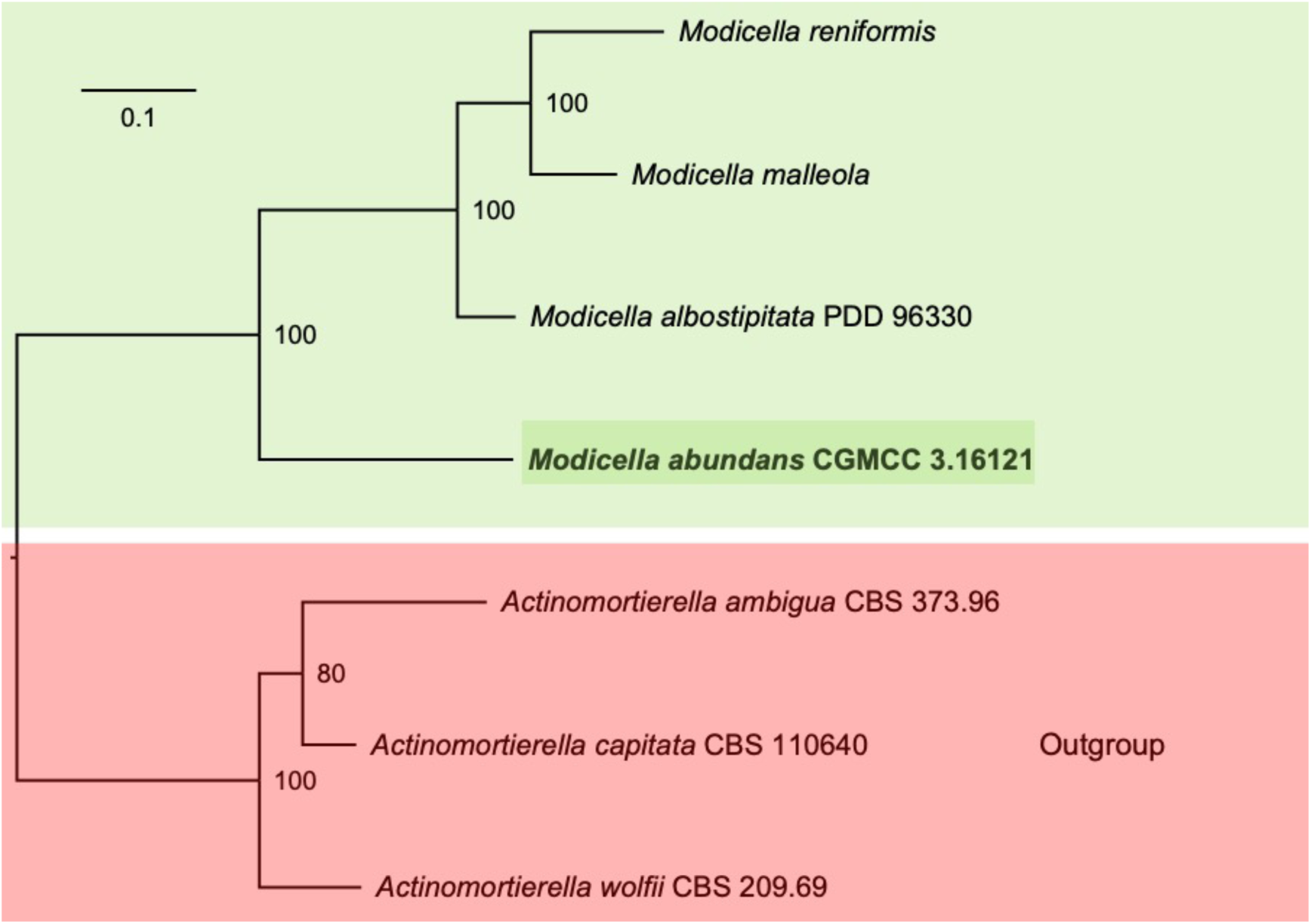
Maximum Likelihood phylogenetic tree of *Modicella* based on ITS rDNA sequences, with *Actinomortierella* as outgroup. One new species, *M. abundans*, is in shade. Maximum Likelihood (ML) bootstrap values (≥50%) of each clade are indicated at node. A scale bar in the upper left indicates substitutions per site.

**Fig. 13.**
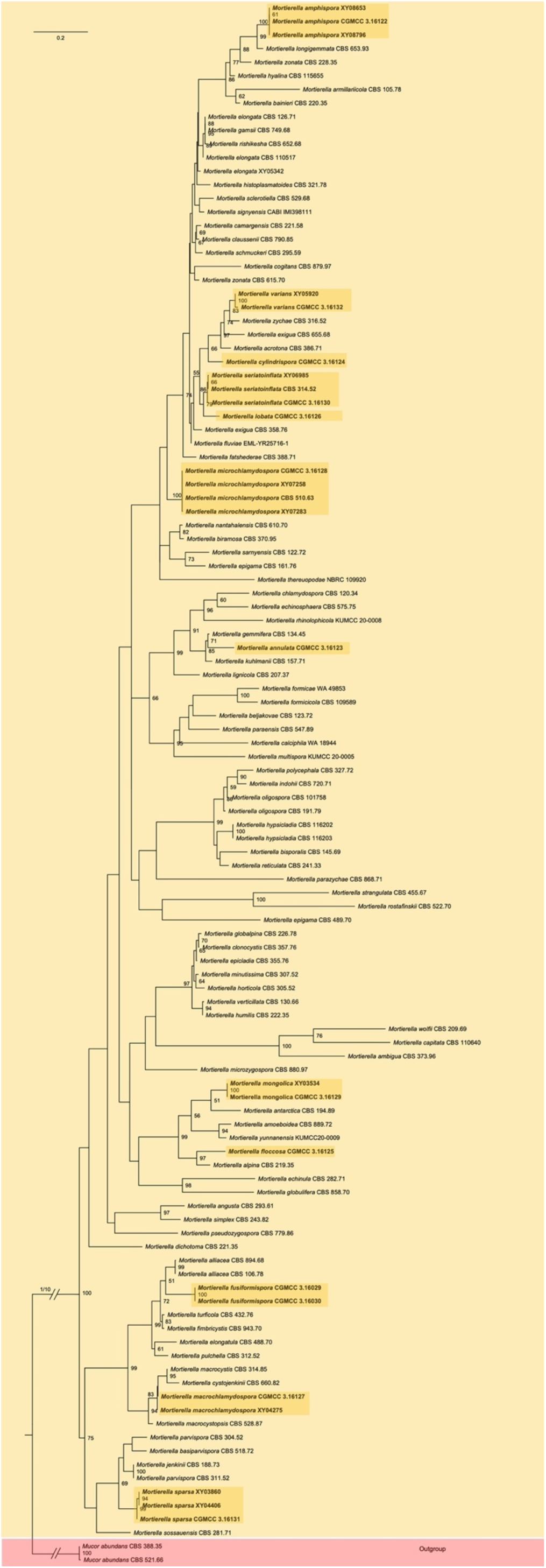
Maximum Likelihood phylogenetic tree of *Mortierella* based on ITS rDNA sequences, with *Mucor abundans* as outgroup. Twelve new species, *M. amphispora*, *M. annulata*, *M. cylindrispora*, *M. floccosa*, *M. fusiformispora*, *M. lobata*, *M. macrochlamydospora*, *M. microchlamydospora*, *M. mongolica*, *M. seriatoinflata*, *M. sparsa* and *M. varians*, are in shade, Maximum Likelihood (ML) bootstrap values (≥50%) of each clade are indicated at nodes. A scale bar in the upper left indicates substitutions per site. Double slashes indicate branch lengths shortened ten-fold to facilitate visualization.

**Fig. 14.**
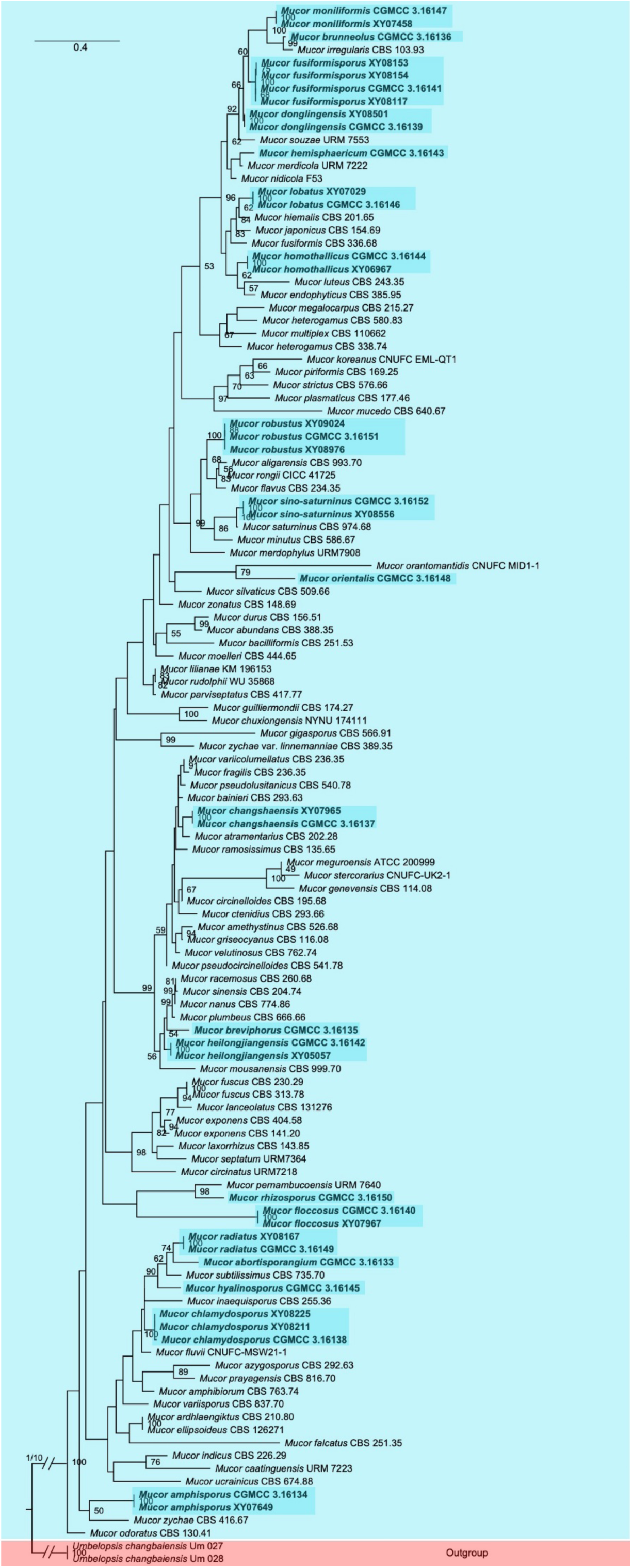
Maximum Likelihood phylogenetic tree of *Mucor* based on ITS rDNA sequences, with *Umbelopsis changbaiensis* as outgroup. Twenty new species, *M. abortisporangium*, *M. amphisporus*, *M. breviphorus*, *M. brunneolus*, *M. changshaensis*, *M. chlamydosporus*, *M. donglingensis*, *M. floccosus*, *M. fusiformisporus*, *M. heilongjiangensis*, *M. hemisphaericum*, *M. homothallicus*, *M. hyalinosporus*, *M. lobatus*, *M. moniliformis*, *M. orientalis*, *M. radiatus*, *M. rhizosporus*, *M. robustus* and *M. sino-saturninus*, are in shade. Maximum Likelihood (ML) bootstrap values (≥50%) of each clade are indicated at nodes. A scale bar in the upper left indicates substitutions per site. Double slashes indicate branch lengths shortened ten-fold to facilitate visualization.

**Fig. 15.**
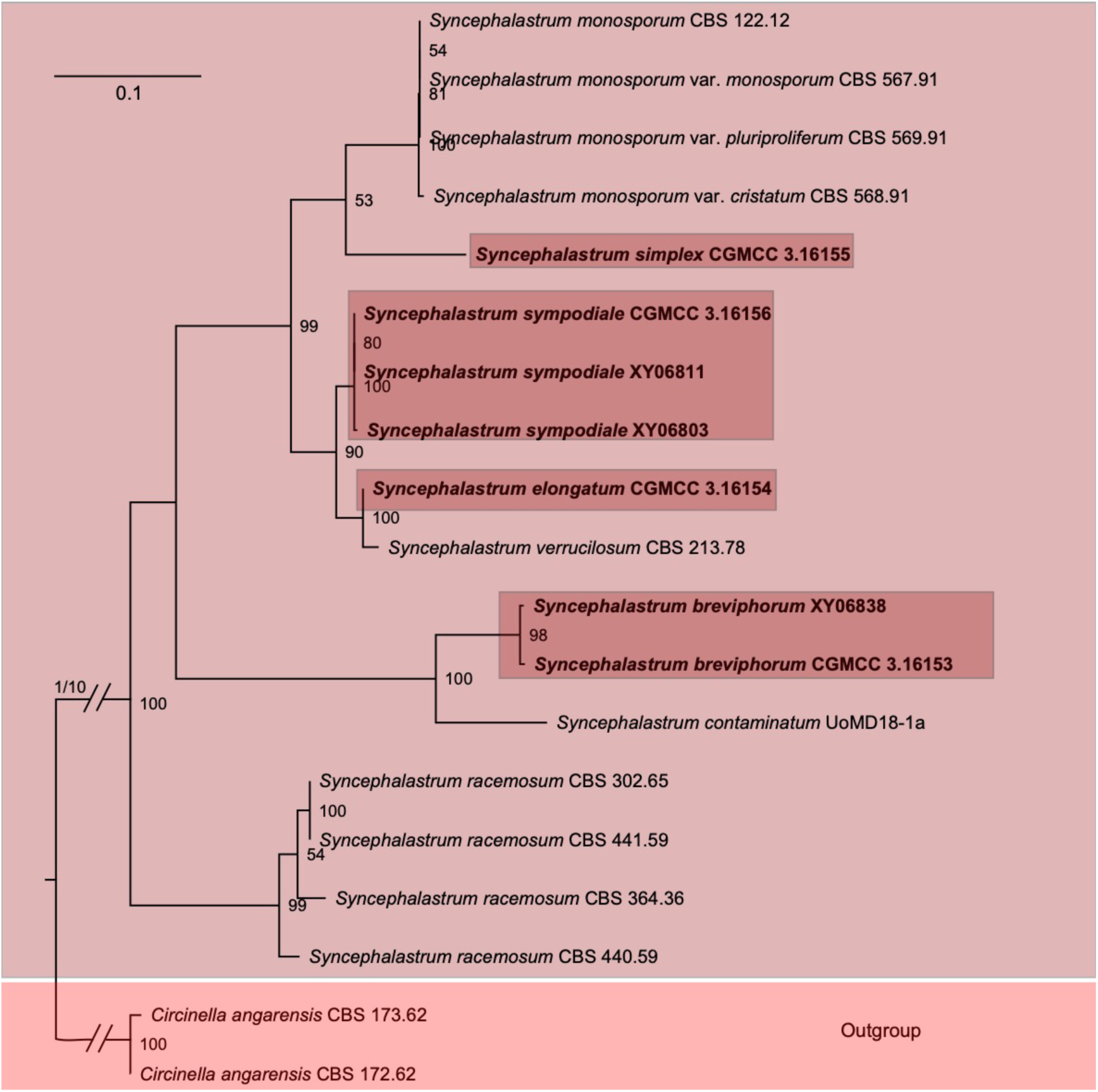
Maximum Likelihood phylogenetic tree of *Syncephalastrum* based on ITS rDNA sequences, with *Circinella angarensis* as outgroup. Four new species, *S. breviphorum*, *S. elongatum*, *S. simplex* and *S. sympodiale*, are in shade. Maximum Likelihood (ML) bootstrap values (≥50%) of each clade are indicated at nodes. A scale bar in the upper left indicates substitutions per site. Double slashes indicate branch lengths shortened ten-fold to facilitate visualization.

**Fig. 16.**
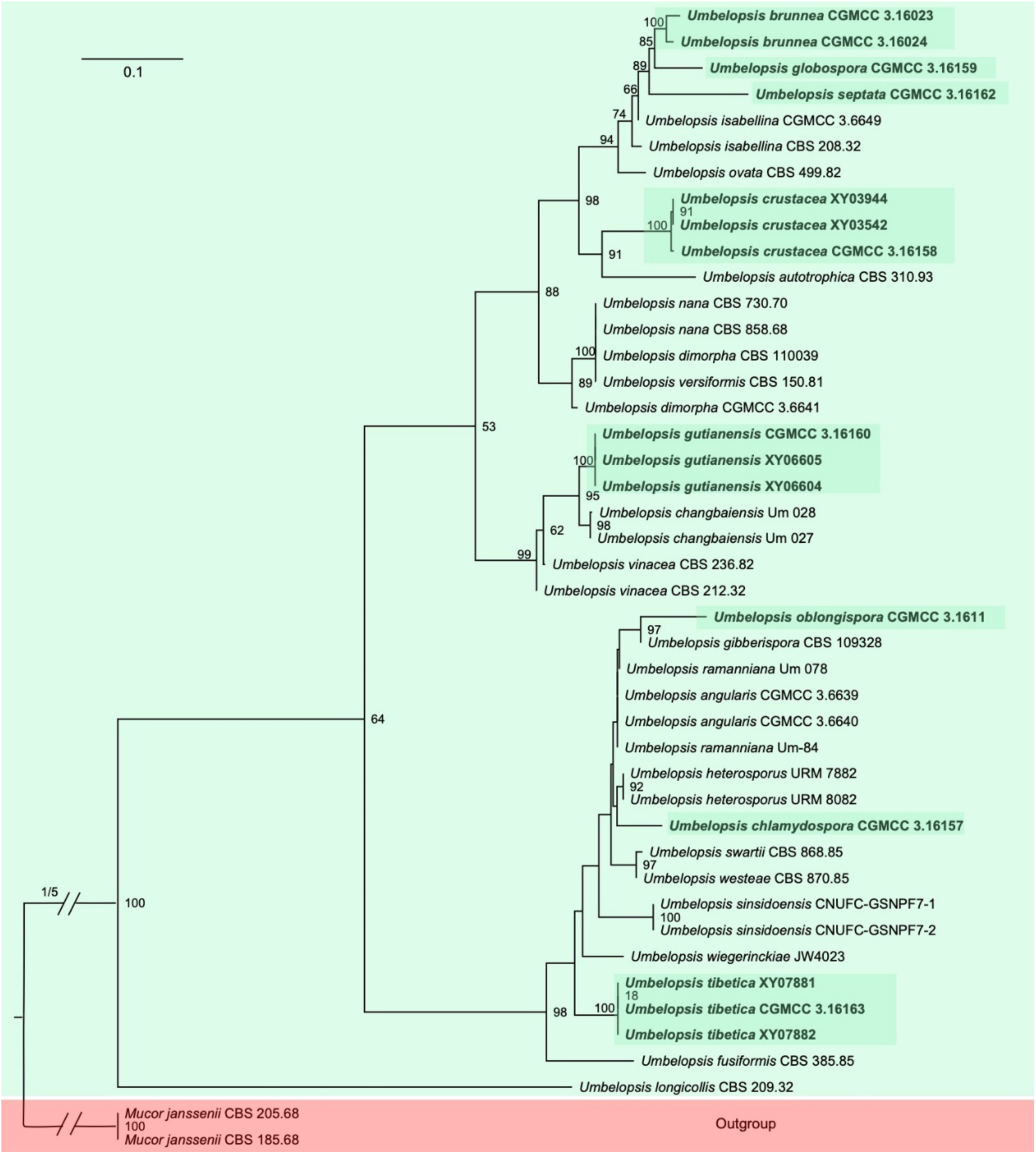
Maximum Likelihood phylogenetic tree of *Umbelopsis* based on ITS rDNA sequences, with *Mucor janssenii* as outgroup. Eight new species, *U. brunnea*, *U. chlamydospora*, *U. crustacea*, *U. globospora*, *U. gutianensis*, *U. oblongispora*, *U. septata* and *U. tibetica*, are in shade. Maximum Likelihood (ML) bootstrap values (≥50%) of each clade are indicated at nodes. A scale bar in the upper left indicates substitutions per site. Double slashes indicate branch lengths shortened ten-fold to facilitate visualization.

### Taxonomy of new species and new combinations

In the present study, we describe 73 new species in the Mortierellomycota, Mucoromycota and Umbelopsidomycota (Table 1). Besides, three new combinations are elevated from a variety status (Table 1). All these new taxa are illustrated below.

**1. *Absidia alpina*** H. Zhao, Y.C. Dai & X.Y. Liu, sp. nov., Fig. 17.

**Fig. 17.**
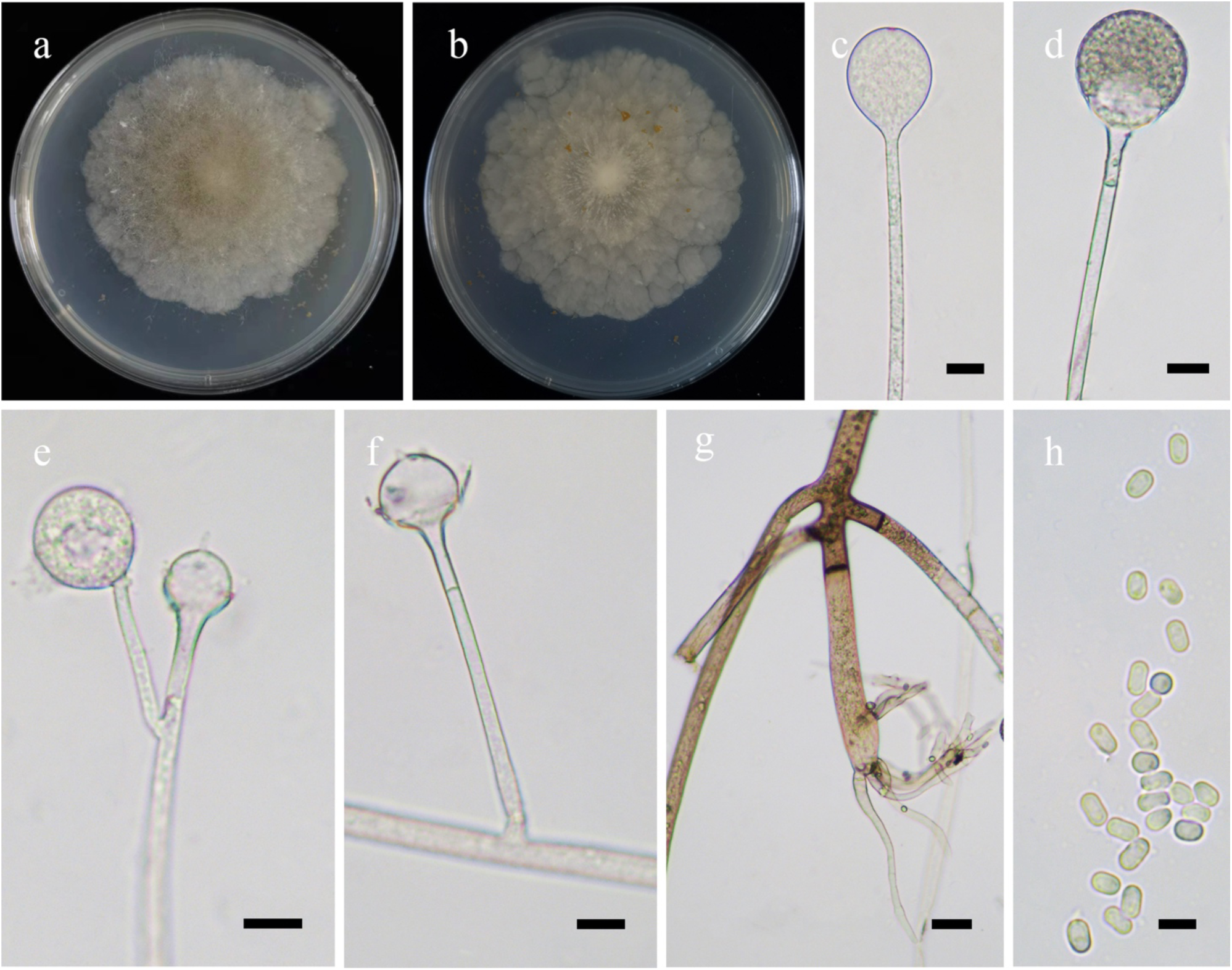
Morphologies of *Absidia alpina* ex-holotype CGMCC 3.16104. **a, b**. Colonies on MEA (**a.** obverse, **b.** reverse); **c–e.** Sporangia; **f.** Columellae; **g.** Rhizoids; **h.** Sporangiospores. — Scale bars: **c–f.** 10 μm, **g.** 20 μm, **h.** 5 μm.

*Fungal Names*: FN570882.

*Etymology*: *alpina* (Lat.) refers to the mountain environment where the type was collected.

*Holotype*: HMAS 351506.

*Colonies* on MEA at 27 °C for 10 days, slow growing, reaching 65 mm in diameter, irregular concentrically zonate with ring, white at first, gradually becoming Light Brown to Sudan Brown. *Hyphae* hyaline at first, brown when mature, aseptate when juvenile, septate with age, 3.0–15.5 µm in diameter. *Stolons* branched, hyaline to brownish, smooth, with few septa. *Rhizoids* root-like, always unbranched, with a septum at the top. *Sporangiophores* erect or slightly bent, 1–5 in whorls, often monopodial, rarely simple unbranched, hyaline, with a septum 11.5–18.0 µm below apophyses, 35.0–300.0 µm or more in length and 2.5–5.0 µm in width. *Sporangia* subglobose, smooth, multi-spored, hyaline when young, brown when old, 12.5–30.0 µm long and 16.0–28.0 µm wide, walls deliquescent. *Apophyses* distinct, slightly pigmented, 3.0–6.0 µm high, 4.0–7.0 µm wide at the base, and 6.5–15.0 µm wide at the top. *Collars* present or absent, and distinct if present. *Columellae* subglobose to ellipsoid, hyaline, smooth, some with a 3.0–7.5 µm projection at the apex, 10.0–18.5 µm long and 12.5–20.0 µm wide. *Sporangiospores* cylindrical, hyaline, smooth, slightly constricted in center, 3.5–5.5 µm long and 2.0–2.5 µm wide. *Zygospores* absent.

*Chlamydospores* absent.

*Material examined*: China, Yunnan Province, Dali, Huacong Mountain, 26°25’21″N, 99°53’41″E, from soil sample, 28 October 2021, Heng Zhao (holotype HMAS 351506, living ex-holotype culture CGMCC 3.16104).

*GenBank accession number*: OL678133.

*Notes*: *Absidia alpina* is closely related to *A. ampullacea* T.K. Zong *et al*. based on phylogenetic analysis of ITS rDNA sequences (Fig. 5). However, morphologically *A. ampullacea* differs from *A. alpina* by ovoid to subglobose sporangiospores, sympodially-branched and swollen sporangiophores.

**2. *Absidia ampullacea*** T.K. Zong, H. Zhao, Y.C. Dai & X.Y. Liu, sp. nov., Fig. 18.

**Fig. 18.**
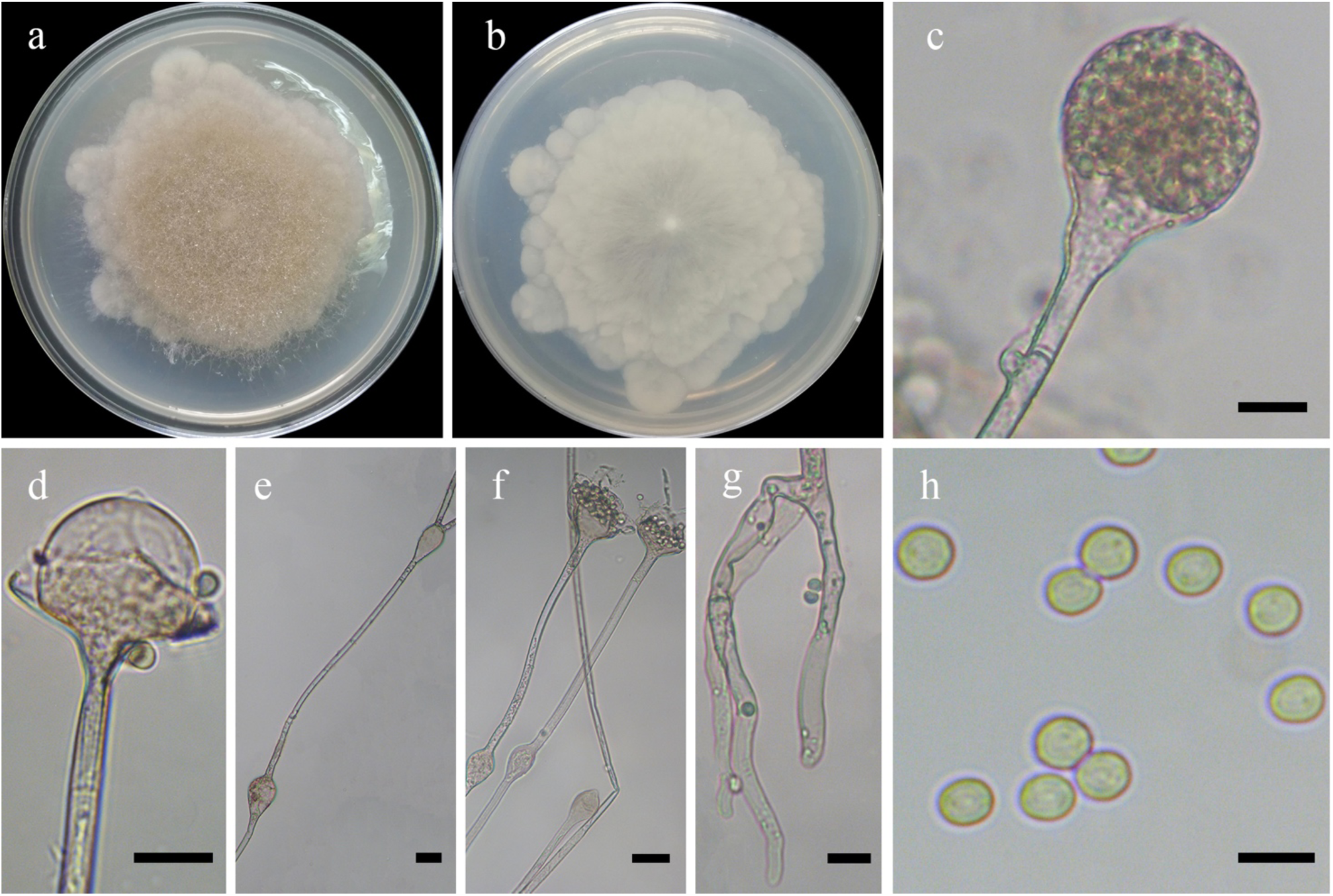
Morphologies of *Absidia ampullacea* ex-holotype CGMCC 3.16054. **a, b**. Colonies on MEA (**a.** obverse, **b.** reverse); **c.** Sporangia; **d.** Columellae; **e, f.** Swelling on sporangiophores and hyphae; **g.** Rhizoids; **h.** Sporangiospores. — Scale bars: **c, d, g.** 10 μm, **e, f.** 20 μm, **h.** 5 μm.

*Fungal Names*: FN570883.

*Etymology*: *ampullacea* (Lat.) refers to the species having ampulliform swollen hyphae and sporangiophores.

*Holotype*: HMAS 350295.

*Colonies* on MEA at 27 °C for 10 days, slow growing, reaching 90 mm in diameter, irregularly zonate, white at first and then Saccardo’s Olive. *Hyphae* hyaline at first, becoming brown when mature, sometimes ampulliform-shaped swollen, 6.0–16.5 µm in diameter. *Stolons* branched, hyaline, smooth, septate, 4.0–9.0 µm in diameter. *Rhizoids* finger-like, mostly twice or more repeatedly, with a septum at the base. *Sporangiophores* erect or slightly bent, 1–6 in whorls, unbranched, simple branched, monopodial, or sympodial, hyaline, with a septum 14.5–21.5 µm below apophyses, sometimes a swelling beneath sporangium, 30.0–320.0 µm long and 2.5–5.5 µm wide. *Sporangia* globose to pyriform, smooth, multi-spored, 17.5–37.5 µm long and 17.5–45.0 µm wide, walls deliquescent. *Apophyses* distinct, bell-shaped, slightly pigmented, 4.0–9.5 µm high, 2.5–8.0 µm wide at the base, and 9.5–20.0 µm wide at the top. *Collars* distinct. *Columellae* hemispherical, hyaline, smooth, occasionally with a 0.5–2.0 µm verrucous projection at the apex, 8.5–22.5 µm long and 10.5–24.0 µm wide. *Sporangiospores* ovoid or subglobose, 3.0–4.5 µm long and 2.5–4.0 µm wide. *Zygospores* absent. *Chlamydospores* absent. No growth at 30 °C.

*Material examined*: China, Beijing, 40°3′49″N, 116°1′26″E, from soil sample, 31 December 2019, Xiao-Yong Liu (holotype HMAS 350295, living ex-holotype culture CGMCC 3.16054).

*GenBank accession number*: MZ354138.

*Notes*: See the notes in *Absidia alpina*.

**3. *Absidia biappendiculata*** (Rall & Solheim) T.K. Zong, H. Zhao, Y.C. Dai & X.Y. Liu, comb. et stat. nov., Fig. 19.

**Fig. 19.**
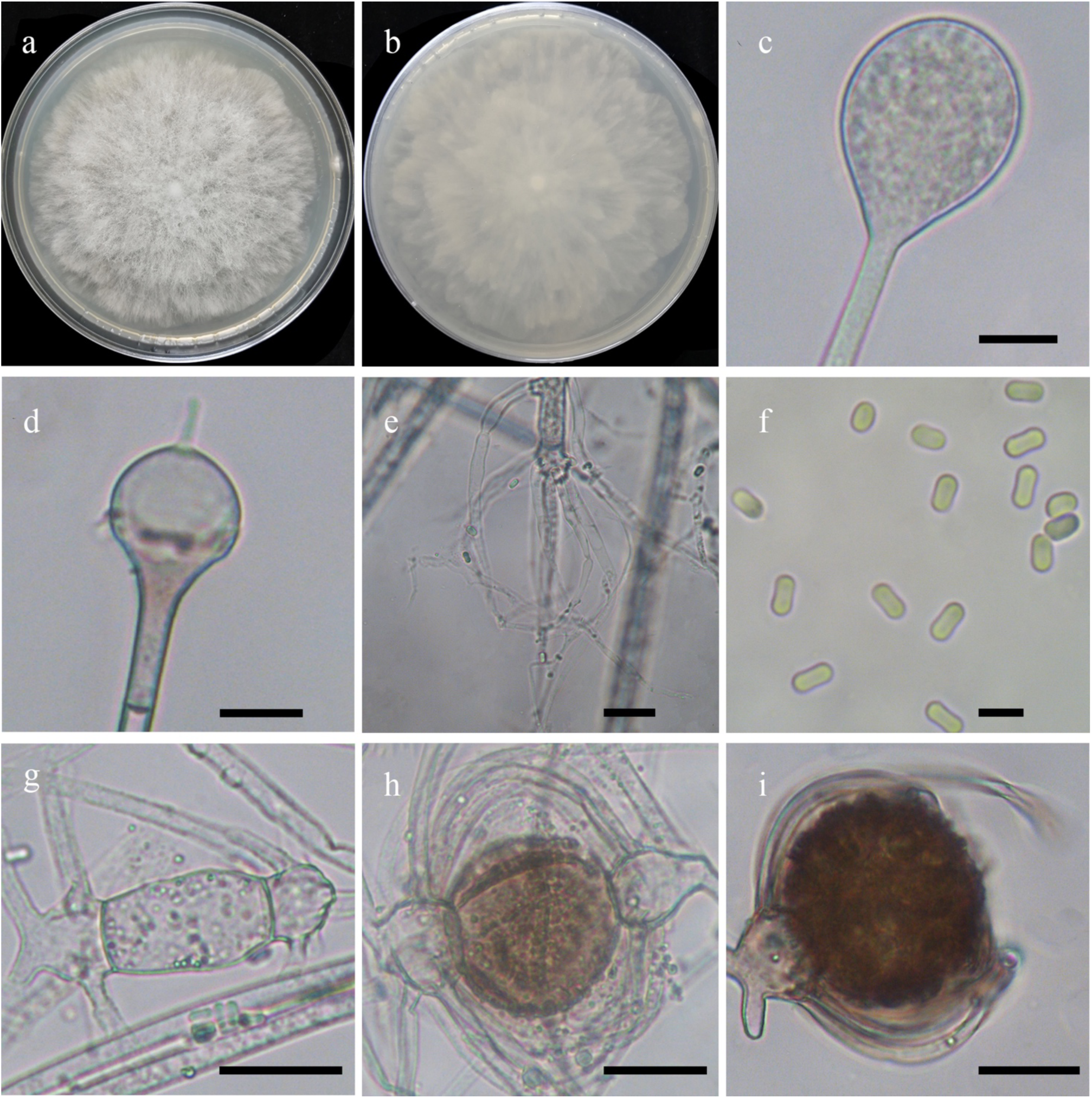
Morphologies of *Absidia biappendiculata* ex-holotype CBS 187.64. **a, b**. Colonies on MEA (**a.** obverse, **b.** reverse); **c.** Sporangia; **d.** Columellae; **e.** Rhizoids; **f.** Sporangiospores; **g–i.** Zygospores. — Scale bars: **c, d.** 10 μm, **f.** 5 μm, **e, g–i.** 20 μm.

*Fungal Names*: FN570884.

Basionym: *Absidia spinosa* var. *biappendiculata* Rall & Solheim, Mycologia, 56(1), 99. 1964. [MB no.: 348993].

*Holotype*: CBS 187.64.

*Colonies* on MEA at 27 °C for 6 days, reaching 90 mm in diameter, regularly zonate, white at first and becoming Pale Mouse Gray with age. *Hyphae* hyaline to slightly pigmented, 5.5–11.0 µm in diameter. *Stolons* branched, hyaline, smooth, septate, 3.5–5.5 µm in diameter. *Rhizoids* root-like, tapering at the end, with a septum at the base. *Sporangiophores* erect or slightly bent, 1–5 in whorls, unbranched or simple branched, monopodial, and sympodial forms absent, hyaline, with a septum 11.0–17.0 µm below apophyses, 65.0–210.0 µm long and 2.5–5.0 µm wide. *Sporangia* globose to pyriform, smooth, multi-spored, 16.0–29.0 µm long and 15.5–27.0 µm wide, walls deliquescent. *Apophyses* distinct, slightly pigmented, 3.5–9.0 µm high, 2.5–4.5 µm wide at the base, and 8.0–20.0 µm wide at the top. *Collars* distinct if present. *Columellae* hemispherical to subglobose, hyaline, smooth, sometimes with a 3.0–7.0 µm spinous projection at the apex, occasionally with two projections, or with a conical projection in some of smaller columellae, 7.5–19.0 µm long and 8.5– 20.0 µm wide. *Sporangiospores* cylindrical to ovoid, slightly constricted in the center, hyaline, smooth, 3.5–4.5 µm long and 2.0–2.5 µm wide. *Zygospores* globose, brown to Dark Brown, rough, 33.0–66.0 µm in diameter. *Suspensor* with appendages, mostly two, nearly equal, parallel or nearly so, sometimes only one, hyaline or brown, 14.0–27.0 µm in diameter. *Chlamydospores* absent. No growth at 35 °C.

*Material examined*: USA, Wyoming, Albany County, from leaves of *Comandra pallida*, August 1960, G. Rall (NRRL 3033, isotype CBS 187.64, isotype HMAS 350310)

*GenBank accession number*: MZ354153

*Notes*: *Absidia biappendiculata* was previously regarded as a synonym of *A. spinosa*. However, morphologically *A. spinosa* differs from *A. biappendiculata* by wider sporangiophores (5.0–10.5 µm wide vs. 2.0–2.5 µm wide), producing one projection on columellae only (Hesseltine & Ellis 1964, Rall & Solheim 1964).

**4. *Absidia brunnea*** T.K. Zong, H. Zhao, Y.C. Dai & X.Y. Liu, sp. nov., Fig. 20.

**Fig. 20.**
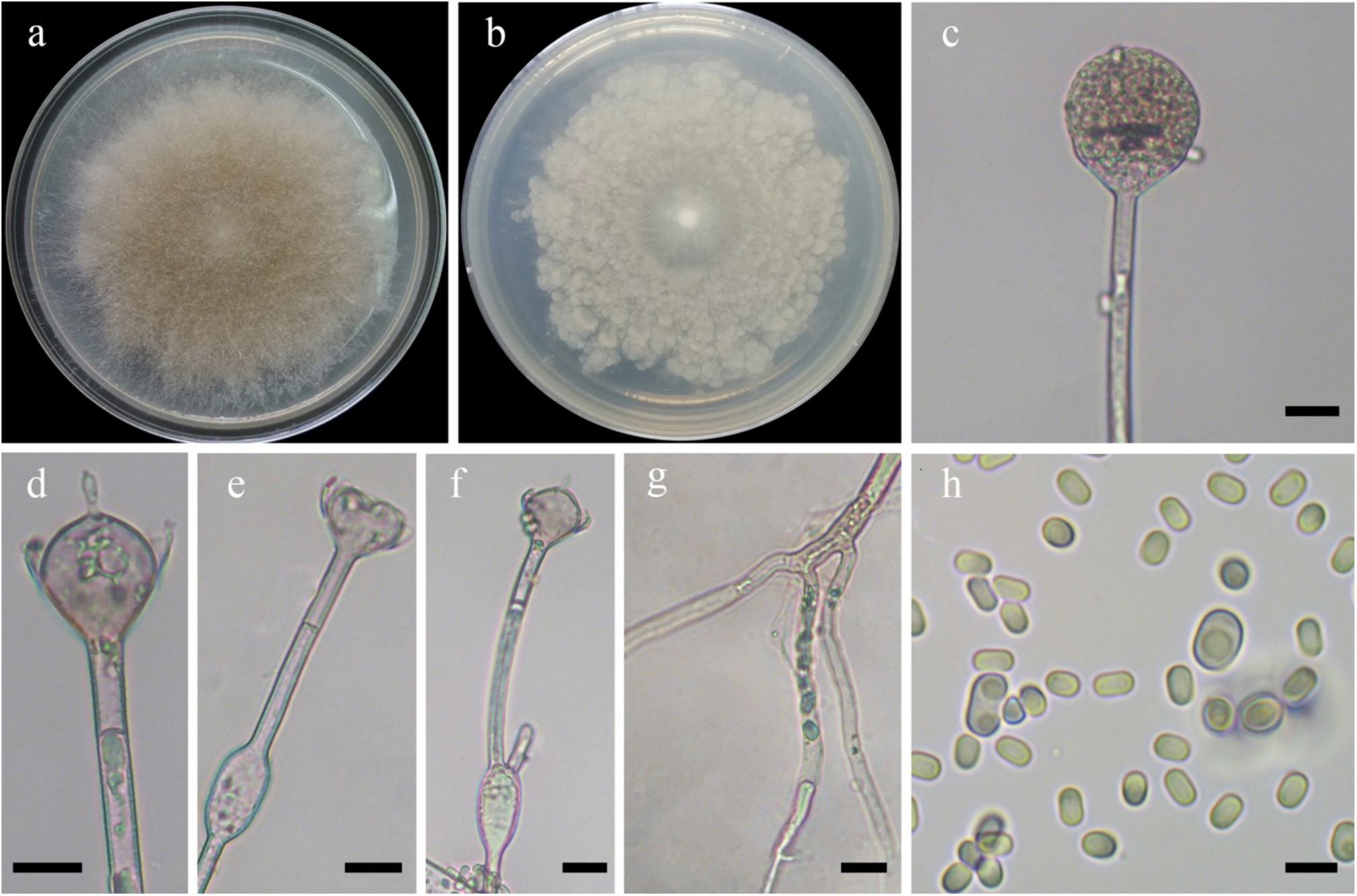
Morphologies of *Absidia brunnea* ex-holotype CGMCC 3.16055. **a, b**. Colonies on MEA (**a.** obverse, **b.** reverse); **c.** Sporangia; **d.** Columellae; **e, f.** Swellings on sporangiophores; **g.** Rhizoids; **h.** Sporangiospores. — Scale bars: **c–g.** 10 μm, **h.** 5 μm.

*Fungal Names*: FN570885.

*Etymology*: *brunnea* (Lat.) refers to the species having brown colonies on MEA.

*Holotype*: HMAS 350296.

*Colonies* on MEA at 27 °C for 10 days, slow growing, reaching 90 mm in diameter, obversely regularly concentrically zonate with ring, white at first and then Sayal Brown or Snuff Brown. *Hyphae* hyaline at first, becoming brown when mature, 6.0–17.5 µm in diameter. *Stolons* branched, hyaline, smooth, septate, 3.5–10.0 µm in diameter. *Rhizoids* rarely branched, with a septum at the base. *Sporangiophores* erect or slightly bent, 1–7 in whorls but mostly 4–6, unbranched, simple branched or monopodial, rarely sympodial, hyaline, with a septum 11.0–17.0 µm below apophyses, sometimes a swelling beneath sporangium, occasionally branched at the swelling, 50.0–420.0 µm long and 3.0–6.5 µm wide. *Sporangia* globose to pyriform, smooth, multi-spored, 17.5–38.0 µm long and 19.0–34.5 µm wide, walls deliquescent. *Apophyses* distinct, funnel-shaped, slightly pigmented, 3.0–10.0 µm high, 2.5–7.5 µm wide at the base, and 10.0–20.0 µm wide at the top. *Collars* distinct. *Columellae* hemispherical, hyaline, smooth, always with a 2.5–6.0 µm clavate projection at the apex, 7.0–18.5 µm long and 10.5–24.0 µm wide. *Sporangiospores* hyaline, smooth, two kinds, cylindrical to ovoid, 3.0–4.0 µm long and 2.0–2.5 µm wide; ovoid with vacuoles, 4.0–7.0 µm long and 3.0–5.0 µm wide. *Zygospores* absent. *Chlamydospores* absent. No growth at 35 °C.

*Material examined*: China, Qinghai Province, Xining, Huangzhong County, from soil sample, 2 August 1999, Long Wang (holotype HMAS 350296, living ex-holotype culture CGMCC 3.16055).

*GenBank accession number*: MZ354139.

*Notes*: *Absidia brunnea* is closely related to *A. soli* V.G. Hurdeal *et al*. based on phylogenetic analysis of ITS rDNA sequences (Fig. 5). However, morphologically *A. soli* differs from *A. brunnea* by sporangiophores up to 6 in whorls, septa distantly below apophyses (21.5–37.5 µm vs. 11.0–17.0 µm), and cylindrical sporangiospores without a vacuole (Hurdeal *et al*. 2021). Moreover, *A. soli* physiologically differs from *A. brunnea* by a slightly higher maximum growth temperature (37 °C vs. 35 °C).

**5. *Absidia chinensis*** T.K. Zong, H. Zhao, Y.C. Dai & X.Y. Liu, sp. nov., Fig. 21.

**Fig. 21.**
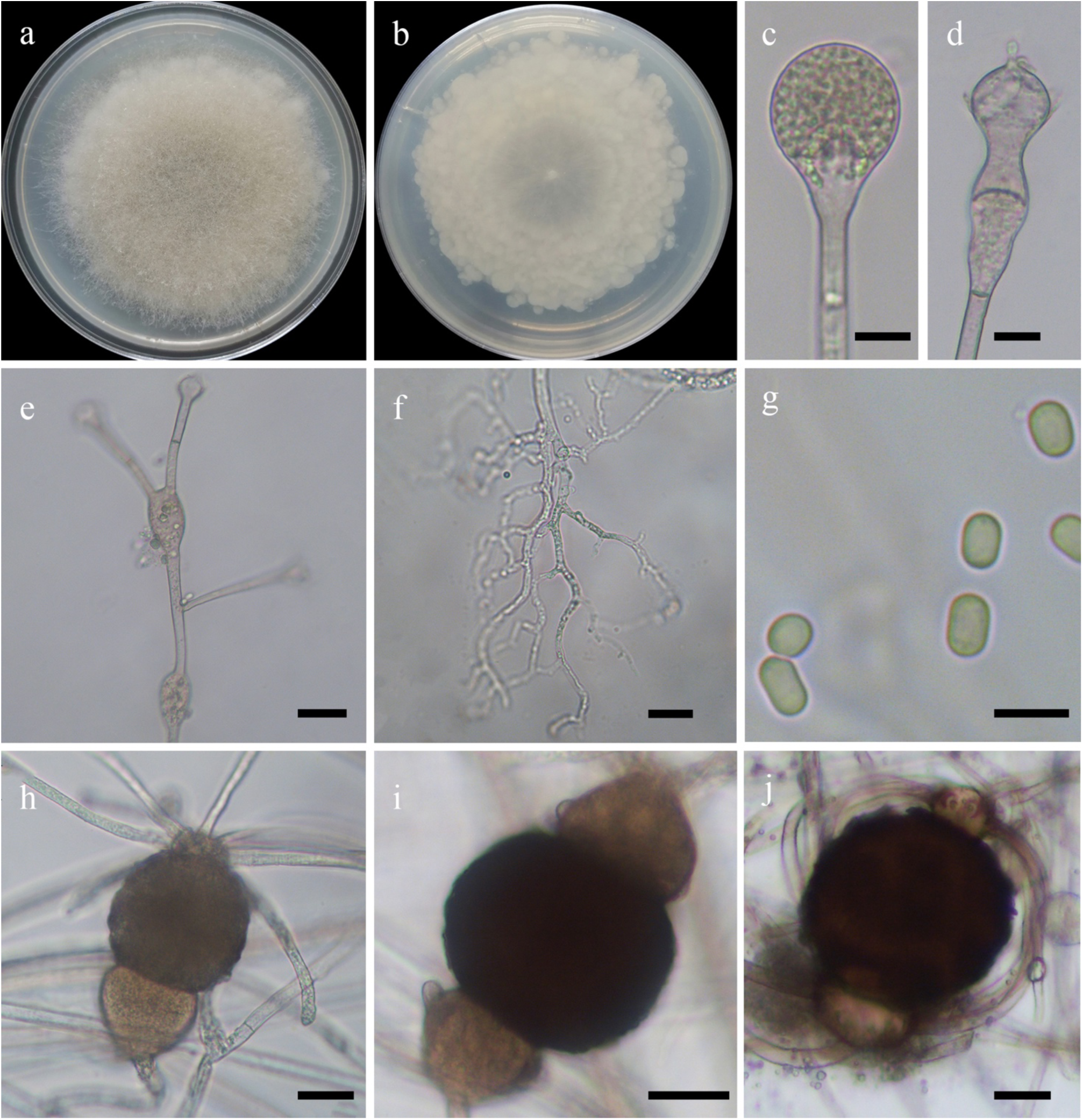
Morphologies of *Absidia chinensis* ex-holotype CGMCC 3.16056. **a, b**. Colonies on MEA (**a.** obverse, **b.** reverse); **c.** Sporangia; **d.** Columellae; **e.** Swellings in sporangiophores; **f.** Rhizoids; **g.** Sporangiospores; **h–j.** Zygospores. — Scale bars: **c, d, f, h–j.** 10 μm, **e.** 20 μm, **g.** 5 μm.

*Fungal Names*: FN570886.

*Etymology*: *chinensis* (Lat.) refers to China where the type was collected.

*Holotype*: HMAS 350297.

*Colonies* on MEA at 27 °C for 9 days, slow growing, reaching 90 mm in diameter, obversely regularly concentrically zonate with ring, white at first and then Drab Gray or Snuff Brown. *Hyphae* hyaline at first, becoming brown when mature, generally swollen, 6.0–15.5 µm in diameter. *Stolons* branched, hyaline, smooth, septate, 3.5–11.0 µm in diameter. *Rhizoids* coralliform, mostly simple or two-branched, with a septum at the base. *Sporangiophores* erect or slightly bent, 1–5 in whorls, mostly unbranched or simple branched, rarely monopodial, sympodial form absent, hyaline, with a septum 13.0–22.0 µm below apophyses, generally a swelling beneath sporangium, sometimes branched at the swelling, 45.0–220.0 µm long and 2.5–6.0 µm wide. *Sporangia* globose or pyriform, smooth, multi-spored, 15.0–39.5 µm long and 15.0–37.5 µm wide, walls deliquescent. *Apophyses* distinct, bell-shaped or hourglass-shaped, slightly pigmented, 3.5–7.0 µm high, 2.5–5.5 µm wide at the base, and 6.0–15.5 µm wide at the top. *Collars* present or absent, but distinct if present. *Columellae* spherical or hemispherical, hyaline, smooth, always with a 2.0–5.5 µm papillary projection at the apex, 5.0–15.5 µm long and 7.5–17.5 µm wide. *Sporangiospores* cylindrical to ovoid, hyaline, smooth, 3.5–4.5 µm long and 2.0–2.5 µm wide. *Zygospores* subglobose, brown or Dark Brown, rough, 37.0–90.0 µm long and 36.0–80.0 µm wide. *Suspensor* unequal, parallel to subparallel, brown, the larger one 27.0–44.0 µm in diameter, the smaller one mostly 20.0–33.0 µm in diameter. *Chlamydospores* absent. No growth at 31 °C.

*Materials examined*: China. Yunnan Province, Jinghong, Mengla County, from soil sample, 4 July 1994, Gui-Qing Chen (holotype HMAS 350297, living ex-holotype culture CGMCC 3.16056). Sichuan Province, Ngawa, Jiuzhaigou County, from soil sample, 23 May 1994, Gui-Qing Chen (living culture CGMCC 3.16057).

*GenBank accession numbers*: MZ354140 and MZ354141.

*Notes*: *Absidia chinensis* is closely related to *A. xinjiangensis* H. Zhao, Y.C. Dai & X.Y. Liu based on phylogenetic analysis of ITS rDNA sequences (Fig. 5). However, morphologically *A. xinjiangensis* differs from *A. chinensis* by the absence of zygospores and projections.

**6. *Absidia cinerea*** T.K. Zong, H. Zhao, Y.C. Dai & X.Y. Liu, sp. nov., Fig. 22.

**Fig. 22.**
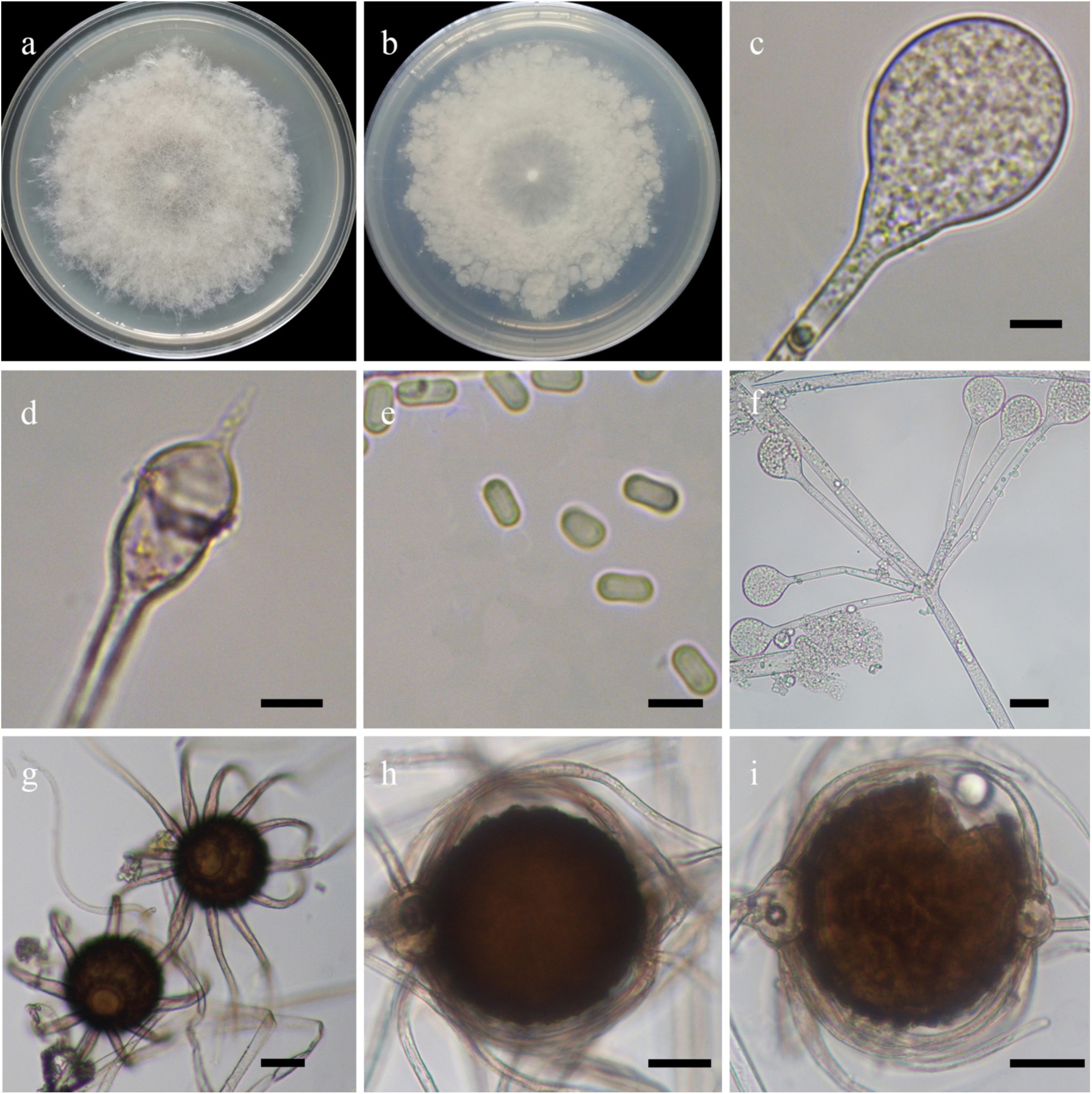
Morphologies of *Absidia cinerea* ex-holotype CGMCC 3.16062. **a, b.** Colonies on MEA (**a.** obverse, **b.** reverse); **c.** Sporangia; **d.** Columellae; **e.** Sporangiospores; **f.** Sporangiophores in whorls; **g–i.** Zygospores. — Scale bars: **c–e.** 5 μm, **f–i.** 20 μm.

*Fungal Names*: FN570887.

*Etymology*: *cinerea* (Lat.) refers to the species having gray colonies on MEA.

*Holotype*: HMAS 350303.

*Colonies* on MEA at 27 °C for 5 days, reaching 90 mm in diameter, regularly concentrically zonate with ring, white at first and then Pale Mouse Gray. *Hyphae* hyaline at first, becoming brown when mature, 5.5–14.5 µm in diameter. *Stolons* branched, hyaline, smooth, septate, 3.5–7.0 µm in diameter. *Rhizoids* branch-shaped and tapering at the end, mostly two branches arising at a place, with a septum at the base. *Sporangiophores* erect or slightly bent, 1–7 in whorls, unbranched, simple branched or monopodial arising from stolons, mostly sympodial arising from aerial mycelia, hyaline, with a septum 11.0–17.0 µm below apophyses, 50.0–150.0 µm long and 2.5–5.0 µm wide. *Sporangia* globose to pyriform, smooth, multi-spored, 7.0–27.5 µm long and 7.0–25.0 µm wide, walls deliquescent. *Apophyses* distinct, bell-shaped or funnel-shaped, slightly pigmented, 2.0–7.5 µm high, 2.5–5.5 µm wide at the base, and 6.0–12.5 µm wide at the top. *Collars* present or absent, but distinct if present. *Columellae* hemispherical, hyaline, smooth, always with a 1.3–5.0 µm rod-shaped or needle-like projection at the apex, 5.0–14.0 µm long and 7.0–15.0 µm wide. *Sporangiospores* cylindrical, slightly constricted in the center, hyaline, smooth, 4.5–6.0 µm long and 2.0–2.5 µm wide. *Zygospores* globose, brown to Dark Brown, rough, 36.0–81.0 µm in diameter. *Suspensor* mostly one, rarely unequal two, brown, with 8–13 appendages on a single suspensor, 17.0–33.0 µm in diameter. *Chlamydospores* absent. No growth at 35 °C.

*Material examined*: China, Beijing, 40°21′30″N, 116°6′27″E, from soil sample, 31 December 2019, Xiao-Yong Liu (holotype HMAS 350303, living ex-holotype culture CGMCC 3.16062).

*GenBank accession number*: MZ354146.

*Notes*: *Absidia cinerea* is closely related to *A. pseudocylindrospora* Hesselt. & J.J. Ellis and *A. stercoraria* Hyang B. Lee *et al*. based on phylogenetic analysis of ITS rDNA sequences (Fig. 5). However, morphologically *A. pseudocylindrospora* differs from *A. cinerea* by shorter cylindrical sporangiospores (2.5 µm long vs. 4.5–6.0 µm long), rhizoids rarely septate, and arising in whorls of up to 11 (Hesseltine & Ellis 1961). *A. stercoraria* differs from *A. cinerea* by rhizoids poorly developed and arising in whorls of up to 5 (Li *et al*. 2016). Moreover, physiologically, *A. pseudocylindrospora* has a maximum growth temperature of 37 °C, but neither *A. cinerea* nor *A. stercoraria* can grow above 35 °C.

**7. *Absidia digitula*** T.K. Zong, H. Zhao, Y.C. Dai & X.Y. Liu, sp. nov., Fig. 23.

**Fig. 23.**
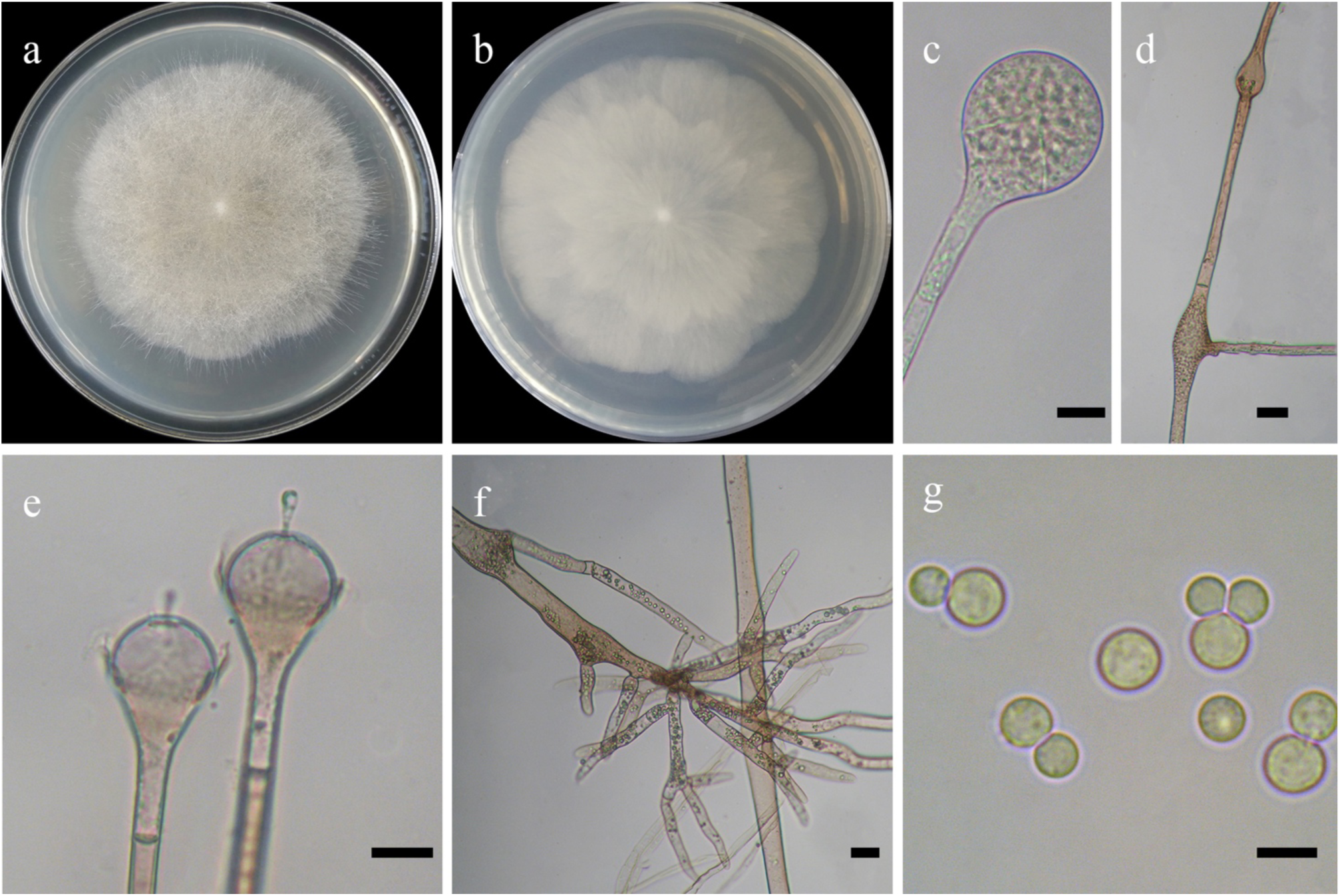
Morphologies of *Absidia digitula* ex-holotype CGMCC 3.16058. **a, b**. Colonies on MEA (**a.** obverse, **b.** reverse); **c.** Sporangia; **d.** Swellings in hyphae; **e.** Columellae; **f.** Rhizoids; **g.** Sporangiospores. — Scale bars: **c, e.** 10 μm, **d, f.** 20 μm, **g.** 5 μm.

*Fungal Names*: FN570888.

*Etymology*: *digitula* (Lat.) refers to the species having finger-like rhizoids.

*Holotype*: HMAS 350299.

*Colonies* on MEA at 27 °C for 9 days, slow growing, reaching 90 mm in diameter regularly flower-shaped zonate, white at first and then becoming Snuff Brown. *Hyphae* hyaline at first, becoming brown when mature, sometimes swollen, 8.0–18.0 µm in diameter. *Stolons* branched, hyaline or Light Brown, smooth, septate, 4.0–9.5 µm in diameter. *Rhizoids* finger-like, mostly twice or more repeatedly, with a septum at the base. *Sporangiophores* erect or slightly bent, 1–6 in whorls, unbranched, not generally simple branched, each sporangiophore in whorls, rarely monopodial or sympodial, hyaline or Light Brown, with a septum 14.5–25.5 µm below apophyses, occasionally a swelling beneath sporangium, 60.08–470.0 µm long and 3.5–8.5 µm wide. *Sporangia* globose to pyriform, smooth, multi-spored, 24.0–64.0 µm long and 26.5–48.5 µm wide, walls deliquescent. *Apophyses* distinct, slightly pigmented, 5.5–13.5 µm high, 4.5–8.5 µm wide at the base, and 12.5–21.0 µm wide at the top. *Collars* distinct if present. *Columellae* hemispherical, hyaline, smooth, sometimes with a 4.0–9.0 µm clavate projection at the apex, occasionally with two projections, slightly bulbous at the end, 10.5–22.0 µm long and 11.5–30.0 µm wide. *Sporangiospores* globose, hyaline, smooth, non-uniform in shape, 3.0–5.0 µm in diameter. *Zygospores* absent. *Chlamydospores* absent. No growth at 32 °C.

*Material examined*: China, Xinjiang Auto Region, Ili, Qapqal Xibe Country, Wusun Mountain, from soil sample, 14 June 2002, Xue-Wei Wang (holotype HMAS 350299, living ex-holotype culture CGMCC 3.16058).

*GenBank accession number*: MZ354142.

*Notes*: *Absidia digitula* is closely related to *A. turgida* T.K. Zong & X.Y. Liu based on phylogenetic analysis of ITS rDNA sequences (Fig. 5). However, morphologically *A. turgida* differs from *A. digitula* by sporangiophores singly or in whorls of up to 4, one projection on the columellae only, and variably shaped sporangiospores such as globose, cylindrical, or irregular (Zong *et al*. 2021).

**8. *Absidia jiangxiensis*** H. Zhao, T.K. Zong, Y.C. Dai & X.Y. Liu, sp. nov., Fig. 24.

**Fig. 24.**
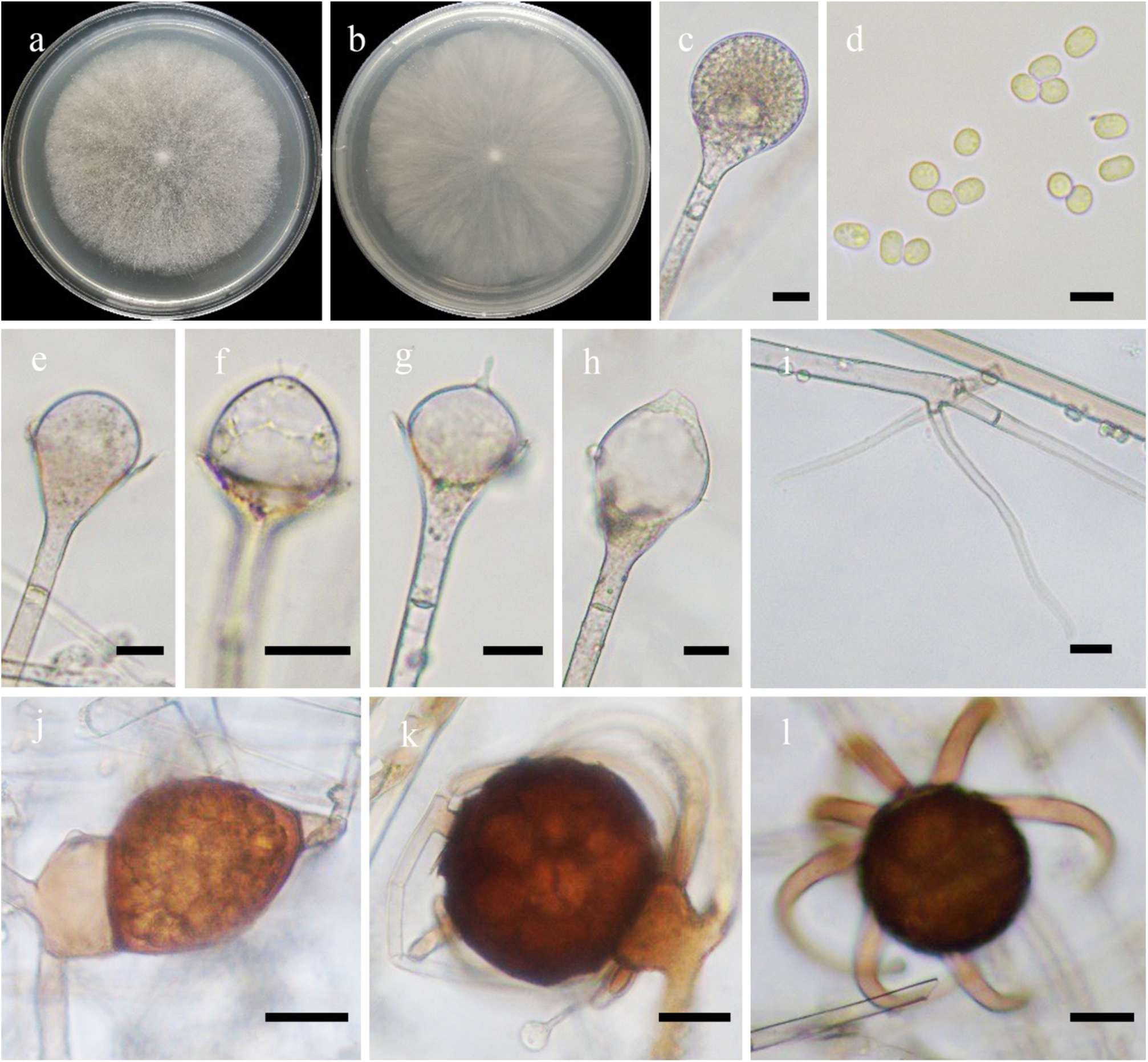
Morphologies of *Absidia jiangxiensis* ex-holotype CGMCC 3.16105. **a, b**. Colonies on MEA (**a.** obverse, **b.** reverse); **c.** Sporangia; **d.** Sporangiospores; **e–h.** Columellae; **i.** Rhizoids; **j–l.** Zygospores. — Scale bars: **c, e–i.** 10 μm, **d.** 5 μm, **j–l.** 20 μm.

*Fungal Names*: FN570889.

*Etymology*: *jiangxiensis* (Lat.) refers to the locality of Jiangxi Province, China, where the type was collected.

*Holotype*: HMAS 351507.

*Colonies* on MEA at 27 °C for 5 days, reaching 68 mm in diameter, higher in the centre than at margin, white at first becoming Dark Citrine when mature. *Hyphae* hyaline at first, becoming Light Brown when mature, 7.0–17.0 µm in diameter. *Stolons* branched, hyaline to Light Brown, smooth, septate, 3.5–8.5 µm in diameter. *Rhizoids* root-like, rarely branched, hyaline, septate. *Sporangiophores* erect or slightly bent, 1–6 in whorls, unbranched, simple, monopodial or sympodial, hyaline to Light Brown, with a septum 10.5–22.5 µm below apophyses, 25.0–280.0 µm long and 3.0–6.0 µm wide. *Sporangia* globose to pyriform, smooth, multi-spored, 16.5–48.0 µm long and 16.5–44.0 µm wide, walls deliquescent. *Apophyses* distinct, funnel-shaped, slightly greenish, 6.0–15.0 µm high, 3.0–7.5 µm wide at the base, and 10.0–20.0 µm wide at the top. *Collars* distinct if present. *Columellae* subspherical to hemispherical, sometimes conical, hyaline, smooth, 10.0–30.0 µm long and 10.5–34.0 µm wide, one projection present on apex of columellae, occasionally two, papillary, rod-like or blunt, 2.5–6.5 µm long. *Sporangiospores* ovoid to cylindrical, sometimes globose to subglobose, hyaline, smooth, uniform, 3.5–4.5 µm long and 2.5– 3.0 µm wide. *Zygospores* globose, brown to Dark Brown, rough, 40.0–85.0 µm in diameter. *Suspensor* mostly one, sometimes two, particularly unequal, hemispherical, brown, 17.0–36.0 µm in diameter. *Appendages* derived from larger one. Homothallic. *Chlamydospores* absent. No growth at 31 °C.

*Material examined*: China, Jiangxi Province, Lushan, from soil sample, 1 October 2013, Xiao-Yong Liu (holotype HMAS 351507, living ex-holotype culture CGMCC 3.16105).

*GenBank accession number*: OL678134.

*Notes*: *Absidia jiangxiensis* is closely related to *A. jindoensis* Hyang B. Lee & T.T.T. Nguyen based on phylogenetic analysis of ITS rDNA sequences (Fig. 5). However, morphologically *A. jindoensis* differs from *Absidia jiangxiensis* by hemispherical, sometimes subglobose columellae with one projection and a collarette (Wanasinghe *et al*. 2018).

**9. *Absidia nigra*** (Hesselt. & J.J. Ellis) T.K. Zong, H. Zhao, Y.C. Dai & X.Y. Liu, comb. et stat. nov., Fig. 25.

**Fig. 25.**
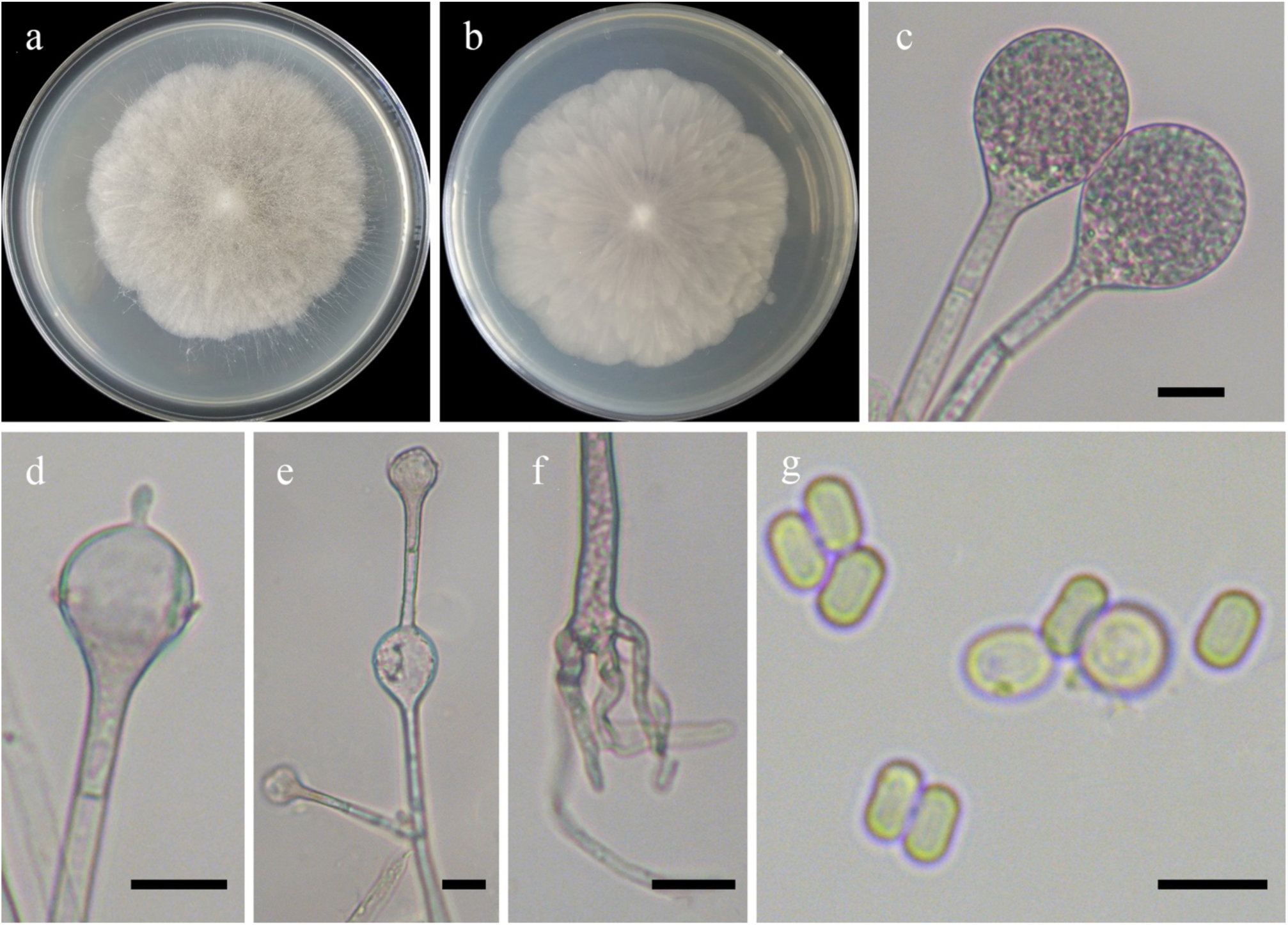
Morphologies of *Absidia nigra* ex-holotype CGMCC 3.16059. **a, b**. Colonies on MEA (**a.** obverse, **b.** reverse); **c.** Sporangia; **d.** Columellae; **e.** Swellings in sporangiophores; **f.** Rhizoids; **g.** Sporangiospores. — Scale bars: **c–f.**10 μm, **g.** 5 μm.

*Fungal Names*: FN570890.

Basionym: *Absidia cylindrospora* var. *nigra* Hesselt. & J.J. Ellis, Mycologia, 56(4): 595, 1964. [MB no.: 353238]

*Holotype*: NRRL 3060.

*Colonies* on MEA at 27 °C for 9 days, slow growing, reaching 62 mm in diameter, regularly zonate, white at first becoming Light Drab. *Hyphae* hyaline at first, becoming brown when mature, sometimes swollen, 8.0–18.5 µm in diameter. *Stolons* branched, hyaline, smooth, septate, 4.0–8.5 µm in diameter. *Rhizoids* finger-like, short, mostly twice or repeatedly, with a septum at the base. *Sporangiophores* erect or slightly bent, 1–5 in whorls, unbranched, simple, rarely monopodial or sympodial, hyaline, with a septum 11.5–21.0 µm below apophyses, sometimes a swelling beneath sporangium, 60.0–380.0 µm long and 2.5–6.5 µm wide. *Sporangia* globose to pyriform, smooth, multi-spored, 17.0–46.5 µm long and 16.5–38.5 µm wide, walls deliquescent. *Apophyses* distinct, slightly greenish, 3.5–8.5 µm high, 3.3–5.9 µm wide at the base, and 9.5–18.5 µm wide at the top. *Collars* distinct if present. *Columellae* spherical or ovoid to hemispherical, hyaline, smooth, sometimes with a 1.5–5.0 µm clavate or slightly swollen projection at the apex, 8.5–19.0 µm long and 9.5–23.0 µm wide. *Sporangiospores* two types: cylindrical hyaline, smooth, 3.0–4.0 µm long and 2.0–2.5 µm wide and globose, 4.0–9.0 µm in diameter. *Zygospores* absent. *Chlamydospores* absent. No growth at 31 °C.

*Materials examined*: USA, Wisconsin, from soil sample, 1940 (holotype NRRL 3060, isotype CBS 127.68, isotype HMAS 350309). China. Jilin Province, Jiaohe, from soil sample, 1 September 1991, Feng-Yan Bai (HMAS 350300, living culture CGMCC 3.16059); Inner Mongolia Auto Region, Arxan, from soil sample, 16 August 1991, Feng-Yan Bai (HMAS 350301, living culture CGMCC 3.16060).

*GenBank accession numbers*: MZ354143, MZ354144 and MZ354152

*Notes*: *Absidia nigra* was previously synonymized with *A. cylindrospora* Hagem (Hesseltine & Ellis 1964), however, this does not receive any support either phylogenetically or morphologically (Fig. 5). Phylogenetically, *A. nigra* is closer to *A. chinensis* rather than *A. cylindrospora*. Physiologically, strains of *A. nigra* and *A. cylindrospora* are not compatible (Hesseltine and Ellis 1964). Morphologically, *A. cylindrospora* differs from *A. nigra* by producing cylindrical sporangiospores only, while there are two types of sporangiospores in *A. nigra* (Hesseltine & Ellis 1964).

**10. *Absidia oblongispora*** T.K. Zong, H. Zhao, Y.C. Dai & X.Y. Liu, sp. nov., Fig. 26.

**Fig. 26.**
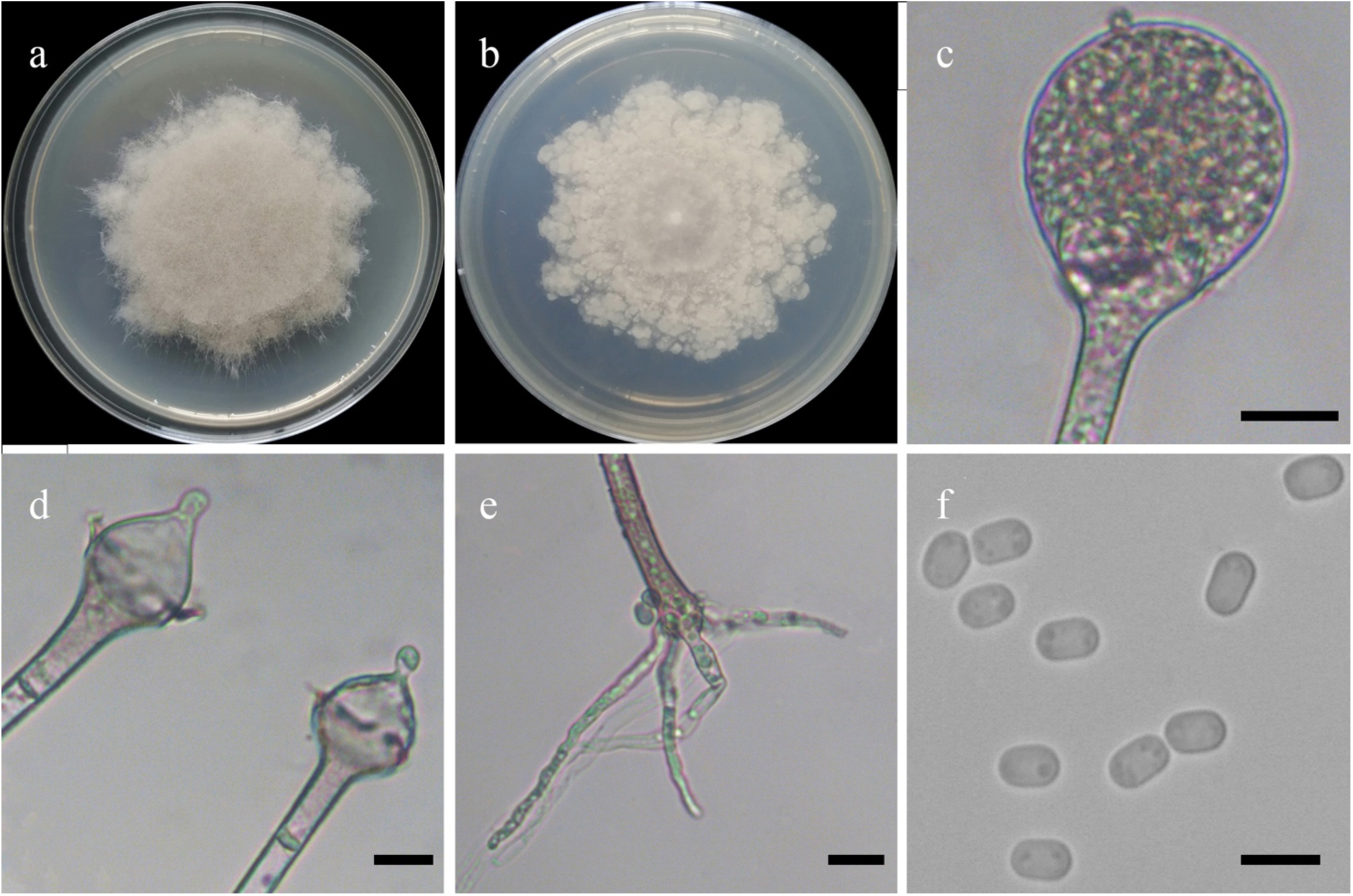
Morphologies of *Absidia oblongispora* ex-holotype CGMCC 3.16061. **a, b**. Colonies on MEA (**a.** obverse, **b.** reverse); **c.** Sporangia; **d.** Columellae; **e.** Rhizoids; **f.** Sporangiospores. — Scale bars: **c–e.** 10 μm, **f.** 5 μm.

*Fungal Names*: FN570891.

*Etymology*: *oblongispora* (Lat.) refers to the species having oblong sporangiospores.

*Holotype*: HMAS 350302.

*Colonies* on MEA at 27 °C for 10 days, slow growing, reaching 90 mm in diameter, slightly irregularly concentrically zonate with ring, white at first and then becoming Pale Mouse Gray to Light Drab. *Hyphae* hyaline at first, becoming brown when mature, 5.5–11.5 µm in diameter. *Stolons* branched, hyaline, smooth, with few septa near the base of sporangiophores, 4.0–9.5 µm in diameter. *Rhizoids* root-like, branched repeatedly, with a septum at the base. *Sporangiophores* erect or slightly bent, 1–5 in whorls, mostly unbranched, rarely simple or monopodial, sympodial form absent, hyaline, with a septum 9.5–16.0 µm below apophyses, occasionally with a septum at the base, 33.0–300.0 µm long and 3.0–5.5 µm wide. *Sporangia* globose, smooth, multi-spored, 13.5– 31.0 µm in diameter, walls deliquescent. *Apophyses* distinct, slightly greenish, 3.5–6.5 µm high, 3.5–7.5 µm wide at the base, and 7.0–19.0 µm wide at the top. *Collars* present or absent, but distinct if present. *Columellae* mostly conical, rarely hemispherical, hyaline, smooth, always with a 3.0–7.5 µm bulbous projection at the apex, 7.0–15.0 µm long and 8.5–16.5 µm wide. *Sporangiospores* oblong, hyaline, smooth, uniform, 3.5–4.5 µm long and 2.5–3.0 µm wide. *Zygospores* absent. *Chlamydospores* absent. No growth at 32 °C.

*Material examined*: China, Yunnan Province, Jinghong, Mengla County, from soil sample, 4 July 1994, Gui-Qing Chen (holotype HMAS 350302, living ex-holotype culture CGMCC 3.16061).

*GenBank accession number*: MZ354145.

*Notes*: *Absidia oblongispora* is closely related to *A. heterospora* Y. Ling based on phylogenetic analysis of ITS rDNA sequences (Fig. 5). However, morphologically *A. heterospora* differs from *A. oblongispora* by wider columellae (10.5–34 µm vs. 8.5–16.5 µm in diameter) and two types of sporangiospores (Hesseltine & Ellis 1964).

**11. *Absidia purpurea*** H. Zhao, T.K. Zong, Y.C. Dai & X.Y. Liu, sp. nov., Fig. 27.

**Fig. 27.**
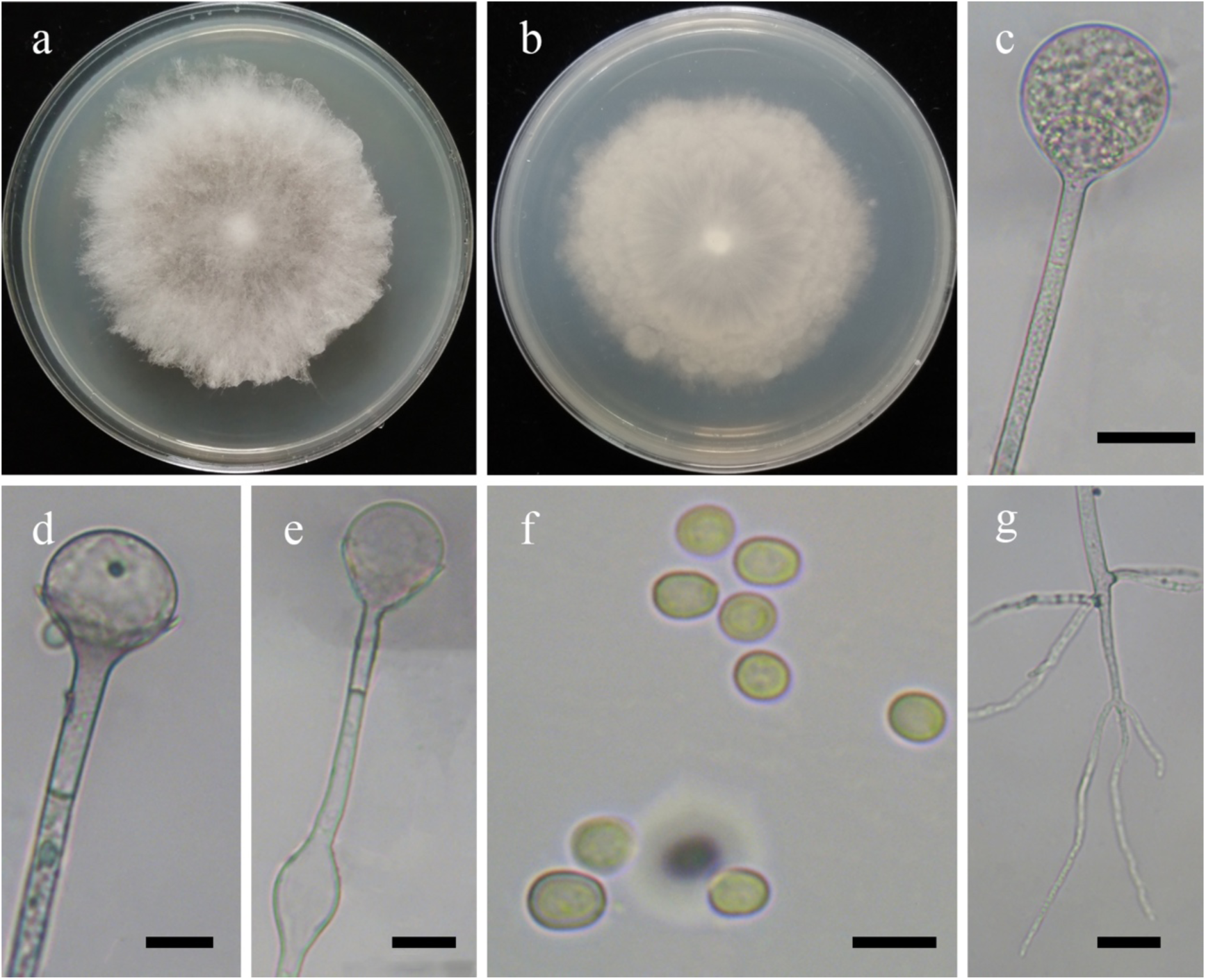
Morphologies of *Absidia purpurea* ex-holotype CGMCC 3.16106. **a, b**. Colonies on MEA (**a.** obverse, **b.** reverse); **c.** Sporangia; **d, e.** Columellae; **f.** Sporangiospores; **g.** Rhizoids. — Scale bars: **c, g.** 20 μm, **d, e.** 10 μm, **f.** 5 μm.

*Fungal Names*: FN570892.

*Etymology*: *purpurea* (Lat.), refers to the species having pale purple colonies, hyphae and stolons.

*Holotype*: HMAS 351508.

*Colonies* on MEA at 27 °C for 8 days, slow growing, reaching 90 mm in diameter, regularly concentrically zonate with ring, higher at margin than in center, white at first and then becoming Pale Quaker Drab. *Hyphae* hyaline at first, becoming violet-blue when mature, 5.0–15.5 µm in diameter. *Stolons* branched, hyaline, smooth, septate, 3.5–6.5 µm in diameter. *Rhizoids* root-like, tapering at the end, septate. *Sporangiophores* erect or slightly bent, 1–5 in whorls, unbranched, simple or monopodial, never sympodial, hyaline, with a septum 14.0–28.5 µm below apophyses, 75.0–285.0 µm long and 3.5–5.5 µm wide. *Sporangia* mostly globose, occasionally pyriform, smooth, multi-spored, 19.0–42.0 µm in diameter, walls deliquescent. *Collars* distinct if present. *Apophyses* distinct, funnel-shaped, slightly greenish, 4.5–6.0 µm high, 3.0–5.5 µm wide at the base, and 9.5–15.0 µm wide at the top. *Columellae* subspherical to hemispherical, hyaline, smooth, without projection, 9.0–24.0 µm long and 13.0–24.0 µm wide. *Sporangiospores* ovoid or cylindrical, hyaline, smooth, 3.0–5.0 µm long and 2.5–4.0 µm wide. *Zygospores* absent. *Chlamydospores* absent. No growth at 30 °C.

*Material examined*: China, Yunnan Province, Lincang, from soil sample, 22 April 2021, Heng Zhao (holotype HMAS 351508, living ex-holotype culture CGMCC 3.16106).

*GenBank accession number*: OL678135.

*Notes*: *Absidia purpurea* is closely related to *A. multispora* T.R.L. Cordeiro *et al*. based on phylogenetic analysis of ITS rDNA sequences (Fig. 5). However, morphologically *A. multispora* differs from *A. purpurea* by producing projections and abortive sporangia, sporangiospores variably shaped, including globose, subglobose, ellipsoid, cylindrical, short-cylindrical, and even irregular, and branched (single or in whorls of 2(–4) vs. 1–5 in whorls; Cordeiro *et al*. 2020).

**12. *Absidia sympodialis*** T.K. Zong, H. Zhao, Y.C. Dai & X.Y. Liu, sp. nov., Fig. 28.

**Fig. 28.**
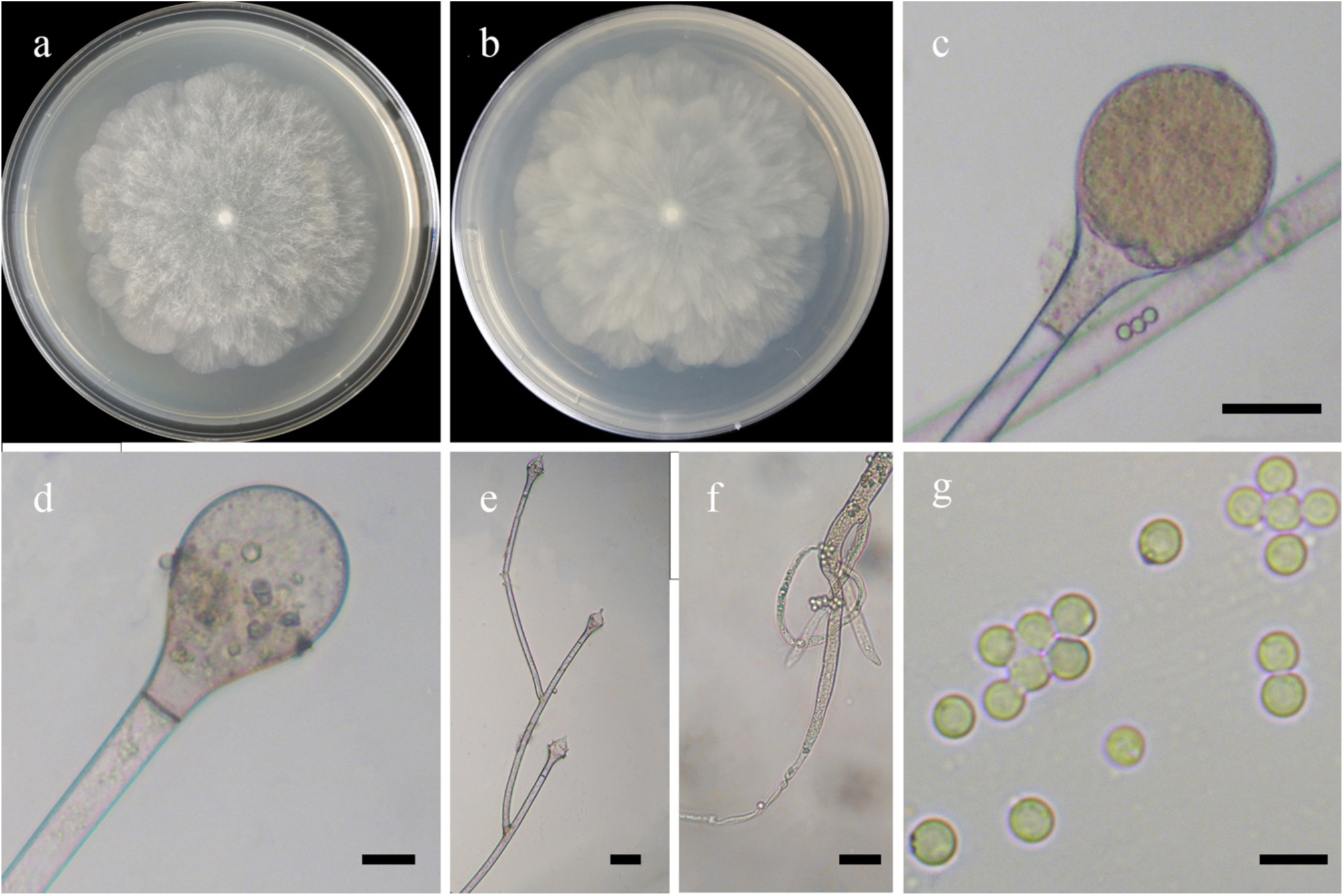
Morphologies of *Absidia sympodialis* ex-holotype CGMCC 3.16064. **a, b**. Colonies on MEA (**a.** obverse, **b.** reverse); **c.** Sporangia; **d.** Columellae; **e.** Sympodial sporangiophores; **f.** Rhizoids; **g.** Sporangiospores. — Scale bars: **c, e, f.** 20 μm, **d.** 10 μm, **g.** 5 μm.

*Fungal Names*: FN570893.

*Etymology*: *sympodialis* (Lat.) refers to the species having sympodial sporangiophores.

*Holotype*: HMAS 350305.

*Colonies* on MEA at 27 °C for 6 days, reaching 90 mm in diameter, regularly zonate, white at first and then Cedar Green. *Hyphae* hyaline at first, becoming Light Brown when mature, 9.0–19.0 µm in diameter. *Stolons* branched, hyaline, smooth, with few septa near the base of sporangiophores, 6.5–12.5 µm in diameter. *Rhizoids* fibrous-root-shaped, tapering at the end, branched mostly repeatedly, with a septum at the base. *Sporangiophores* erect or slightly bent, 1–6 in whorls, unbranched, simple, monopodial or sympodial, hyaline or Light Brown, with a septum 7.5–17.5 µm below apophyses, 90.0–630.0 µm long and 4.5–10.5 µm wide. *Sporangia* globose to pyriform, smooth, multi-spored, 20.0–63.0 µm long and 18.5–63.0 µm wide, walls deliquescent. *Apophyses* distinct, slightly pigmented, 6.0–13.7 µm high, 6.0–14.1 µm wide at the base, and 12.0–31.0 µm wide at the top. *Collars* present or absent, but distinct if present. *Columellae* spherical, ovoid to hemispherical or chestnut-shaped, hyaline, smooth, sometimes with a 2.0–4.0 µm papillary or pointed projection at the apex, 8.0–40.0 µm long and 9.5–39.0 µm wide. *Sporangiospores* globose, hyaline, smooth, uniform, 2.5–3.5 µm in diameter. *Zygospores* absent. *Chlamydospores* absent. No growth at 33 °C.

*Materials examined*: China. Shaanxi Province, Hanzhong, Foping County, from soil sample, 13 October 2002, Xue-Wei Wang (holotype HMAS 350305, living ex-holotype culture CGMCC 3.16064). Beijing, 40°1′32″N, 116°8′54″E, from soil sample, 31 December 2019, Xiao-Yong Liu (living culture CGMCC 3.16063).

*GenBank accession numbers*: MZ354148 and MZ354147.

*Notes*: *Absidia sympodialis* is closely related to *A. virescens* T.K. Zong *et al*. based on phylogenetic analysis of ITS rDNA sequences (Fig. 5). However, morphologically *A. virescens* differs from *A. sympodialis* by root-like rhizoids, hemispherical columellae, and whorls of sporangiophores (6 vs. 4). In addition, *A. virescens* produces sporangiospores which are non-uniform in size and columellae with two projections, some projections up to 6.5 µm in length, whereas the sporangiospores of the new species are uniform in size, and both columellae have only one projection less than 5.0 µm in length.

**13. *Absidia varians*** T.K. Zong, H. Zhao, Y.C. Dai & X.Y. Liu, sp. nov., Fig. 29.

**Fig. 29.**
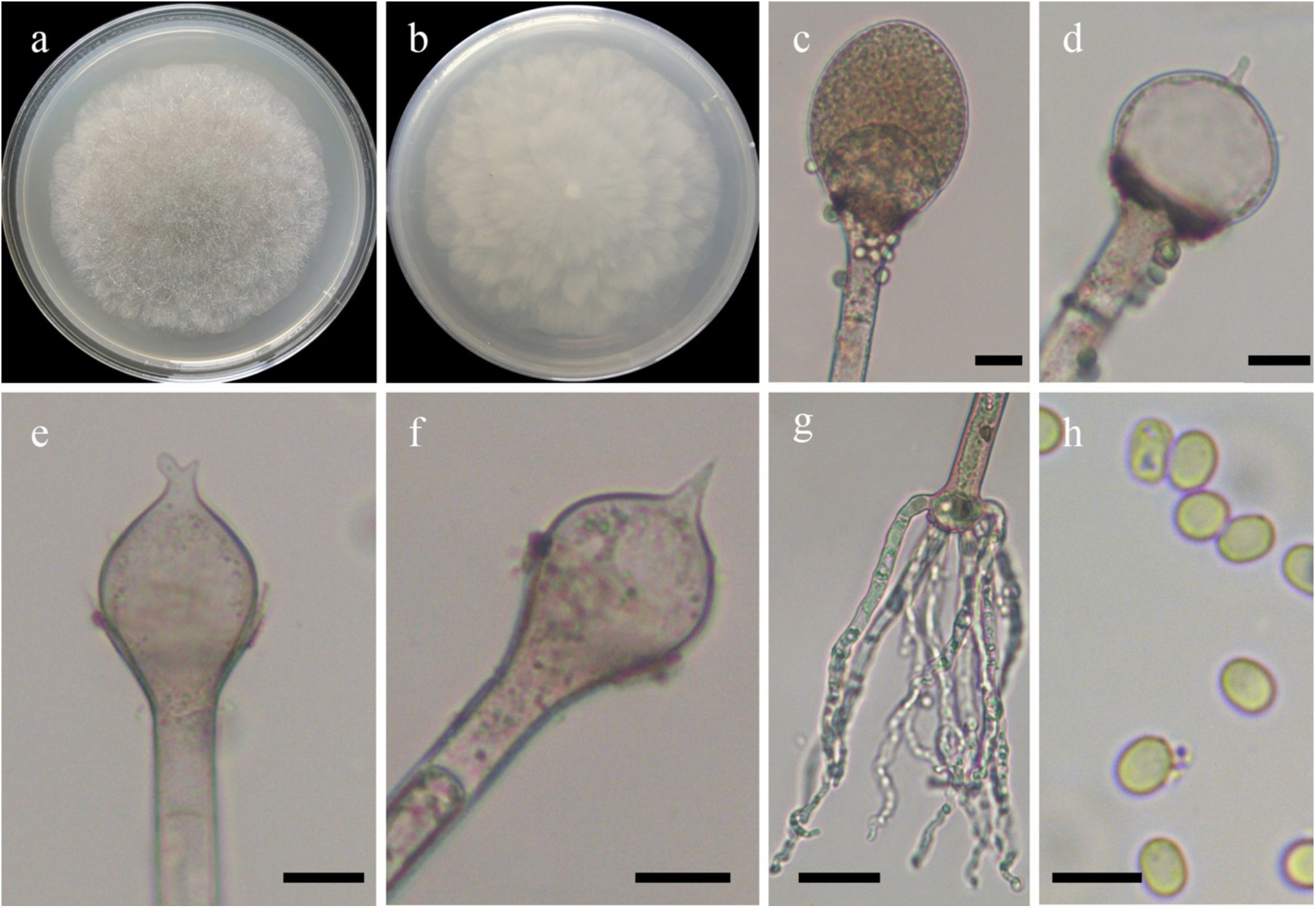
Morphologies of *Absidia varians* ex-holotype CGMCC 3.16065. **a, b**. Colonies on MEA (**a.** obverse, **b.** reverse); **c.** Sporangia; **d–f.** Columellae; **g.** Rhizoids; **h.** Sporangiospores. — Scale bars: **c–f.** 10 μm, **g.** 20 μm, **h.** 5 μm.

*Fungal Names*: FN570894.

*Etymology*: *varians* (Lat.) refers to the species having various projections on columellae.

*Holotype*: HMAS 350306.

*Colonies* on MEA at 27 °C for 5 days, slow growing, reaching 90 mm in diameter, regularly zonate, white at first and then Pale Green-Blue Gray. *Hyphae* hyaline to slightly brownish, 8.5–12.0 µm in diameter. *Stolons* branched, hyaline, smooth, septate, 5.0–8.5 µm in diameter. *Rhizoids* root-like, multi-branched, aseptate. *Sporangiophores* erect or slightly bent, 1–4 in whorls but mostly 2–3 borne from stolons, unbranched, simple, monopodial or sympodial, hyaline, with a septum 13.5–19.0 µm below apophyses, 100.0–480.0 µm long and 4.5–9.0 µm wide. *Sporangia* subglobose to ellipsoid, smooth, multi-spored, 26.0–58.0 µm long and 22.5–50.0 µm wide, walls deliquescent. *Apophyses* distinct, 6.0–12.0 µm high, 5.5–11.0 µm wide at the base, and 10.5–22.5 µm wide at the top. *Collars* distinct if present. *Columellae* spherical or hemispherical, hyaline, smooth, always with several kinds of projections at the apex, mostly one, occasionally two, bulbous swelling, papillary, clavate or spinous, 11.0–30.5 µm long and 12.5–35.0 µm wide. *Sporangiospores* globose, ovoid, short cylindrical, hyaline, smooth, 3.0–4.0 µm long and 2.0–3.0 µm wide. *Zygospores* absent. *Chlamydospores* absent. No growth at 29 °C.

*Material examined*: China, Yunnan Province, Dehong, Mangshi, 24°32′36″N, 98°40′29″E, from soil sample, 30 April 2021, Heng Zhao (holotype HMAS 350306, living ex-holotype culture CGMCC 3.16065).

*GenBank accession number*: MZ354149.

*Notes*: *Absidia varians* is closely related to *A. glauca* Hagem based on phylogenetic analysis of ITS rDNA sequences (Fig. 5). However, morphologically *A. glauca* differs from *A. varians* by globose sporangiospores (Ellis & Hesseltine 1965, Schipper 1990). Besides, *A. varians* is morphologically similar to *A. repens* Tiegh. by sharing different types of projections on the columellae. However, *A. repens* differs from *A. varians* by olive-gray colonies and two types of sporangiospores (Hesseltine & Ellis 1966).

**14. *Absidia virescens*** T.K. Zong, H. Zhao, Y.C. Dai & X.Y. Liu, sp. nov., Fig. 30.

**Fig. 30.**
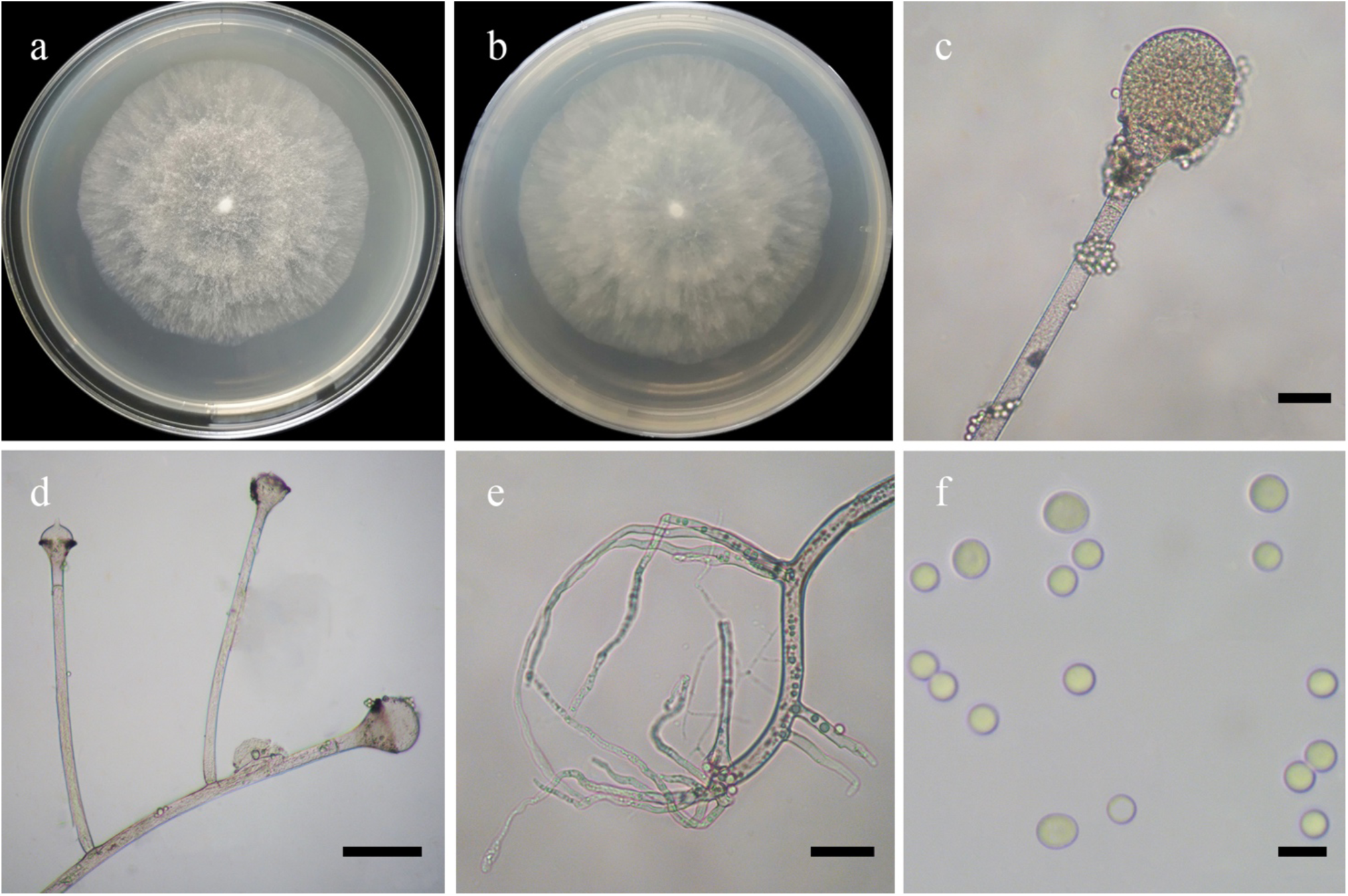
Morphologies of *Absidia virescens* ex-holotype CGMCC 3.16067. **a, b**. Colonies on MEA (**a.** obverse, **b.** reverse); **c.** Sporangia; **d.** Monopodial sporangiophores with columellae; **e.** Rhizoids; **f.** Sporangiospores. — Scale bars: **c, e.** 10 μm, **d.** 20 μm, **f.** 5 μm.

*Fungal Names*: FN570895.

*Etymology*: *virescens* (Lat.) refers to the species having a greenish colony on MEA.

*Holotype*: HMAS 350307.

*Colonies* on MEA at 27 °C for 6 days, reaching 90 mm in diameter, regularly disc-shaped, white at first and then Ring Elm Green. *Hyphae* hyaline at first, becoming brown when mature, 6.0–11.5 µm in diameter. *Stolons* branched, hyaline, smooth, with few septa near the base of sporangiophores, 4.5–9.0 µm in diameter. *Rhizoids* root-like, branched mostly twice or three times, with a septum at the base. *Sporangiophores* erect or slightly bent, 1–4 in whorls, mostly unbranched or simple, rarely monopodial or sympodial, hyaline, with a septum 10.0–22.5 µm below apophyses, occasionally with a septum at the base, sometimes a swelling beneath sporangium, 75.0–480.0 µm long and 5.5–11.5 µm wide. *Sporangia* globose to pyriform, smooth, multi-spored, 15.5–51.5 µm long and 15.5–45.5 µm wide, walls deliquescent. *Apophyses* distinct, sometimes slightly greenish, 5.5–14.0 µm high, 6.5––15.5 µm wide at the base, and 12.5–25.5 µm wide at the top. *Collars* present or absent, but distinct if present. *Columellae* spherical or ovoid to hemispherical, hyaline, smooth, sometimes with one or two 3.0–6.5 µm projections at the apex, 10.5–40.5 µm long and 11.5–45.0 µm wide. *Sporangiospores* mostly globose, occasionally subglobose, hyaline, smooth, 3.0–5.5 µm in diameter, but mostly 3.0–3.5 µm in diameter. *Zygospores* absent. *Chlamydospores* absent. No growth at 32 °C.

*Materials examined*: China, Yunnan Province. Xishuangbanna, from soil sample, 9 July 1994, Gui-Qing Chen (holotype HMAS 350307, living ex-holotype culture CGMCC 3.16067). Kumming, from soil sample, 27 August 1995, Ying-Lan Guo (living culture CGMCC 3.16066).

*GenBank accession numbers*: MZ354151 and MZ354150.

*Notes*: See notes of *Absidia sympodialis*.

**15. *Absidia xinjiangensis*** H. Zhao, Y.C. Dai & X.Y. Liu, sp. nov., Fig. 31.

**Fig. 31.**
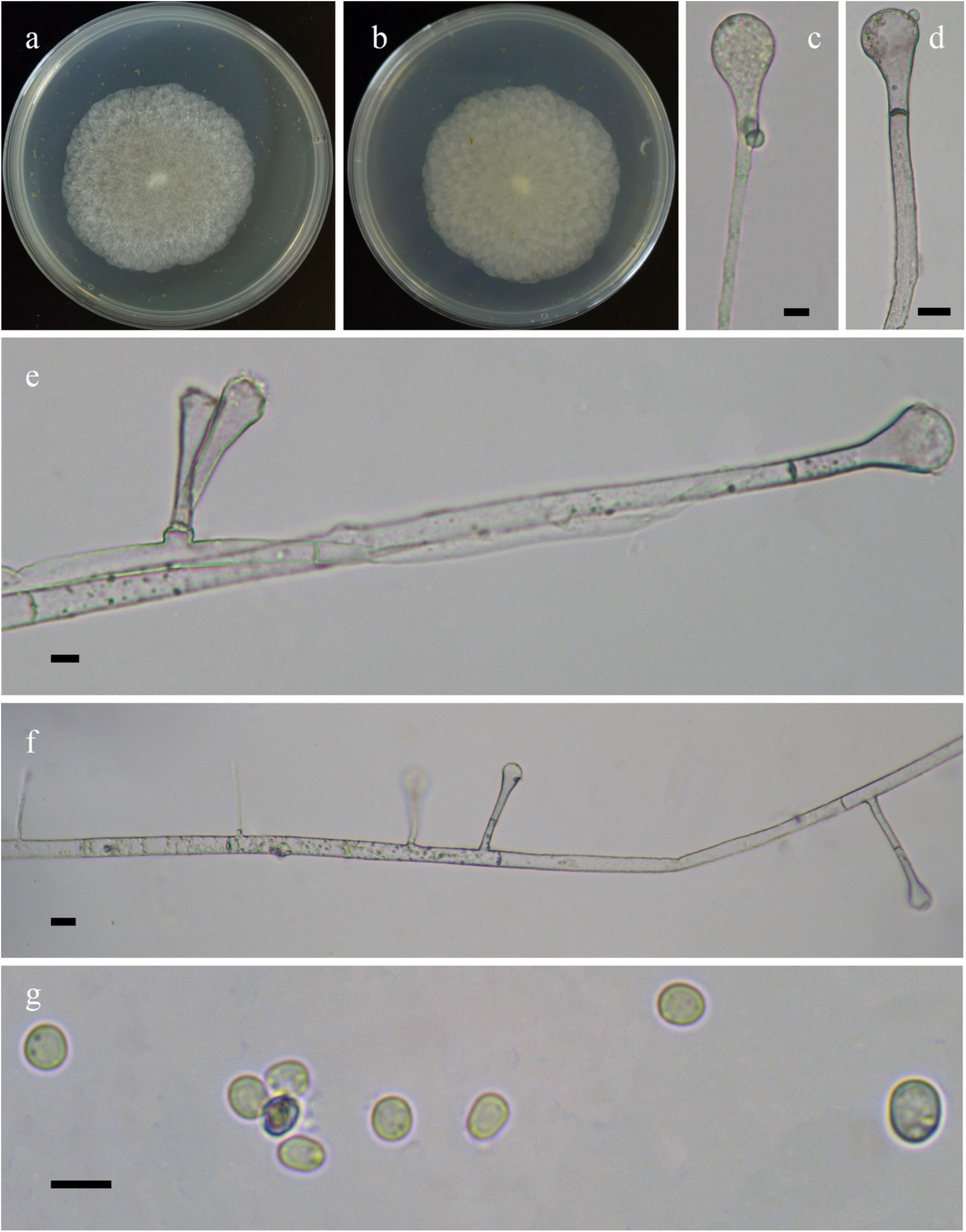
Morphologies of *Absidia xinjiangensis* ex-holotype CGMCC 3.16107 **a, b**. Colonies on MEA (**a.** obverse, **b.** reverse); **c.** Sporangia; **d, e.** Columellae; **f.** Monopodial sporangiophores; **g.** Sporangiospores. — Scale bars: **c–e, g.** 5 μm, **f.** 10 μm.

*Fungal Names*: FN570896.

*Etymology*: *xinjiangensis* (Lat.) refers to the locality of Xinjiang Auto Region, China, where the type was collected.

*Holotype*: HMAS 351509.

*Colonies* on MEA at 27 °C for 7 days, slow growing, reaching 45 mm in diameter, irregular concentrically zonate with ring, white at first and then gradually becoming Light Brown. *Hyphae* hyaline at first, brown when mature, aseptate when juvenile, septate with age, 2.5–13.5 µm in diameter. *Stolons* branched, hyaline, brownish, smooth, with septa. *Rhizoids* coralliform, branched or unbranched. *Sporangiophores* erect or slightly bent, 1–4 in whorls, often monopodial, simple, unbranched, rarely sympodial, hyaline, sometimes with one septum or two septa 12.0–22.0 µm below apophyses, 25.0–170.0 µm long and 2.5–6.0 µm wide. *Apophyses* distinct, 3.5–7.0 µm high, 3.5–6.0 µm wide at the base, and 5.0–11.0 µm wide at the top. *Sporangia* subglobose to ellipsoid, multi-spored, hyaline when young, brownish with age, 14.5–26.5 µm long and 13.5–24.0 µm wide, walls deliquescent. *Columellae* hemispherical, subglobose to globose, clavate, hyaline and brownish, smooth, 6.0–17.0 µm long and 6.5–17.5 µm wide. *Collars* usually absent, and rarely distinct if present. *Sporangiospores* always cylindrical to ellipsoid, or rarely irregular, subhyaline, smooth, 3.0–6.0 µm long and 2.5–5.0 µm wide. *Zygospores* absent. *Chlamydospores* absent.

*Material examined*: China, Xinjiang Auto Region, Altay, Burqin Country, 28°37’1″N, 87°2’58″E, from soil sample, 31 October 2021, Heng Zhao (holotype HMAS 351509, living ex-holotype culture CGMCC 3.16107).

*GenBank accession number*: OL678136.

*Notes*: See notes of *Absidia chinensis*.

**16. *Backusella dichotoma*** H. Zhao, Y.C. Dai & X.Y. Liu, sp. nov. Fig. 32.

**Fig. 32.**
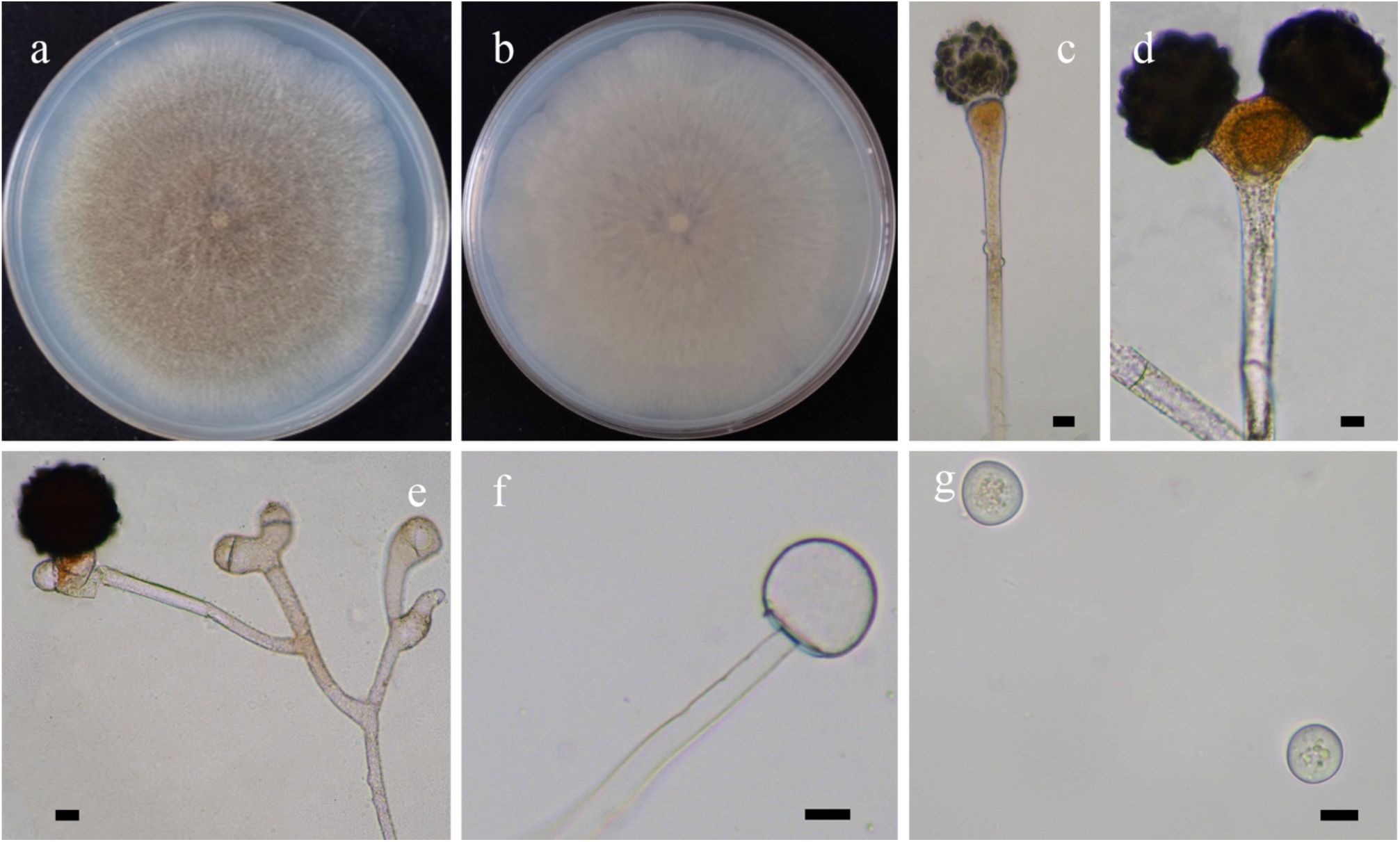
Morphologies of *Backusella dichotoma* ex-holotype CGMCC 3.16108. **a, b.** Colonies on PDA (**a.** obverse, **b.** reverse); **c–e.** Azygospores; **f.** Columellae of terminal sporangia; **g.** Sporangiospores. — Scale bars: **c–g.** 10 μm.

*Fungal Names*: FN570897.

*Etymology*: *dichotoma* (Lat.) refers to the species producing a dichotomous form of azygospores.

*Holotype*: HMAS 351510.

*Colonies* on PDA at 27 °C for 7 days, reaching 90 mm in diameter, less than 5 mm high, flat, granulate, initially white, soon becoming Sudan Brown to Brussels Brown, irregular at margin. *Odor* none. *Hyphae* sparse, hyaline, aseptate at first, septate with age, 2.5–9.5 μm in diameter. *Rhizoids* absent*. Stolons* absent. *Sporangiophores* arising directly from substrate mycelia, usually with large terminal sporangia, erect, forming lateral circinate branches, bent or curved, simple or divided once to several times. *Terminal sporangia* subglobose to globose, hyaline or brownish, smooth, multi-spored, more than 10 sporangiospores per sporangium, walls deliquescent. *Lateral circinate branches* simple or sympodial, ending with a multi-spored sporangiolum or a uni-sporangiolum. *Lateral sporangiola* globose, with numerous spines, hyaline, containing 2–6 sporangiospores, or rarely uni-spored, 10.5–28.5 μm in diameter, walls deliquescent. *Apophyses* absent. *Collars* absent. *Columellae* hyaline, always hemispherical in terminal sporangia, 16.0–25.5 μm long and 14.5–23.5 μm wide, always conical or hemispherical in lateral sporangiola, 5.5–11.5 μm long and 6.5–12.5 μm wide. *Sporangiospores* globose, subglobose or ovoid, hyaline, with droplets, 11.0–15.5 μm long and 10.5–13.5 μm wide. *Azygosporangia* mostly globose, occasionally subglobose, initially hyaline, soon becoming Light Brown to black, with conical projections on walls, 51.0–104.0 μm in diameter. *Suspensor cells* erect, bent, usually dichotomous, Light Brown. *Chlamydospores* absent. *Zygospores* absent.

*Materials examined*: China. Yunnan Province, Lincang, Cangyuan Country, from soil sample, 22 July 2021, Heng Zhao (holotype HMAS 351510, living ex-holotype culture CGMCC 3.16108, and living culture XY07504).

*GenBank accession numbers*: OL678137 and OL678138.

*Notes*: *Backusella dichotoma* is closely related to *B. psychrophila* Urquhart & Douch based on phylogenetic analysis of ITS rDNA sequences (Fig. 6). However, morphologically *B. psychrophila* differs from *B. dichotoma* by producing collars, variably shaped sporangiospores, including globose, broadly ellipsoid to ellipsoid, and the absence of azygospores (Urquhart *et al*. 2021).

**17. *Backusella moniliformis*** H. Zhao, Y.C. Dai & X.Y. Liu, sp. nov., Fig. 33.

**Fig. 33.**
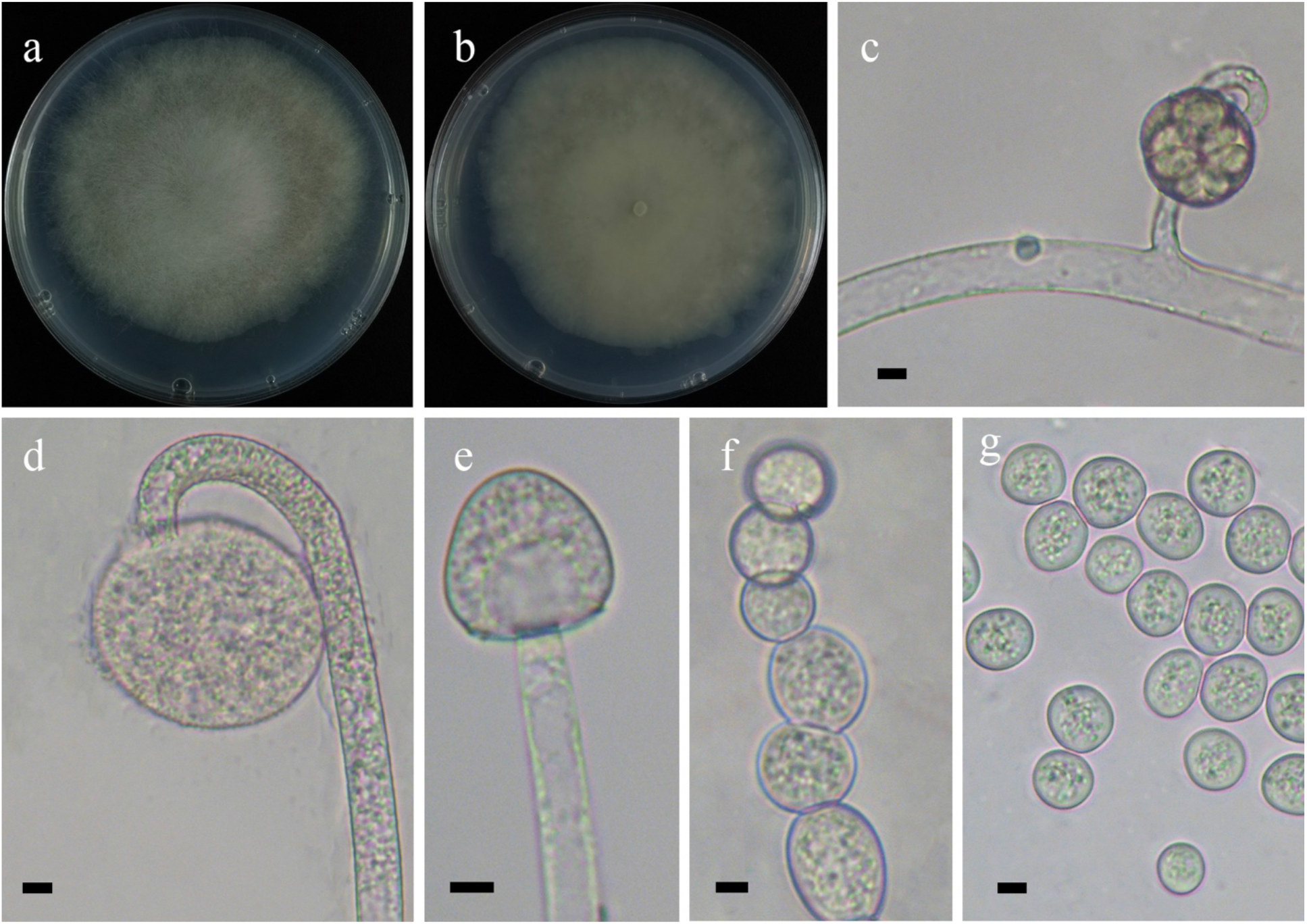
Morphologies of *Backusella moniliformis* ex-holotype CGMCC 3.16109. **a, b.** Colonies on PDA (**a.** obverse, **b.** reverse); **c.** Lateral uni-spored sporangiolum; **d.** Lateral uni-spored sporangiolum; **e.** Columellae; **f.** Chlamydospores; **g.** Sporangiospores. — Scale bars: **c–g. 5** μm.

*Fungal Names*: FN570898.

*Etymology*: *moniliformis* (Lat.) refers to the species having moniliform chlamydospores in the substrate hyphae.

*Holotype*: HMAS 351511.

*Colonies* on PDA at 27 °C for 5 days, reaching 90 mm in diameter, 15 mm high, floccose, white, irregular at margin. *Odor* none. *Hyphae* flourishing, aseptate at first, septate with age, hyaline, 4.5–13.5 μm in diameter. *Rhizoids* absent. *Stolons* absent. *Sporangiophores* arising directly from substrate hyphae, usually with a large terminal sporangium, erect or slightly curved, forming lateral circinate branches, simple or divided once. *Terminal sporangia* mostly globose, occasionally subglobose, rough with spines, Light Brown to black, multi-spored with more than 30 sporangiospores per sporangium, 51.0–61.5 μm in diameter, walls deliquescent. *Lateral circinate branches* simple or sympodial, ending with multi-spored sporangiola or uni-sporangiola. *Lateral sporangiola* globose, with spines, usually multi-spored with 3–20 sporangiospores and Light Brown to black, rarely uni-spored and yellow to Light Brown, 22.0–43.5 μm in diameter, walls deliquescent. *Apophyses* absent. *Collars* usually absent, rarely small if present. *Columellae* frequently hemispherical or conical, hyaline or brownish, 17.0–27.5 μm long and 18.0–26.5 μm wide. *Sporangiospores* subglobose to globose, ovoid, hyaline, with droplets, 7.5–13.0 μm long and\ 5.5–12.0 μm wide. *Chlamydospores* abundant in substrate hyphae, moniliform when young, becoming ovoid, subglobose or globose when mature, hyaline, with droplets, 9.5–21.0 μm long and 11.5–15.5 μm wide. *Zygospores* unknown.

*Material examined*: China, Shaanxi Province, Baoji, 107°46’40.03′′E, 37°0’0.51′′N, from soil sample, 1 October 2013, Xiao-Yong Liu (holotype HMAS 35151, living ex-holotype culture CGMCC 3.16109).

*GenBank accession number*: OL678139

*Notes*: *Backusella moniliformis* is closely related to *Backusella* “group X” Urquhart & Douch based on phylogenetic analysis of ITS rDNA sequences (Fig. 6). However, morphologically *Backusella* ‘group X’ differs from *B. moniliformis* by the variably shaped columellae, including globose, ellipsoid and applanate, and the absence of chlamydospores (Urquhart *et al*. 2021).

**18. *Backusella ovalispora*** H. Zhao, Y.C. Dai & X.Y. Liu, sp. nov., Fig. 34.

**Fig. 34.**
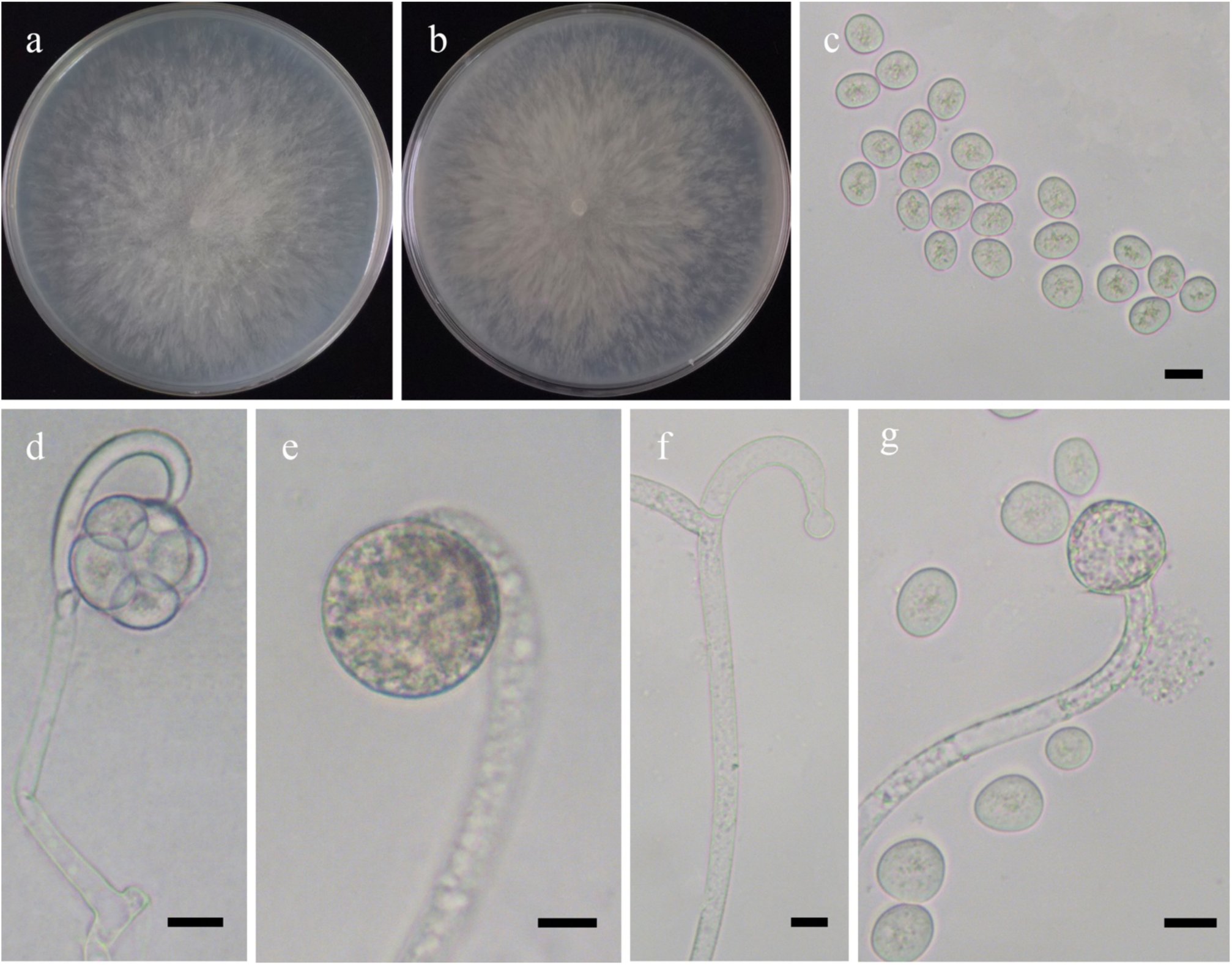
Morphologies of *Backusella ovalispora* ex-holotype CGMCC 3.16110. **a, b.** Colonies on PDA (**a.** obverse, **b.** reverse); **c.** Sporangiospores; **d.** Lateral multi-spored sporangiola; **e.** Lateral uni-spored sporangiolum; **f, g.** Columellae — Scale bars: **c–g.** 10 μm.

*Fungal Names*: FN570899.

*Etymology*: *ovalispora* (Lat.) refers to the species having ovoid sporangiospores.

*Holotype*: HMAS 351512.

*Colonies* on PDA at 27 °C for 3 days, fast growing, reaching 90 mm in diameter, 20 mm high, cottony, initially white, gradually becoming Grayish White. *Odor* none. *Hyphae* branched, hyaline, aseptate at first, septate with age, 4.5–18.0 μm in diameter. *Rhizoids* absent. *Stolons* absent. *Sporangiophores* arising from substrate or aerial hyphae, frequently tapering slightly toward the sporangia, usually with a large terminal sporangium, straight or slightly curved, and then forming lateral circinate branches, bent or curved, simple or divided once to several times successively and terminating in a sporangiolum. *Terminal sporangia* mostly globose, occasionally subglobose, hyaline or pigmented, smooth, multi-spored, usually with more than 50 sporangiospores per sporangium, 52.5–89.5 μm in diameter, walls deliquescent. *Lateral circinate branches* simple, ending with a multi-spored sporangiolum or rarely with a uni-sporangiolum. *Lateral sporangiola* mostly globose, occasionally subglobose, smooth, hyaline, three to fifteen sporangiospores per sporangiolum, 16.0–37.0 μm in diameter, walls deliquescent. *Apophyses* absent. *Collars* frequently absent, rarely small. *Columellae* always globose or subglobose in terminal sporangia, usually conical, subglobose, globose or depressed globose in lateral sporangiola, hyaline or brown, 6.0–59.5 μm long and 7.0–60.5 μm wide. *Sporangiospores* with striations, mainly ovoid, rarely irregular, 8.5–18.5 μm long and 7.0–16.0 μm wide. *Chlamydospores* absent. *Zygospores* absent.

*Materials examined*: China, Guangxi Auto Region, Fangchenggang, 21°44’27″N, 108°3’29″E, from soil sample, 15 July 2021, Heng Zhao (holotype HMAS 351512, living ex-holotype culture CGMCC 3.16110, and living culture XY07481).

*GenBank accession numbers*: OL678140 and OL678141.

*Notes*: *Backusella ovalispora* is closely related to *B. gigacellularis* J.I. Souza *et al*. based on phylogenetic analysis of ITS rDNA sequences (Fig. 6). However, morphologically *B. gigacellularis* differs from *B. ovalispora* by abundant giant cells, ellipsoid, cylindrical and rarely pyriform columellae, ellipsoid sporangiospores, and multi-spored sporangiola consisting of 3‒4 sporangiospores (de Souza *et al*. 2014).

**19. *Circinella homothallica*** H. Zhao, Y.C. Dai & X.Y. Liu, sp. nov., Fig. 35.

**Fig. 35.**
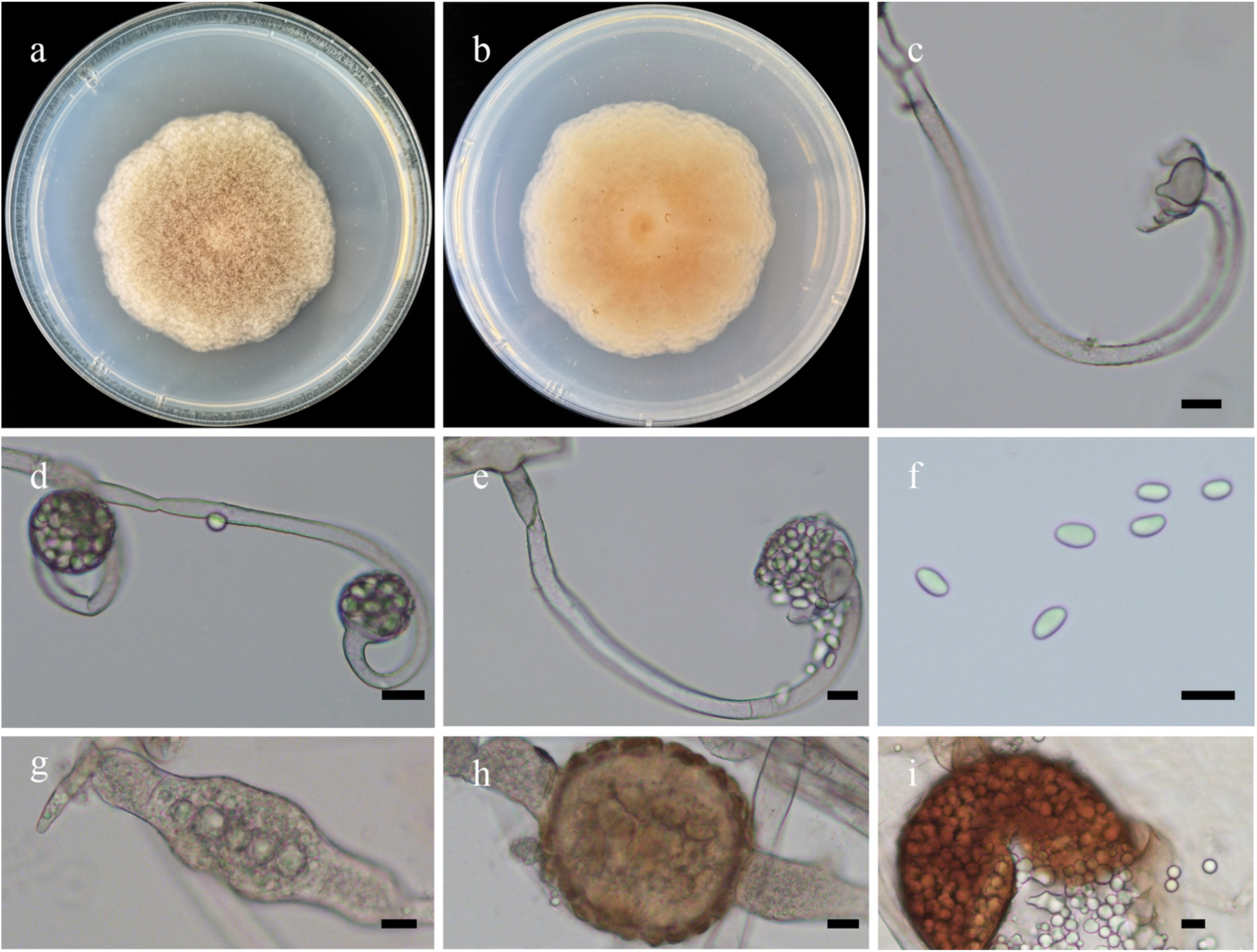
Morphologies of *Circinella homothallica* ex-holotype CGMCC 3.16026. **a, b**. Colonies on PDA (**a,** obverse, **b,** reverse); **c,** Columellae with a projection and collars; **d, e.** Circinate lateral sporangiophores**; h.** Sporangiospores; **g–i.** Zygospores. — Scale bars: **c–i,** 10 μm.

*Fungal Names*: FN570831.

*Etymology*: *homothallica* (Lat.) refers to the species producing zygospores by homothallism.

*Holotype*: HMAS 249879.

*Colonies* on PDA at 27 °C for 9 days, slow growing, reaching 50 mm in diameter, carpet-like, initially white, soon becoming Mars Yellow, irregular at margin. *Hyphae* branched, aseptate when juvenile, septate with age, 4.5–10.5 μm in diameter. *Rhizoids* absent. *Stolons* absent. *Sporangiophores* always laterally circinate, and apically erect. *Sporangia* globose to ellipsoid, smooth, 17.0–43.0 μm long and 14.5–43.5 μm wide, walls deliquescent. *Apophyses* present. *Collars* always present. *Columellae* globose to conical, smooth, 8.0–31.5 μm long and 11.0–29.5 μm wide. *Projections* sometimes present. *Sporangiospores* ellipsoid, 5.5–8.5 μm long and 2.5–4.5 μm wide. *Chlamydospores* absent. *Zygospores* homothallic, hyaline when juvenile, gradually brownish with age, surrounded by droplets, 53.0–90.0 μm long and 55.0–89.0 μm wide.

*Material examined*: China, Hainan Province, Shihuashui Cave, from soil sample, 12 November 2020, Jia-Jia Chen (holotype HMAS 249879, living ex-holotype culture CGMCC 3.16026).

*GenBank accession number*: MW580599.

*Notes*: *Circinella homothallica* is distinguished from all other species of the genus by producing zygospores through a homothallic mating. Bainier (1903) first reported zygospores in *Circinella*, however, zygospores have not been observed by any other researchers (Hesseltine & Fennell 1955, Zheng *et al*. 2017). *C. homothallica* is closely related to *C. mucoroides* Saito based on phylogenetic analysis of ITS rDNA sequences (Fig. 7). However, morphologically *C. mucoroides* differs from *C. homothallica* by producing globose, shortly ovoid sporangiospores, ovoid, cylindrical, or pyriform columellae, lacking projections (Zheng *et al*. 2017).

**20. *Cunninghamella arrhiza*** H. Zhao, Y.C. Dai & X.Y. Liu, sp. nov., Fig. 36.

**Fig. 36.**
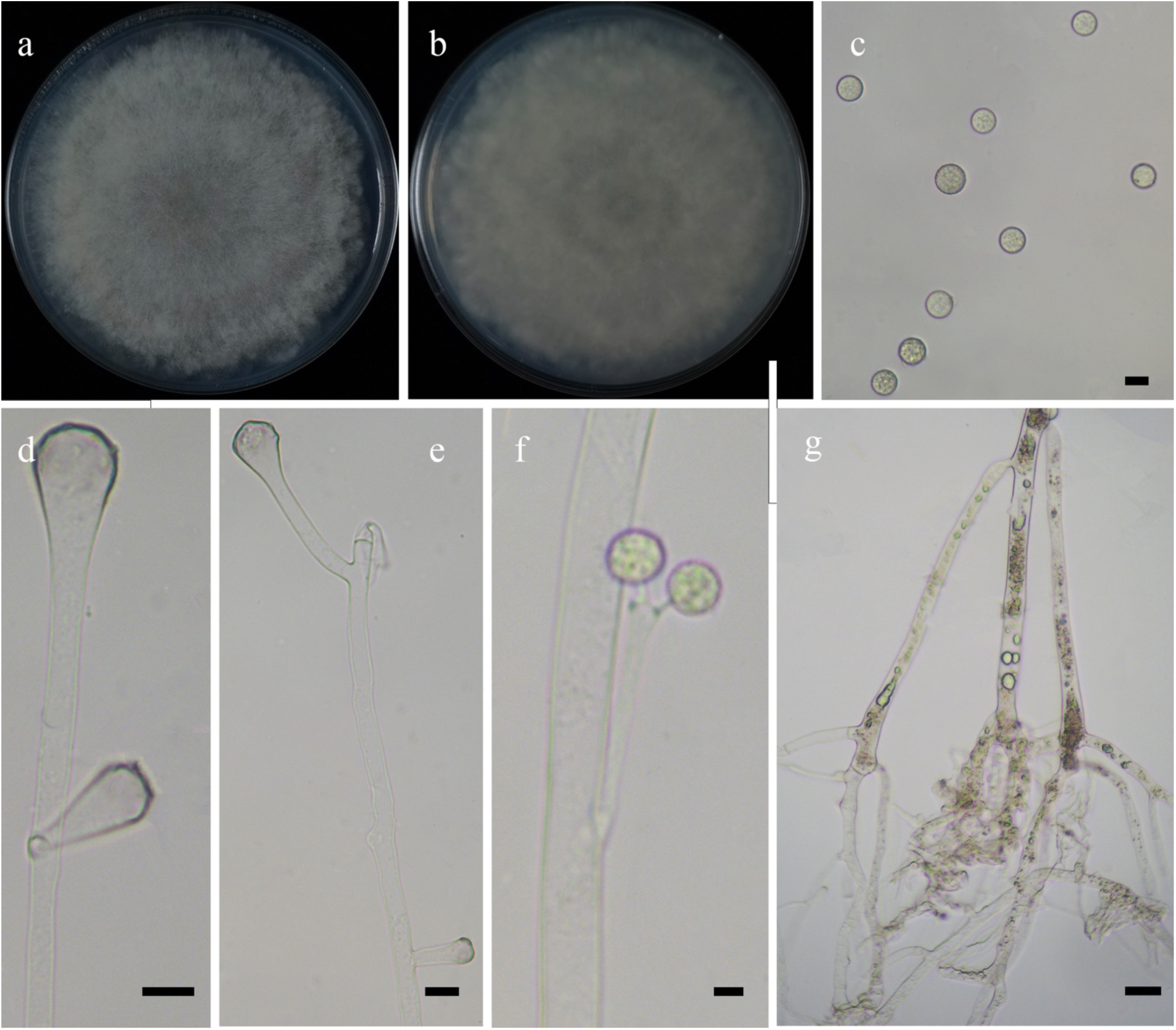
Morphologies of *Cunninghamella arrhiza* ex-holotype CGMCC 3.16111. **a, b.** Colonies on PDA (**a.** obverse, **b.** reverse); **c.** Sporangiola; **d–f.** Sporangiophores showing characteristic branching patterns with vesicles; **g.** Rhizoids. — Scale bars: **c–f.** 10 μm, **g.** 20 μm.

*Fungal Names*: FN570900.

*Etymology*: *arrhiza* (Lat.) refers to the species rarely producing rhizoids.

*Holotype*: HMAS 351513.

*Colonies* on PDA at 27 °C for 5 days, reaching 90 mm in diameter, more than 20 mm high, initially white, gradually becoming Light Gray, floccose, irregular at margin. *Hyphae* flourishing, branched, aseptate when young, septate when old, 1.5–12.0 μm in diameter. *Rhizoids* infrequent, root-like, always branched. *Stolons* present. *Sporangiophores* arising from aerial hyphae, erect, straight, always broadening upwards, unbranched or simple branched, in pairs, monopodial, but never verticillate. *Septa* absent. *Vesicles* subglobose, clavate, hyaline or subhyaline, rough, sometimes with brownish tints, 7.0–36.5 μm long and 5.0–29.5 μm wide. *Apophyses* present. *Pedicels* 3.0–4.5 μm long. *Sporangiola* borne on pedicels on vesicles, globose with thick spines, 7.5–13.0 μm in diameter. *Chlamydospores* absent. *Zygospores* unknown.

*Materials examined*: China. Hunan Province, Changsha, 28°8’4″N, 112°57’19″E, from soil sample, 28 August 2021, Heng Zhao (holotype HMAS 351513, living ex-holotype culture CGMCC 3.16111). Beijing, 40°39’35″N, 117°16’35″E, from soil sample, 31 July 2021, Heng Zhao (living culture XY08047).

*GenBank accession numbers*: OL678142 and OL678143.

*Notes*: *Cunninghamella arrhiza* is closely related to *C. guizhouensis* Zhi.Y. Zhang *et al*. based on phylogenetic analysis of ITS rDNA sequences (Fig. 7). However, morphologically *C. guizhouensis* differs from *C. arrhiza* by producing monopodial and verticillate sporangiophores, and globose to subglobose versicles (Zhang *et al*. 2020).

**21. *Cunninghamella guttata*** H. Zhao, Y.C. Dai & X.Y. Liu, sp. nov., Fig. 37.

**Fig. 37.**
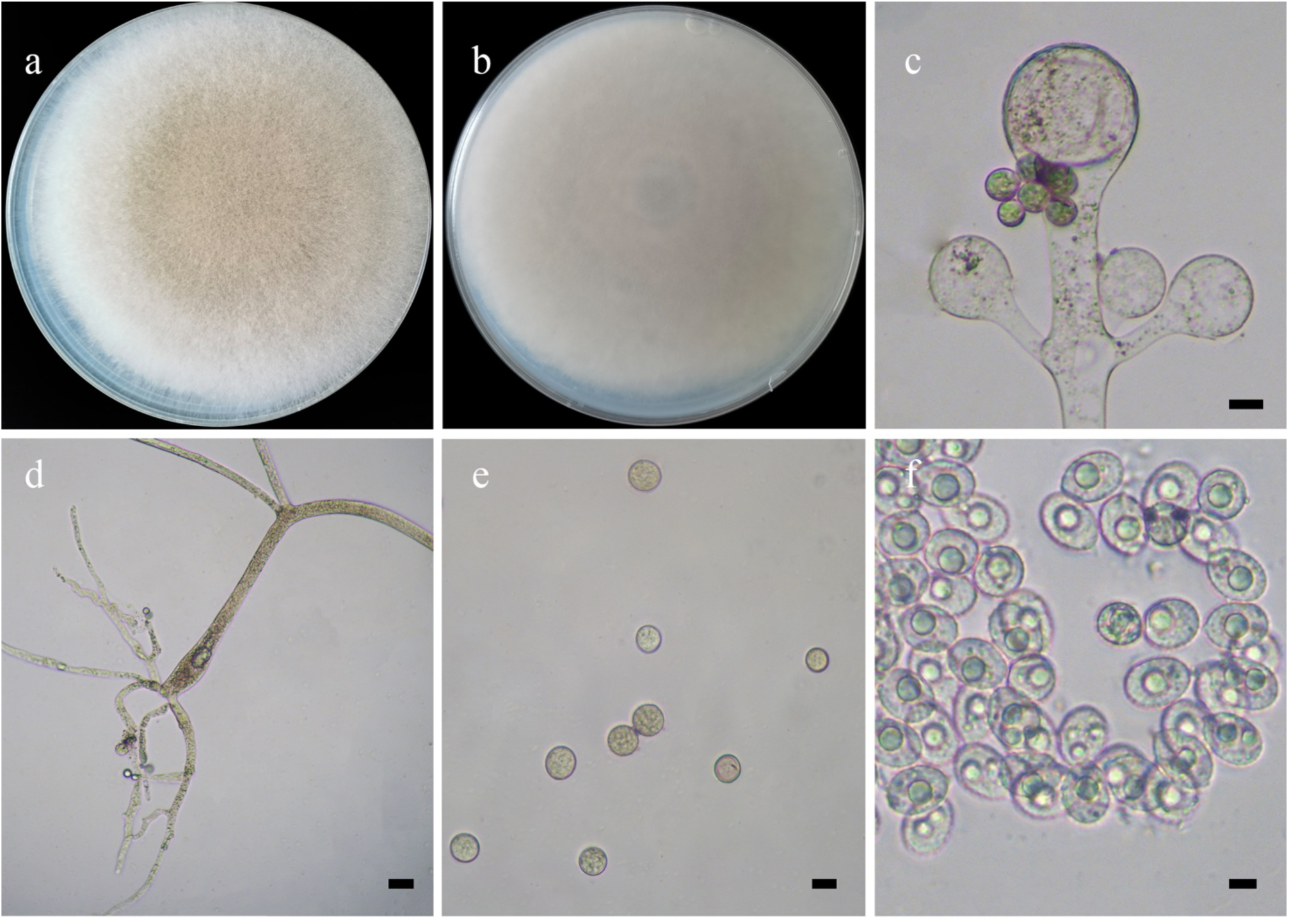
Morphologies of *Cunninghamella guttata* ex-holotype CGMCC 3.16112. **a, b.** Colonies on PDA (**a.** obverse, **b.** reverse); **c.** Sporangiophores showing characteristic branching patterns; **d.** Rhizoids; **e, f.** Sporangiola. — Scale bars: **c, e.** 10 μm, **d.** 20 μm, **f. 5** μm.

*Fungal Names*: FN570901.

*Etymology*: *guttata* (Lat.) refers to the species producing lipid droplets in sporangiola.

*Holotype*: HMAS 351514.

*Colonies* on PDA at 27 °C for 6 days, reaching 90 mm in diameter, 15 mm high, initially white, becoming Drab Gray with age, floccose, in reverse regular at margin. *Hyphae* branching, aseptate when young, rarely septate with age, 4.5–20.5 μm in diameter. *Rhizoids* abundant, root-like, branched. *Stolons* present. *Sporangiophores* arising from stolons or aerial hyphae, erect, straight or bent, 0–4 branched, single, in pairs or 3 or 4 verticillate, often broadening upwards. *Septa* if present, often at the upper part of the sporophores. *Vesicles* subglobose or globose, usually hyaline or subhyaline, sometimes with brownish tints, 13.5–40.5 μm long and 12.5–40.5 μm wide. *Apophyses* present. *Pedicels* 2.0–3.0 μm long. *Sporangiola* borne on pedicels on vesicles, ovoid, with droplets when young, subhyaline or hyaline, 9.0–12.5 μm long and 7.5–9.0 μm wide, mostly globose, occasionally subglobose, dark, with a few spines, 8.5–15.0 μm in diameter.

*Chlamydospores* absent. *Zygospores* unknown.

*Material examined*: China, Beijing, 40°30’29.16″N, 116°48’29.34″E, from soil sample, 28 July 2021, Heng Zhao (holotype HMAS 351514, living ex-holotype culture CGMCC 3.16112).

*GenBank accession number*: OL678144.

*Notes*: *Cunninghamella guttata* is closely related to *C. varians* H. Zhao *et al*. based on phylogenetic analysis of ITS rDNA sequences (Fig. 8). However, morphologically *C. varians* differs from *C. guttata* by producing two types of sporangiola (ovoid or globose vs. mostly globose, occasionally subglobose), and short pedicels (1.0–1.5 μm long vs. 2.0–3.0 μm long).

**22. *Cunninghamella irregularis*** H. Zhao, Y.C. Dai & X.Y. Liu, sp. nov., Fig. 38.

**Fig. 38.**
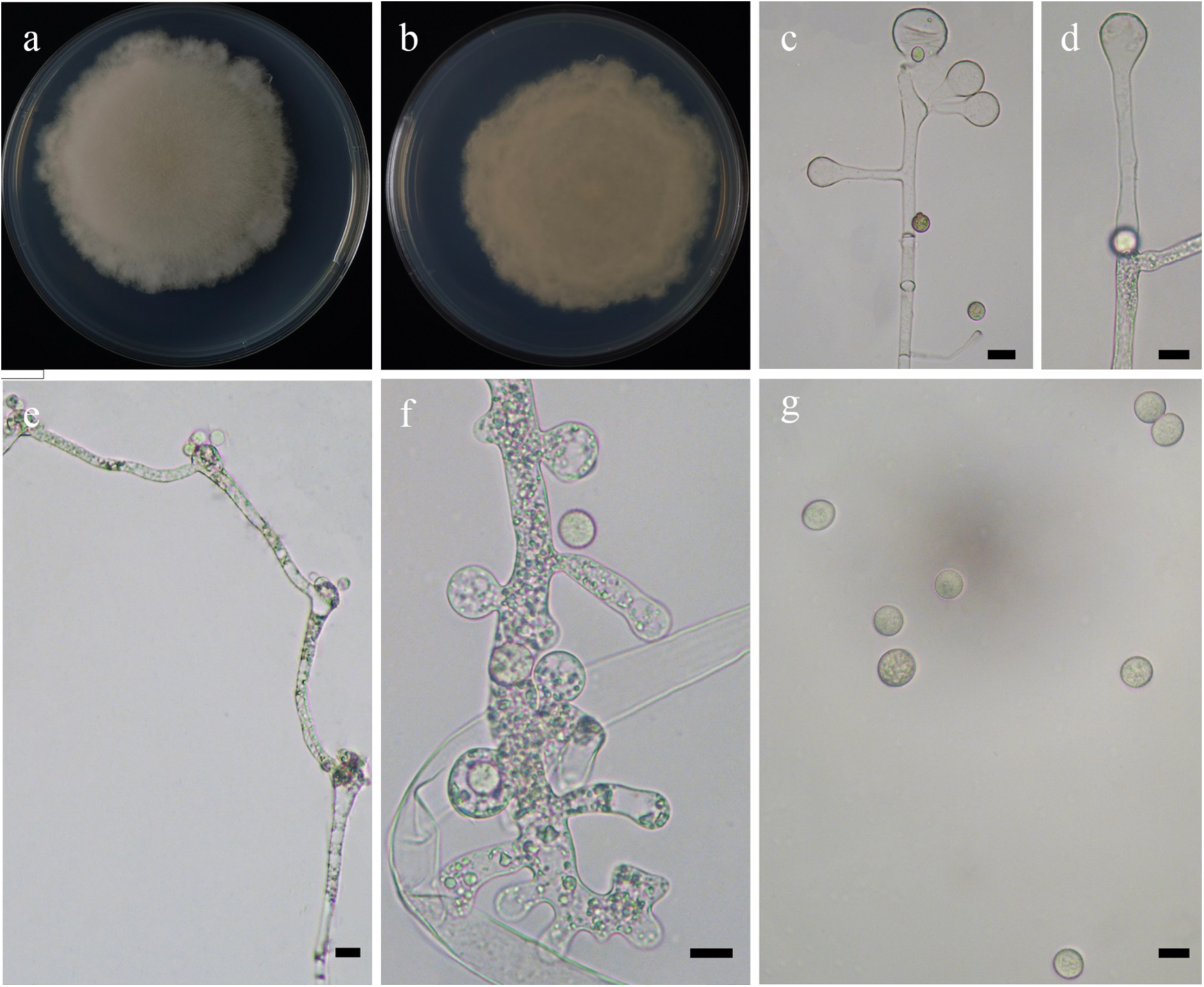
Morphologies of *Cunninghamella irregularis* ex-holotype CGMCC 3.16113. **a, b.** Colonies on PDA (**a.** obverse, **b.** reverse); **c–e.** Sporangiophores showing characteristic branching patterns with vesicles; **f.** Rhizoids; **g.** Sporangiola. — Scale bars: **c–g.** 10 μm.

*Fungal Names*: FN570902.

*Etymology*: *irregularis* (Lat.) refers to the colonies having an irregular margin.

*Holotype*: HMAS 351515.

*Colonies* on PDA at 27 °C for 7 days, slow growing, reaching 70 mm in diameter, 10 mm high, initially white, gradually becoming Light Gray, floccose, irregular at margin. *Hyphae* flourishing, branched, aseptate when young, septate when old, 3.0–13.5 μm in diameter. *Rhizoids* abundant, root-like, always branched, sometimes swollen. *Stolons* present. *Sporangiophores* arising from aerial hyphae, erect, straight, or few slightly bent, always broadening upwards, unbranched, or simple branched, in pairs, sympodial, but never verticillate. *Septa* if present, usually one to several below the vesicles on the sporangiophores. *Vesicles* ovoid, subglobose, globose and rarely clavate, even irregular, rough, hyaline, sometimes with brownish tints, 3.0–40.0 μm long and 2.5–34.5 μm wide. *Apophyses* present. *Pedicels* short, usually less than 3.0 μm long. *Sporangiola* borne on pedicels on vesicles, two types: with thick spines, ovoid, 10.0–12.5 μm long and 8.0–10.5 μm wide, and globose, 9.0–16.5 μm in diameter. *Chlamydospores* absent. *Zygospores* unknown.

*Materials examined*: China, Inner Mongolia Auto Region. Xilingol, Erenhot, 43°24’53″N, 112°7’55″E, from soil sample, 31 July 2021, Heng Zhao (holotype HMAS 351515, living ex-holotype culture CGMCC 3.16113, and living culture XY07657); Ulanqab, Chahar, 41°50’8″N, 113°4’10″E, from soil sample, 31 July 2021, Heng Zhao (living culture XY07683).

*GenBank accession numbers*: OL678145, OL678146 and OL678147.

*Notes*: *Cunninghamella irregularis* is closely related to *C. varians* H. Zhao *et al*. based on phylogenetic analysis of ITS rDNA sequences (Fig. 8). However, morphologically *C. varians* differs from *C. irregularis* by verticillate sporangiophores.

**23. *Cunninghamella nodosa*** (R.Y. Zheng) H. Zhao, Y.C. Dai & X.Y. Liu, comb. et stat. nov., Fig. 39.

**Fig. 39.**
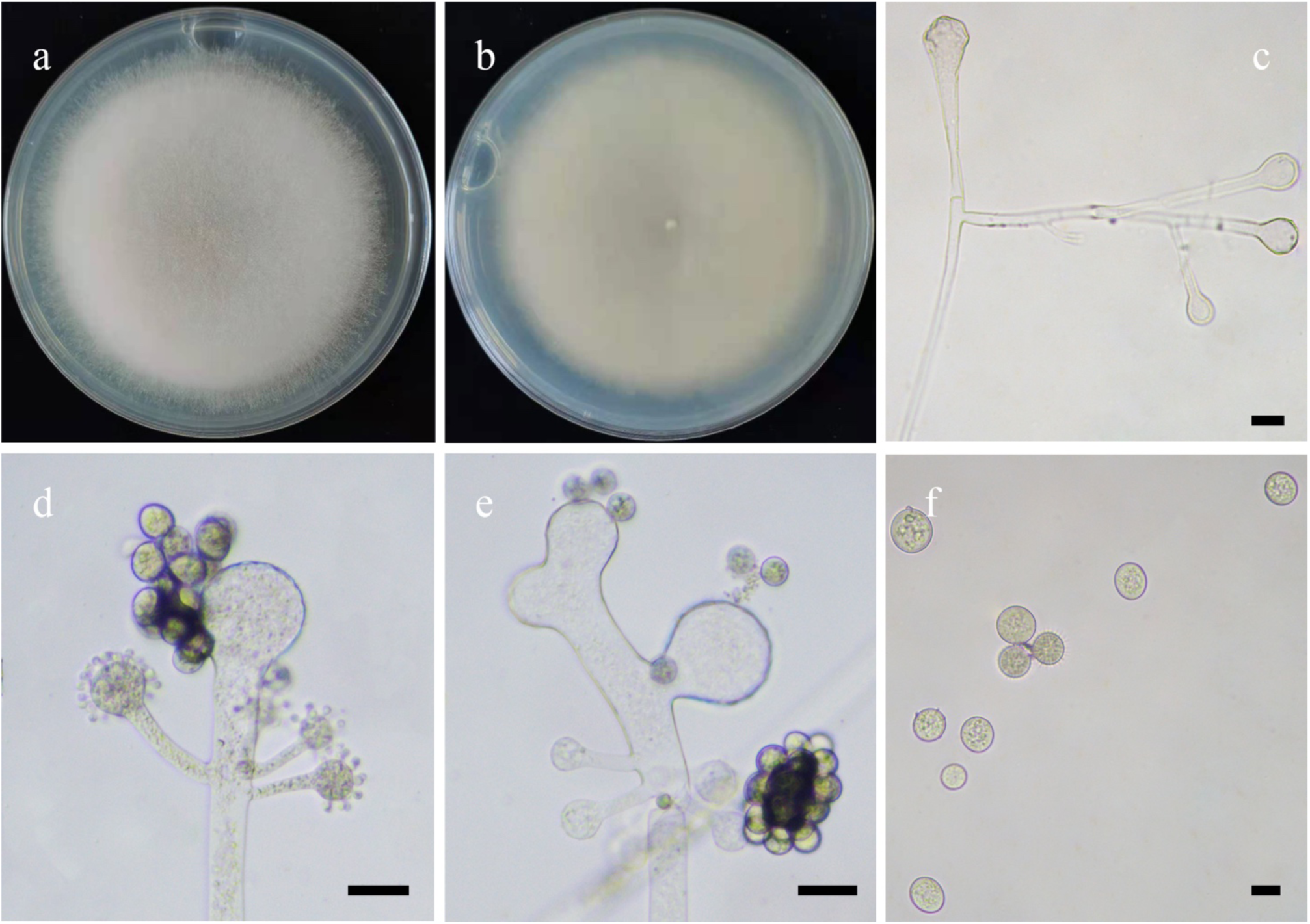
Morphologies of *Cunninghamella nodosa* ex-holotype CGMCC 3.5628. **a, b.** Colonies on PDA (**a.** obverse, **b.** reverse); **c–e.** Sporangiophores showing characteristic branching patterns; **f.** Sporangiola. — Scale bars: **c–f.** 10 μm.

*Fungal Names*: FN570903.

Basionym: *Cunninghamella echinulata* var. *nodosa* R.Y. Zheng, in Zheng & Chen, Mycotaxon 80: 35, 2001 [MycoBank no.: 474177]

*Holotype*: HMAS 80705

*Colonies* on SMA at 27 °C for 6 days, reaching 90 mm in diameter, 10 mm high, white at first, finally Avellaneous or Colonial Buff, in reverse Yellowish Cream, initially floccose, soon granulate. *Hyphae* branched, aseptate when young, septate with age, 3.5**–**14.0 µm in diameter. *Stolons* sometimes present. *Rhizoids* common, simple or repeatedly branched. *Sporangiophores* arising from stolons, sometimes opposite rhizoids, or directly from aerial hyphae, erect, stolon-like, straight or recumbent, verticillate or pseudoverticillate, mostly simple, occasionally in pairs, rarely repeated, typically short and 10.0**–**75.0 µm long, occasionally reaching 250.0 µm long, usually broadening upwards, sometimes with nodulous swellings, 7.5**–**25.0 µm in diameter. *Septa* sometimes in sporangiophores. *Vesicles* forming on the top of the main axes of sporangiophores, slightly depressed-globose, subglobose, 22.5**–**70.0 µm in diameter, or forming on lateral branches, usually globose, occasionally broadly ovoid, 8.5**–**27.5 µm in diameter, smooth and hyaline, occasionally proliferating. *Pedicels* numerous, arising over entire surface of vesicles, 2.5**–**4 µm long. *Sporangiola* borne on pedicels on vesicles, globose, broadly ellipsoid, ovoid or bluntly pointed at one end, hyaline, 6.0–17.5 µm long 5.5–13.5 µm wide, or borne singly or intermixed with the hyaline ones on the same vesicle, globose, dark, 11.5–17.5 µm in diameter, exclusively monosporous, all evidently echinulate and with long thick spines. *Sporangiospores* detachable, but usually remaining within the sporangiola, similar in shape and size to the sporangiola, spineless, tuberculate, hyaline. *Chlamydospores* absent. *Zygosporangia* mostly globose, occasionally subglobose, Dark Brown when mature, with pointed wart-like projections, 26.5–53.5 µm in diameter. *Zygospores* heterothallic, mainly globose, rarely broad ellipsoid, hyaline, usually very thick-walled, smooth or slightly crenulate, with indistinct ridges, 23.0–48.0 µm in diameter.\ *Suspensors* equal, not inflated, 12.5–35.5 µm in long and 9.0–29.5 µm wide, slightly attenuated or typically remaining straight or broadening toward the base, aseptate or septate at the basal part, not incrusted and hyaline or sometimes incrusted and brownish (Zheng and Chen 2001).

*Material examined*: China, Beijing, from flower sample, 28 May 2021, Gui-Qing Chen, 15 August 1986, (holotype HMAS 80705, living ex-holotype CGMCC 3.5628).

*GenBank accession number*: AF346407.

*Notes*: *Cunninghamella nodosa* was initially established as a variety of *C. echinulata* (Zheng and Chen 2001). However, morphologically *C. nodosa* has two intergrading kinds of sporangiophores and dark giant sporangiola in old cultures, being distinguished from these three allied species by processing nodulous sporophores, proliferating vesicles, long and slender pedicels (Zheng and Chen 2001).

**24. *Cunninghamella regularis*** H. Zhao, Y.C. Dai & X.Y. Liu, sp. nov., Fig. 40.

**Fig. 40.**
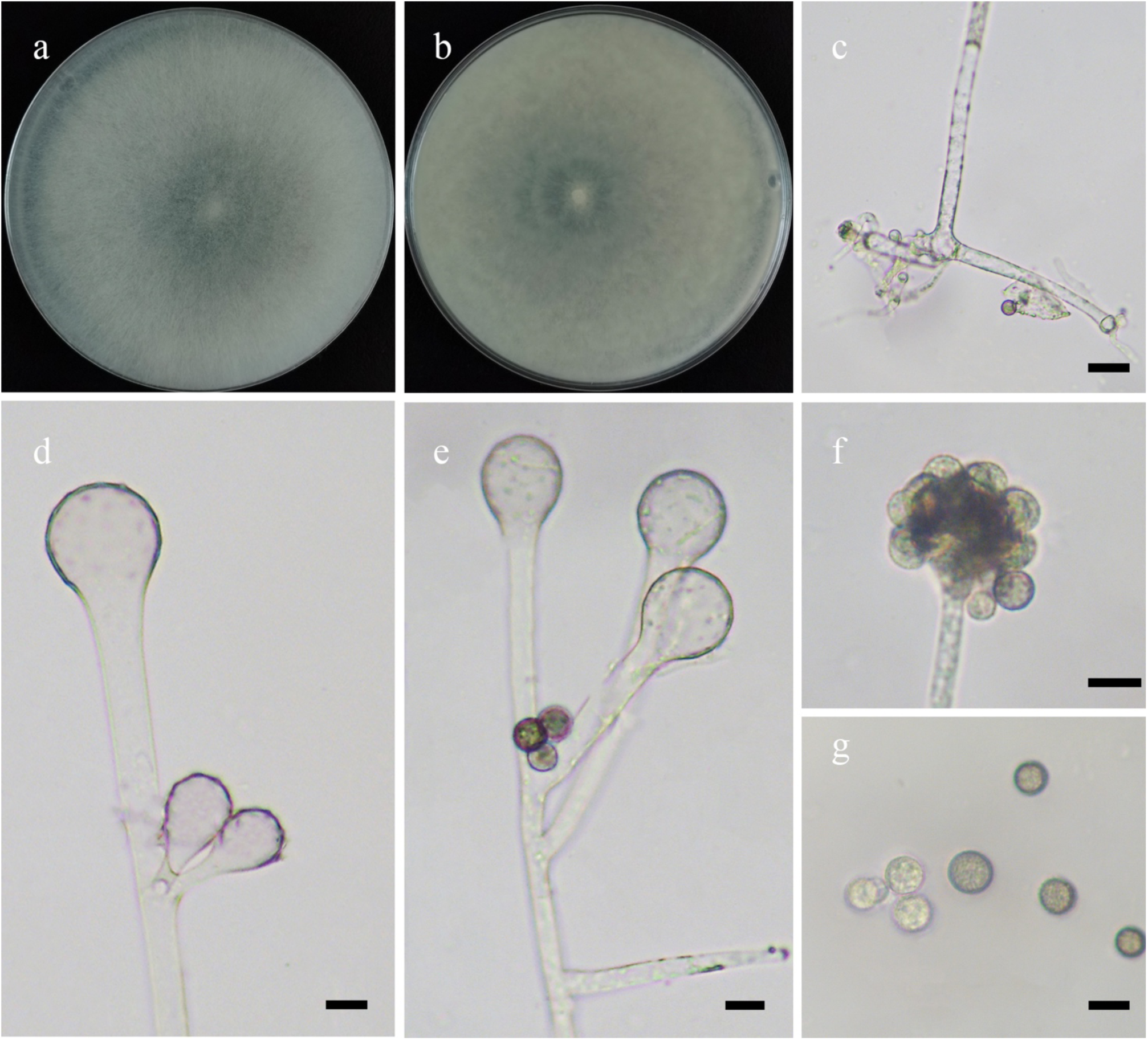
Morphologies of *Cunninghamella regularis* ex-holotype CGMCC 3.16114. **a, b.** Colonies on PDA (**a.** obverse, **b.** reverse); **c.** Rhizoids; **d, e.** Sporangiophores showing characteristic branching patterns; **f.** Lateral branches of sporangiophores with sporangiola; **g.** Sporangiola. — Scale bars: **c.** 20 μm, **d–g.** 10 μm.

*Fungal Names*: FN570904.

*Etymology*: *regularis* (Lat.) refers to the species having uniform shape and size of sporangiola.

*Holotype*: HMAS 351516.

*Colonies* on PDA at 27 °C for 4 days, fast growing, reaching 90 mm in diameter, more than 20 mm high, initially white, soon becoming Light Gray, floccose. *Hyphae* flourishing, branched, aseptate when young, septate with age, 3.5–12.0 μm in diameter. *Rhizoids* abundant, root-like or finger-like, branched or unbranched. *Stolons* present. *Sporangiophores* arising from stolons or aerial hyphae, erect, straight or slightly bent, often broadening upwards, mainly unbranched and simple branched, abundant when borne laterally, very rarely dichotomous, in pairs or sympodial. *Septa* if present, at the upper part of the sporangiophores. *Vesicles* ovoid or globose, hyaline or pigmented, rough, sometimes with brownish tints, 8.5–27.0 μm long and 7.5–28.0 μm wide. *Apophyses* present. *Pedicels* 4.5–7.5 μm long. *Sporangiola* borne on pedicels on vesicles, globose, hyaline when young, brown or black when old, with few spines, 6.5–10.0 μm in diameter. *Chlamydospores* absent. *Zygospores* unknown.

*Materials examined*: China, Yunnan Province, Lincang, Cangyuan Country, from soil sample, 22 July 2021, Heng Zhao (holotype HMAS 351516, living ex-holotype culture CGMCC 3.16114, and living culture XY07510, XY07512 and XY07516).

*GenBank accession numbers*: OL678148, OL678149, OL678150 and OL678151.

*Notes*: *Cunninghamella regularis* is closely related to *C. binariae* Naumov based on phylogenetic analysis of ITS rDNA sequences (Fig. 8). However, morphologically *C. binariae* differs from *C. regularis* by the variably shaped sporangiola, ovoid to ellipsoid, globose and lacrymoid, compared to only one type of sporangiola in *C. regularis* (Zheng & Chen 2001). Moreover, in *C. binariae*, sporangiophores are pseudoverticillate and verticillate, while they are absent in *C. regularis* (Zheng & Chen 2001).

**25. *Cunninghamella subclavata*** H. Zhao, Y.C. Dai & X.Y. Liu, sp. nov., Fig. 41.

**Fig. 41.**
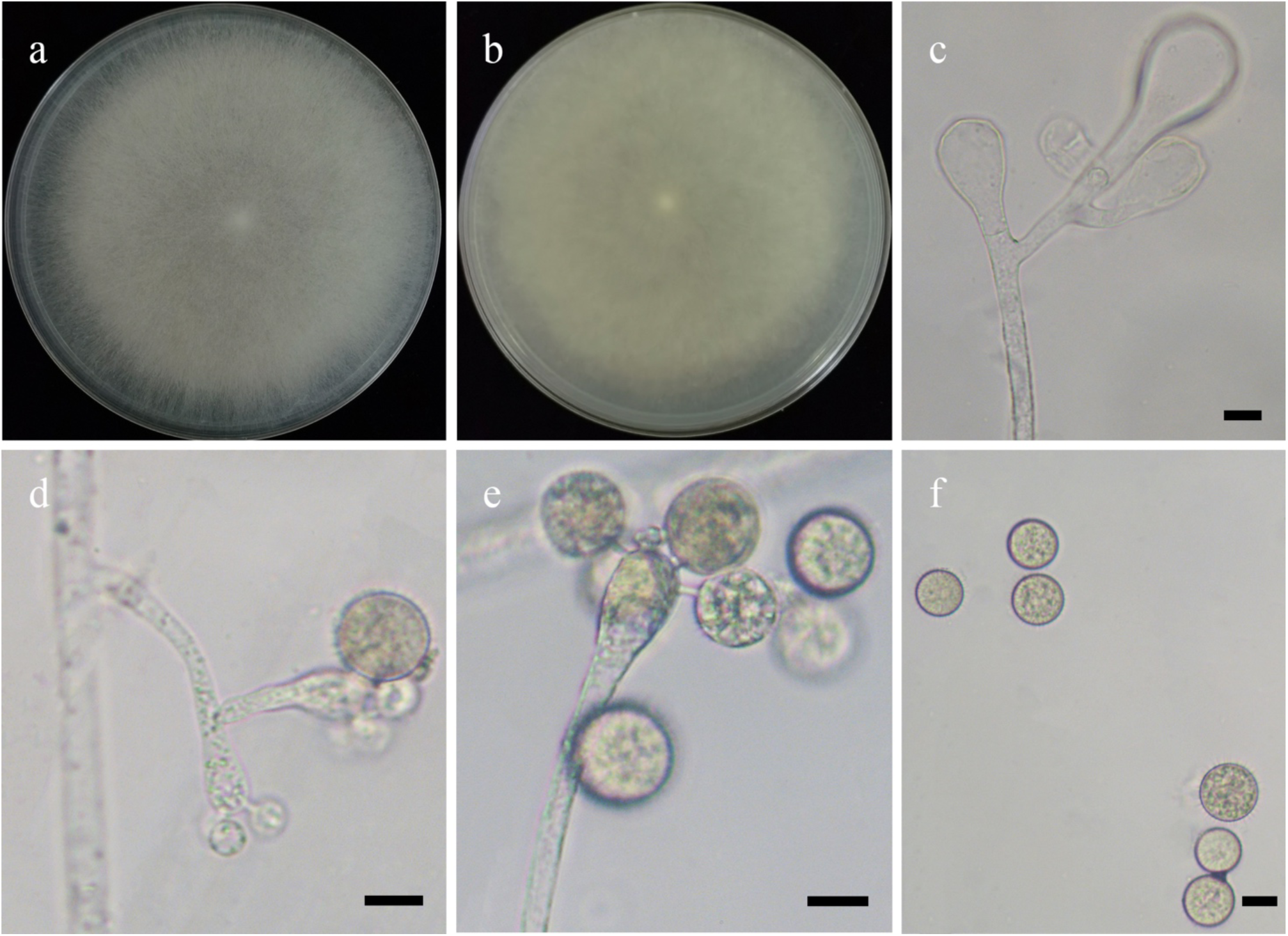
Morphologies of *Cunninghamella subclavata* ex-holotype CGMCC 3.16115. **a, b.** Colonies on PDA (**a.** obverse, **b.** reverse); **c–e.** Sporangiophores showing characteristic branching patterns; **f.** Sporangiola. — Scale bars: **c–f.** 10 μm.

*Fungal Names*: FN570905.

*Etymology*: *subclavata* (Lat.) refers to the species being similar to *Cunninghamella clavata*. *Holotype*: HMAS 351517.

*Colonies* on PDA at 27 °C for 4 days, fast growing, reaching 90 mm in diameter, 20 mm high, at first white, finally Light Gray. *Hyphae* flourishing, branched, aseptate when young, septate with age, 4.5–17.0 μm in diameter. *Rhizoids* abundant, root-like, always branched. *Stolons* present. *Sporangiophores* arising from aerial hyphae, erect, straight, always broadening upwards, unbranched or simple branched, in pairs, sympodial or dichotomous. *Septa* absent. *Vesicles* ovoid or globose on main sporangiophores, clavate or subclavate on lateral sporangiophores, hyaline, rough, sometimes with brownish tints, 9.5–33.5 μm long and 7.5–29.5 μm wide. *Apophyses* present. *Pedicels* 1.5–4.0 μm long. *Sporangiola* borne on pedicels on vesicles, globose, hyaline when young, Light Brown with age, with many thin (10.0–17.0 μm in diameter) spines on the surface. *Chlamydospores* absent. *Zygospores* unknown.

*Materials examined*: China, Yunnan Province, Xishuangbanna, 21°55’70″N, 101°16’16″E, from soil sample, 31 July 2021, Heng Zhao (holotype HMAS 351517, living ex-holotype culture CGMCC 3.16115, and living culture XY07766).

*GenBank accession numbers*: OL678152 and OL678153.

*Notes*: *Cunninghamella subclavata* is closely related to *C. clavata* R.Y. Zheng & G.Q. Chen based on phylogenetic analysis of ITS rDNA sequences (Fig. 8). However, morphologically *C. clavata* differs from *C. subclavata* by longer pedicels (2.5–11.0 μm long vs. 1.5–4.0 μm long), and typically clavate vesicles (Zheng & Chen 1998, 2001).

**26. *Cunninghamella varians*** H. Zhao, Y.C. Dai & X.Y. Liu, sp. nov., Fig. 42.

**Fig. 42.**
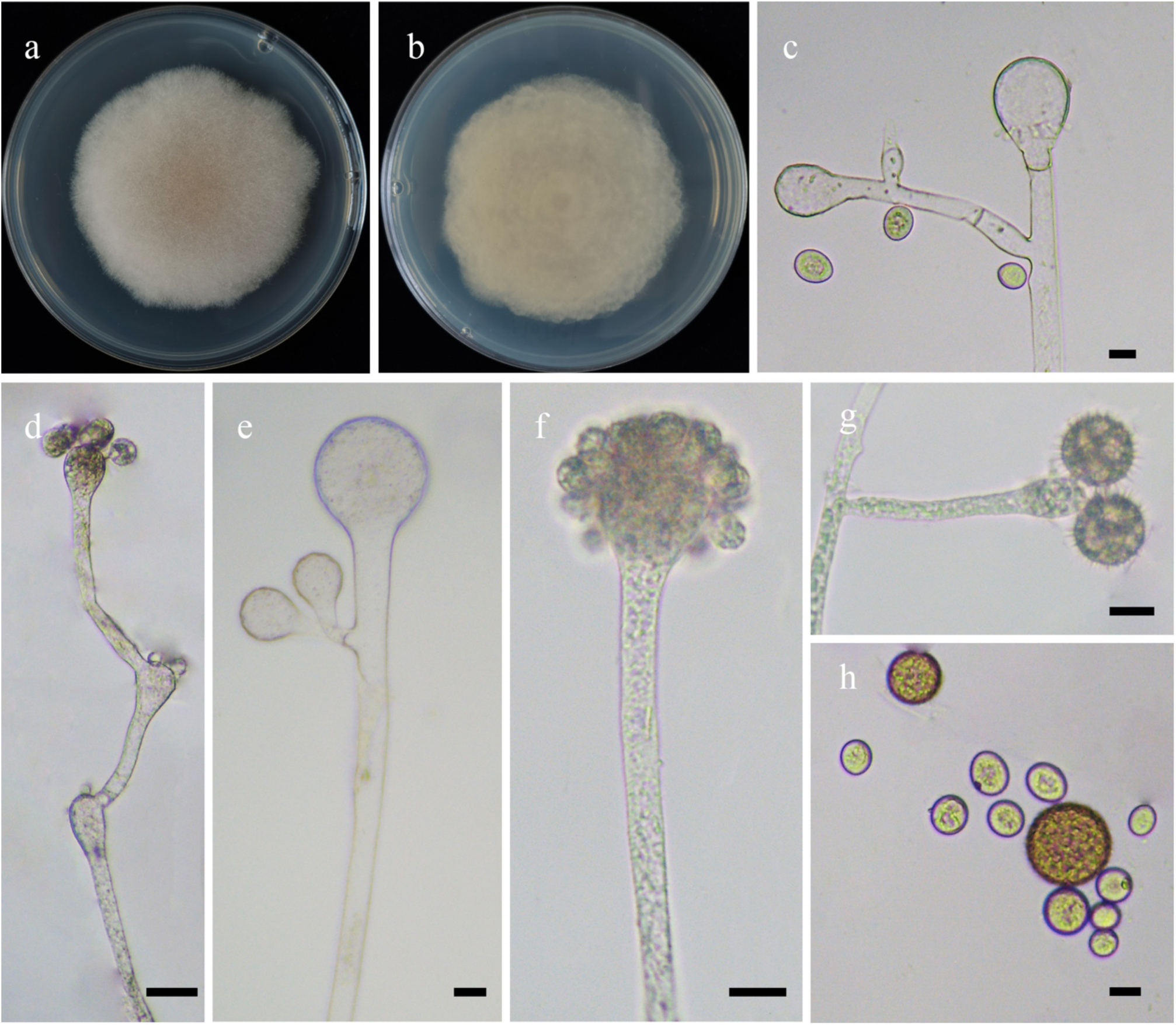
Morphologies of *Cunninghamella varians* ex-holotype CGMCC 3.16116. **a, b.** Colonies on PDA (**a.** obverse, **b.** reverse); **c–g.** Sporangiophores showing characteristic branching patterns; **h.** Sporangiola. — Scale bars: **c–h.** 10 μm.

*Fungal Names*: FN570906.

*Etymology*: *varians* (Lat.) refers to the species producing sporangiola of various shapes and sizes.

*Holotype*: HMAS 351518.

*Colonies* on PDA at 27 °C for 7 days, reaching 70 mm in diameter, 10–15 mm high, initially white, soon becoming Drab Gray, floccose, reverse irregular at margin. *Hyphae* branched, aseptate when young, septate when old, 2.5–15.0 μm in diameter. *Rhizoids* frequent, root-like, branched. *Stolons* present. *Sporangiophores* arising from stolons or aerial hyphae, erect, straight or recumbent, sometimes slightly broadening upwards in main axes, branched solitarily, in pairs, sympodially or 2–5 verticillate, variable in length, 11.5–251.5 μm in long. *Vesicles* ovoid or globose, usually hyaline, sometimes with brownish tints, 6.0–44.0 μm long and 5.0–40.0 μm wide. *Septa* often below the vesicles in the sporangiophores if present. *Apophyses* present. *Pedicels* 1.0–1.5 μm long. *Sporangiola* borne on pedicels on vesicles, two types: ovoid or globose, 7.0–14.5 μm long and 6.5– 12.5 μm wide, and dark, exclusively globose, with thick (11.0–21.0 μm in diameter) spines. *Chlamydospores* absent. *Zygospores* unknown.

*Materials examined*: China, Jiangsu Province, Xinghua, 33°8’5″N, 120°6’14″E, from soil sample, 28 May 2021, Heng Zhao (holotype HMAS 351518, living ex-holotype culture CGMCC 3.16116, and living culture XY06999).

*GenBank accession numbers*: OL678154 and OL678155.

*Notes*: See notes of *Cunninghamella irregularis*.

**27. *Gamsiella globistylospora*** H. Zhao, Y.C. Dai & X.Y. Liu, sp. nov., Fig. 43.

**Fig. 43.**
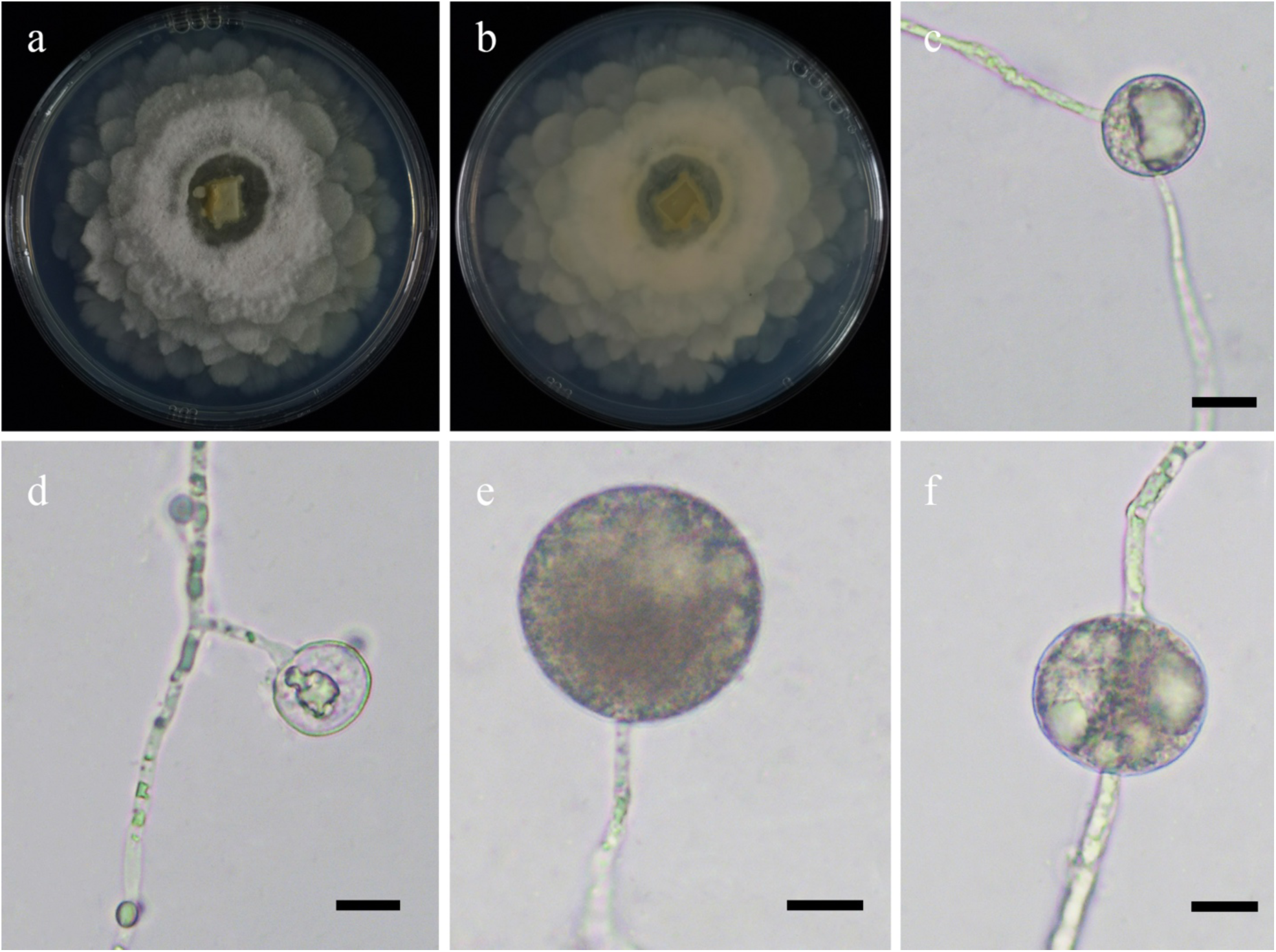
Morphologies of *Mortierella globistylospora* ex-holotype CGMCC 3.16117. **a, b.** Colonies on PDA (**a.** obverse, **b.** reverse); **c, f.** Stylospores on the intercalarily produced; **d, e.** Stylospores terminally produced. — Scale bars: **c–f.** 10 μm.

*Fungal Names*: FN570907.

*Etymology*: *globistylospora* (Lat.) refers to the species having globose stylospores.

*Holotype*: HMAS 351519.

*Colonies* on PDA at 20 °C for 10 days, slow growing, reaching 90 mm in diameter, less than 5 mm high, broadly lobed and zonate, white, irregular margin and Pale Chalcedony Yellow. *Hyphae* branched, dense in the center, 1.5–5.0 μm in diameter. *Stylospores* borne on aerial hyphae, mostly globose, occasionally subglobose, abundant after two weeks, terminal and intercalary, with droplets, subhyaline to hyaline when young, gradually becoming light brow to black with age, smooth, 12.0– 42.0 μm in diameter. *Rhizoids*, s*tolons*, *collar*, *columellae*, *sporangia*, *sporangiospores*, *sporangiophores*, *chlamydospores* and *zygospores* absent.

*Material examined*: China, Tibet Auto Region, Linzhi, Bome County, from soil sample, 28 August 2021, Heng Zhao (holotype HMAS 351519, living ex-holotype culture CGMCC 3.16117).

*GenBank accession number*: OL678156.

*Notes*: *Gamsiella globistylospora* is closely related to *G. stylospora* (Dixon-Stew.) Vandepol & Bonito based on phylogenetic analysis of ITS rDNA sequences (Fig. 9). However, morphologically *G. stylospora* differs from *G. globistylospora* by stylospores bearing spines (Dixon-Stewart 1932, Vandepol *et al*. 2020).

**28. *Gongronella chlamydospora*** H. Zhao, Y.C. Dai & X.Y. Liu, sp. nov., Fig. 44.

**Fig. 44.**
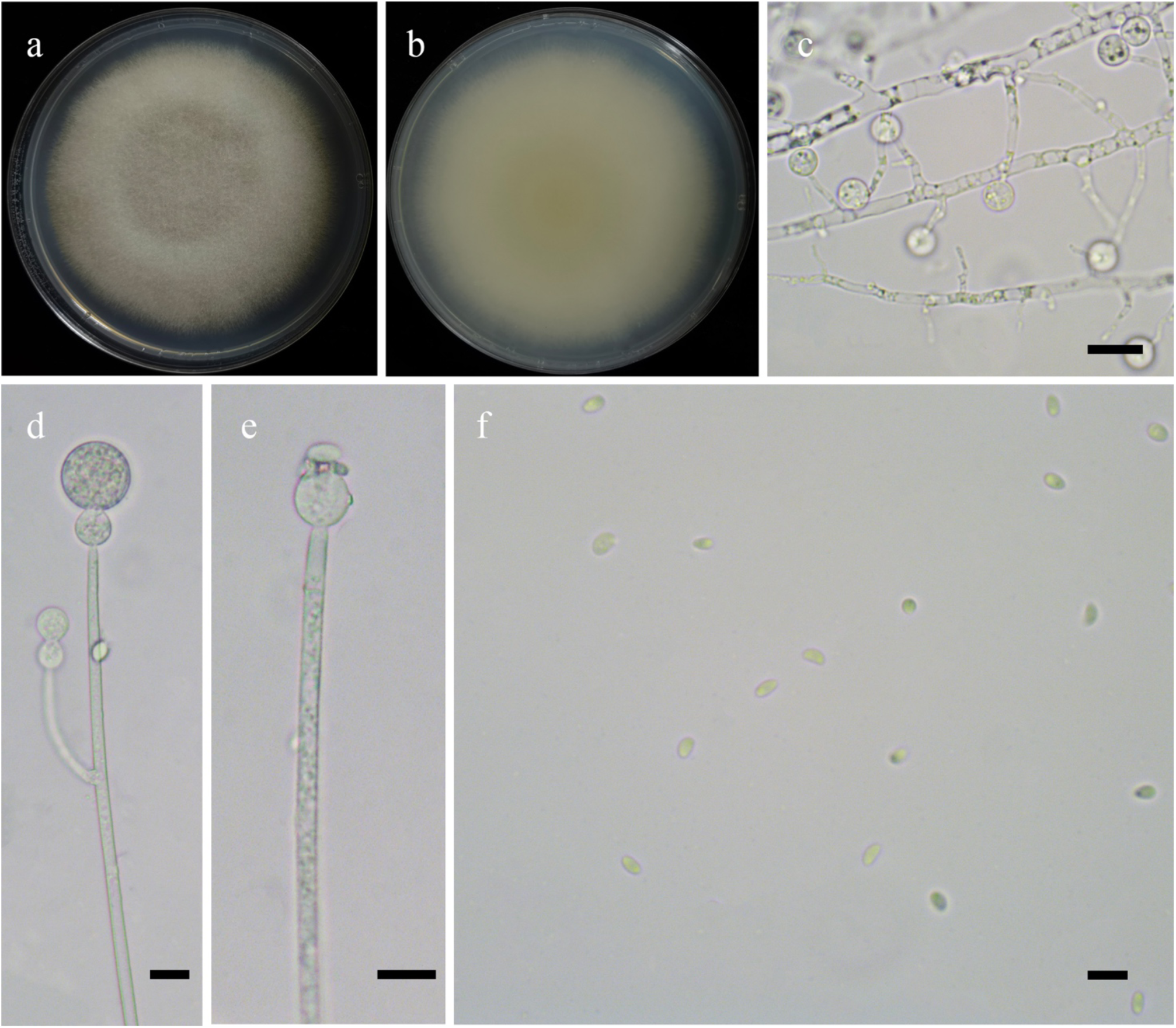
Morphologies of *Gongronella chlamydospora* ex-holotype CGMCC 3.16118. **a, b.** Colonies on PDA (**a.** obverse, **b.** reverse); **c.** Chlamydospores; **d.** Simple branches of sporangiophores; **e.** Columellae; **f.** Sporangiospores. — Scale bars: **c, f.** 5 μm, **d, e.** 10 μm.

*Fungal Names*: FN570908.

*Etymology*: *chlamydospora* (Lat.) refers to the species producing chlamydospores in substrate hyphae.

*Holotype*: HMAS 351520.

*Colonies* on PDA at 27 °C for 11 days, slow growing, reaching 90 mm in diameter, floccose, first white, gradually becoming Drab Gray. *Hyphae* flourishing, branched, aseptate when juvenile, septate with age, 2.5–5.5 μm in diameter. *Rhizoids* and *stolons* present. *Sporangiophores* arising from substrate hyphae or stolons, erect, straight, or bent, unbranched or simple branched, hyaline, slightly constricted at the top, sometimes one septum below the apophysis. *Sporangia* globose, multi-spored, hyaline or subhyaline, 8.5–17.0 μm in diameter. *Apophyses* urn-shaped to subglobose, hyaline or subhyaline, 6.0–12.0 μm long and 6.0–10.0 μm wide. *Collars* present. *Columellae* ovoid to depressed subglobose, hyaline, smooth, 3.0–5.5 μm long and 3.5–6.5 μm wide. *Sporangiospores* ellipsoid, reniform or irregular, 2.0–3.0 μm long and 1.0–2.0 μm wide. *Chlamydospores* in substrate hyphae, mostly globose, occasionally irregular, abundant, 2.0–3.0 μm in diameter. *Zygospores* unknown.

*Material examined*: China, Jiangxi Province, Nanchang, from soil sample, 28 October 2013, Xiao-Yong Liu (holotype HMAS 351520, living ex-holotype culture CGMCC 3.16118).

*GenBank accession number:* OL678157.

*Notes*: *Gongronella chlamydospora* is closely related to *G. butleri* (Lendn.) Peyronel & Dal Vesco based on phylogenetic analysis of ITS rDNA sequences (Fig. 10). However, morphologically *G. butleri* differs from *G. chlamydospore* by larger columellae (4–11 μm in diameter vs. 3.0–5.5 μm long and 3.5–6.5 μm wide), larger sporangia (7–32 μm in diameter vs. 8.5–17.0 μm in diameter), larger chlamydospores (4–9 μm in diameter vs. 2.0–3.0 μm in diameter; Hesseltine & Ellis 1964). In addition, zygospores are produced in *G. butleri*, but not in *G. chlamydospora* (Hesseltine & Ellis 1964).

**29. *Gongronella multispora*** H. Zhao, Y.C. Dai & X.Y. Liu, sp. nov., Fig. 45.

**Fig. 45.**
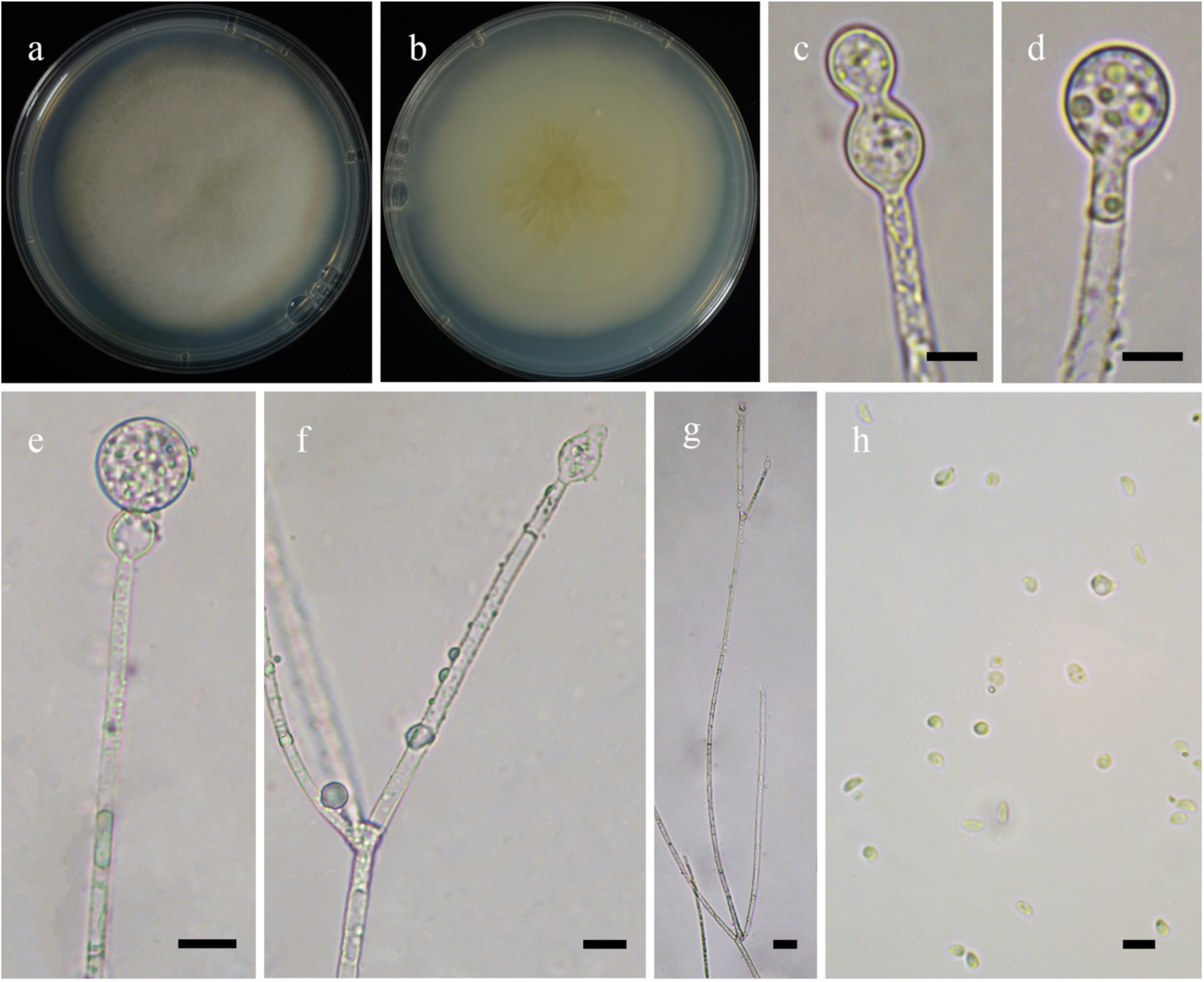
Morphologies of *Gongronella multispora* ex-holotype CGMCC 3.16119. **a, b.** Colonies on PDA (**a.** obverse, **b.** reverse); **c–e.** Sporangia; **f.** Columellae with apophyses; **g.** Sporangiophores showing branches; **h.** Sporangiospores. — Scale bars: **c, d, h.** 5 μm, **e, f.** 10 μm, **g.** 20 μm.

*Fungal Names*: FN570909.

*Etymology*: *multispora* (Lat.) refers to the species having multiple sporangiospores.

*Holotype*: HMAS 351521.

*Colonies* on PDA at 27 °C for 10 days, slow growing, reaching 80 mm in diameter, less than 10 mm high, floccose, first white, gradually becoming Yellowish Glaucous, crusty in reverse, Picric Yellow. *Hyphae* flourishing, branched, aseptate when juvenile, septate with age, 2.0–5.0 μm in diameter. *Rhizoids* rarely present, root-like. *Stolons* present. *Sporangiophores* arising from substrate hyphae or stolons, erect, straight, or slightly bent, unbranched or simple branched, sympodial, 2–3 in whorls and swollen on the base, slightly constricted at the top, often with one septum to several septa below the apophyses. *Sporangia* globose, multi-spored, hyaline or subhyaline, 12.0–17.0 μm in diameter. *Apophyses* pyriform to subglobose, hyaline, 8.0–12.0 μm long and 7.0–9.5 μm wide. *Collars* absent. *Columellae* degenerated, hemispherical, hyaline, smooth, 2.0–4.5 μm long and 2.0– 4.0 μm wide. *Sporangiospores* variably shaped, ellipsoid, fusiform, cylindrical, reniform subglobose to globose, or irregular, 2.5–3.5 μm long and 1.5–2.5 μm wide. *Chlamydospores* absent. *Zygospores* unknown.

*Material examined*: China, Beijing, from soil sample, 28 October 2021, Zhi-Kang Zhang (holotype HMAS 351521, living ex-holotype culture CGMCC 3.16119).

*GenBank accession number*: OL678158.

*Notes*: *Gongronella multispora* is closely related to *G. chlamydospora* H. Zhao *et al*. based on phylogenetic analysis of ITS rDNA sequences (Fig. 10). However, morphologically *G. chlamydospora* differs from *G. multispora* by producing globose chlamydospores on the top of substrate hyphae.

**30. *Lichtheimia alba*** H. Zhao, T.K. Zong, Y.C. Dai & X.Y. Liu, sp. nov., Fig. 46.

**Fig. 46.**
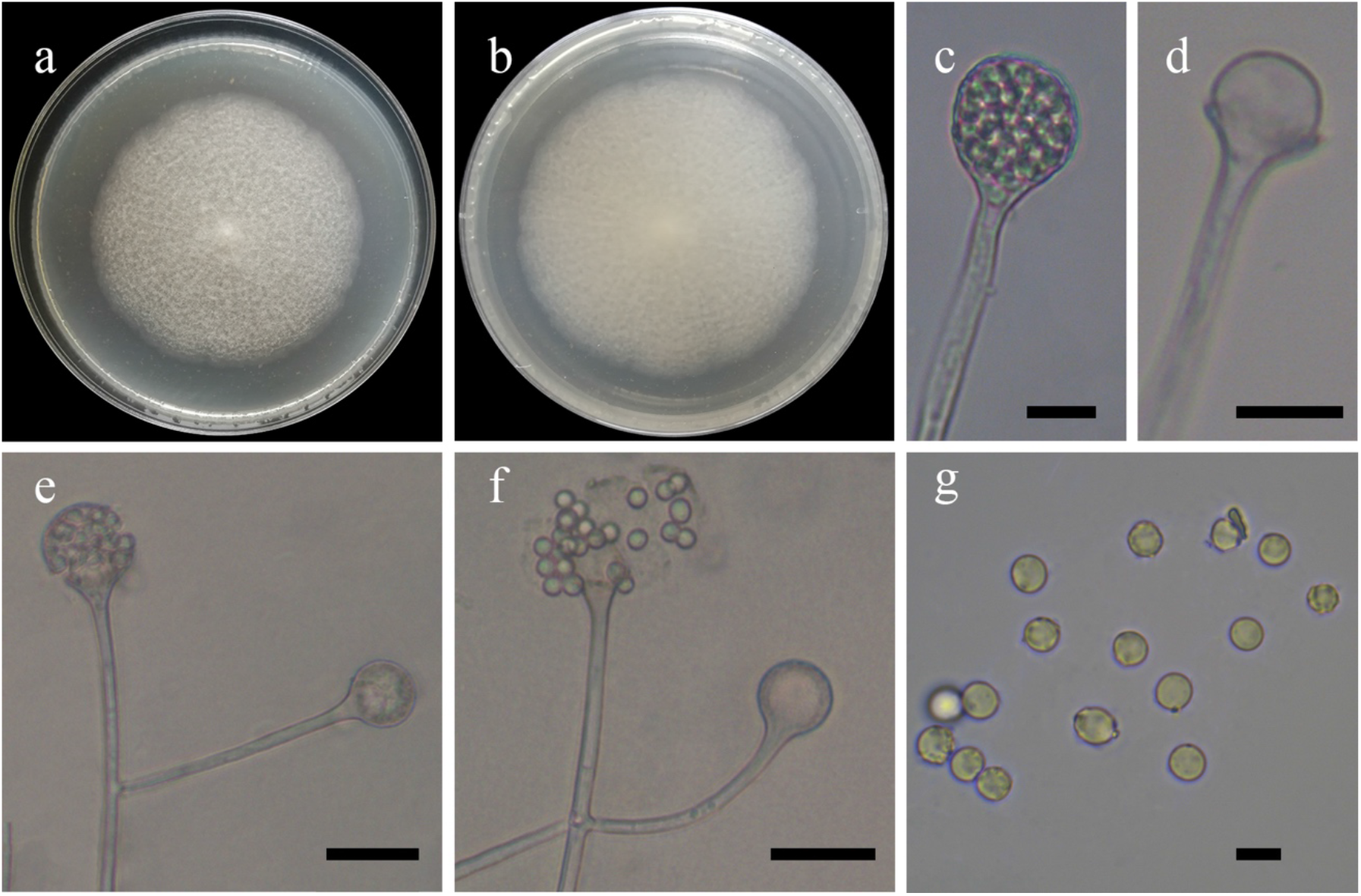
Morphologies of *Lichtheimia alba* ex-holotype CGMCC 3.16120. **a, b.** Colonies on MEA (**a.** obverse, **b.** reverse); **c.** Sporangia; **d.** Columellae; **e, f.** Sporangiophores branched; **g.** Sporangiospores. — Scale bars: **c, d.** 10 μm, **e, f.** 20 μm, **g.** 5 μm.

*Fungal Names*: FN570910.

*Etymology*: *alba* (Lat.) refers to the species forming white colonies.

*Holotype*: HMAS 351522.

*Colonies* on MEA at 30 °C for 8 days, slow growing, reaching 60 mm in diameter, floccose, flat, regular at margin, always white, sporulating profusely yet with sparse aerial hyphae. *Hyphae* hyaline at first, becoming slightly brownish with age, 5.0–11.0 µm wide. *Stolons* branched, hyaline or slightly brownish, smooth, 3.5–7.5 µm in diameter. *Rhizoids* infrequent because of the proliferation of substrate hyphae. *Sporangiophores* erect or slightly bent, never curved or circinate, solitary or in pairs, never in whorls, unbranched, simple, monopodial or sympodial, hyaline, sometimes with a septum 15.0–60.0 µm below apophyses, 50.0–260.0 long and 2.5–6.5 µm wide. *Apophyses* sometimes distinct, funnel-shaped, 6.0–22.0 µm in diameter at the widest point. *Sporangia* globose to ellipsoid, occasionally subpyriform, smooth, multi-spored, 13.5–40.0 long and 13.5–38.5 µm wide, walls deliquescent. *Collars* short if present. *Columellae* spherical to hemispherical, or spatulate, hyaline, smooth, without projections, 8.0–35.0 long and 8.0–32.0 µm wide. *Sporangiospores* mostly globose, occasionally subglobose, hyaline, smooth, uniform, 3.5–5.0 µm in diameter. *Zygospores*, *chlamydospores* and *giant cell* absent. Maximum growth temperature 40 °C.

*Material examined*: China, Beijing, 1 October 2013, from soil sample Xiao-Yong Liu (holotype HMAS 351522 living ex-holotype culture CGMCC 3.16120).

*GenBank accession number*: OL678159.

*Notes*: *Lichtheimia alba* is closely related to *L. corymbifera* (Cohn) Vuill., *L. ornata* (A.K. Sarbhoy) Alastr.-Izq. & Walther and *L. romosa* (Zopf) Vuill. based on phylogenetic analysis of ITS rDNA sequences (Fig. 11). However, physiologically *L. corymbifera*, *L. ornata* and *L. romosa* can grow above 43 °C (Hoffmann 2010).

**31. *Lichtheimia globospora*** H. Zhao, Y.C. Dai & X.Y. Liu, sp. nov., Fig. 47.

**Fig. 47.**
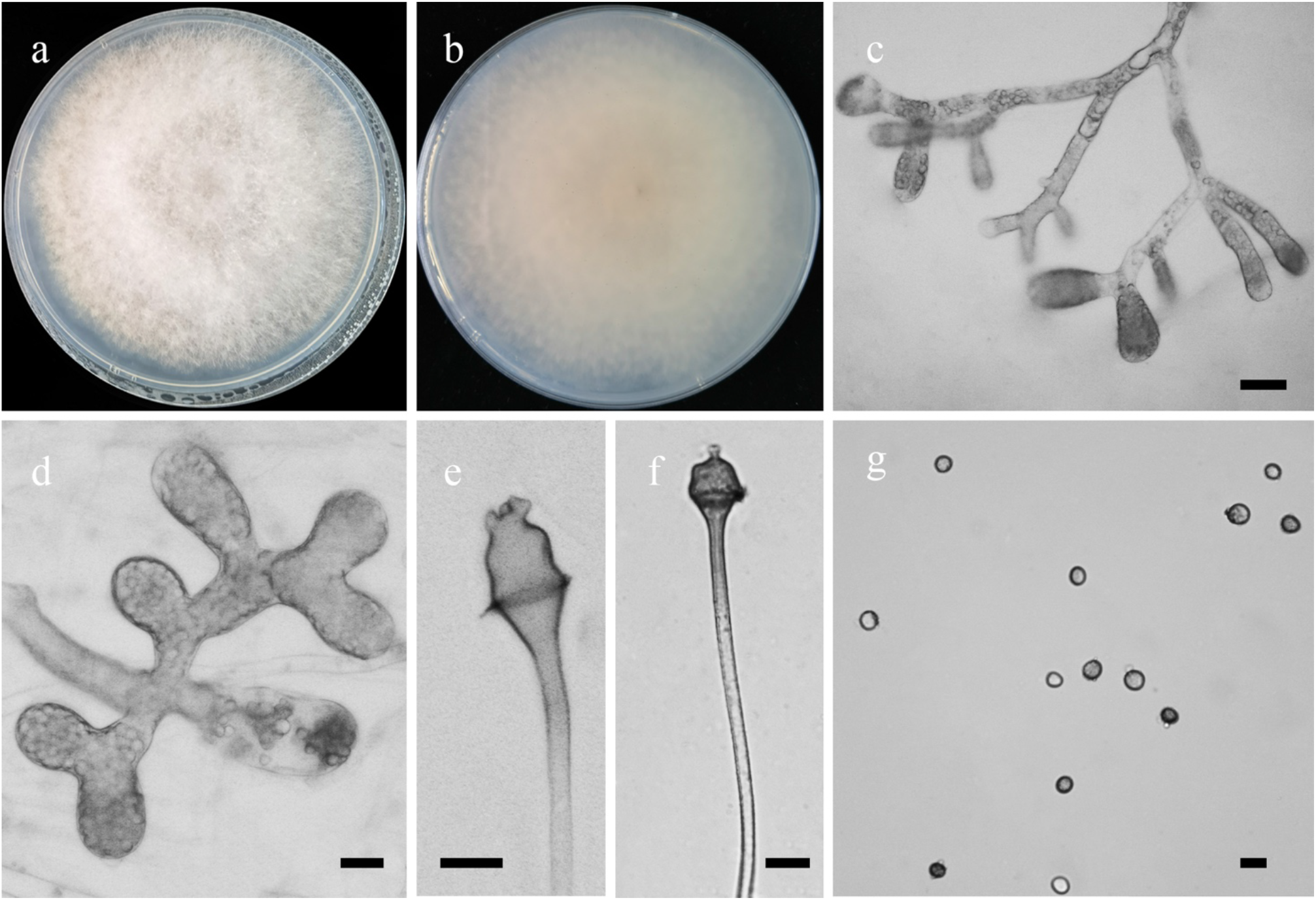
Morphologies of *Lichtheimia globospora* ex-holotype CGMCC 3.16025. **a, b.** Colonies on PDA (**a.** obverse, **b.** reverse); **c, d.** Giant cells in aerial hyphae; **e, f.** Columellae and collars; **g.** Sporangiospores. — Scale bars: **c.** 20 μm, **d–g.** 10 μm.

*Fungal Names*: FN570829.

*Etymology*: *globospora* (Lat.) refers to the species having globose sporangiospores.

*Holotype*: HMAS 249878.

*Colonies* on PDA at 27 °C for 8 days, slow growing, reaching 90 mm in diameter, floccose, initially white, becoming Yellow Ocher with age. *Hyphae* simple branched, aseptate when juvenile, septate with age, with irregularly swollen giant cells, surrounded by liquid droplets, 3.0–20.0 μm in diameter. *Rhizoids* absent. *Stolons* absent. *Sporangiophores* arising from aerial hyphae, not or simple branched, erect, bent or circinate, 50.5–95.5 μm long. *Sporangia* irregular or subglobose, Dark Brown, rough, 19.5–30.5 μm long and 16.0–35.0 μm wide, walls deliquescent. *Apophyses* obvious, 3.5–12.5 μm long and 2.5–11.5 μm wide. *Collars* present. *Columellae* brown, variable, bell-shaped, 4.0–12.5 μm long and 6.5–11.5 μm wide. *Projections* always present, one or two, cylindrical or irregular. *Sporangiospores* hyaline or pigmented, globose, 5.0–7.5 μm in diameter. *Chlamydospores* absent. *Zygospores* absent.

*Material examined*: China, Hainan Province, Shihuashui Cave, from soil sample, 12 November 2020, Jia-Jia Chen (holotype HMAS 249878, living ex-holotype culture CGMCC 3.16025)

*GenBank accession number*: MW580598.

*Notes*: *Lichtheimia globospora* is closely related to *L. hyalospora* (Saito) Kerst. Hoffm. *et al*. based on phylogenetic analysis of ITS rDNA sequences (Fig. 11). However, morphologically *L. hyalospora* differs from *L. globospora* by producing stolons and rhizoids, regularly pyriform sporangia, longer columellae (up to 42.0 μm long vs. 4.0–12.5 μm long), and hemispherical columellae (Hoffmann *et al*. 2009b).

**32. *Modicella abundans*** H. Zhao, Y.C. Dai & X.Y. Liu, sp. nov., Fig. 48.

**Fig. 48.**
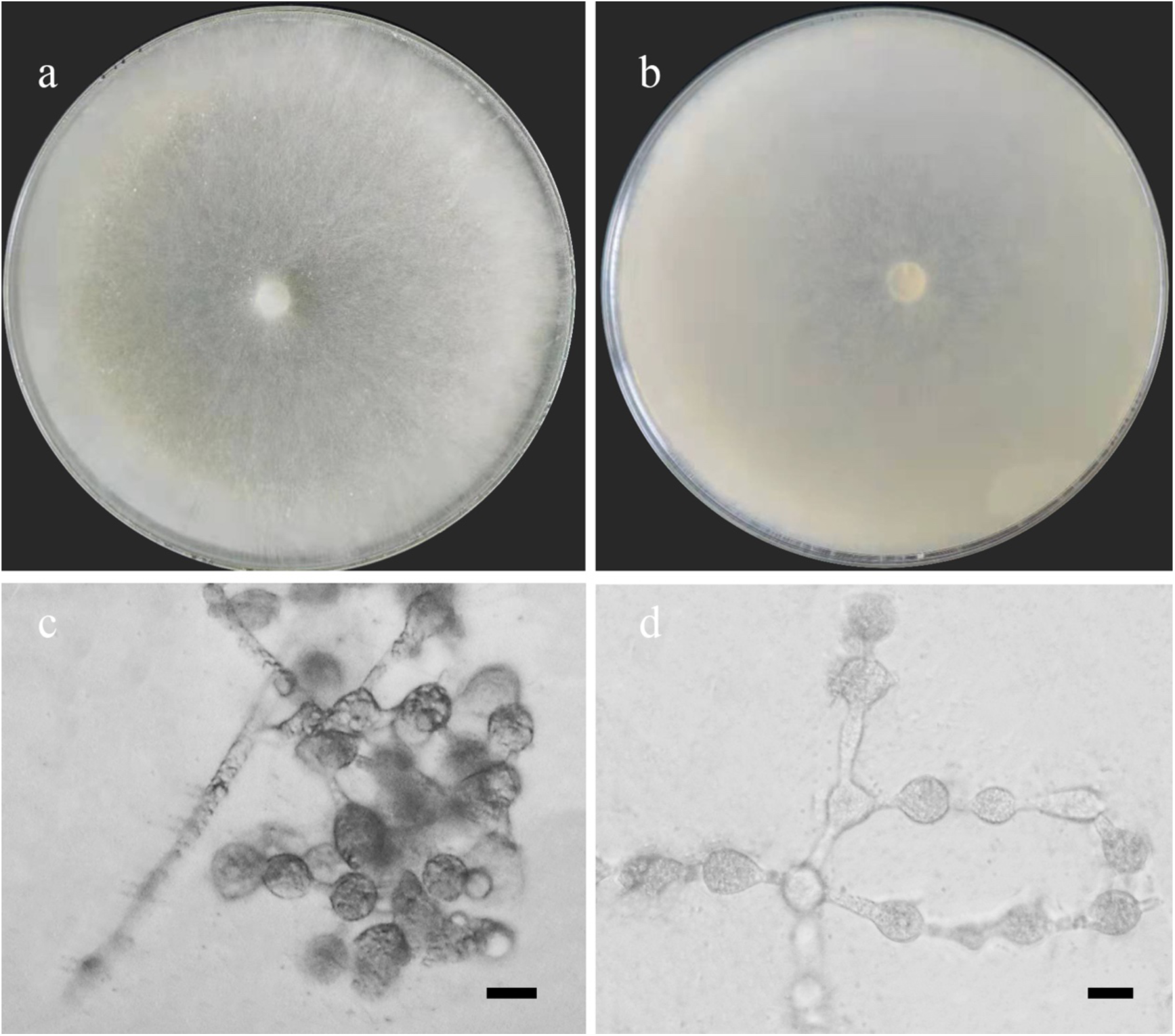
Morphologies of *Modicella abundans* ex-holotype CGMCC 3.16121. **a, b.** Colonies on PDA (**a.** obverse, **b.** reverse); **c, d.** Chlamydospores. — Scale bars: **c, d.** 10 μm.

*Fungal Names*: FN570911.

*Etymology*: *abundans* (Lat.) refers to the species having abundant chlamydospores.

*Holotype*: HMAS 351523.

*Colonies* on PDA at 20 °C for 3 days, fast growing, reaching 90 mm in diameter, devoid of visible growth rings, white to Light Gray, with dense aerial mycelia at edge. *Hyphae* flourishing, white, branched, 2.5–4.5 μm in diameter. *Rhizoids* rarely present, branched, thinner than hyphae, less than 2 μm in diameter. S*tolons* present. *Sporangiophores*, s*porangia*, and s*porangiospores* absent even after one month of growth. *Chlamydospores* variable, ovoid, ovoid, ellipsoid, subglobose, in chains, hyaline or subhyaline, simple branched, 1.5–3.0 μm long and 1.5–2.0 μm wide. *Zygospores* and *sporocarps* unknown.

*Material examined*: China, Anhui Province, Hefei, Lu’an, 115°43’24′′E, 31°27’36′′N, from soil sample, 28 May 2021, Heng Zhao (holotype HMAS 351523, living ex-holotype culture CGMCC 3.16121).

*GenBank accession number*: OL678160.

*Notes*: In this paper, we isolated a culture from a soil sample, which phylogenetically is closer to *Modicella* based on the ITS rDNA sequences (Fig. 12). We describe it as *Modicella abundans*. It is characterized by chlamydospores in chains and by an absence of sporangiospores or sporocarps. These characteristics are unique in members of *Modicella* (Smith *et al*. 2013, Cooper & Park 2020).

**33. *Mortierella amphispora*** H. Zhao, Y.C. Dai & X.Y. Liu, sp. nov., Fig. 49.

**Fig. 49.**
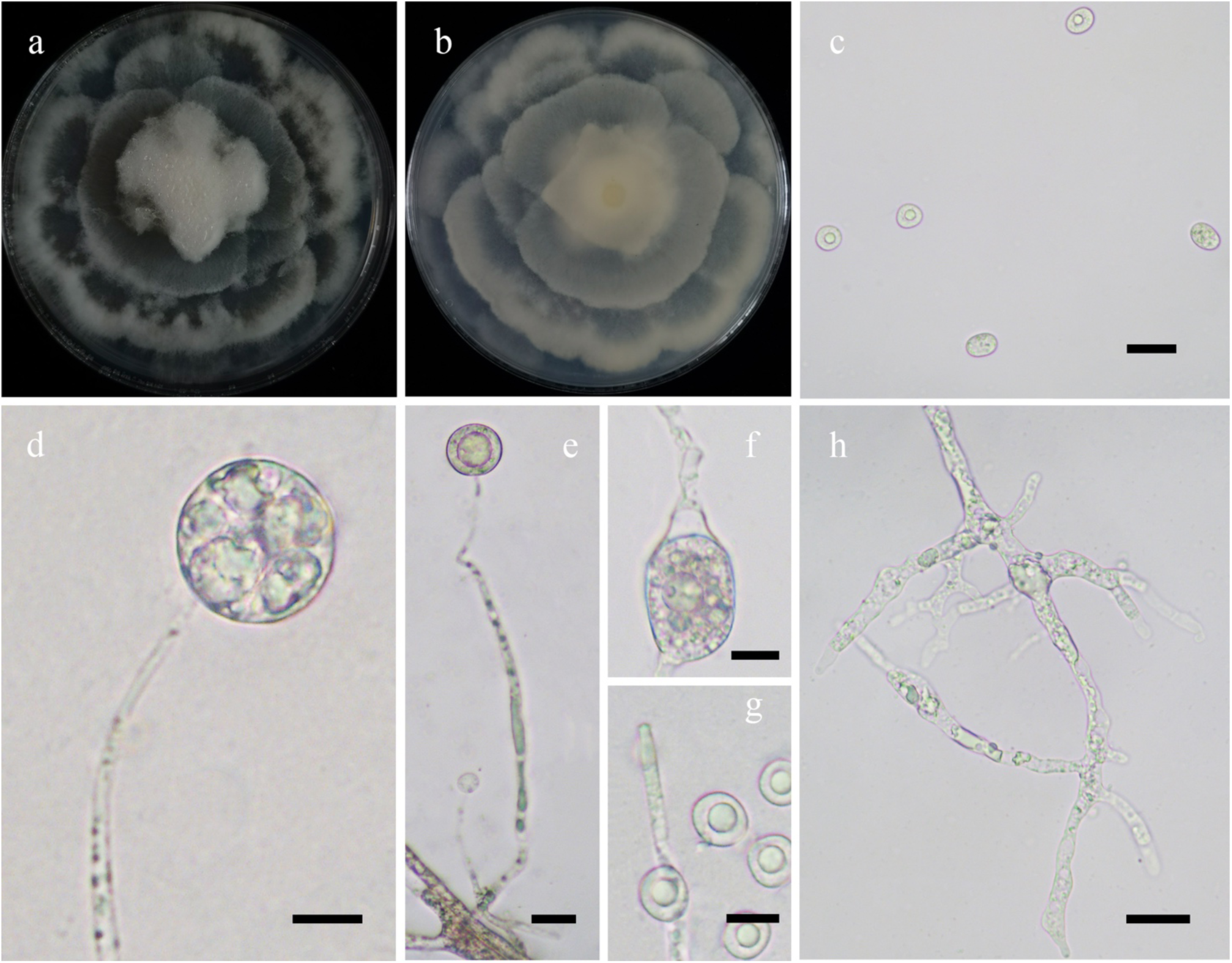
Morphologies of *Mortierella amphispora* ex-holotype CGMCC 3.16122. **a, b.** Colonies on PDA (**a.** obverse, **b.** reverse); **c.** Sporangiospores; **d, e.** Sporangiophores with sporangia; **f.** Chlamydospores; **g.** Degenerated columellae without collars; **h.** Rhizoids. — Scale bars: **c, e, h.** 20 μm, **d, f, g.** 10 μm.

*Fungal Names*: FN570912.

*Etymology*: *amphispora* (Lat.) refers to the species having two kinds of sporangiospores.

*Holotype*: HMAS 351524.

*Colonies* on PDA at 20 °C for 7 days, slow growing, reaching 90 mm in diameter, broadly lobed and concentrically zonate, with dense aerial mycelia in two weeks, white, irregular at margin. *Hyphae* flourishing, branched, 3.0–8.5 μm in diameter. *Rhizoids* rarely present, root-like, branched. S*tolons* present. *Sporangiophores* arising from aerial hyphae, erect, usually unbranched, occasionally simple branched, monopodial, generally constricted at the top. *Sporangia* mostly globose, occasionally subglobose, hyaline or subhyaline, smooth, multi-spored or uni-spored, 14.0– 24.5 μm in diameter, walls deliquescent. *Apophyses* absent. *Collars* absent. *Columellae* degenerated. *Sporangiospores* ovoid or globose, hyaline with a large droplet, or subhyaline with very small droplets, 9.5–15.0 μm long and 8.0–11.5 μm wide. *Chlamydospores* bearing in aerial hyphae, elongate, subhyaline, 16.0–27.5 μm long and 10.5–19.0 μm wide. *Zygospores* unknown.

*Materials examined*: China, Yunnan Province. Diqing, Pudacuo National Park, 27°49’40″N, 99°57’45″E, from soil sample, 24 October 2021, Heng Zhao (holotype HMAS 351524, living ex-holotype culture CGMCC 3.16122, and living culture XY08653). Lijiang, Ninglang Country, 27°31’23″N, 100°44’32″E, from soil sample, 24 October 2021, Heng Zhao (living culture XY08796).

*GenBank accession numbers*: OL678161, OL678162 and OL678292.

*Notes*: *Mortierella amphispora* is closely related to *M. longigemmata* Linnem. based on phylogenetic analysis of ITS rDNA sequences (Fig. 13). However, morphologically *M. longigemmata* differs from *M. amphispora* by shorter sporangiospores (5–9 μm in long vs. 9.5–15.0 μm long), larger chlamydospores (up to 60 μm in diameter vs. 16.0–27.5 μm long and 10.5–19.0 μm wide), and the absence of rhizoids (Zycha *et al*. 1969).

**34. *Mortierella annulata*** H. Zhao, Y.C. Dai & X.Y. Liu, sp. nov., Fig. 50.

**Fig. 50.**
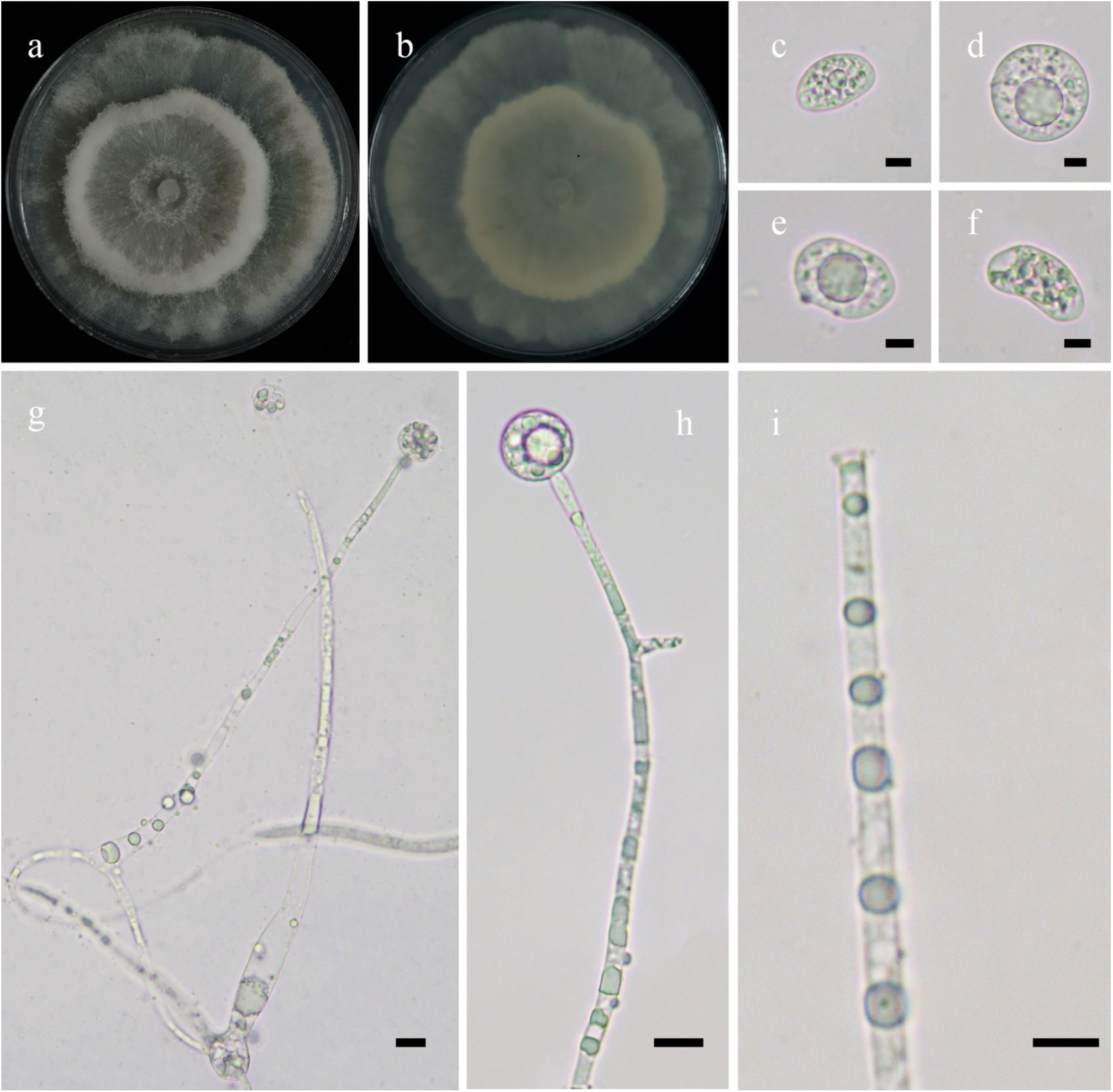
Morphologies of *Mortierella annulata* ex-holotype CGMCC 3.16123. **a, b.** Colonies on PDA (**a.** obverse, **b.** reverse); **c–f.** Sporangiospores; **g, h.** Sporangia; **i**. Sporangiophores without collars and columellae. — Scale bars: **c–f.** 5 μm, **g–i.** 10 μm.

*Fungal Names*: FN570913.

*Etymology*: *annulata* (Lat.) refers to the species producing rings of colonies.

*Holotype*: HMAS 351525.

*Colonies* on PDA at 20 °C for 7 days, reaching 90 mm in diameter, broadly lobed and concentrically zonate, with dense aerial mycelia along the growth ring, white, irregular at margin. *Hyphae* flourishing, branched, 2.5–10.5 μm in diameter. *Rhizoids* absent. *Stolons* absent. *Sporangiophores* arising from aerial hyphae, erect or slightly bent, unbranched or monopodial, usually swollen at the bottom and constricted at the top. *Sporangia* mostly globose, occasionally subglobose, hyaline when young, greenish when old, persistent-walled, 15.0–33.5 μm in diameter. *Apophyses* absent. *Collars* absent. *Columellae* degenerated. *Sporangiospores* ellipsoid, reniform, subglobose, globose or irregular, hyaline or subhyaline, always with droplets, 14.5–21.0 μm long and 9.5–12.5 μm wide. *Chlamydospores* absent. *Zygospores* unknown.

*Material examined*: China, Yunnan Province, Diqing, Pudacuo National Park, 27°49’40″N, 99°57’45″E, from soil sample, 24 October 2021, Heng Zhao (holotype HMAS 351525, living ex-holotype culture CGMCC 3.16123).

*GenBank accession number*: OL678163.

*Notes*: *Mortierella annulata* is closely related to *M. gemmifera* M. Ellis based on phylogenetic analysis of ITS rDNA sequences (Fig. 13). However, morphologically *M. gemmifera* differs from *M. annulata* by producing zygospores on malt agar, and sporangiophores sometimes branched cymosely (Ellis 1940).

**35. *Mortierella cylindrispora*** H. Zhao, Y.C. Dai & X.Y. Liu, sp. nov., Fig. 51.

**Fig. 51.**
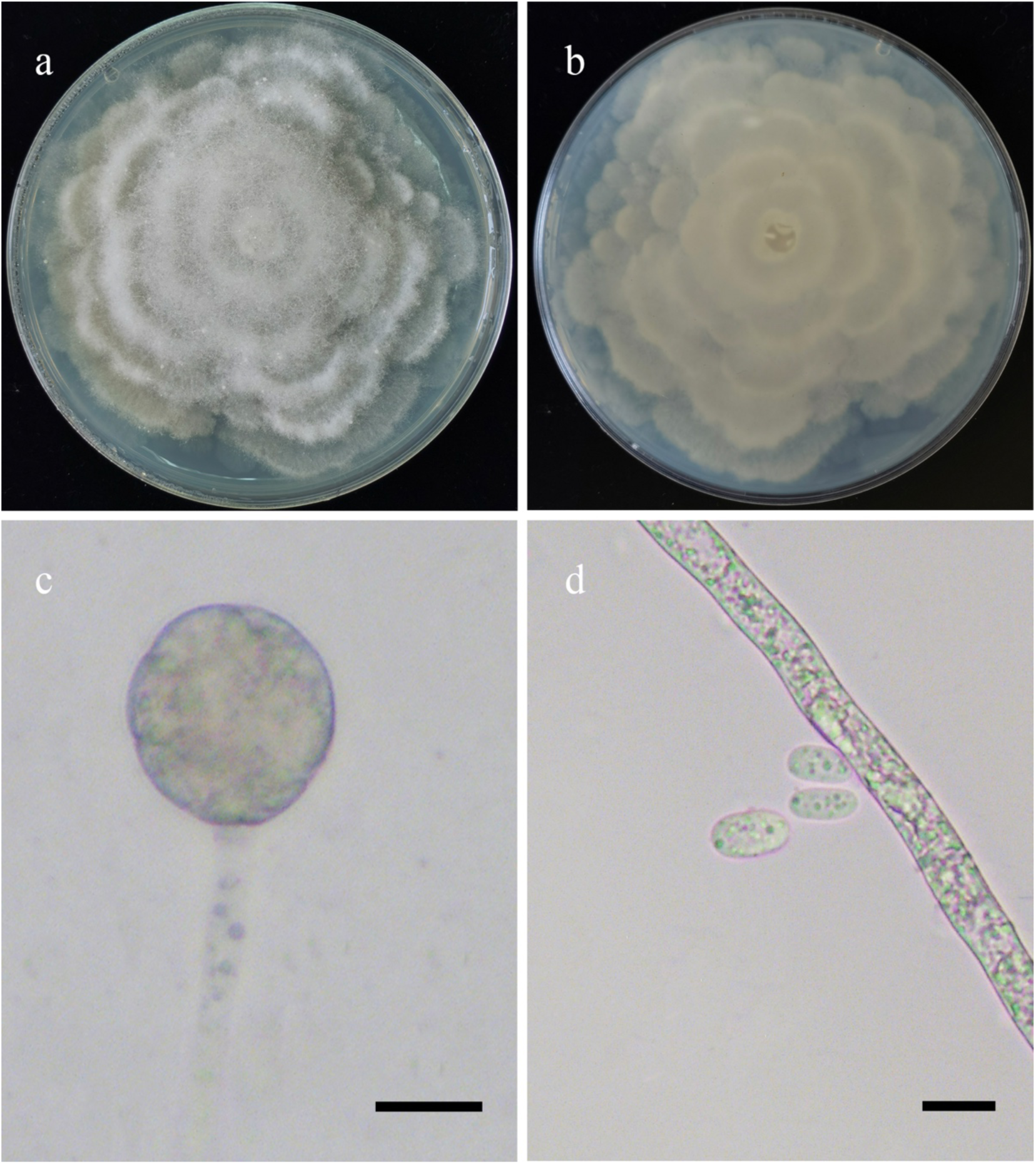
Morphologies of *Mortierella cylindrispora* ex-holotype CGMCC 3.16124. **a, b.** Colonies on PDA (**a.** obverse, **b.** reverse); **c.** Sporangia; **d.** Sporangiospores with hyphae. — Scale bars: **c, d.** 10 μm.

*Fungal Names*: FN570914.

*Etymology*: *cylindrispora* (Lat.) refers to the species having cylindrical sporangiospores.

*Holotype*: HMAS 351526

*Colonies* on PDA at 20 °C for 9 days, slow growing reaching 90 mm in diameter, forming rosettes of dense lobes, white, with dense aerial mycelia. *Hyphae* flourishing, white, branched, aseptate when young, septate when old, 2.0–6.5 μm in diameter. *Rhizoids* present, root-like. S*tolons* present, abundant, simple branched, slightly thin. *Sporangiophores* arising from aerial or substrate hyphae, erect, unbranched, very long. *Sporangia* subglobose or globose, smooth, 15.0–24.5 μm long and 15.0–26.5 μm wide, walls deliquescent. *Apophyses* and *Collars* absent*. Columellae* degenerated. *Sporangiospores* ellipsoid or cylindrical, hyaline, 9.0–16.0 μm long and 4.5–8.0 μm wide. *Chlamydospores* absent. *Zygospores* unknown.

*Material examined*: China, Jiangsu Province, Lianyungang, 119°30’1′′E, 34°38’10′′N, from soil sample, 28 May 2021, Heng Zhao (holotype HMAS 351526, living ex-holotype culture CGMCC 3.16124).

*GenBank accession number*: OL678164.

*Notes*: *Mortierella cylindrispora* is closely related to *M. zychae* Linnem. based on phylogenetic analysis of ITS rDNA sequences (Fig. 13). However, morphologically *M. zychae* differs from *M. cylindrispora* by wider sporangia (30–35 μm wide vs. 5.0–26.5 μm wide), shorter sporangiospores (9.2 μm long vs. 9.0–16.0 μm long), and globose to oblong chlamydospores in the aerial hyphae (Linnemann 1941).

**36. *Mortierella floccosa*** H. Zhao, Y.C. Dai & X.Y. Liu, sp. nov., Fig. 52.

**Fig. 52.**
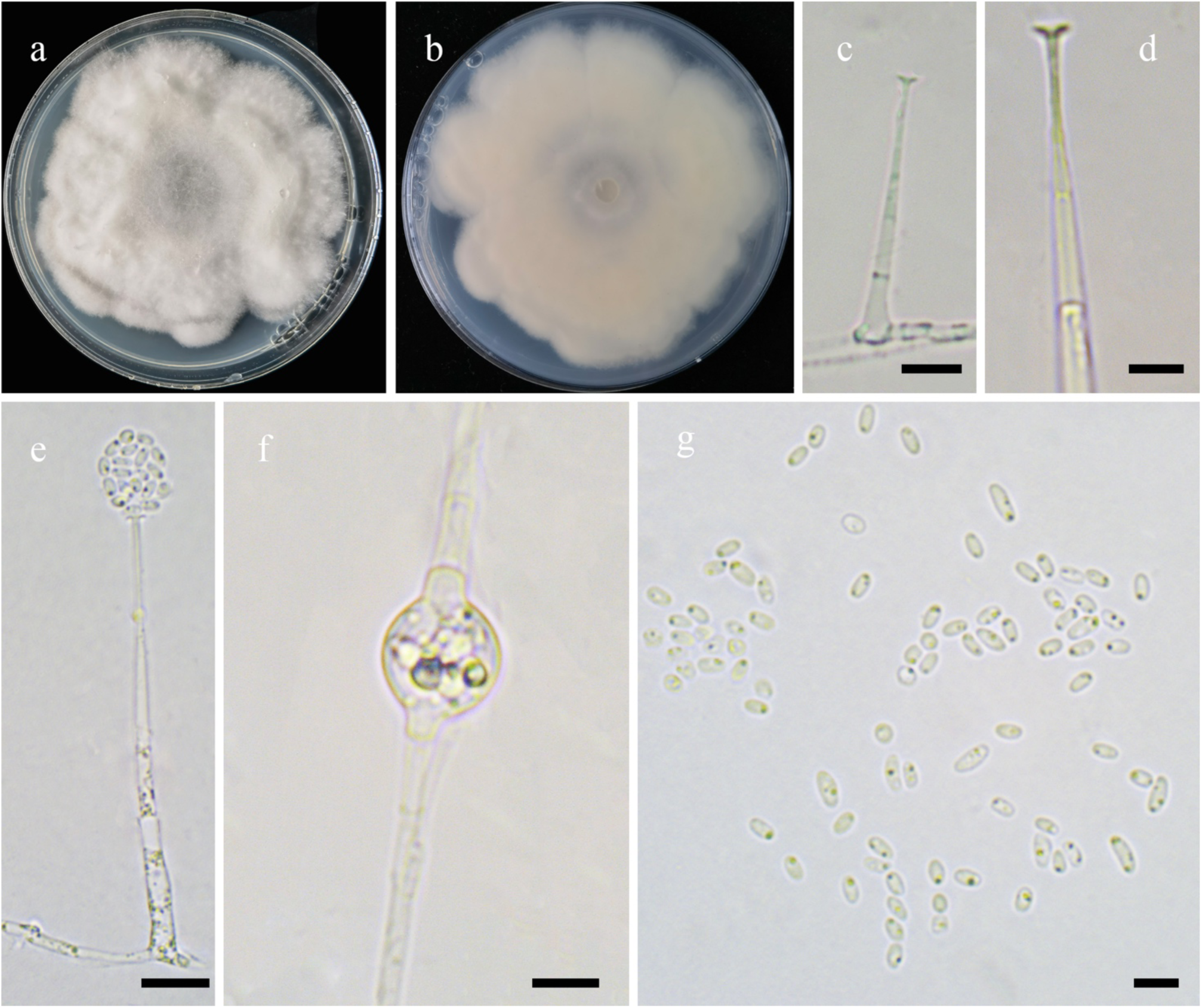
Morphologies of *Mortierella floccosa* ex-holotype CGMCC 3.16125. **a, b.** Colonies on PDA (**a.** obverse, **b.** reverse); **c, d.** Sporangiophores with collars; **e.** Sporangia; **f.** Chlamydospores; **g**. Sporangiospores. — Scale bars: **c, e.** 10 μm, **d, f, g.** 5 μm.

*Fungal Names*: FN570915.

*Etymology*: *floccosa* (Lat.) refers to the species having floccose colonies.

*Holotype*: HMAS 351527.

*Colonies* on PDA at 20 °C for 15 days, slow growing, reaching 90 mm in diameter, broadly lobed and concentrically zonate, floccose, white. *Hyphae* simply branched, hyaline, 1.5–3.0 μm in diameter. *Rhizoids* absent. *Stolons* absent. *Sporangiophores* arising from aerial hyphae, erect, unbranched, usually swollen at bottom and tapering very slightly from the bottom upwards. *Sporangia* mostly globose, occasionally subglobose, hyaline, smooth, multi-spored, 6.5–11.5 μm in diameter, walls deliquescent. *Apophyses* absent. *Collars* present. *Columellae* degenerated. *Sporangiospores* ovoid, cylindrical, fusiform or oblong, hyaline, with some droplets, 2.0–4.5 μm long and 1.0–2.0 μm wide. *Chlamydospores* regular, subglobose or globose, hyaline, always with droplets, 4.5–9.5 μm long and 3.0–9.5 μm wide. *Zygospores* absent.

*Material examined*: China, Jiangsu Province, Lianyungang, 34°38’9″N, 119°29’58″E, from soil sample, 28 May 2021, Heng Zhao (holotype HMAS 351527, living ex-holotype culture CGMCC 3.16125).

*GenBank accession number:* OL678165.

*Notes*: *Mortierella floccosa* is closely related to *M. alpina* Peyronel based on phylogenetic analysis of ITS rDNA sequences (Fig. 13). However, morphologically *M. alpina* differs from *M. floccosa* by producing rhizoids and the absence of chlamydospores (Zycha *et al*. 1969), while rhizoids are absent and chlamydospores are present in *M. floccosa*.

**37. *Mortierella fusiformispora*** H. Zhao, Y.C. Dai & X.Y. Liu, sp. nov., Fig. 53.

**Fig. 53.**
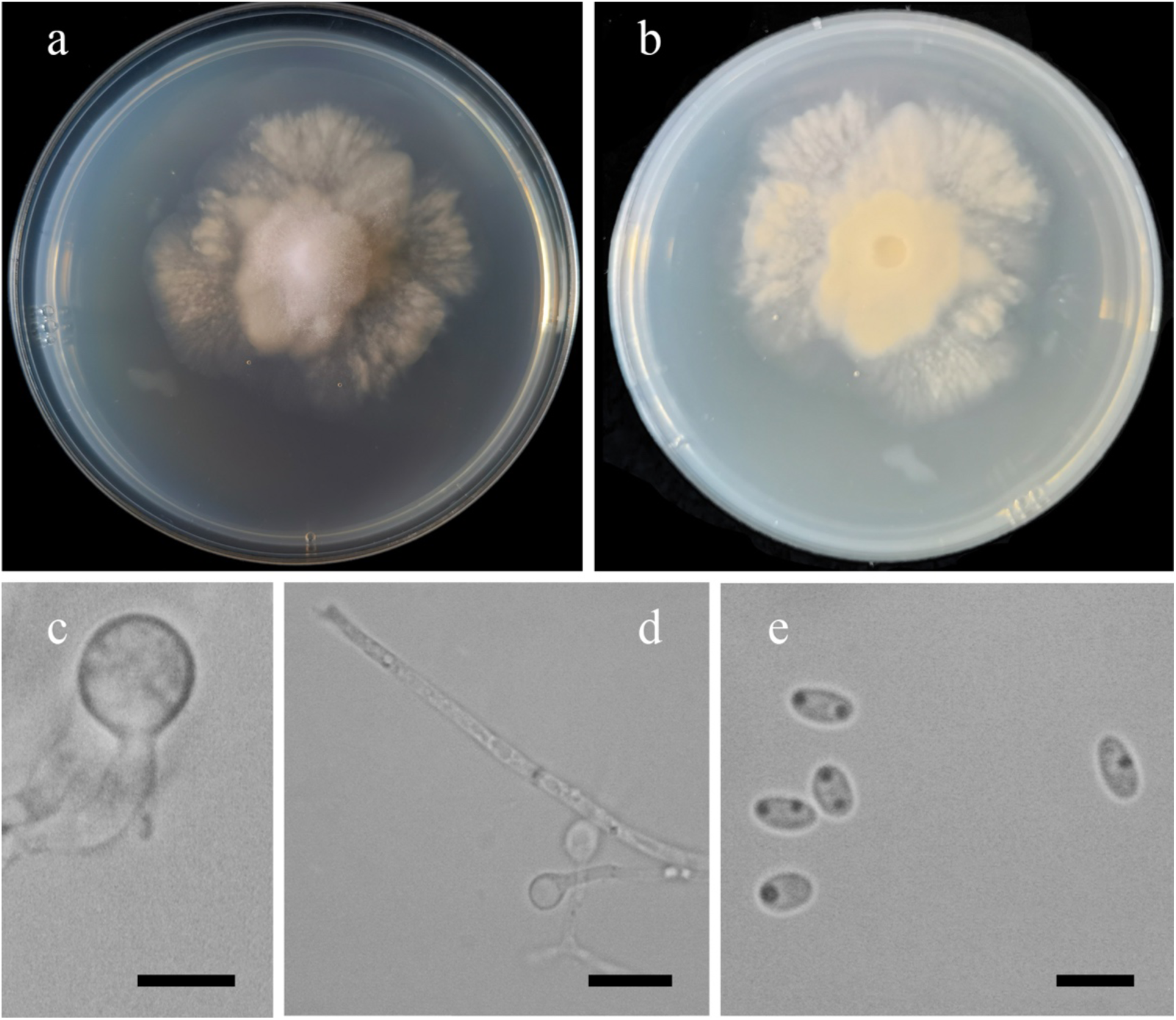
Morphologies of *Mortierella fusiformispora* ex-holotype CGMCC 3.16029. **a, b**. Colonies on PDA (a, obverse, b, reverse); **c.** Immature sporangia; **d**. Collars (red arrow) and sporangiophores; **e**. Sporangiospores showing one or two pigmented spots inside. — Scale bars: **c, e.** 5 μm, **d.** 10 μm.

*Fungal Names*: FN570828.

*Etymology*: *fusiformispora* (Lat.) refers to the species having fusiform sporangiospores.

*Holotype*: HMAS 249877.

*Colonies* on PDA at 20 °C for 10 days, slow growing, reaching 60 mm in diameter, crust-like, white, reverse Pale Yellow-Orange Light to Orange-Yellow. *Hyphae* simple branched, aseptate when juvenile, septate with age, 1.0–4.5 μm in diameter. *Rhizoids* absent. *Stolons* absent. *Sporangiophores* arising from aerial hyphae, erect or bent. *Sporangia* contained in a droplet of water, globose or sub-globose, smooth, 5.0–8.0 μm long and 5.5–7.5 μm wide, walls deliquescent. *Apophyses* absent. *Collars* present. *Columellae* degenerated. *Sporangiospores* fusiform, hyaline, smooth, with one or two droplets, 3.0–4.0 μm long and 1.5–2.5 μm wide. *Chlamydospores* absent. *Zygospores* absent.

*Materials examined*: Antarctica, Fields Peninsula, from soil sample, 27 May 2020, Ze Liu (holotype HMAS 249877, living ex-holotype culture CGMCC 3.16029, and living culture CGMCC 3.16030).

*GenBank accession numbers*: MW580602 and MW580603.

*Notes*: *Mortierella fusiformispora* is closely related to *M. alliacea* Linnem. based on phylogenetic analysis of ITS rDNA sequences (Fig. 13). However, morphologically *M. alliacea* differs from *M. fusiformispora* by the presence of chlamydospores and the absence of sporangiospores (Gams 1977).

**38. *Mortierella lobata*** H. Zhao, Y.C. Dai & X.Y. Liu, sp. nov., Fig. 54.

**Fig. 54.**
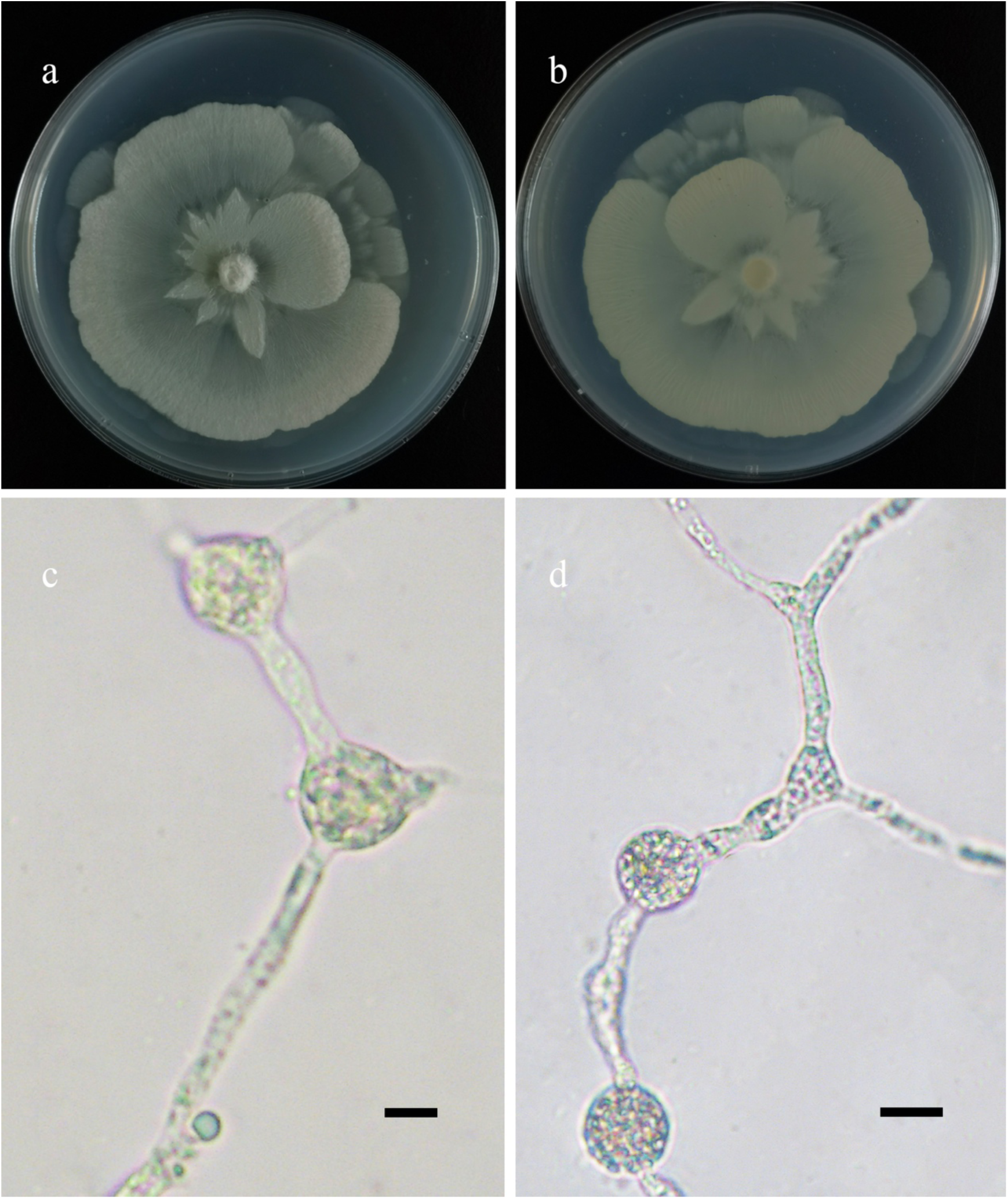
Morphologies of *Mortierella lobata* ex-holotype CGMCC 3.16126. **a, b.** Colonies on PDA (**a.** obverse, **b.** reverse); **c, d.** Chlamydospores. — Scale bars: **c.** 5 μm, **d.** 10 μm.

*Fungal Names*: FN570916.

*Etymology*: *lobata* (Lat.) refers to the species having lobate colonies.

*Holotype*: HMAS 351528.

*Colonies* on PDA at 20 °C for 7 days, slow growing, reaching 70 mm in diameter, broadly lobed and concentrically zonate, with dense aerial mycelia in two weeks, white, irregular at margin *Hyphae* sparse in one week, flourishing in two weeks, branched, hyaline, 1.5–6.0 μm in diameter. *Rhizoids* absent. *Stolons* absent. *Sporangiophores*, s*porangia*, and s*porangiospores* absent even after one month. *Chlamydospores* mainly globose, occasionally subglobose, 6.0–13.5 μm in diameter, rarely ovoid or irregular, 8.0–20 μm long and 6.5–13.5 μm wide, subhyaline or Light Brown, sometimes with papillate appendages, always with droplets. *Zygospores* unknown.

*Material examined*: China, Beijing, 115°25’54′′E, 39°58’6′′N, 6 July 2021, from soil sample, Heng Zhao (holotype HMAS 351528, living ex-holotype culture CGMCC 3.16126).

*GenBank accession number*: OL678166.

*Notes*: *Mortierella lobata* is closely related to *M. seriatoinflata* H. Zhao *et al*. based on phylogenetic analysis of ITS rDNA sequences (Fig. 13). However, morphologically *M. seriatoinflata* differs from *M. lobata* by a faster growth rate under the same conditions (90 mm in diameter vs. 70 mm in diameter) and producing sporangiospores.

**39. *Mortierella macrochlamydospora*** H. Zhao, Y.C. Dai & X.Y. Liu, sp. nov., Fig. 55.

**Fig. 55.**
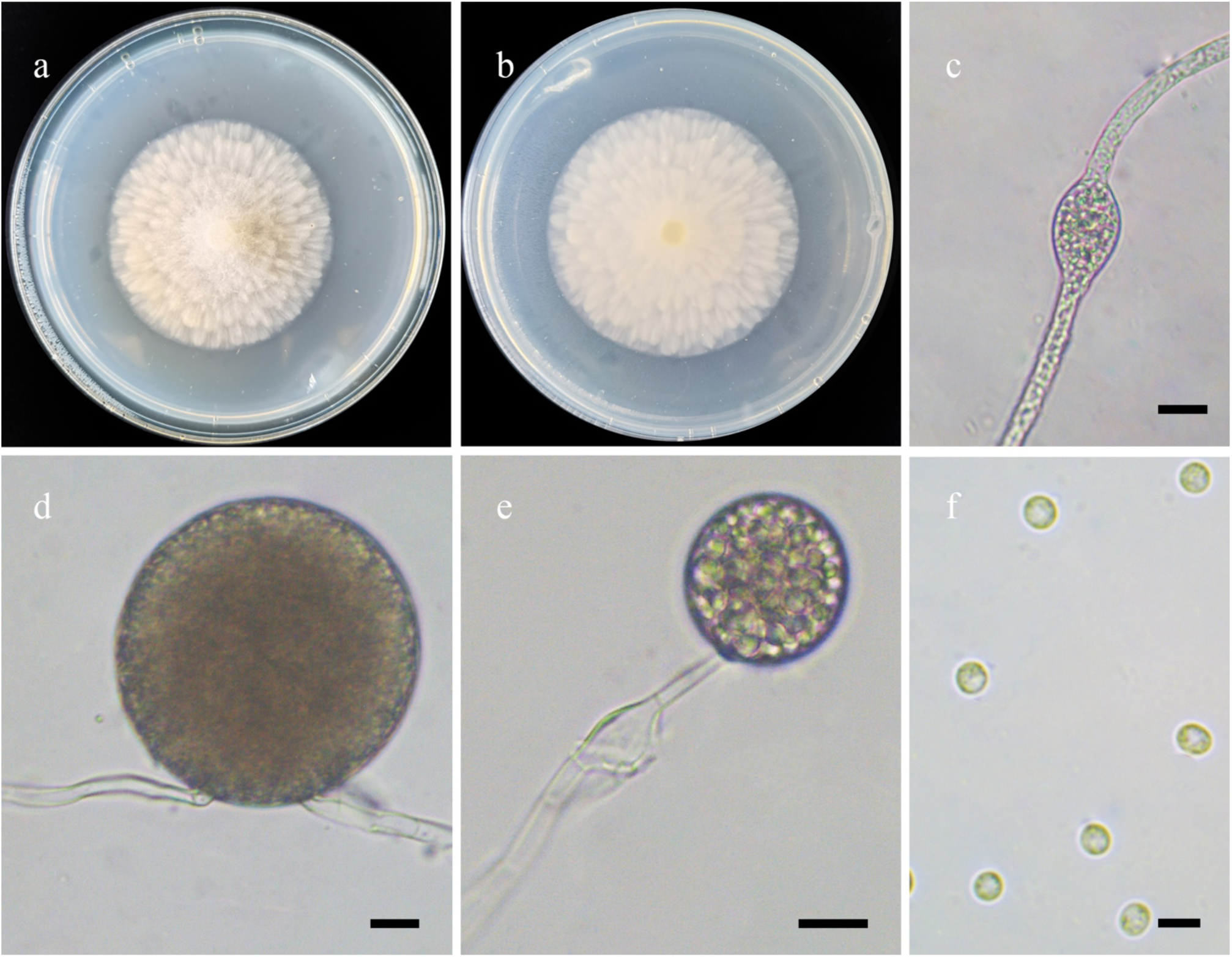
Morphologies of *Mortierella macrochlamydospora* ex-holotype CGMCC 3.16127. **a, b.** Colonies on PDA (**a.** obverse, **b.** reverse); **c.** Hyphal swellings; **d.** Chlamydospores; **e.** Sporangia; **f.** Sporangiospores. — Scale bars: **c–f.** 10 μm.

*Fungal Names*: FN570917.

*Etymology*: *macrochlamydospora* (Lat.) refers to the species having big chlamydospores.

*Holotype*: HMAS 351529.

*Colonies* on PDA at 20 °C for 7 days, slow growing, reaching 55 mm in diameter, scaly and white, reverse regular and Baryta Yellow. *Hyphae* sparse, white, branched, aseptate when young, septate when old, 2.0–7.5 μm in diameter. *Rhizoids* absent. *Stolons* absent. *Sporangiophores* arising from aerial hyphae, erect, unbranched, aseptate. *Sporangia* mostly globose, occasionally subglobose, persistent-walled, smooth, 14.5–30.0 μm in diameter. *Apophyses* absent. *Collars* absent. *Columellae* degenerated. *Sporangiospores* mostly globose, occasionally subglobose, 2.0–3.5 μm in diameter. *Chlamydospores* abundantly produced in media, mostly globose, occasionally subglobose, 22.0–67.0 μm in diameter. *Zygospores* absent.

*Materials examined*: China, Inner Mongolia Auto Region, Hulun Buir, Genhe County, 122°15’48″E, 52°8’21″N, from plant debris, 30 Mar 2018, Peng-Cheng Deng (holotype HMAS 351529, living ex-holotype culture CGMCC 3.16127, and living culture XY04275).

*GenBank accession numbers*: OL678167 and OL678168.

*Notes*: *Mortierella macrochlamydospora* is closely related to *M. macrocystis* W. Gams based on phylogenetic analysis of ITS rDNA sequences (Fig. 13). However, morphologically *M. macrocystis* differs from *M. macrochlamydospora* by larger chlamydospores (20.0–300.0 μm in diameter vs. 22.0–67.0 μm in diameter; Gams 1961, 1977).

**40. *Mortierella microchlamydospora*** H. Zhao, Y.C. Dai & X.Y. Liu, sp. nov., Fig. 56.

**Fig. 56.**
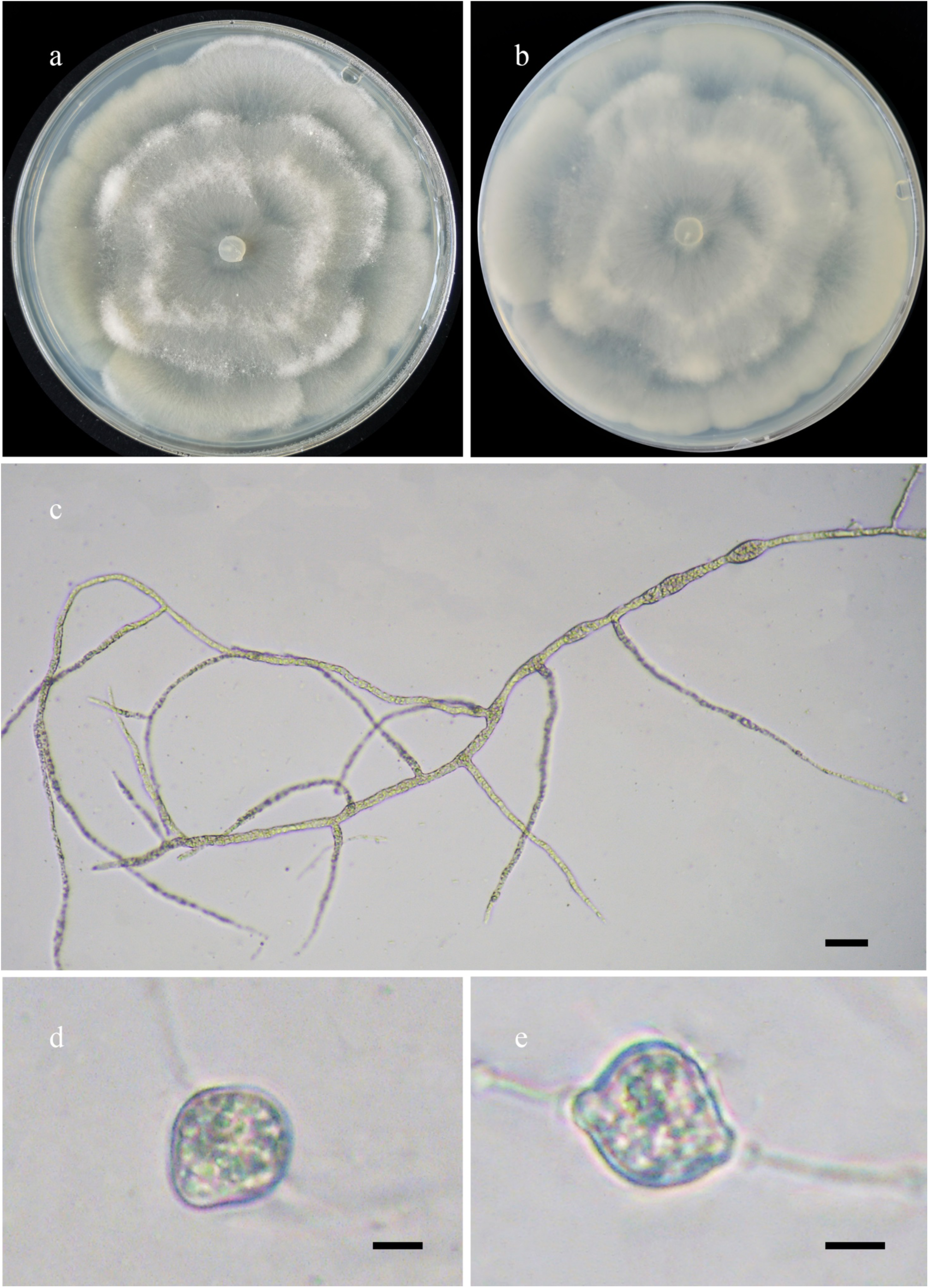
Morphologies of *Mortierella microchlamydospora* ex-holotype CGMCC 3.16128. **a, b.** Colonies on PDA (**a.** obverse, **b.** reverse); **c.** Hyphae branched; **d, e.** Chlamydospores. — Scale bars: **c.** 20 μm, **d, e.** 5 μm.

*Fungal Names*: FN570918.

*Etymology*: *microchlamydospora* (Lat.) refers to the species having the small chlamydospores.

*Holotype*: HMAS 351530.

*Colonies* on PDA at 20 °C for 6 days, reaching 90 mm in diameter, broadly lobed and concentrically zonate, white, with dense aerial mycelia at edge. *Hyphae* flourishing, white, branched, aseptate when young, septate when old, sometimes swollen, 0.5–5.5 μm in diameter. *Rhizoids* absent. *Stolons* absent. *Sporangiophores*, s*porangia*, s*porangiospores* absent after one month of incubation. *Chlamydospores* globose or subglobose or irregular, hyaline or subhyaline, sometimes with spines, obviously thick-walled when mature, 9.5–17.5 μm long and 10–13.5 μm wide. *Zygospores* unknown.

*Materials examined*: China. Yunnan Province, Dehong, Mangshi, 98°40’3′′E, 24°32’38′′N, from soil sample, 28 May 2021, Heng Zhao (holotype HMAS 351530, living ex-holotype culture CGMCC 3.16128). Beijing, 115°25’54′′E, 39°58’6′′N, from soil sample, 30 June 2021, Heng Zhao (culture XY07258); 115°34’1′′E, 39°47’49′′N, from soil sample, 30 June 2021, Heng Zhao (culture XY07283).

*GenBank accession numbers*: OL678169, OL678170 and OL678171.

*Notes*: We isolated three strains (XY06926, XY07258, XY07283) from China, which are closely related to the strain CBS 510.63 in ITS rDNA phylogeny (Fig. 13). The CBS 510.63 was identified as *M. exigua* Linnem. Two other strains of *M. exigua* CBS 655.68 (type of *M. sterilis* B.S. Mehrotra & B.R. Mehrotra) and CBS 358.76, were also deposited in GenBank with ITS rDNA sequences. According to the protologue of *M. exigua*, the sporangiophores are 300 μm long, gradually narrowing, from 5 μm to 1.5 μm wide; the sporangia are globose with deliquescent walls, the sporangiospores are globose or ovoid, 4–7 μm in diameter, and the chlamydospores are globose or hemispherical, 20 μm in diameter (Linnemann 1941). In this study, we proposed *M. microchlamydospora* for these three Chinese strains based on their distinctive characteristics, viz., the absence of sporangiophores, sporangia and sporangiospores and the presence of chlamydospores which are globose, subglobose, irregularly shaped, 9.5–17.5 μm long and 10–13.5 μm wide. Consequently, we treated the strain CBS 510.63 as *M. microchlamydospora* rather than *M. exigua*.

**41. *Mortierella mongolica*** H. Zhao, Y.C. Dai & X.Y. Liu, sp. nov., Fig. 57.

**Fig. 57.**
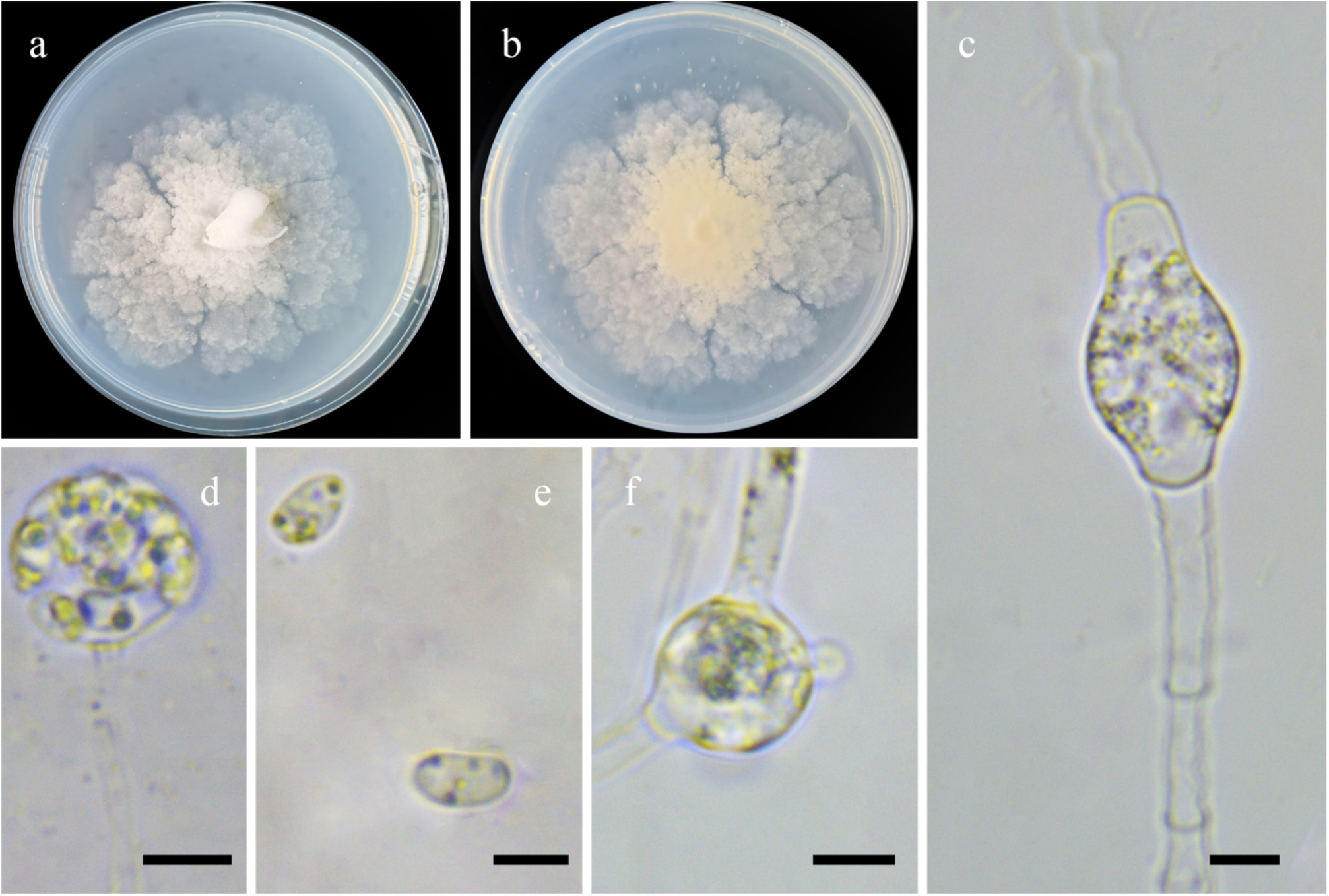
Morphologies of *Mortierella mongolica* ex-holotype CGMCC 3.16129. **a, b.** Colonies on PDA (**a.** obverse, **b.** reverse); **c, f.** Chlamydospores; **d.** Sporangia; **e.** Sporangiospores. — Scale bars: **c–f.** 5 μm.

*Fungal Names*: FN570919.

*Etymology*: *mongolica* (Lat.) refers to the Inner Mongolia Auto Region, China, where the type was collected.

*Holotype*: HMAS 351531.

*Colonies* on PDA at 20 °C for 12 days, slow growing, reaching 80 mm in diameter, lobed and scaly, white, reverse Pale Yellow-Orange. *Hyphae* flourishing, white, branched, sometimes swollen, aseptate when young, septate when old, 1.5–6.0 μm in diameter. *Sporangia* infrequent, subglobose to globose, hyaline or subhyaline, smooth, several-spored, 9.0–11.5 μm long and 8.0–12.5 μm wide, walls deliquescent. *Apophyses* and *Collars* absent. *Columellae* degenerated. *Sporangiospores* ovoid, hyaline, 3.5–6.5 μm long and 3.0–3.5 μm wide. *Chlamydospores* ovoid or globose, hyaline or subhyaline, sometimes with papillate appendages, 9.0–21.0 μm long and 9.5–15.0 μm wide. *Zygospores* unknown.

*Materials examined*: China, Inner Mongolia Auto Region. Zhalantun, 121°39’26′′E, 47°37’45′′N, from soil sample, 7 March 2018, Yu-Chuan Bai (holotype HMAS 351531, living ex-holotype culture CGMCC 3.16129); Arxan, 119°41’16′′E, 47°18’26’’N, from soil sample, 29 January 2018, Yu-Chuan Bai (living culture XY03534).

*GenBank accession numbers*: OL678172 and OL678173.

*Notes*: *Mortierella mongolica* is closely related to *M. antarctica* Linnem. based on phylogenetic analysis of ITS rDNA sequences (Fig. 13). However, morphologically *M. antarctica* differs from *M. mongolica* by subglobose sporangiospores, and globose chlamydospores (Zycha *et al*. 1969, Gams 1977).

**42. *Mortierella seriatoinflata*** H. Zhao, Y.C. Dai & X.Y. Liu, sp. nov., Fig. 58.

**Fig. 58.**
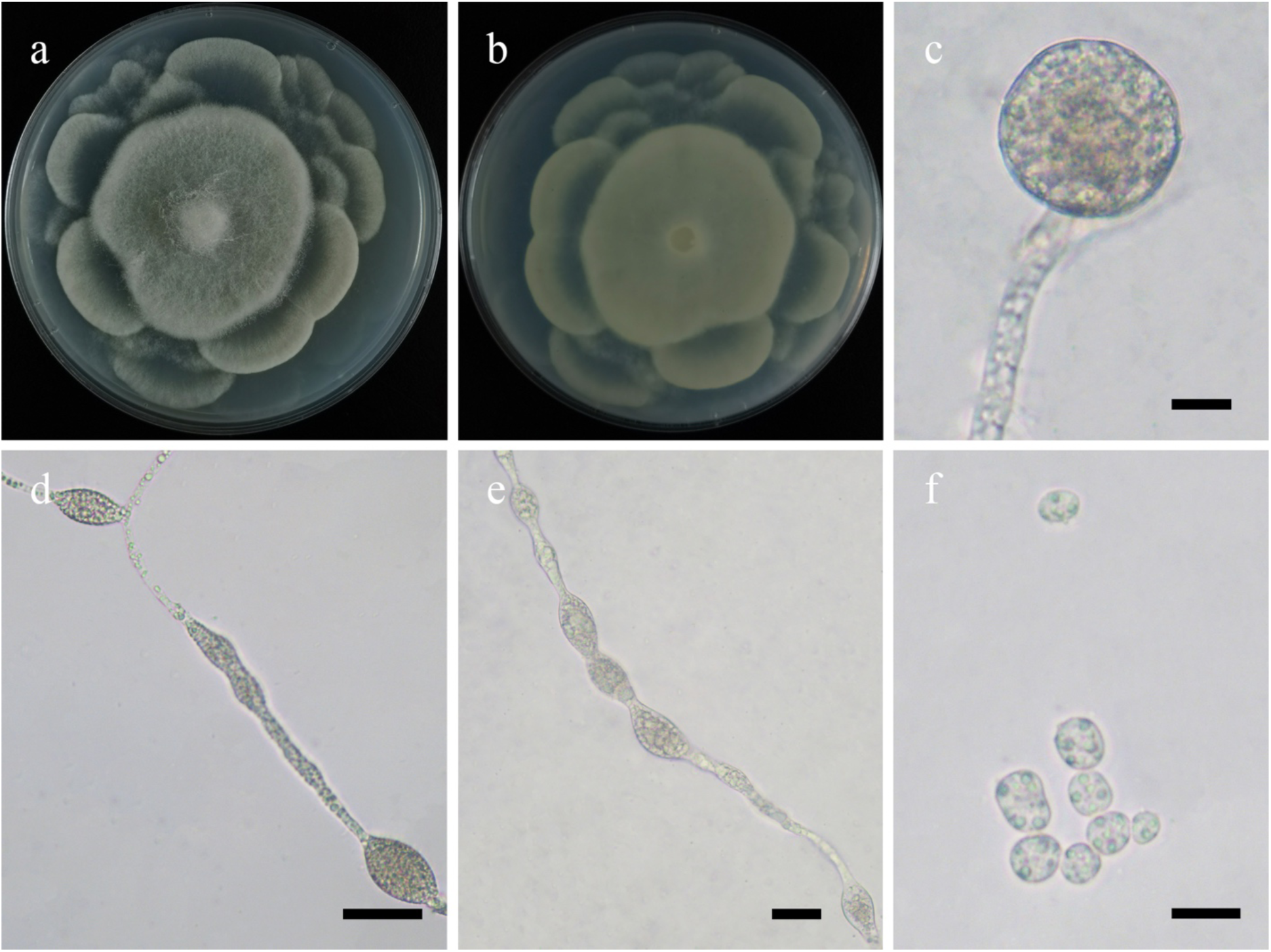
Morphologies of *Mortierella seriatoinflata* ex-holotype CGMCC 3.16130. **a, b.** Colonies on PDA (**a.** obverse, **b.** reverse); **c.** Sporangia; **d, e.** Swellings in hyphae; **f.** Sporangiospores. — Scale bars: **c, f.** 10 μm, **d, e.** 20 μm.

*Fungal Names*: FN570920.

*Etymology*: *seriatoinflata* (Lat.) refers to the species having chains of swellings in the hyphae.

*Holotype*: HMAS 351532.

*Colonies* on PDA at 20 °C for 7 days, reaching 90 mm in diameter, broadly lobed and concentrically zonate, white, with some aerial mycelia mainly in center, sparse growth within one week but abundant within two weeks. *Hyphae* simply branched, hyaline, producing a chain of swellings in both aerial and substrate hyphae, 1.0–7.0 μm in diameter. *Rhizoids* absent. *Stolons* absent. *Sporangiophores* arising from aerial hyphae, erect or slightly bent, unbranched, tapering slightly from bottom upwards. *Sporangia* contain droplets, mostly globose, occasionally subglobose, hyaline or subhyaline, 15.0–30.0 μm in diameter, walls deliquescent or persistent. *Apophyses* absent. *Collars* absent. *Columellae* degenerated. *Sporangiospores* ovoid or subglobose, hyaline, with some droplets, 5.5–11.5 μm long and 4.5–8.5 μm wide. *Chlamydospores* absent. *Zygospores* absent.

*Materials examined*: China. Beijing, 115°25’54′′E, 39°58’6′′N, from plant debris sample, 6 July 2021, Heng Zhao (holotype HMAS 351532, living ex-holotype culture CGMCC 3.16130). Anhui Province, Chuzhou, Quanjiao Country, 117°55’8′′E, 32°0’25′′N, from soil sample, 28 May 2021, Heng Zhao (living culture XY06985). Germany, from forest soil sample, G. Linnemann (living culture CBS 314.52).

*GenBank accession numbers*: OL678174, OL678175 and MH868591.

*Notes*: Two strains XY07264 and XY06985 from China are grouped together with CBS 314.52, which was treated as *Mortierella gamsii* Milko based on phylogenetic analysis of ITS rDNA sequences (Fig. 13). However, type strain of *M. gamsii*, CBS 749.68, is distantly related to these three strains. Therefore, the identification of CBS 314.52 was incorrect. Historically, CBS 314.52 was initially a syntype of *M. spinosa*, but it is incongruent with the typical features of *M. spinosa* Linnem., such as collars and columellae (Linnemann 1941). Consequently, *M. seriatoinflata* was proposed for CBS 314.52 and two Chinese strains.

**43. *Mortierella sparsa*** H. Zhao, Y.C. Dai & X.Y. Liu, sp. nov., Fig. 59.

**Fig. 59.**
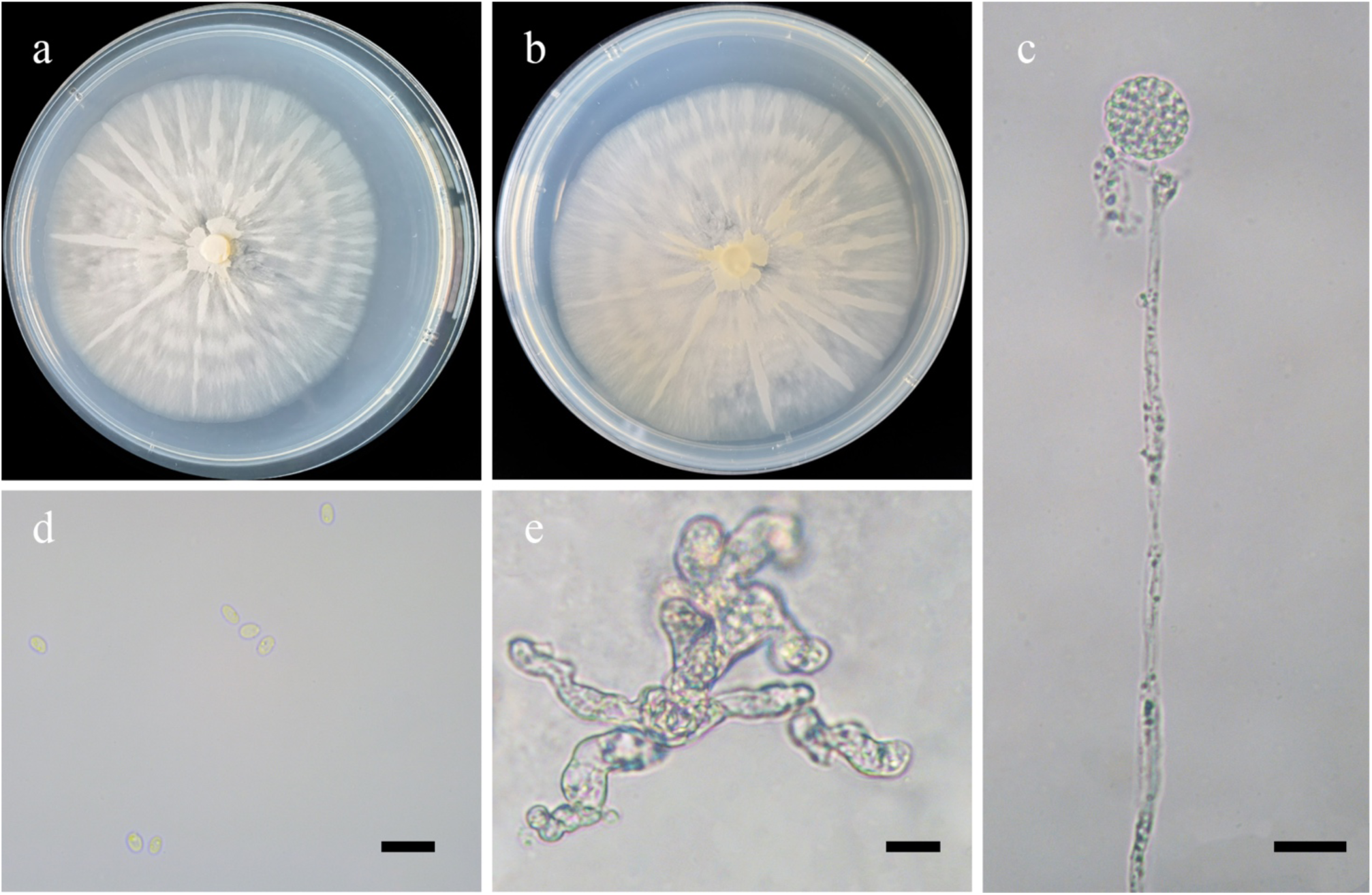
Morphologies of *Mortierella sparsa* ex-holotype CGMCC 3.16131. **a, b.** Colonies on PDA (**a.** obverse, **b.** reverse); **c.** Sporangia; **d.** Sporangiophores; **e.** Swellings in substrate hyphae. — Scale bars: **c.** 20 μm, **d, e.** 10 μm.

*Fungal Names*: FN570921.

*Etymology*: *sparsa* (Lat.) refers to the species having sparse aerial hyphae.

*Holotype*: HMAS 351533.

*Colonies* on PDA at 20 °C for 7 days, slow growing, reaching 60 mm in diameter, radial, broadly lobed, initially white, then Baryta Yellow. *Substrate hyphae* swollen. *Aerial hypha*e sparse, 1.5–5.0 μm in diameter. *Rhizoids* absent. *Stolons* absent. *Sporangiophores* arising from aerial hyphae, erect, unbranched, aseptate. *Sporangia* mostly globose, occasionally subglobose, hyaline or subhyaline, smooth, 15.5–20.0 μm in diameter, walls deliquescent. *Apophyses* and *collars* absent. *Columellae* degenerated. *Sporangiospores* ellipsoid, cylindrical or ovoid, hyaline or subhyaline, 3.0–4.0 μm long and 2.0–2.5 μm wide. *Chlamydospores* absent. *Zygospores* absent.

*Materials examined*: China. Heilongjiang Province, Mohe Country, 52°29′27′′N, 122°31′41′′E, from leaves sample, 21 January 2018, Peng-Cheng Deng (holotype HMAS 35153, living ex-holotype culture CGMCC 3.16131); Huma Country, 52°21′23′′N, 123°59′15′′E, from leaves sample, 6 February 2018, Peng-Cheng Deng (living culture XY03860). Inner Mongolia Auto Region, Argun, 51°53′57′′N, 120°54′33′′E, from leaves sample, 8 April 2018, Peng-Cheng Deng (living culture XY04406).

*GenBank accession numbers*: OL678176, OL678177 and OL678291.

*Notes*: *Mortierella sparsa* is closely related to *M. basiparvispora* W. Gams & Grinb., *M. jenkinii* (A.L. Sm.) Naumov and *M. parvispora* Linnem. based on phylogenetic analysis of ITS rDNA sequences (Fig. 13). However, morphologically *M. basiparvispora* and *M. parvispora* differ from *M. sparsa* by subglobose to globose of sporangiospores (Gams 1976, 1977). *M. jenkinii* differs from *M. sparsa* by producing lemon-shaped chlamydospores (Gams 1977).

**44. *Mortierella varians*** H. Zhao, Y.C. Dai & X.Y. Liu, sp. nov., Fig. 60.

**Fig. 60.**
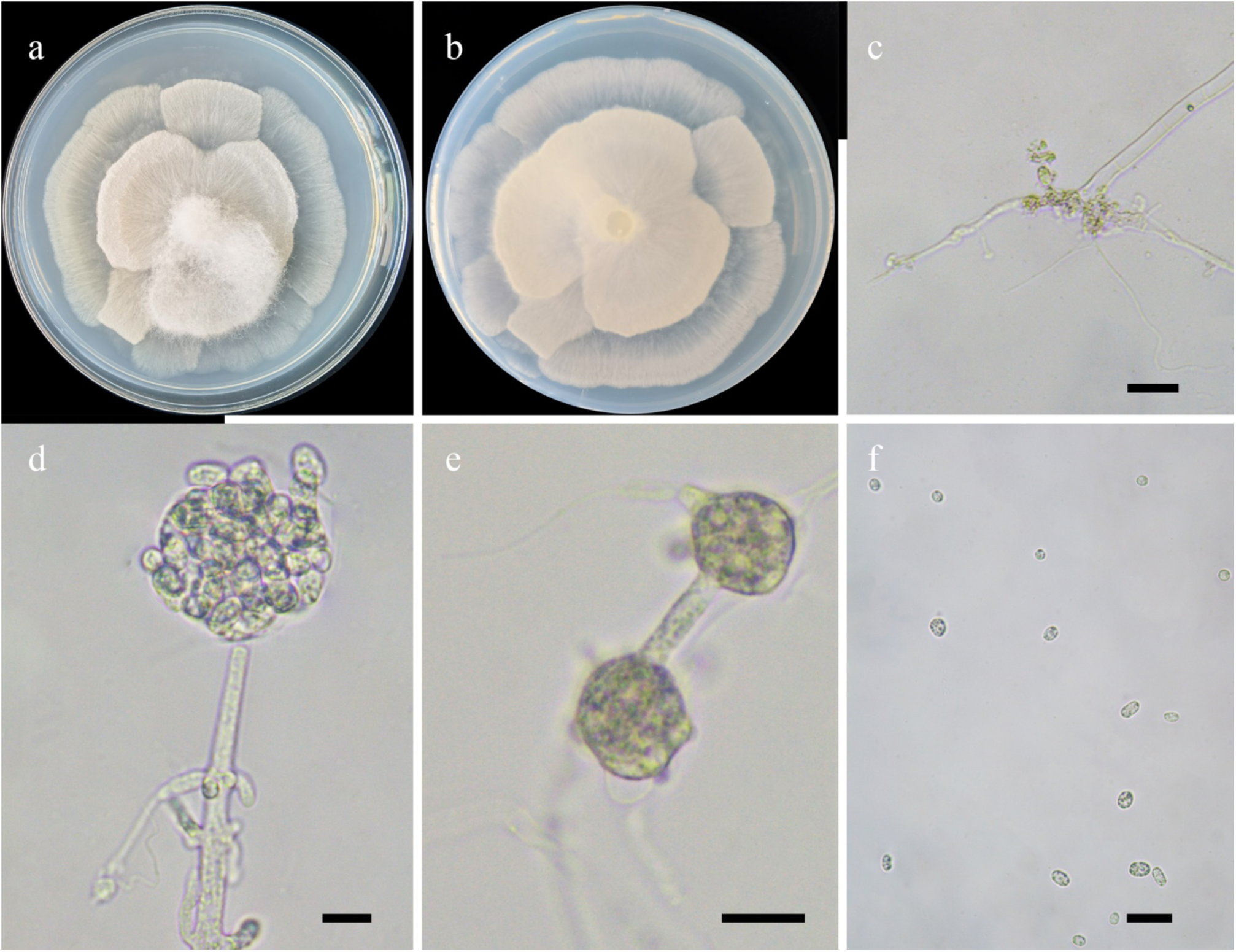
Morphologies of *Mortierella varians* ex-holotype CGMCC 3.16132. **a, b.** Colonies on PDA (**a.** obverse, **b.** reverse); **c.** Rhizoids; **d.** Sporangia; **e.** Chlamydospores; **f.** Sporangiospores. — Scale bars: **c, f.** 20 μm, **d, e.** 10 μm.

*Fungal Names*: FN570922.

*Etymology*: *varians* (Lat.) refers to the species having various sporangiospores in shape.

*Holotype*: HMAS 351534.

*Colonies* on PDA at 20 °C for 7 days, slow growing, reaching 80 mm in diameter, broadly lobed zonate or umbrella-shaped, white. *Hyphae* sparse when young, flourishing when old, white, branched, septate with age, 1.5–13.5 μm in diameter. *Rhizoids* root-like, simple branched, slightly thin. *Sporangiophores* arising from aerial hyphae, erect or slightly bent, unbranched, aseptate. *Sporangia* subglobose or globose, hyaline or subhyaline, smooth, 20.5–37.5 μm long and 22.0–36.0 μm wide, walls deliquescent. *Apophyses* and *Collars* absent*. Columellae* degenerated. *Sporangiospores* variable, ovoid or ellipsoid or globose or subglobose, 5.0–14.0 μm long and 4.0– 8.0 μm wide. *Chlamydospores* irregular or subglobose or ovoid, 9.0–15.0 μm long and 7.0–13.0 μm wide. *Zygospores* absent.

*Materials examined*: China, Tibet Auto Region, Naqu, from soil sample, 21 Dec 2020, Heng Zhao (holotype HMAS 351534, living ex-holotype culture CGMCC 3.16132, and living culture XY05920).

*GenBank accession numbers*: OL678178 and OL678179.

*Notes*: *Mortierella varians* is closely related to *M. acrotona* W. Gams based on phylogenetic analysis of ITS rDNA sequences (Fig. 13). However, morphologically *M. acrotona* differs from *M. varians* by sporangia containing 1–4 sporangiospores which are variable in size (11–24 μm in diameter; Gams 1976), while sporangia contain abundant sporangiospores which are stable in size (17–20 μm in diameter) in *M. varians*.

**45. *Mucor abortisporangium*** H. Zhao, Y.C. Dai & X.Y. Liu, sp. nov., Fig. 61.

**Fig. 61.**
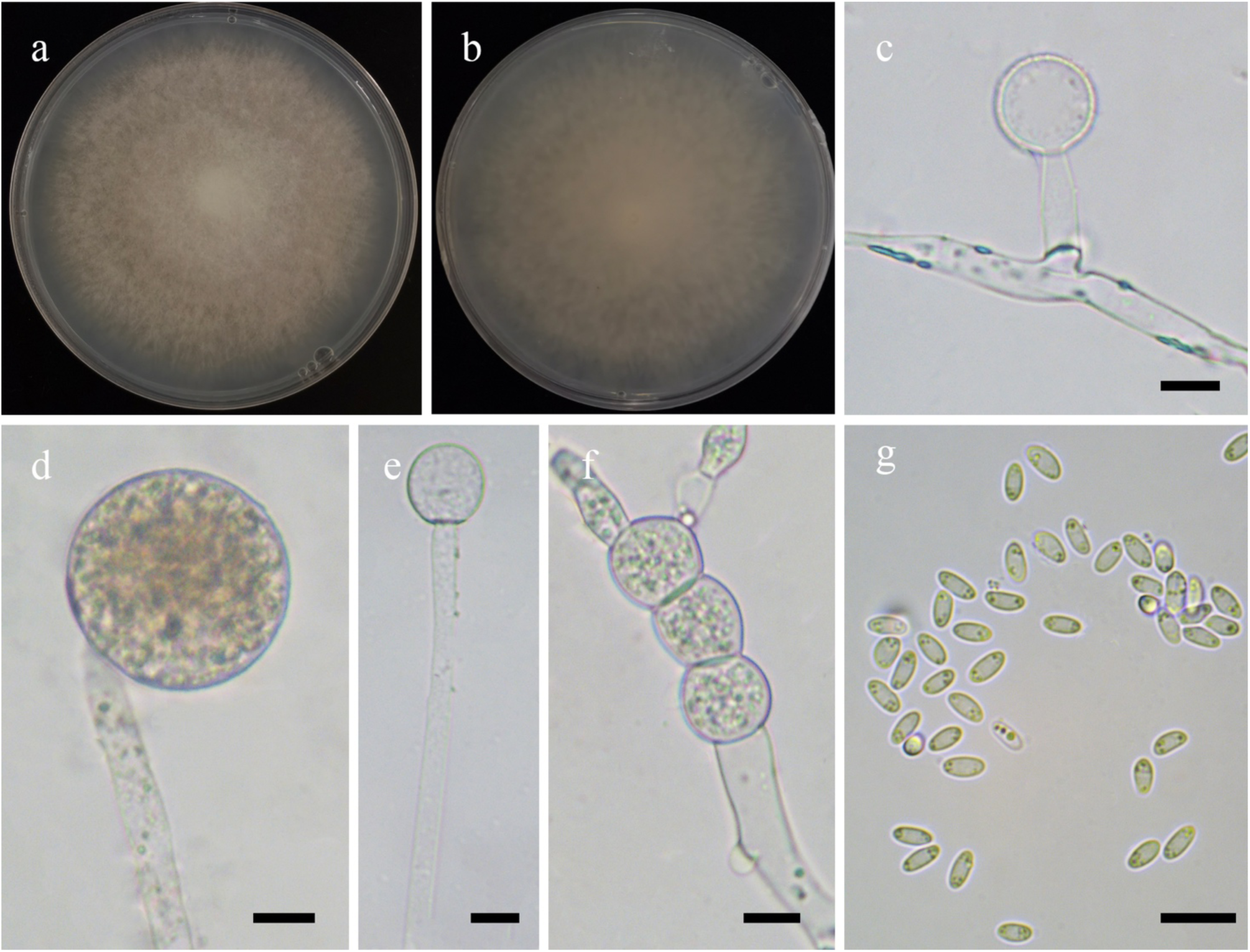
Morphologies of *Mucor abortisporangium* ex-holotype CGMCC 3.16133. **a, b.** Colonies on PDA (**a.** obverse, **b.** reverse); **c.** Aborted lateral sporangia; **d.** Sporangia; **e.** Columellae; **f.** Chlamydospores; **g.** Sporangiospores. — Scale bars: **c–g.** 10 μm.

*Fungal Names*: FN570923.

*Etymology*: *abortisporangium* (Lat.) refers to the species having sporangia failing to mature.

*Holotype*: HMAS 351535.

*Colonies* on PDA at 27 °C for 7 days, reaching 80 mm in diameter, first white, gradually becoming Cinnamon-Drab, floccose, irregular at margin. *Hyphae* flourishing, branched, aseptate when juvenile, septate with age, 2.0–10.0 μm in diameter. *Rhizoids* absent. *Stolons* absent.*Sporangiophores* arising from substrate and aerial hyphae, erect or slightly curved, unbranched. *Fertile sporangia* globose, smooth, 19.0–30.0 μm in diameter, walls deliquescent. *Aborted sporangia* laterally formed on aerial hyphae, hyaline. *Apophyses* absent. *Collars* absent. *Columellae* subglobose or globose, hyaline, 11.0–25.5 μm long and 10.5–24.0 μm wide. *Sporangiospores* fusiform or ellipsoid, with droplets, 4.5–5.5 μm long and 2.0–3.0 μm wide. *Chlamydospores* produced in substrate hyphae, in chains, cylindrical, ellipsoid, ovoid, subglobose, globose or irregular, 5.0–22.5 μm long and 6.5–15.5 μm wide. *Zygospores* unknown.

*Material examined*: China, Guangxi Auto Region, Guigang, from soil sample, 16 September 2021, Heng Zhao (holotype HMAS 351535 living ex-holotype culture CGMCC 3.16133).

*GenBank accession number*: OL678180.

*Notes*: *Mucor abortisporangium* is closely related to *M. radiatus* H. Zhao *et al*. based on phylogenetic analysis of ITS rDNA sequences (Fig. 14). However, morphologically *M. radiatus* differs from *M. abortisporangium* by a slower growth rate (60 mm within 10 days vs. 80 mm within 7 days), abundant chlamydospores borne in the substrate hyphae and aerial hyphae (in substrate hyphae only in *M. abortisporangium*).

**46. *Mucor amphisporus*** H. Zhao, Y.C. Dai & X.Y. Liu, sp. nov., Fig. 62.

**Fig. 62.**
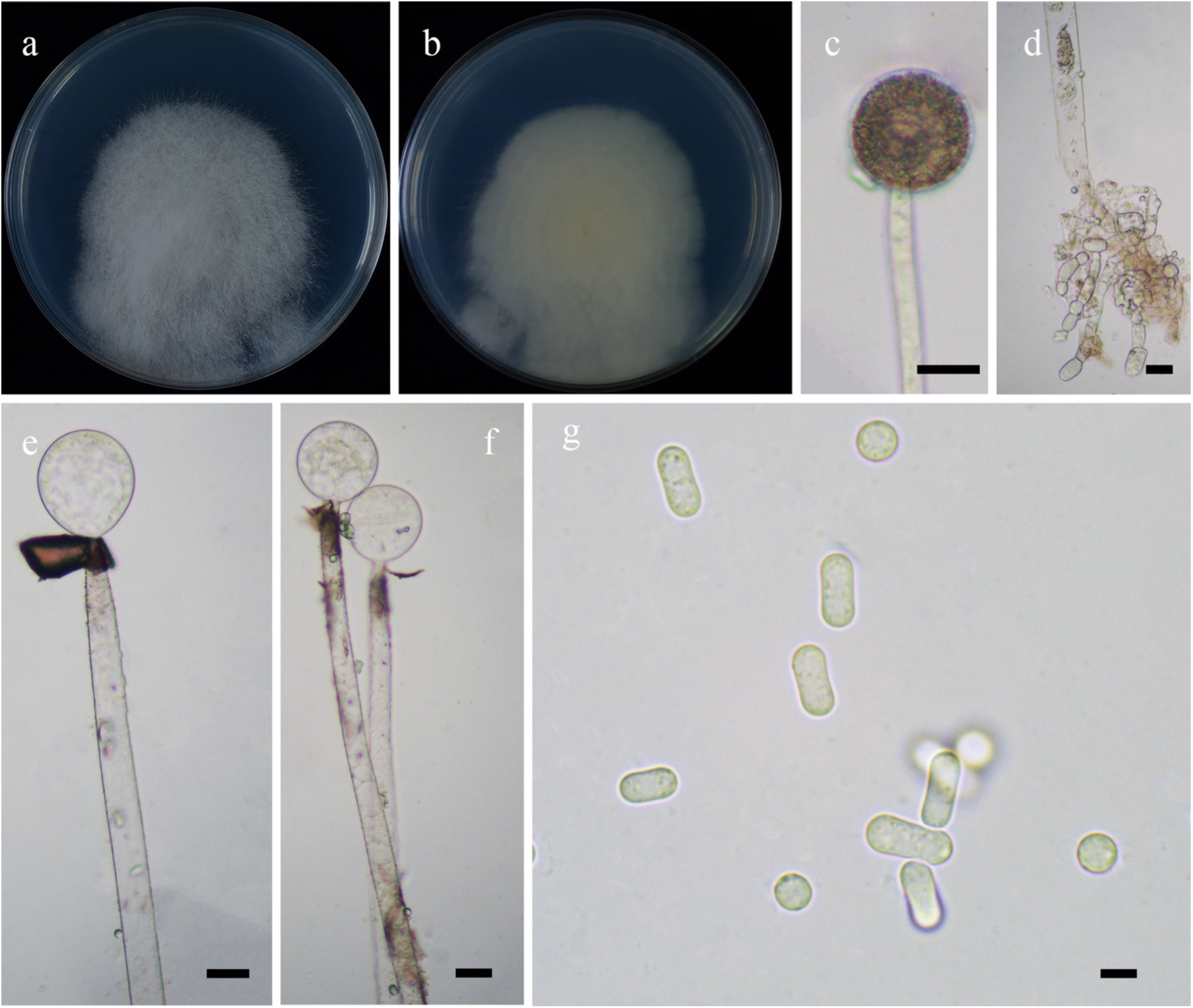
Morphologies of *Mucor amphisporus* ex-holotype CGMCC 3.16134. **a, b.** Colonies on PDA (**a.** obverse, **b.** reverse); **c.** Sporangia; **d.** Substrate hyphae with swollen; **e, f.** Columellae with collar; **g.** Sporangiospores. — Scale bars: **c–f.** 20 μm, **g.** 5 μm.

*Fungal Names*: FN570924.

*Etymology*: *amphisporus* (Lat.) refers to the species producing two types of sporangiospores.

*Holotype*: HMAS 351536.

*Colonies* on PDA at 27 °C for 9 days, slow growing, reaching 60 mm in diameter, more than 20 mm high, floccose, white, reverse Light Yellow. *Hyphae* flourishing, always unbranched, substrate hyphae abundant, usually swollen, 9.5–23.5 μm in diameter. *Rhizoids* absent. *Stolons* absent. *Sporangiophores* arising from substrate and aerial hyphae, erect or slightly bent, unbranched, generally constricted. *Sporangia* globose, Light Brown to black, smooth, 38.0–77.5 μm in diameter, walls deliquescent. *Apophyses* absent. *Collar* distinct and large. *Columellae* subglobose and globose, brownish, smooth, 17.0–58.5 μm long and 11.5–57.5 μm wide. *Sporangiospores* of two kinds: cylindrical and constricted in the middle, 5.5–9.5 μm long and 2.5–4.0 μm wide, and globose, 4.0–6.0 μm in diameter. *Chlamydospores* absent. *Zygospores* unknown.

*Materials examined*: China, Inner Mongolia Auto Region. 42°27’45″N, 113°5’15″E, from soil sample, 29 July 2021, Heng Zhao (holotype HMAS 351536, living ex-holotype culture CGMCC 3.16134); 42°59’17 ″N, 112°29’34″E, from soil sample, 29 July 2021, Heng Zhao (living culture XY07649).

*GenBank accession numbers*: OL678181 and OL678182.

*Notes*: *Mucor amphisporus* is closely related to *M. zychae* Baijal & B.S. Mehrotra based on phylogenetic analysis of ITS rDNA sequences (Fig. 14). However, morphologically *M. zychae* differs from *M. amphisporus* by larger sporangiospores (12–31.5 μm long and 5.2–15.7 μm wide vs. 5.5–9.5 μm long and 2.5–4.0 μm wide), smaller columellae (17.5–35 μm long and 14–25.5 μm wide vs. 17.0–58.5 μm long and 11.5–57.5 μm wide), producing conical columellae and various-shaped sporangiospores, including globose, broadly ellipsoid and reniform (Baijal & Mehrotra 1965).

**47. *Mucor breviphorus*** H. Zhao, Y.C. Dai & X.Y. Liu, sp. nov., Fig. 63.

**Fig. 63.**
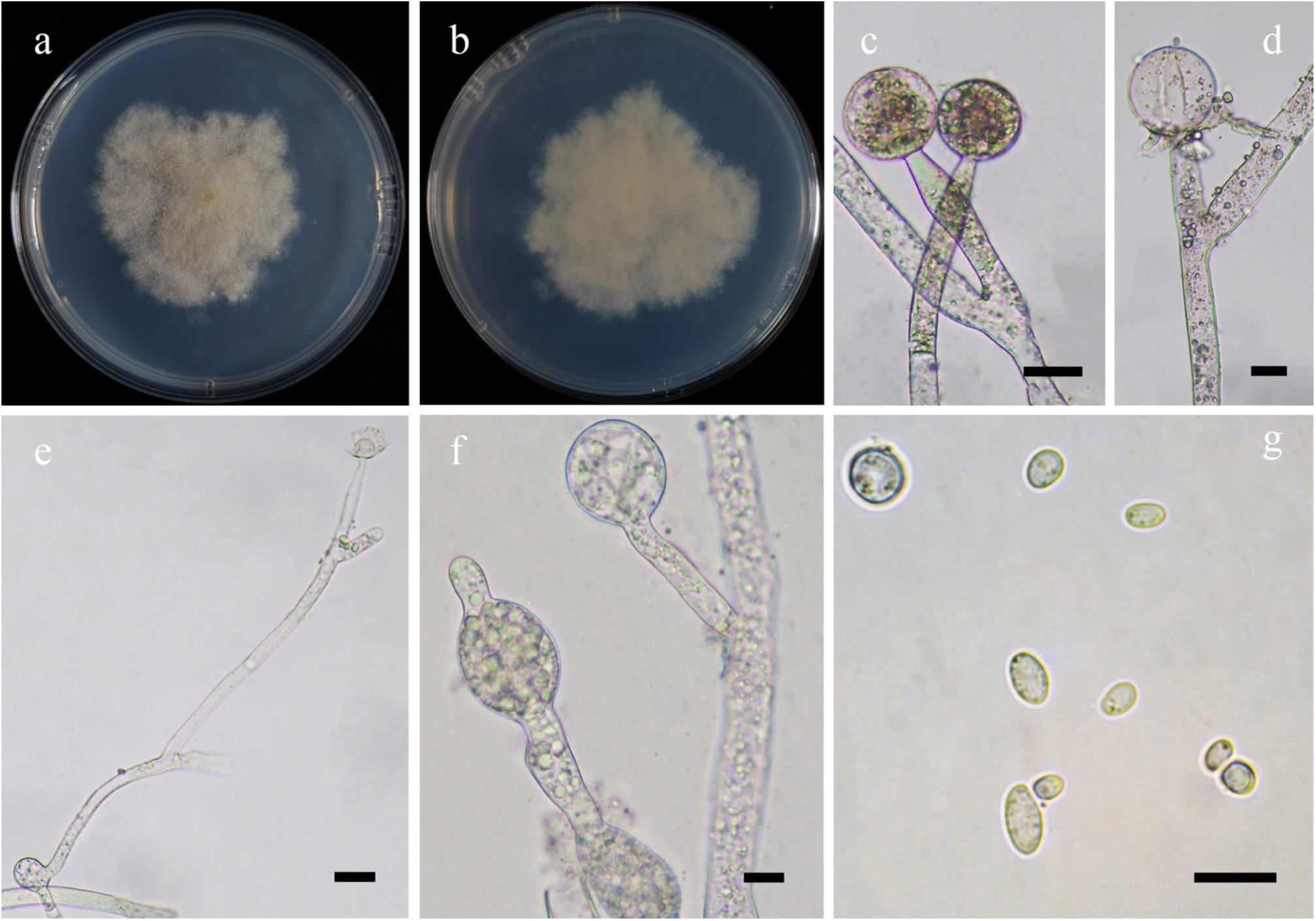
Morphologies of *Mucor breviphorus* ex-holotype CGMCC 3.16135. **a, b.** Colonies on PDA (**a.** obverse, **b.** reverse); **c.** Sporangia; **d.** Columellae with collars; **e.** Sympodial sporangiophores; **f.** Swellings in substrate hyphae; **g.** Sporangiospores. — Scale bars: **c–e.** 20 μm, **f, g.** 10 μm.

*Fungal Names*: FN570925.

*Etymology*: *breviphorus* (Lat.) refers to the species having short lateral sporangiophores.

*Holotype*: HMAS 351537.

*Colonies* on PDA at 27 °C for 12 days, slow growing, reaching 65 mm in diameter, less than 5 mm high, initially white, gradually becoming Pale Yellow-Orange with age, granulate, regular at margin. *Hyphae* flourishing, simply branched, aseptate when juvenile, septate with age, 5.5–19.5 μm in diameter. *Substrate hyphae* abundant, always swollen. *Rhizoids* absent. *Stolons* absent. *Sporangiophores* arising directly from substrate hyphae, erect or recumbent, long or short, unbranched, simply branched or sympodial, always septate below columellae, slightly constricted on the top. *Sporangia* mostly globose, occasionally pigmented, multi-spored, 17.0–52.5 μm in diameter, walls deliquescent but rough. *Apophyses* absent. *Collars* present, always distinct, rarely small. *Columellae* hemispherical or globose, hyaline, smooth, 13.0–42.5 μm long and 13.0–36.5 μm wide. *Sporangiospores* mainly ovoid or fusiform, rarely subglobose, globose or irregular, 4.0–10.5 μm long and 2.5–7.5 μm wide, or 7.0–8.5 μm in diameter. *Chlamydospores* absent. *Zygospores* unknown.

*Material examined*: China, Yunnan Province, Diqing, Deqen County, 28°27’38″N, 98°46’15″E, from soil sample, 10 October 2021, Heng Zhao (holotype HMAS 351537, living ex-holotype culture CGMCC 3.16135).

*GenBank accession number*: OL678183.

*Notes*: *Mucor breviphorus* is closely related to *M. heilongjiangensis* H. Zhao *et al*. based on phylogenetic analysis of ITS rDNA sequences (Fig. 14). However, morphologically *M. heilongjiangensis* differs from *M. breviphorus* by a faster growth rate (90 mm in diameter with 7 days vs. 65 mm in diameter with 12 days), smaller sporangia (18.0–32.0 μm in diameter vs. 17.0– 52.5 μm in diameter), ellipsoid to reniform sporangiospores, and producing chlamydospores.

**48. *Mucor brunneolus*** H. Zhao, Y.C. Dai & X.Y. Liu, sp. nov., Fig. 64.

**Fig. 64.**
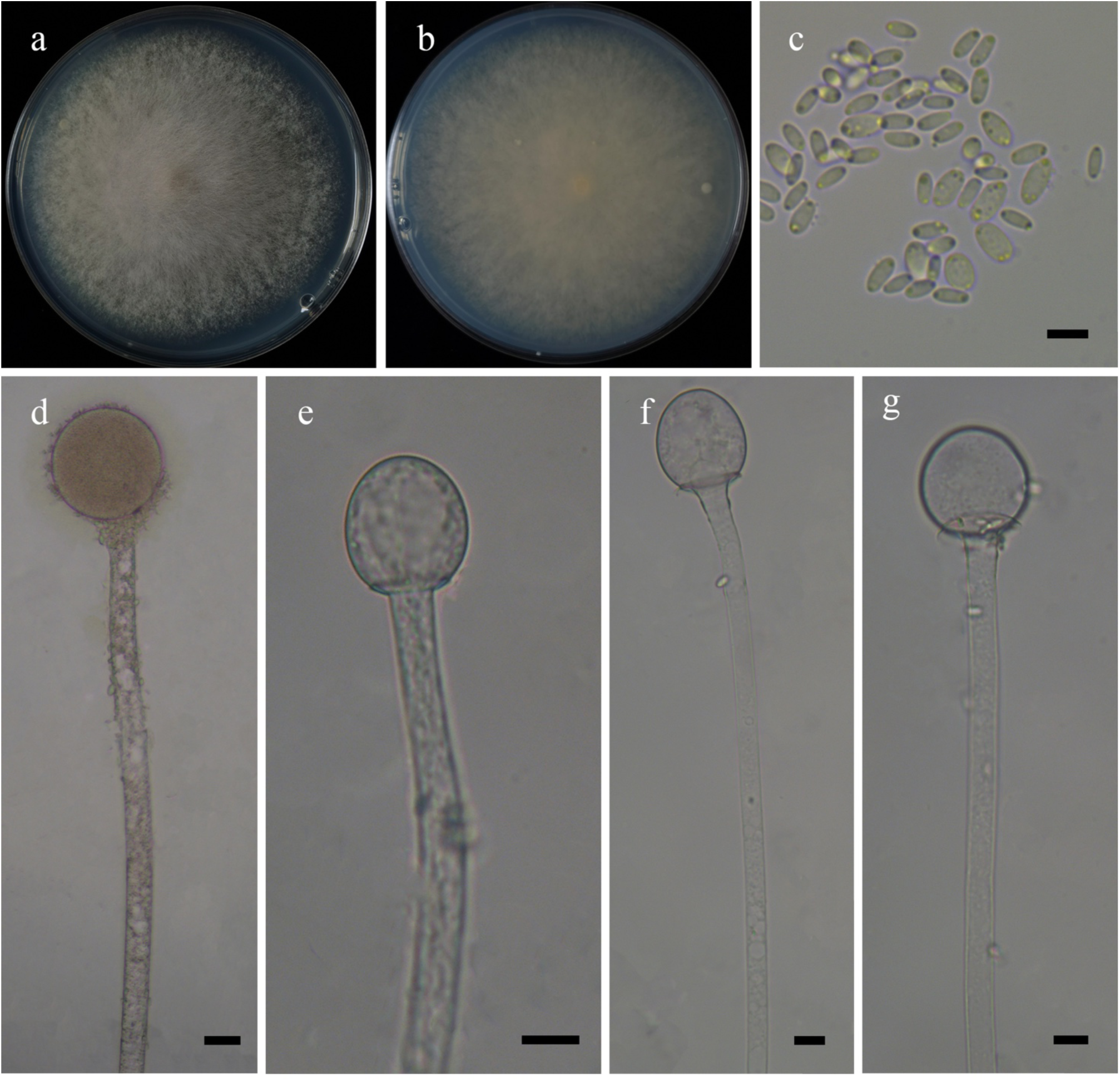
Morphologies of *Mucor brunneolus* ex-holotype CGMCC 3.16136. **a, b.** Colonies on PDA (**a.** obverse, **b.** reverse); **c.** Sporangiospores; **d.** Sporangia; **e–g.** Columellae. — Scale bars: **c.** 5 μm, **d.** 20 μm, **e–g.** 10 μm.

*Fungal Names*: FN570926.

*Etymology*: *brunneolus* (Lat.) refers to the species having pale brown sporangia.

*Holotype*: HMAS 351538.

*Colonies* on PDA at 27 °C for 4 days, fast growing, reaching 90 mm in diameter, more than 10 mm high, floccose, initially white, soon becoming Sulphur Yellow. *Hyphae* flourishing, unbranched, 4.6–16.0 μm in diameter. *Rhizoids* absent. *Stolons* absent. *Sporangiophores* arising from substrate hyphae, erect or bent, unbranched. *Sporangia* globose, hyaline when young, Pale Brown when old, smooth, 40.5–63.0 μm in diameter, walls deliquescent. *Apophyses* absent. *Collar* small, or absent. *Columellae* conical, ovoid, subglobose and globose, hyaline or subhyaline, smooth, 15.5–40.0 μm long and 14.0–40.0 μm wide. *Sporangiospores* mainly fusiform, rarely ovoid and cylindrical, hyaline, 2.5–7.0 μm long and 1.0–3.5 μm wide. *Chlamydospores* absent. *Zygospores* unknown.

*Material examined*: China, Guangxi Auto Region, Fangchenggang, 21°44’46″N, 108°5’13″E, from soil sample, 22 July 2021, Heng Zhao (holotype HMAS 351538, living ex-holotype culture CGMCC 3.16136).

*GenBank accession number*: OL678184.

*Notes*: *Mucor brunneolus* is closely related to *M. irregularis* Stchigel (replaced synonym *Rhizomucor variabilis* var. *variabilis*) based on phylogenetic analysis of ITS rDNA sequences (Fig. 14). However, morphologically *M. irregularis* differs from *M. brunneolus* by sporangiospores and columellae which are both variable and irregular in shape and size, and producing rhizoids and stolons (Zheng & Chen 1991, Alvarez *et al*. 2011).

**49. *Mucor changshaensis*** H. Zhao, Y.C. Dai & X.Y. Liu, sp. nov., Fig. 65.

**Fig. 65.**
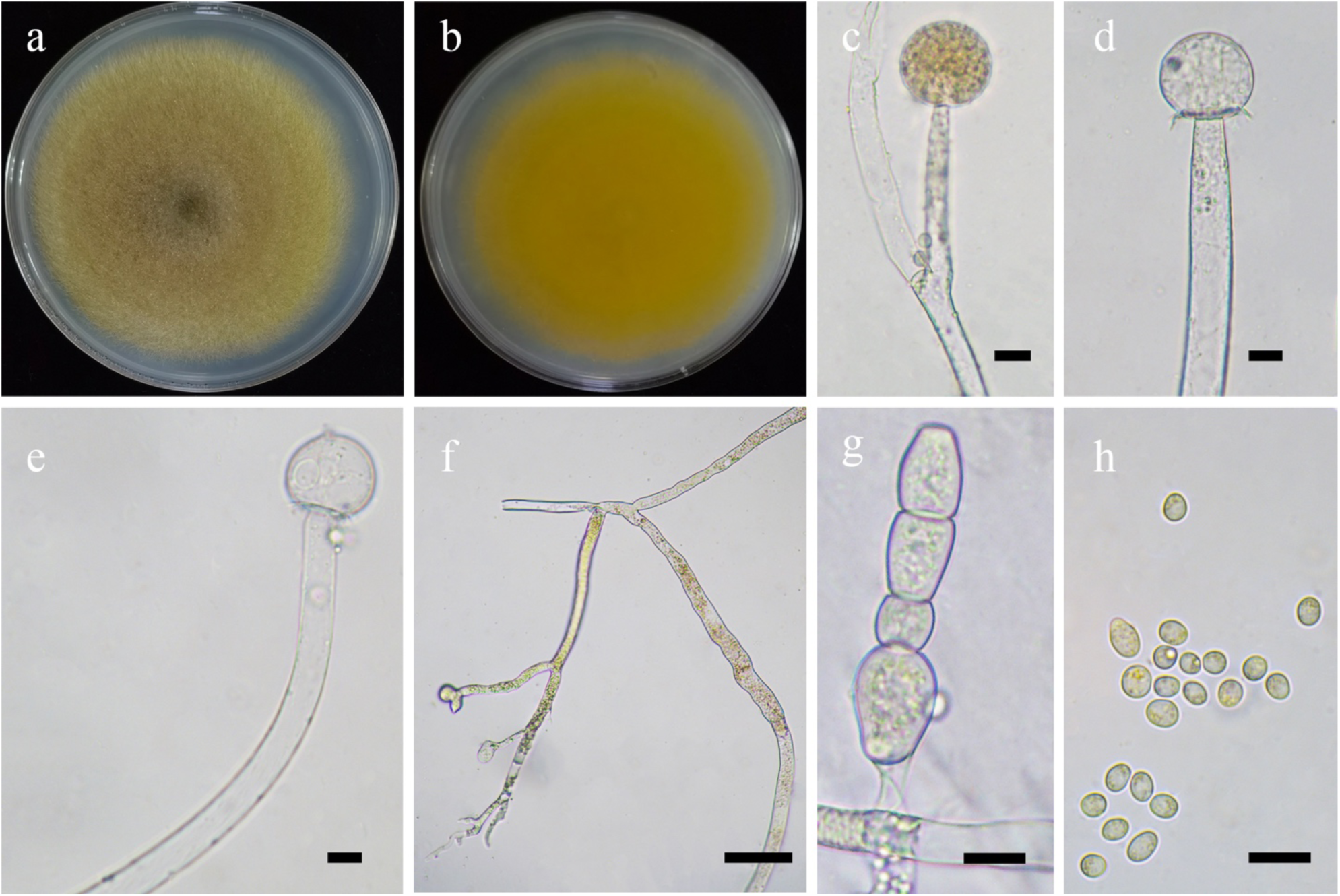
Morphologies of *Mucor changshaensis* ex-holotype CGMCC 3.16137. **a, b.** Colonies on PDA (**a.** obverse, **b.** reverse); **c.** Sporangia; **d, e.** Columellae with collars; **f.** Rhizoids; **g.** Chlamydospores; **h.** Sporangiospores. — Scale bars: **c–e, g, h.** 10 μm, **f.** 50 μm.

*Fungal Names*: FN570927.

*Etymology*: *changshaensis* (Lat.) refers to Changsha, Hunan Province, China, where the type was collected.

*Holotype*: HMAS 351539.

*Colonies* on PDA at 27 °C for 8 days, slow growing, reaching 90 mm in diameter, first light yellow, soon becoming Strontian Yellow, floccose, granulate, irregular at margin. *Hyphae* flourishing, branched, aseptate when juvenile, septate with age, 4.5–15.0 μm in diameter. *Rhizoids* present, root-like, branched. *Stolons* present. *Sporangiophores* arising from substrate or aerial hyphae, erect or bent, unbranched or repeatedly branched, sometimes slightly constricted at the top. *Sporangia* globose, Light Brown to black, smooth, 23.5–52.0 μm in diameter, walls deliquescent. *Apophyses* absent. *Collars* always present. *Columellae* subglobose to globose, hyaline, 10.0–28.5 μm long and 10.5–28.0 μm wide. *Sporangiospores* ovoid to subglobose, 4.0–7.0 μm long and 3.0– 5.0 μm wide. *Chlamydospores* abundant in substrate hyphae, in chains, variable in shape, cylindrical, ellipsoid, ovoid or irregular, 8.5–20.0 μm long and 7.0–16.5 μm wide, or globose, 8.5– 14.0 μm in diameter. *Zygospores* unknown.

*Materials examined*: China. Hunan Province, Changsha, 28°8’4″N, 112°53’10″E, from soil sample, 28 August 2021, Heng Zhao (holotype HMAS 351539, living ex-holotype culture CGMCC 3.16137, and living culture XY07965).

*GenBank accession numbers*: OL678185 and OL678186.

*Notes*: *Mucor changshaensis* is closely related to *M. atramentarius* L. Wagner & G. Walther based on phylogenetic analysis of ITS rDNA sequences (Fig. 14). However, morphologically *M. atramentarius* differs from *M. changshaensis* by larger sporangia (up to 90 μm in diameter vs. 23.5–52.0 μm in diameter), subglobose to globose sporangiospores, and the absence of chlamydospores (Wagner *et al*. 2020).

**50. *Mucor chlamydosporus*** H. Zhao, Y.C. Dai & X.Y. Liu, sp. nov., Fig. 66.

**Fig. 66.**
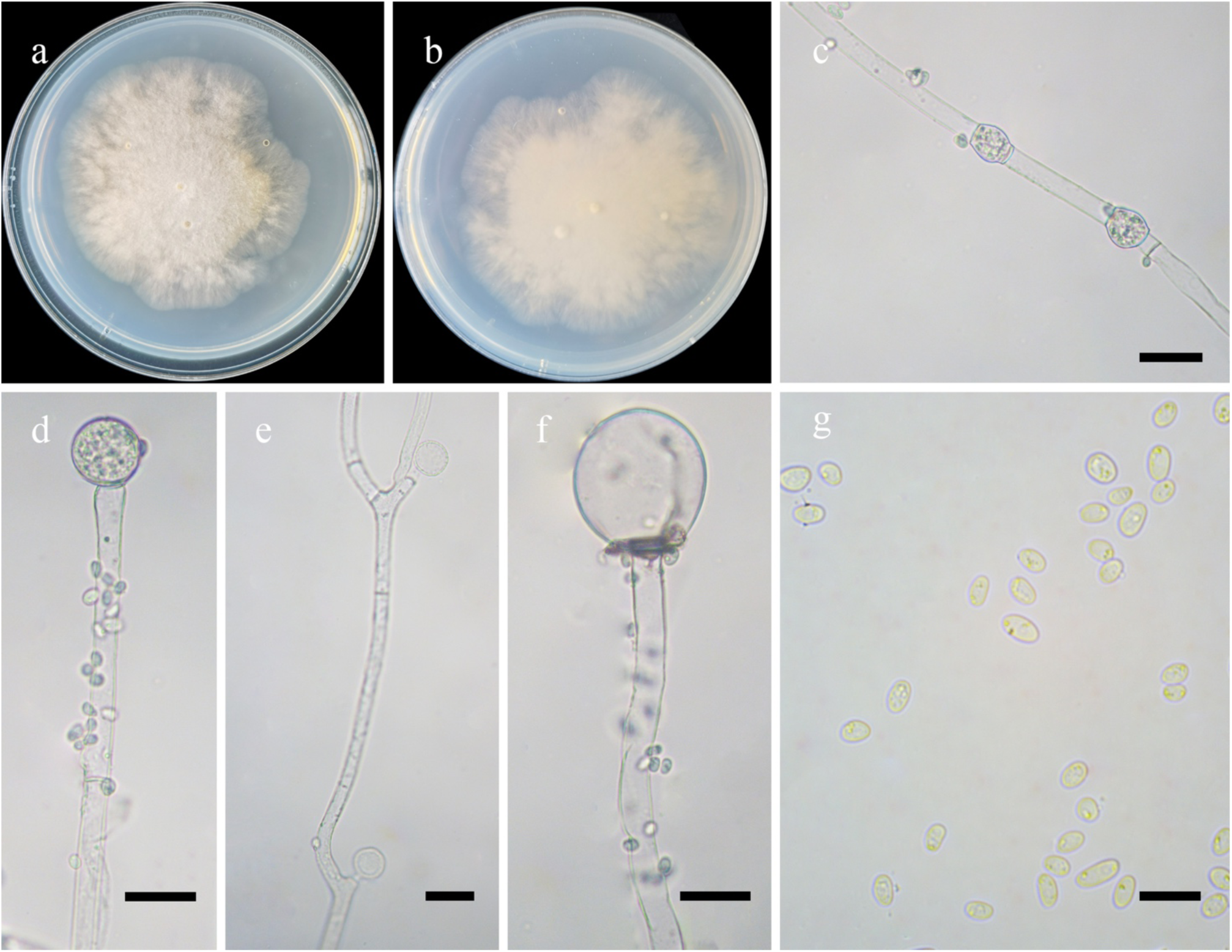
Morphologies of *Mucor chlamydosporus* ex-holotype CGMCC 3.16138. **a, b.** Colonies on PDA (**a.** obverse, **b.** reverse); **c.** Chlamydospores formed in aerial hyphae; **d.** Immature fertile sporangia; **e.** Aborted sporangia; **f.** Columellae; **g.** Sporangiospores. — Scale bars: **c–f.** 20 μm, **g.** 10 μm.

*Fungal Names*: FN570928.

*Etymology*: *chlamydosporus* (Lat.) refers to the species having chlamydospores.

*Holotype*: HMAS 351540.

*Colonies* on PDA at 27 °C for 7 days, slow growing, reaching 70 mm in diameter, floccose, initially white, becoming Pale Yellow-Orange when mature. *Hyphae* flourishing, branched, aseptate when juvenile, septate with age, 3.5–13.5 μm in diameter. *Rhizoids* absent. *Stolons* absent. *Sporangiophores* arising from substrate hyphae, erect, simple and unbranched, or sympodial, septate, slightly constricted. *Fertile sporangia* always terminal on sporangiophores, subglobose or globose, walls deliquescent. *Aborted sporangia* laterally formed on aerial hyphae, globose, hyaline, smooth, 10.0–53.0 μm in diameter. *Apophyses* absent. *Collars* rarely present. *Columellae* mostly globose, occasionally subglobose, smooth, 7.5–24.0 μm in diameter. *Projections* absent. *Sporangiospores* ellipsoid or ovoid, 4.0–7.5 μm long and 2.5–3.5 μm wide. *Chlamydospores* irregular, 10.5–16.0 μm long and 8.5–13.5 μm wide. *Zygospores* absent.

*Materials examined*: China. Hebei Province, Shijiazhuang, 38°43’16″N, 113°50’39″E, from soil sample, 14 Jan 2021, Tong-Kai Zong (holotype HMAS 351540, living ex-holotype culture CGMCC 3.16138). Yunnan Province, Chuxiong, 25°18’53″N, 101°25’17″E, from soil sample, 14 October 2021, Heng Zhao (living culture XY08211 and XY08225).

*GenBank accession numbers*: OL678187, OL678188 and OL678189.

*Notes*: *Mucor chlamydosporus* is closely related to *M. fluvii* Hyang B. Lee *et al*. based on phylogenetic analysis of ITS rDNA sequences (Fig. 14). However, morphologically *M. fluvii* differs from *M. chlamydosporus* by smaller sporangia (8.0–45 μm in diameter vs. 10.0–53.0 μm), bigger columellae (9.5–31.5 μm in diameter vs. 7.5–24.0 μm in diameter; Wanasinghe *et al*. 2018).

**51. *Mucor donglingensis*** H. Zhao, Y.C. Dai & X.Y. Liu, sp. nov., Fig. 67.

**Fig. 67.**
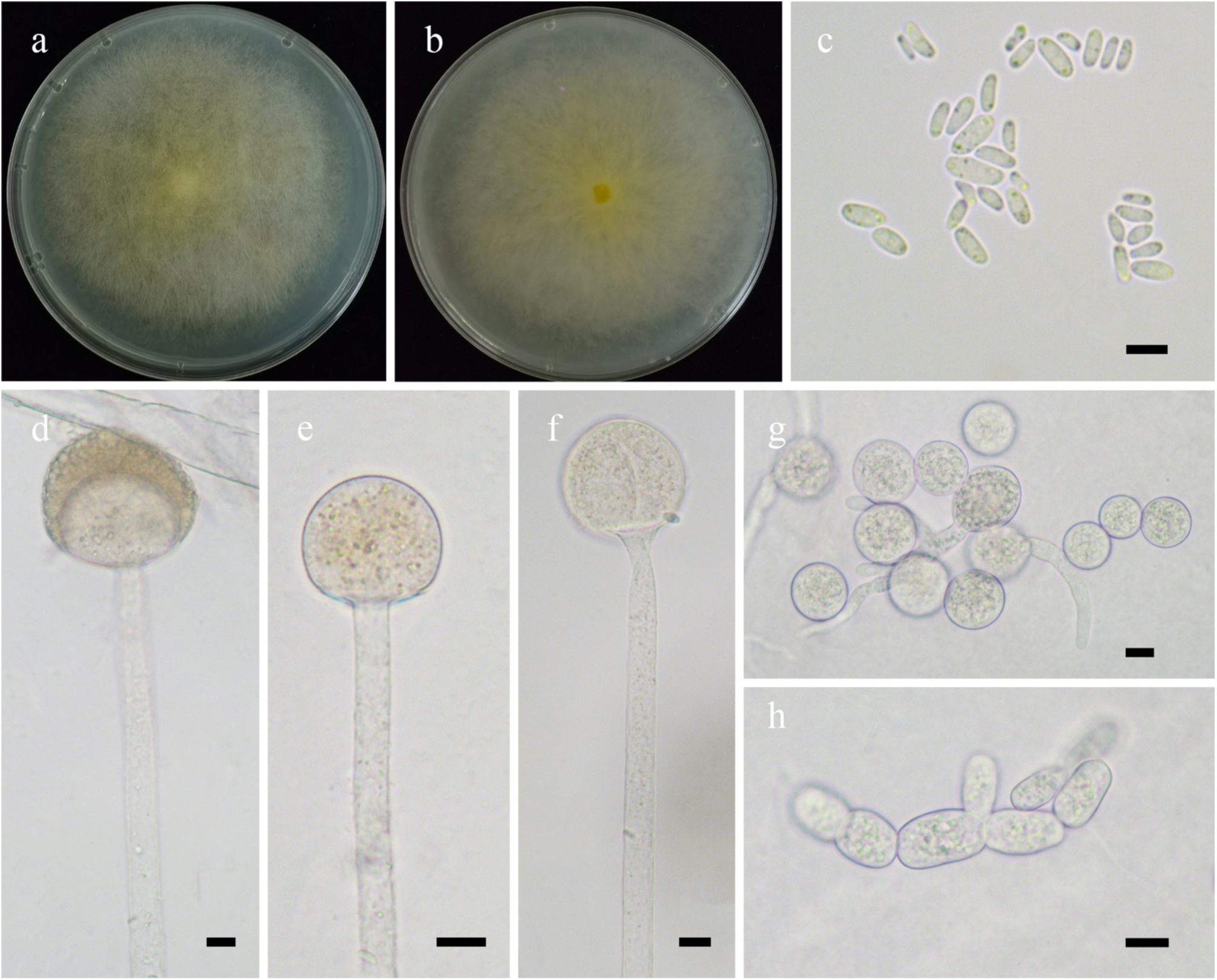
Morphologies of *Mucor donglingensis* ex-holotype CGMCC 3.16139. **a, b.** Colonies on PDA (**a.** obverse, **b.** reverse); **c.** Sporangiospores; **d.** Sporangia; **e, f.** Columellae; **g, h.** Chlamydospores. — Scale bars: **c.** 5 μm, **d–h.** 10 μm.

*Fungal Names*: FN570929.

*Etymology*: *donglingensis* (Lat.) refers to Dongling Mountain, Beijing, China, where the type was collected.

*Holotype*: HMAS 351541.

*Colonies* on PDA at 27 °C for 5 days, fast growing, reaching 90 mm in diameter, floccose, first white, soon becoming Naphthalene Yellow to Citron Yellow. *Hyphae* flourishing, branched, 6.5–16.5 μm in diameter. *Rhizoids* absent. *Stolons* absent. *Sporangiophores* arising from substrate hyphae, erect, unbranched, slightly constricted at the top. *Sporangia* globose, hyaline when young, brown when old, smooth, 39.5–55.0 μm in diameter, walls deliquescent. *Apophyses* absent. *Collars* absent. *Columellae* globose to subglobose, hyaline or subhyaline, 19.5–49.5 μm long and 20.0–44.5 μm wide. *Sporangiospores* oblong to ovoid, hyaline or subhyaline, 2.5–8.0 μm long and 1.0–4.5 μm wide. *Chlamydospores* abundantly in substrate hyphae, in chains, ellipsoid, oblong, ovoid, subglobose, globose or irregular, 11.0–33.5 μm long and 6.5–15.5 μm wide, or 15.0–24.5 μm in diameter. *Zygospores* unknown.

*Materials examined*: China. Beijing, Dongling Mountain, 115°25’48′′E, 39°22’12′′N, from plant debris sample, 28 August 2021, Heng Zhao (holotype HMAS 351541, living ex-holotype culture CGMCC 3.16139). Yunnan Province, Diqing, Deqen Country, 98°46’15′′E, 28°27’38′′N, from soil sample, 9 October 2021, Heng Zhao (living culture XY08501).

*GenBank accession numbers*: OL678190 and OL678191.

*Notes*: *Mucor chlamydosporus* is closely related to *M. fluvii* based on phylogenetic analysis of ITS rDNA sequences (Fig. 14). However, morphologically *M. fluvii* differs from *M. donglingensis* by simple or sympodial sporangiophores, and one or two septa below the sporangia (Crous *et al*. 2018).

**52. *Mucor floccosus*** H. Zhao, Y.C. Dai & X.Y. Liu, sp. nov., Fig. 68.

**Fig. 68.**
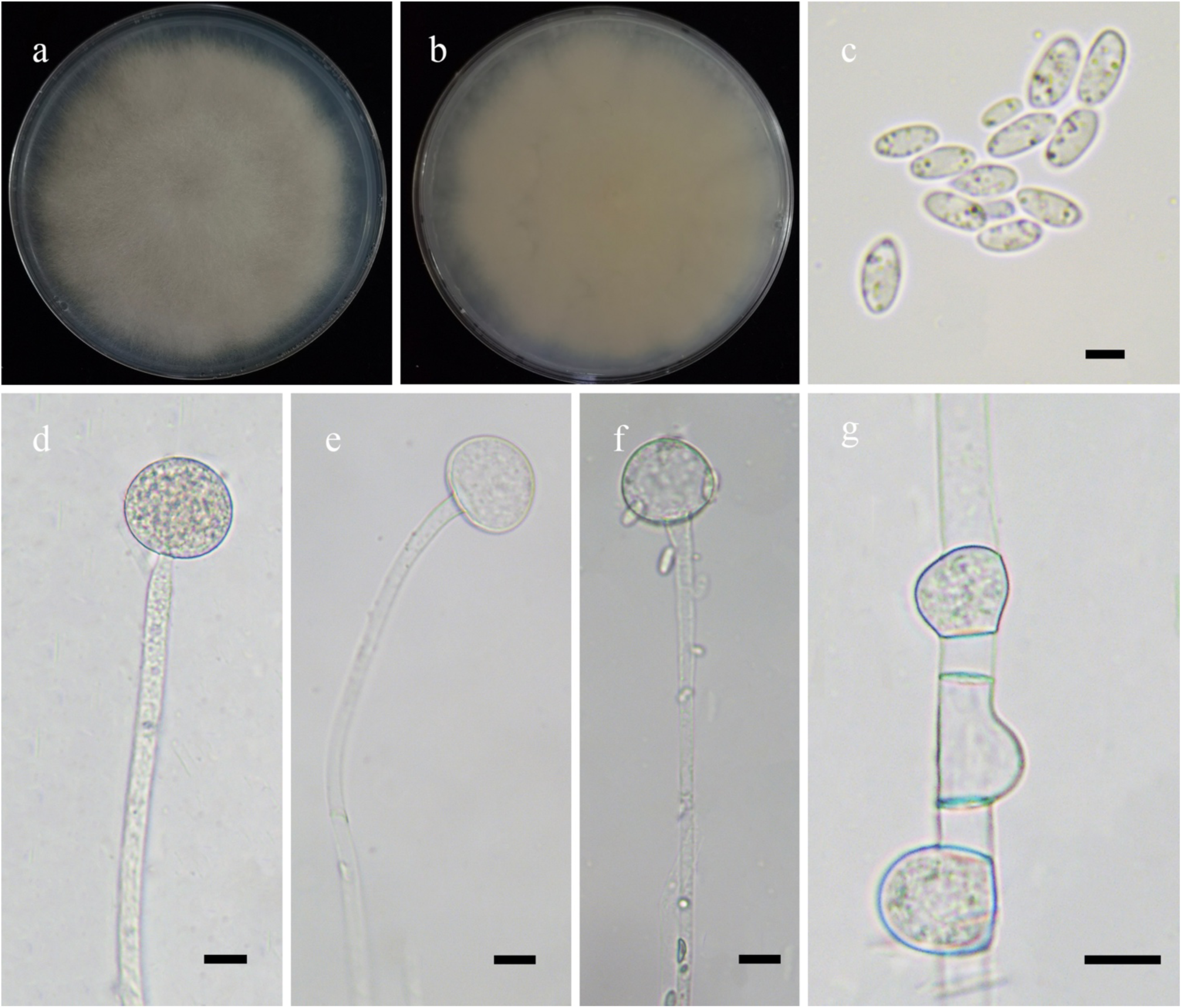
Morphologies of *Mucor floccosus* ex-holotype CGMCC 3.16140. **a, b.** Colonies on PDA (**a.** obverse, **b.** reverse); **c.** Sporangiospores; **d.** Sporangia; **e, f.** Columellae without collar; **g.** Chlamydospores. — Scale bars: **c.** 5 μm, **d–g.** 10 μm.

*Fungal Names*: FN570930.

*Etymology*: *floccosus* (Lat.) refers to the species having floccose colonies.

*Holotype*: HMAS 351542.

*Colonies* on PDA at 27 °C for 7 days, reaching 90 mm in diameter, broadly lobed and concentrically zonate, first white, gradually becoming Light Gray, floccose. *Hyphae* flourishing, branched, aseptate when juvenile, septate with age, 3.0–13.0 μm in diameter. *Rhizoids* absent*. Stolons* absent. *Sporangiophores* arising from aerial hyphae, erect, straight or bent, unbranched, slightly expanded at the top. *Sporangia* globose, hyaline when young, brown when old, smooth, 19.5–43.5 μm in diameter, walls deliquescent. *Apophyses* absent. *Collars* absent. *Columellae* hemispherical or depressed globose, hyaline, smooth, 9.0–23.5 μm long and 11.0–26.0 μm wide. *Sporangiospores* fusiform or ellipsoid, 4.0–7.5 μm long and 1.0–4.0 μm wide. *Chlamydospores* in aerial hyphae, hemispherical, subglobose, globose, or irregular, 9.5–19.0 μm long and 5.5–15.0 μm wide. *Zygospores* unknown.

*Materials examined*: China, Hunan Province, Changsha, 28°8’4″N, 112°53’10″E, from soil sample, 28 August 2021, Heng Zhao (holotype HMAS 351542, living ex-holotype culture CGMCC 3.16140, and living culture XY07967).

*GenBank accession numbers*: OL678192 and OL678193.

*Notes*: *Mucor floccosus* is closely related to *M. rhizosporus* H. Zhao *et al*. and *M. pernambucoensis* C.L. Lima *et al*. based on phylogenetic analysis of ITS rDNA sequences (Fig. 14). However, morphologically *M. rhizosporus* differs from *M. floccosus* by abundant chlamydospores in rhizoids, Pale Yellow-Orange colonies, and ovoid to oblong columellae. *M. pernambucoensi* differs from *M. floccosus* by sympodially-branched sporangiophores (de Lima *et al*. 2018).

**53. *Mucor fusiformisporus*** H. Zhao, Y.C. Dai & X.Y. Liu, sp. nov., Fig. 69.

**Fig. 69.**
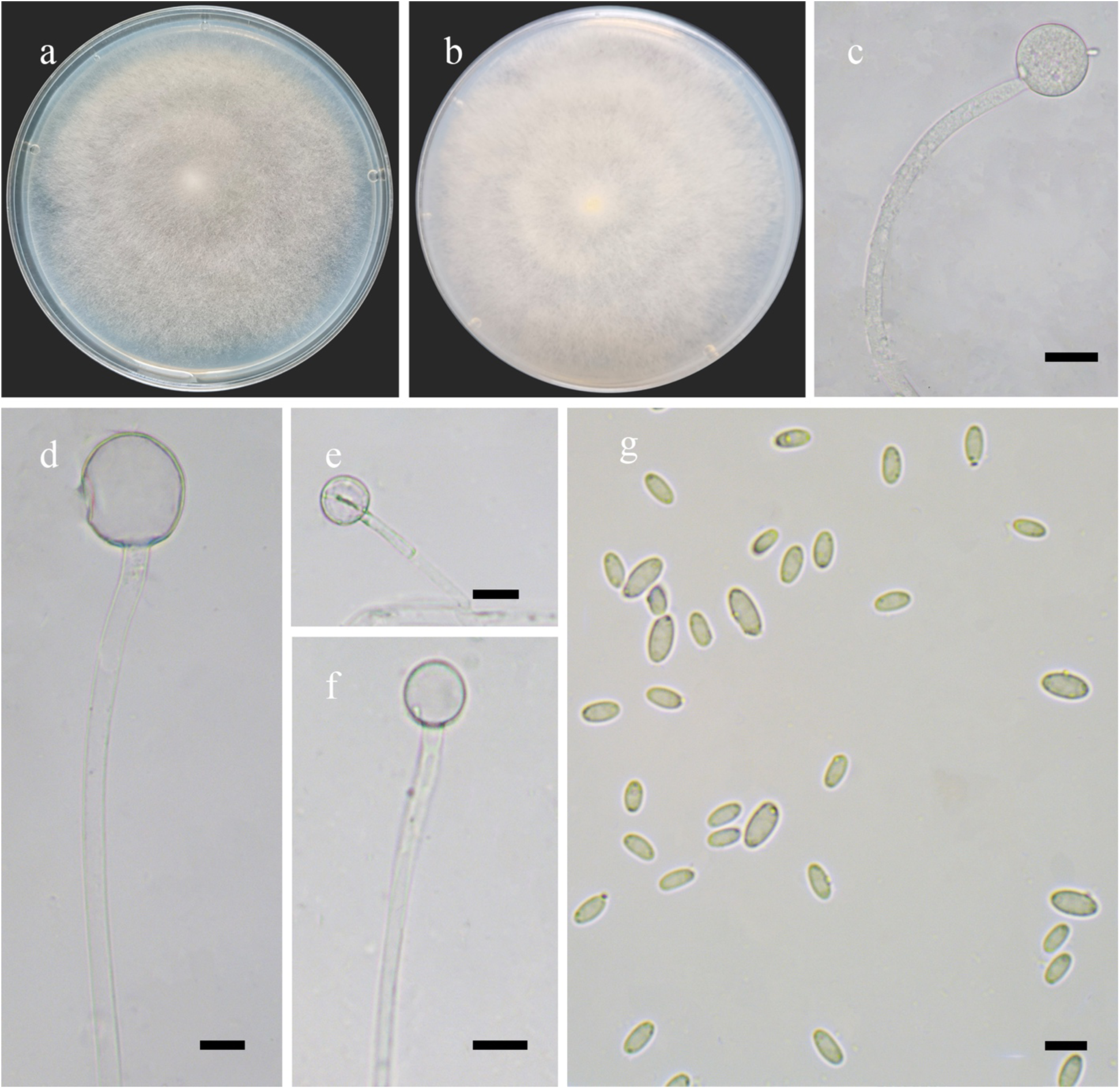
Morphologies of *Mucor fusiformisporus* ex-holotype CGMCC 3.16141. **a, b.** Colonies on PDA (**a.** reverse, **b.** obverse); **c.** Sporangia; **d–f.** Sporangiospores with columellae; **g.** Sporangiospores. — Scale bars: **c.** 20 μm, **d–f.** 10 μm, **g.** 5 μm.

*Fungal Names*: FN570931.

*Etymology*: *fusiformisporus* (Lat.) refers to the species having fusiform sporangiospores.

*Holotype*: HMAS 351543.

*Colonies* on PDA at 27 °C for 4 days, fast growing, reaching 90 mm in diameter, 10 mm high, floccose, granulate, initially white, soon becoming Marguerite Yellow, reverse regular at margin. *Hyphae* flourishing, branched, aseptate when juvenile, septate with age, 4.0–21.5 μm in diameter. *Rhizoids* absent*. Stolons* absent. *Sporangiophores* arising from aerial or substrate hyphae, erect or bent, unbranched. *Sporangia* globose, hyaline when young, Light Brown to Dark Brown when old, smooth, multi-spored, 15.0–40.5 μm in diameter, walls deliquescent. *Apophyses* absent. *Collars* usually absent and occasionally small. *Columellae* subglobose or globose, hyaline or subhyaline, smooth, 8.0–37.5 μm long and 8.5–32.0 μm wide. *Sporangiospores* fusiform, hyaline or subhyaline, smooth, 3.0–7.0 μm long and 1.5–3.5 μm wide. *Chlamydospores* absent. *Zygospores* unknown.

*Materials examined*: China. Zhejiang Province, Hangzhou, 30°07’15″N, 119°58’46″E, from soil sample, 4 June 2021, Jia-Jia Chen (holotype HMAS 351543, living ex-holotype culture CGMCC 3.16141). Yunnan Province, Dali, from soil sample, 28 October 2021, Heng Zhao (living culture XY08153 and XY08154). Zhejiang Province, Hangzhou, from soil sample, 10 October 2021, Heng Zhao (living culture XY08117).

*GenBank accession numbers*: OL678194, OL678195, OL678196 and OL678197.

*Notes*: *Mucor fusiformisporus* is closely related to *M. donglingensis* H. Zhao *et al*. and *M. souzae* C.L. Lima *et al*. based on phylogenetic analysis of ITS rDNA sequences (Fig. 14). However, morphologically *M. donglingensis* differs from *M. fusiformisporus* by chlamydospores that are variable in shape. *M. souzae* differs from *M. fusiformisporus* by sporangiospores that are variable in shape, including ellipsoid, ellipsoid to fusoid, reniform and irregular, and by simple or sympodial sporangiophores (Crous *et al*. 2018).

**54. *Mucor heilongjiangensis*** H. Zhao, Y.C. Dai & X.Y. Liu, sp. nov., Fig. 70.

**Fig. 70.**
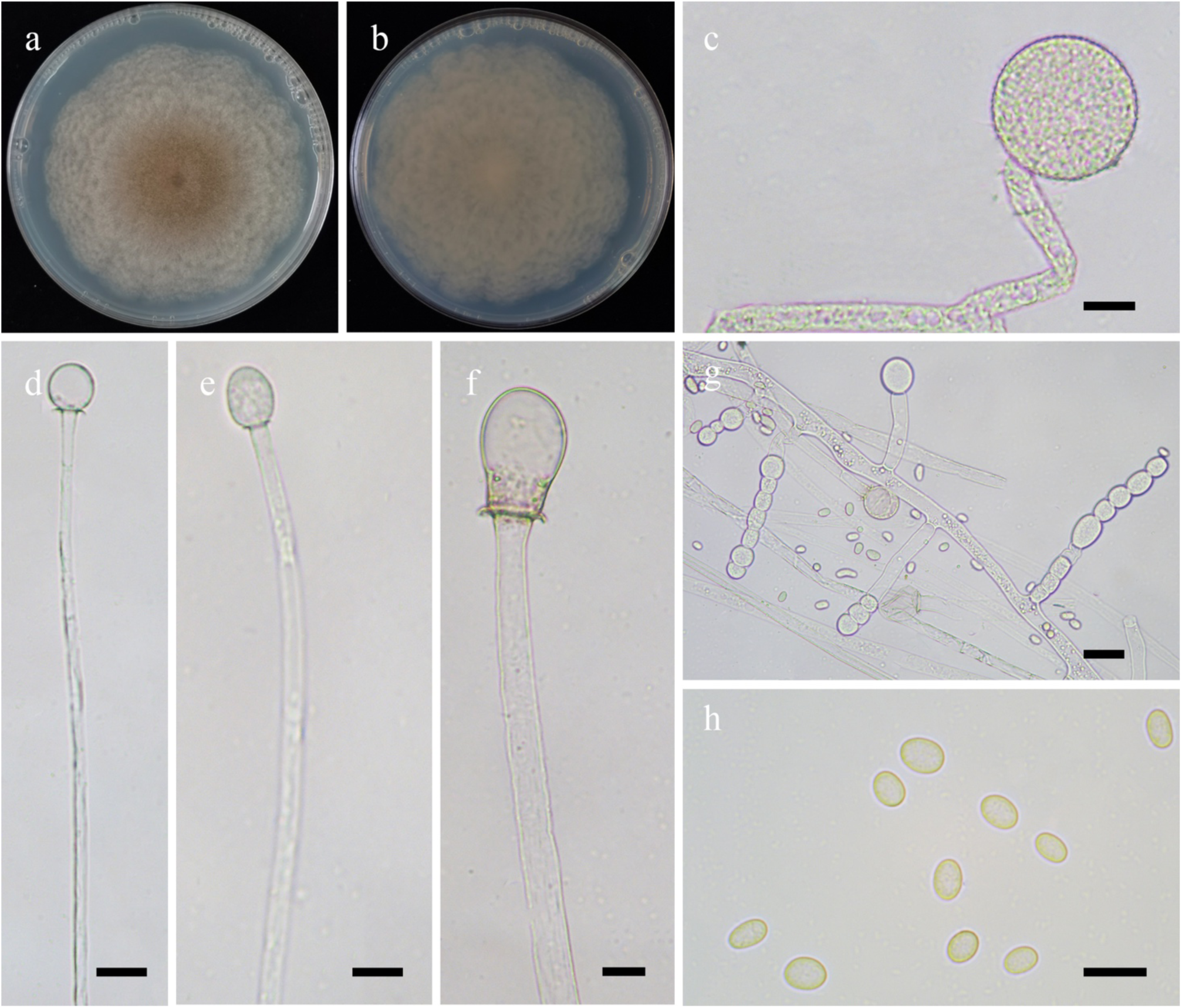
Morphologies of *Mucor heilongjiangensis* ex-holotype CGMCC 3.16142. **a, b.** Colonies on PDA (**a.** obverse, **b.** reverse); **c.** Sporangia; **d–f.** Columellae; **g.** Chlamydospores; **h.** Sporangiospores. — Scale bars: **c–f, h.** 10 μm, **g.** 20 μm.

*Fungal Names*: FN570932.

*Etymology*: *heilongjiangensis* (Lat.) refers to Heilongjiang, a province in China, where the type was collected.

*Holotype*: HMAS 351544.

*Colonies* on PDA at 27 °C for 7 days, reaching 90 mm in diameter, more than 10 mm high, broadly lobed and concentrically zonate, floccose, initially white, gradually becoming Pinkish Cinnamon to Verona Brown. *Hyphae* flourishing, always unbranched when young, branched when old, 2.0–15.5 μm in diameter. *Rhizoids* absent*. Stolons* absent. *Sporangiophores* arising from substrate and aerial hyphae, erect, unbranched, sometimes slightly expanded at the top. *Sporangia* globose, hyaline when young, Light Brown to brown when old, smooth, 18.0–32.0 μm in diameter, walls deliquescent. *Apophyses* absent. *Collars* usually absent, occasionally present but small. *Columellae* subglobose, globose, ellipsoid, or pyriform, hyaline or subhyaline, sometimes pigmented, smooth, 6.0–29.5 μm long and 5.0–21.5 μm wide. *Sporangiospores* usually ellipsoid, occasionally reniform, 3.5–10.0 μm long and 2.5–6.0 μm wide. *Chlamydospores* abundant in substrate hyphae, in chains, ellipsoid, ovoid, subglobose, globose or irregular, 5.0–18.0 μm long and 5.5–15.5 μm wide. *Zygospores* unknown.

*Materials examined*: China, Heilongjiang Province, Heihe, 48°4’37″N, 127°6’54″E, from soil sample, 18 April 2018, Ze Liu (holotype HMAS 351544, living ex-holotype culture CGMCC 3.16142, and living culture XY05057).

*GenBank accession numbers*: OL678198 and OL678199.

*Notes*: See notes of *Mucor breviphorus*.

**55. *Mucor hemisphaericum*** H. Zhao, Y.C. Dai & X.Y. Liu, sp. nov., Fig. 71.

**Fig. 71.**
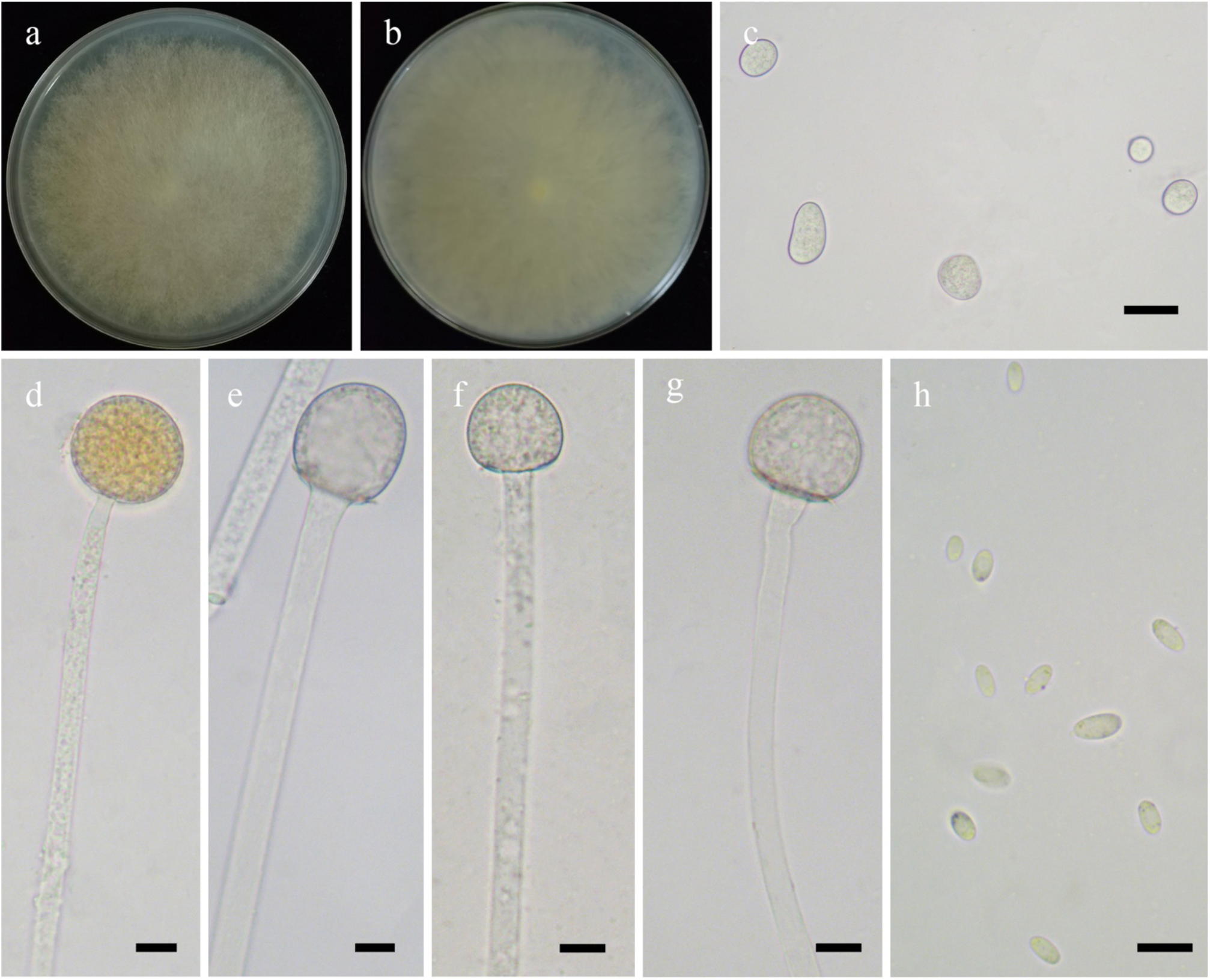
Morphologies of *Mucor hemisphaericum* ex-holotype CGMCC 3.16143. **a, b.** Colonies on PDA (**a.** obverse, **b.** reverse); **c.** Chlamydospores; **d.** Sporangia; **e–g.** Columellae with small collars; **h.** Sporangiospores. — Scale bars: **c.** 20 μm, **d–h.** 10 μm.

*Fungal Names*: FN570933.

*Etymology*: *hemisphaericum* (Lat.) refers to the species having hemispherical columellae.

*Holotype*: HMAS 351545.

*Colonies* on PDA at 27 °C for 4 days, fast growing, reaching 90 mm in diameter, floccose, irregular at margin, first white, soon becoming Ivory Yellow. *Hyphae* flourishing, always unbranched and aseptate when young, branched and septate when old, 1.5–23.5 μm in diameter. *Rhizoids* absent. *Stolons* absent. *Sporangiophores* arising from substrate and aerial hyphae, erect or slightly bent, unbranched, occasionally slightly expanded at the top. *Sporangia* globose, hyaline when young, Light Brown to brown when old, smooth, 17.5–42.5 μm in diameter, walls deliquescent. *Apophyses* absent. *Collars* usually absent, small if present. *Columellae* hemispherical or depressed globose, hyaline or subhyaline, 5.5–20.5 μm long and 6.0–20.0 μm wide. *Sporangiospores* ovoid, 3.5–7.5 μm long and 1.5–3.5 μm wide. *Chlamydospores* abundant in substrate hyphae, in chains, variable in shape, cylindrical, ellipsoid, oblong, ovoid, subglobose, globose or irregular, 9.0–23.5 μm long and 7.0–16.5 μm wide, or 9.0–16.5 μm in diameter. *Zygospores* unknown.

*Material examined*: China, Tibet Auto Region, Rikaze, Yadong County, from soil sample, 22 August 2021, Heng Zhao (holotype HMAS 351545, living ex-holotype culture CGMCC 3.16143).

*GenBank accession number*: OL678200.

*Notes*: *Mucor hemisphaericum* is closely related to *M. merdicola* C.A.F. de Souza & A.L. Santiago based on phylogenetic analysis of ITS rDNA sequences (Fig. 14). However, morphologically *M. merdicola* differs from *M. hemisphaericum* by the wider columellae (30.0–35.0 μm wide vs. 6.0–20.0 μm wide) and sympodially branched sporangiophores (Li *et al*. 2016).

**56. *Mucor homothallicus*** H. Zhao, Y.C. Dai & X.Y. Liu, sp. nov., Fig. 72.

**Fig. 72.**
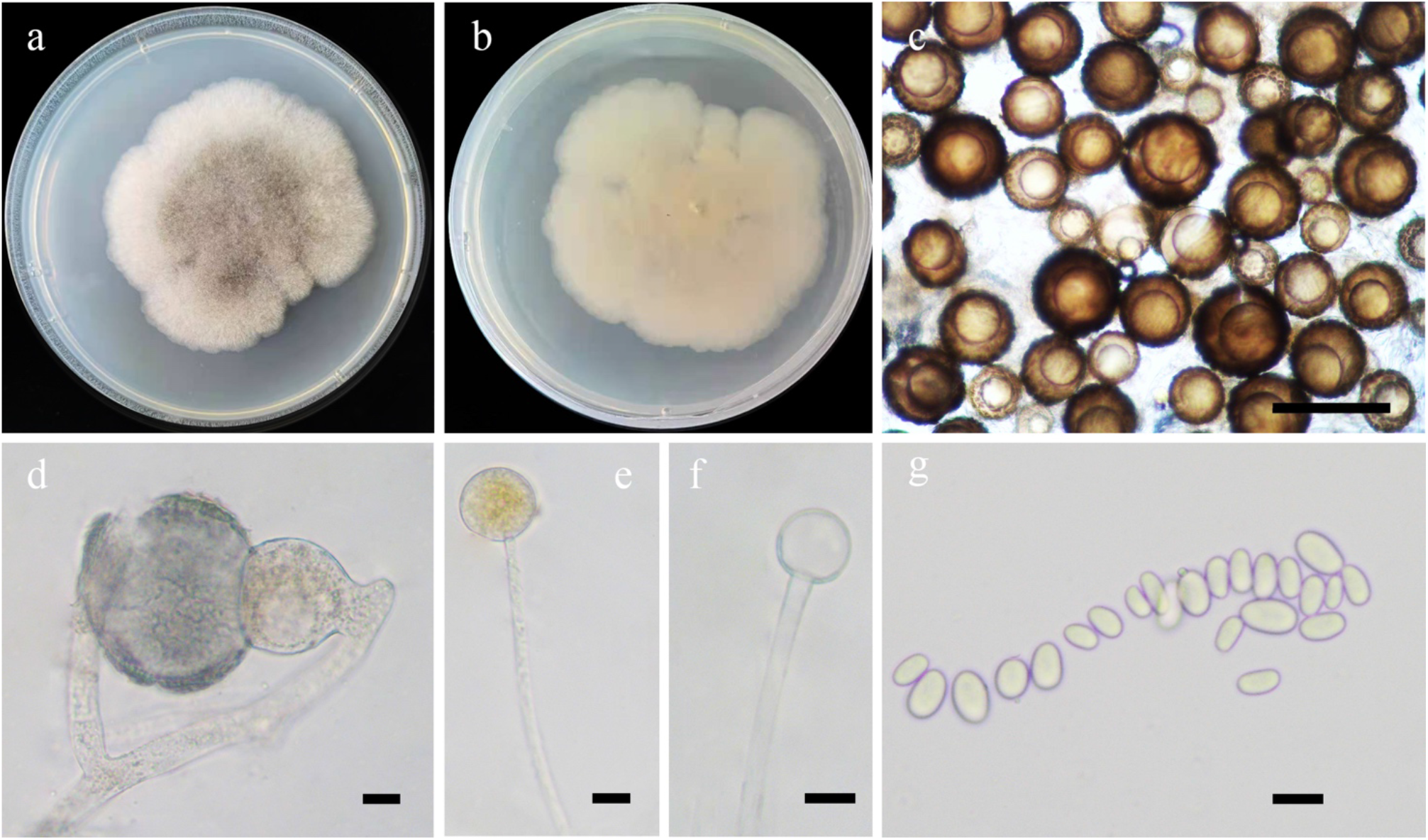
Morphologies of *Mucor homothallicus* ex-holotype CGMCC 3.16144. **a, b.** Colonies on PDA (**a.** obverse, **b.** reverse); **c, d.** Zygospores; **e.** Sporangia; **f.** Columellae; **g.** Sporangiospores. — Scale bars: **c.** 100 μm, **d–g.** 10 μm.

*Fungal Names*: FN570934.

*Etymology*: *homothallicus* (Lat.) refers to the species producing zygospores by homothallism.

*Holotype*: HMAS 351546.

*Colonies* on PDA at 25 °C for 9 days, slow growing, reaching 55 mm in diameter, 10 mm high, floccose, granulate, lobed at edge, initially white, then becoming Light Black, reverse irregular at margin. *Hyphae* flourishing, branched, aseptate when juvenile, septate with age, 5.0–17.0 μm in diameter. *Rhizoids* absent. *Stolons* absent. *Sporangiophores* arising from aerial hyphae, erect or slightly bent, unbranched. *Sporangia* globose, smooth, Light Brown or Dark Brown, 10.0–22.5 μm in diameter, walls deliquescent. *Apophyses* absent. *Collars* absent. *Columellae* oblong, subglobose and globose, hyaline or subhyaline, smooth, 6.0–13.5 μm long and 6.5–14.5 μm wide. *Sporangiospores* ellipsoid or subglobose, 7.0–11.0 μm long and 3.0–7.0 μm wide. *Chlamydospores* absent. *Zygospores* homothallic, mainly globose, hyaline when juvenile, gradually Dark Brown with age, surrounded by liquid droplets, finally Light Brown, hyaline or subhyaline, 39.0–80.0 μm in diameter.

*Materials examined*: China, Jiangsu Province, Lianyungang, 34°38’9″N, 119°29’58″E, from soil sample, 28 May 2021, Heng Zhao (holotype HMAS 351546, living ex-holotype culture CGMCC 3.16144, and living culture XY06967).

*GenBank accession numbers*: OL678201 and OL678202.

*Notes*: *Mucor homothallicus* is closely related to *M. endophyticus* (R.Y. Zheng & H. Jiang) J. Pawłowska & Walthe based on phylogenetic analysis of ITS rDNA sequences (Fig. 14). However, morphologically *M. endophyticus* differs from *M. homothallicus* by abundant rhizoids, collars, subglobose, ovoid, limoniform, or irregular chlamydospores, larger sporangia (38.0–80.0 μm in diameter vs. 10.0–22.5 μm in diameter), larger columellae (19.0–50.5 μm long and 18.0–46.0 μm or 13.5–37.0 μm in diameter vs. 6.0–13.5 μm long and 6.5–14.5 μm wide), and sporangiospores variable in shape, including narrowly to broadly ellipsoid, ovoid, subglobose, triangular, allantoid, and irregular (Zheng & Jiang 1995, Hoffmann *et al*. 2013).

**57. *Mucor hyalinosporus*** H. Zhao, Y.C. Dai & X.Y. Liu, sp. nov., Fig. 73.

**Fig. 73.**
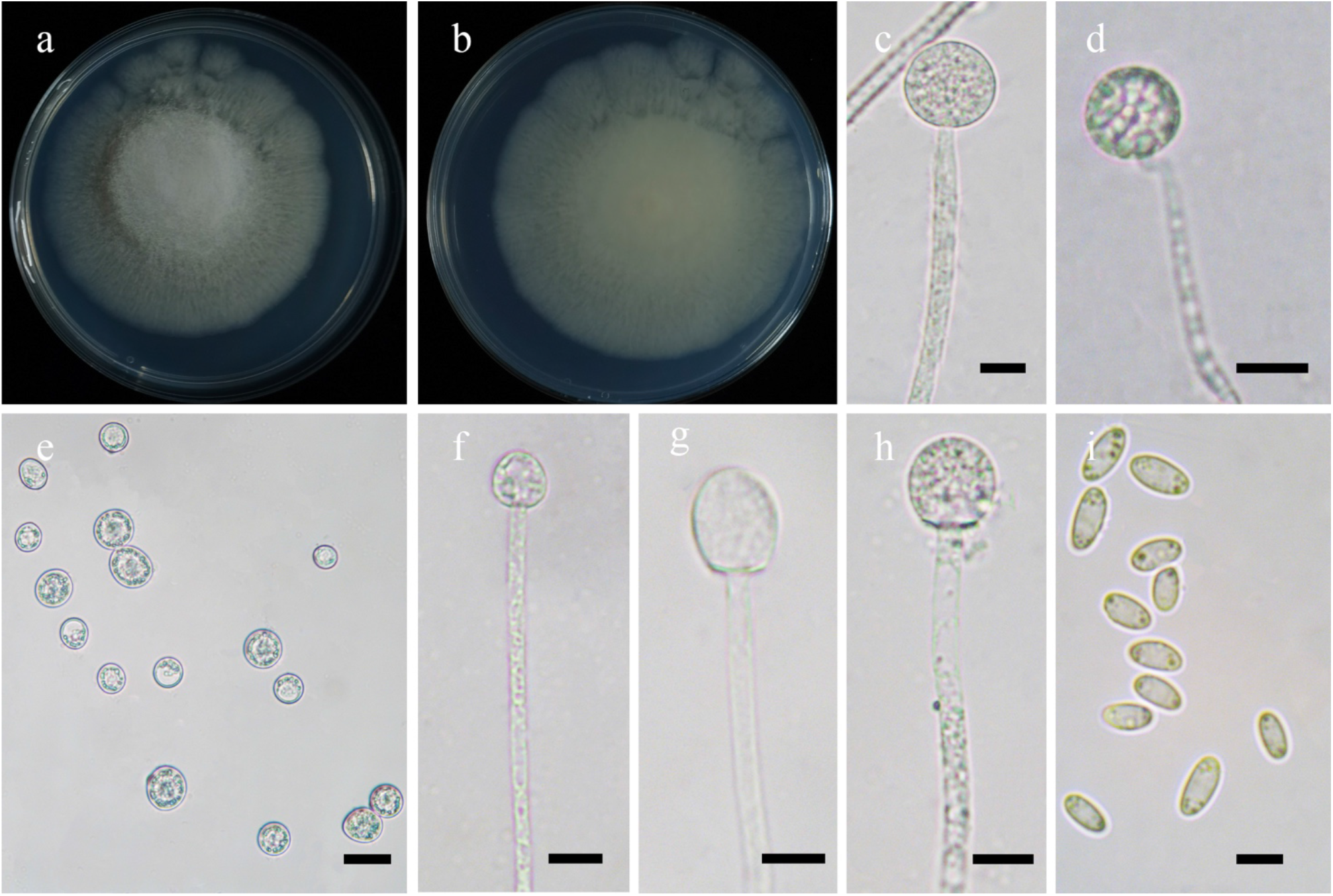
Morphologies of *Mucor hyalinosporus* ex-holotype CGMCC 3.16145. **a, b.** Colonies on PDA (**a.** obverse, **b.** reverse); **c, d.** Sporangia; **e.** Chlamydospores; **f–h.** Columellae without collar; **i.** Sporangiospores. — Scale bars: **c, d, f–h.** 10 μm, **e.** 10 μm, **i.** 5 μm.

*Fungal Names*: FN570935.

*Etymology*: *hyalinosporus* (Lat.) refers to the species having hyaline chlamydospores.

*Holotype*: HMAS 351547.

*Colonies* on PDA at 27 °C for 7 days, reaching 70 mm in diameter, initially white, soon becoming Sea-Foam Yellow, floccose, broadly lobed. *Hyphae* flourishing, firstly mainly in the center, then abundant throughout when old, 1.5–4.5 μm in diameter. *Rhizoids* absent*. Stolons* absent. *Sporangiophores* arising directly from substrate or aerial hyphae, erect or slightly bent, unbranched, always septate, slightly constricted on the top. *Sporangia* mostly globose, occasionally subglobose, brown when older, smooth, 13.0–31.0 μm in diameter. *Apophyses* absent. *Collars* absent, walls deliquescent. *Columellae* conical to cylindrical, hyaline or subhyaline, smooth, 9.5–21.0 μm long and 9.5–17.0 μm wide. *Sporangiospores* fusiform, hyaline, with droplets, 4.5–7.5 μm long and 2.0–4.0 μm wide. *Chlamydospores* borne on substrate hyphae, in chains, moniliform, ovoid, globose, rarely irregular, hyaline, with droplets, 10.5–20.5 μm long and 10.0–17.5 μm wide. *Zygospores* unknown.

*Material examined*: China, Beijing, from soil sample, 28 October 2021, Heng Zhao (holotype HMAS 351547, living ex-holotype culture CGMCC 3.16145).

*GenBank accession number*: OL678203.

*Notes*: *Mucor hyalinosporus* is closely related to *M. subtilissimus* Oudem. based on phylogenetic analysis of ITS rDNA sequences (Fig. 14). However, morphologically *M. subtilissimus* differs from *M. hyalinosporus* by producing collars (Hurdeal *et al*. 2021).

**58. *Mucor lobatus*** H. Zhao, Y.C. Dai & X.Y. Liu, sp. nov., Fig. 74.

**Fig. 74.**
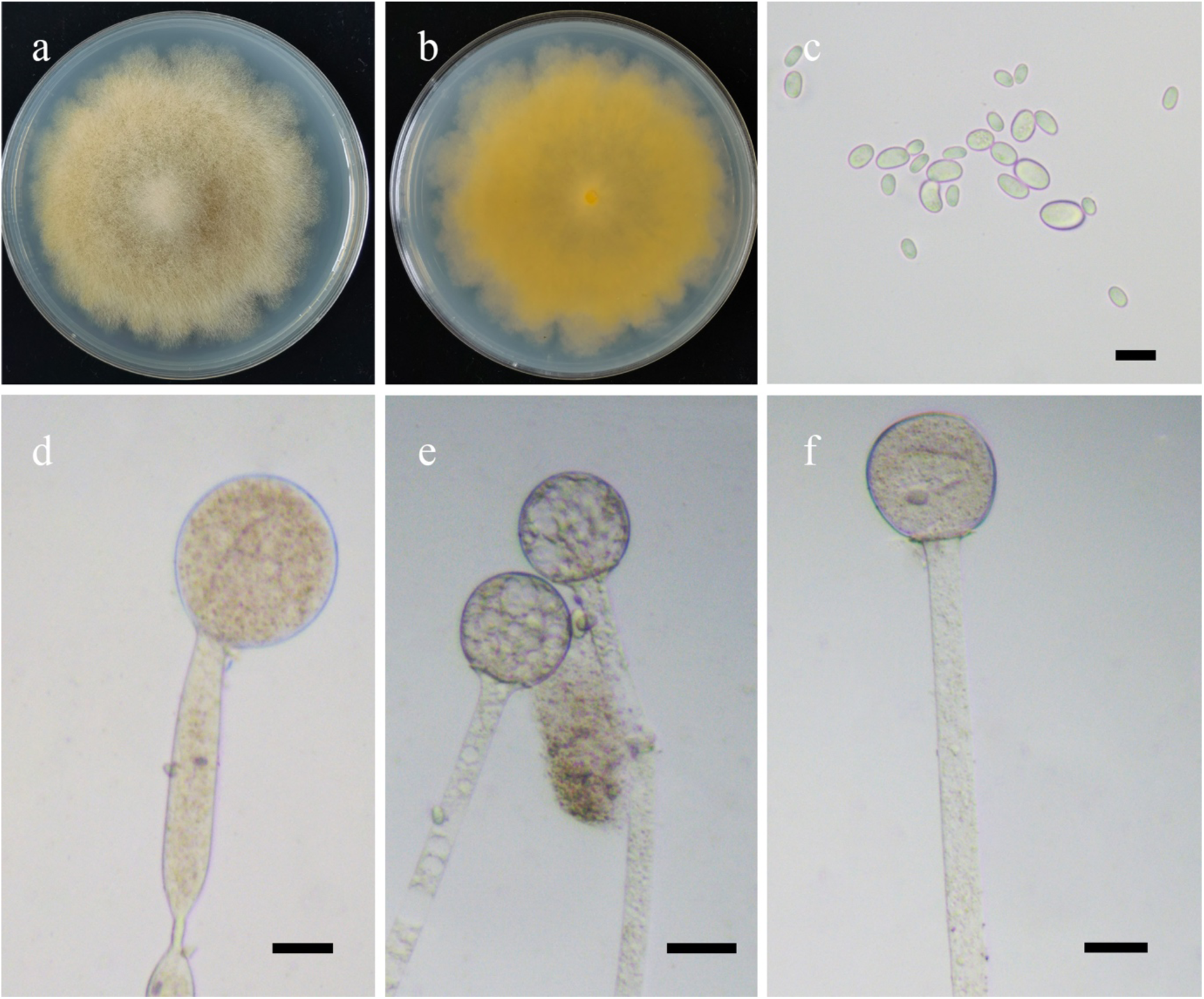
Morphologies of *Mucor lobatus* ex-holotype CGMCC 3.16146. **a, b.** Colonies on PDA (**a.** obverse, **b.** reverse); **c.** Sporangiospores; **d.** Immature sporangia; **e, f.** Columellae with sporangiophores. — Scale bars: **c.** 10 μm, **d–f.** 20 μm.

*Fungal Names*: FN570936.

*Etymology*: *lobatus* (Lat.) refers to the species having lobate colonies.

*Holotype*: HMAS 351548.

*Colonies* on PDA at 27 °C for 6 days, reaching 80 mm in diameter, 10 mm high, floccose, lobed, initially white, soon becoming Baryta Yellow, reverse irregular at margin and Capucine Yellow. *Hyphae* flourishing, unbranched or simply branched, aseptate, 7.5–19.5 μm in diameter. *Rhizoids* absent. *Stolons* absent. *Sporangiophores* arising from substrate hyphae, erect or bent, unbranched, aseptate, sometimes slightly constricted, very long. *Sporangia* globose, subhyaline, smooth, brown, 36.5–70.0 μm in diameter, walls deliquescent. *Apophyses* absent. *Collars* absent. *Columellae* subglobose or globose, hyaline or subhyaline, smooth, 14.0–32.5 μm long and 15.5– 31.0 μm wide. *Sporangiospores* ovoid, ellipsoid, 3.5–11.0 μm long and 3.0–7.5 μm wide. *Chlamydospores* absent. Z*ygospores* unknown.

*Materials examined*: China. Jiangsu Province, Yancheng, 33°41’55″N, 120°23’38″E, from soil sample, 28 May 2021, Heng Zhao (holotype HMAS 351548, living ex-holotype culture CGMCC 3.16146). Hubei Province, Huanggang, 30°59’46″N, 114°32’18″E, from soil sample, 31 May 2021, Heng Zhao (culture XY07029).

*GenBank accession numbers*: OL678204 and OL678203.

*Notes*: *Mucor lobatus* is closely related to *M. fusiformis* Walther & de Hoog, *M. hiemalis* Wehmer, and *M. japonicus* (Komin.) Walther & de Hoog based on phylogenetic analysis of ITS rDNA sequences (Fig. 14). However, morphologically *M. fusiformis*, *M. hiemalis* and *M. japonicus* differ from *M. lobatus* by producing zygospores (Hesseltine *et al*. 1959, Schipper 1986b, Walther *et al*. 2013).

**59. *Mucor moniliformis*** H. Zhao, Y.C. Dai & X.Y. Liu, sp. nov., Fig. 75.

**Fig. 75.**
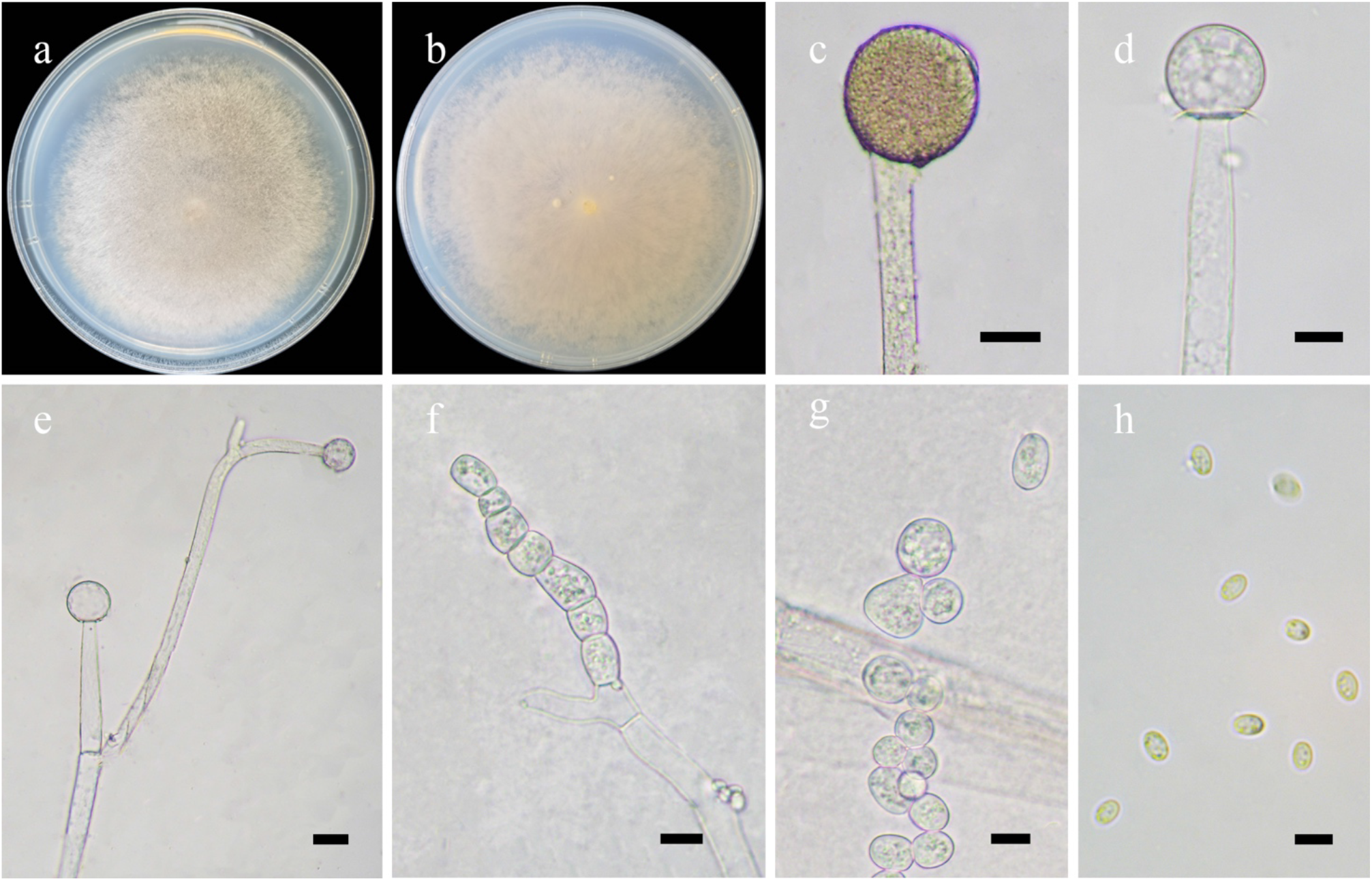
Morphologies of *Mucor moniliformis* ex-holotype CGMCC 3.17147. **a, b.** Colonies on PDA (**a.** obverse, **b.** reverse); **c.** Sporangia; **d.** Columellae with collars; **e.** Sporangiophores; **f, g.** Chlamydospores; **h.** Sporangiospores. — Scale bars: **c, e.** 20 μm, **d, f, g.** 10 μm, **h.** 5 μm.

*Fungal Names*: FN570937.

*Etymology*: *moniliformis* (Lat.) refers to the species having moniliform chlamydospores.

*Holotype*: HMAS 351549.

*Colonies* on PDA at 27 °C for 5 days, reaching 90 mm in diameter, 10 mm high, floccose, granulate, initially white, soon becoming Buff-Yellow, reverse irregular at margin. *Hyphae* flourishing, branched, aseptate when juvenile, septate with age, 4.0–23.5 μm in diameter. *Rhizoids* absent. *Stolons* absent. *Sporangiophores* arising from aerial or substrate hyphae, erect or slightly bent, simple branched, somewhere slightly constricted, 6.0–13.5 μm in diameter. *Sporangia* globose, hyaline when young, Light Brown or Dark Brown when old, smooth, 18.0–64.5 μm in diameter, walls deliquescent. *Apophyses* absent. *Collar* distinct present, or absent. *Columellae* subglobose and globose, hyaline or subhyaline, smooth, 15.0–30.5 μm long and 17.0–32.0 μm wide. *Sporangiospores* ovoid, hyaline or subhyaline, 3.5–5.5 μm long and 2.5–3.5 μm wide. *Chlamydospores* abundantly in substrate hyphae, in chains, ellipsoid, ovoid, subglobose, globose or irregular, 5.5–17.0 μm long and 7.0–15.5 μm wide. *Zygospores* unknown.

*Materials examined*: China. Zhejiang Province, Hangzhou, 30°07’15″N, 119°58’47″E, from soil sample, 4 June 2021, Jia-Jia Chen (holotype HMAS 351549, living ex-holotype culture CGMCC 3.16147). Hubei Province, Tianmen Mountain, from soil sample, 22 July 2021, Heng Zhao (living culture XY07458).

*GenBank accession numbers*: OL678206 and OL678207.

*Notes*: *Mucor moniliformis* is closely related to *M. brunneolus* H. Zhao *et al*. based on phylogenetic analysis of ITS rDNA sequences (Fig. 14). However, morphologically *M. brunneolus* differs from *M. moniliformis* by unbranched sporangiophores and fusiform sporangiospores.

**60. *Mucor orientalis*** H. Zhao, Y.C. Dai & X.Y. Liu, sp. nov., Fig. 76.

**Fig. 76.**
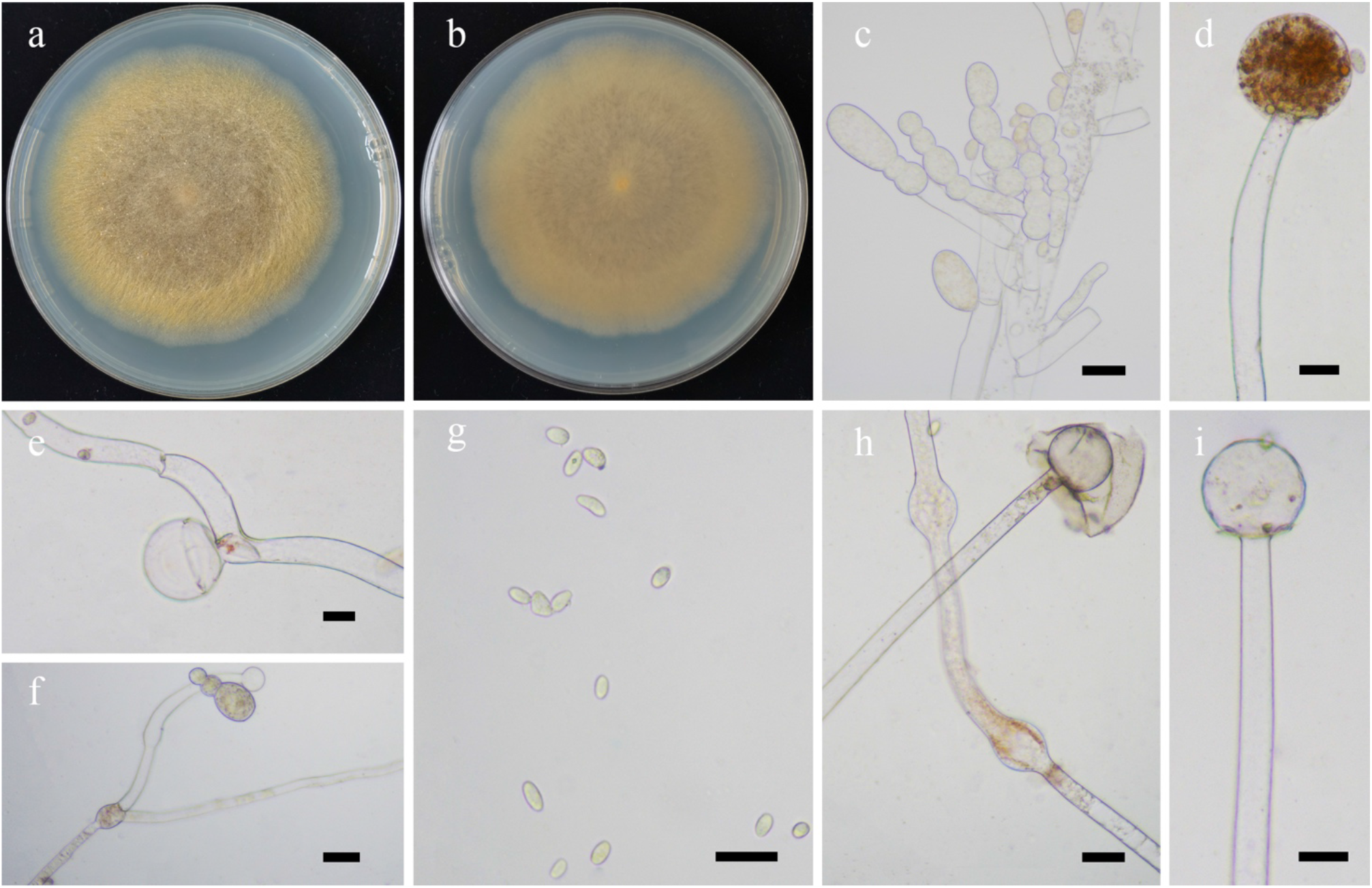
Morphologies of *Mucor orientalis* ex-holotype CGMCC 3.16148. **a, b.** Colonies on PDA (**a.** obverse, **b.** reverse); **c.** Chlamydospores; **d.** Fertile sporangia; **e, f.** Aborted sporangia on sporangiophores; **g.** Sporangiospores; **h, i.** Columellae with collars. — Scale bars: **c–i.** 20 μm.

*Fungal Names*: FN570938.

*Etymology*: *orientalis* (Lat.) refers to the locality of the holotype, Anhui Province, East Asia.

*Holotype*: HMAS 351550.

*Colonies* on PDA at 27 °C for 7 days, reaching 90 mm in diameter, initially white, soon becoming Wax Yellow, floccose, granulate, regular at margin. *Hyphae* flourishing, simply branched, aseptate when juvenile, septate with age, 7.0–31.0 μm in diameter. *Substrate hyphae* abundant, 7.0–25.0 μm in diameter. *Substrate hyphae* abundant. *Rhizoids* absent*. Stolons* absent. *Sporangiophores* arising directly from substrate or aerial hyphae, erect or recumbent, mainly unbranched, sometimes septate, slightly constricted, long or short. *Fertile sporangia* always terminally borne on the main axes, mostly globose, occasionally subglobose, light yellow to brown, smooth, 38.0–56.5 μm in diameter, walls deliquescent. *Aborted sporangia* borne laterally or intercalarily on sporangiophores. *Apophyses* absent. *Collars* sometimes present, always distinct, occasionally large. *Columellae* mostly globose, occasionally subglobose, hyaline, smooth, 17.0– 49.5 μm in diameter. *Sporangiospores* ovoid, ellipsoid, fusiform or irregular, 3.0–5.5 μm long and 2.0–3.5 μm wide. *Chlamydospores* arising from substrate hyphae, moniliform. *Zygospores* unknown.

*Material examined*: China, Anhui Province, Hefei, Lu’an, 31°27’36″N, 115°43’24″E, from soil sample, 28 May 2021, Heng Zhao (holotype HMAS 351550, living ex-holotype culture CGMCC 3.16148).

*GenBank accession number*: OL678208.

*Notes*: *Mucor orientalis* is closely related to *M. orantomantidis* Hyang B. Lee *et al*. based on phylogenetic analysis of ITS rDNA sequences (Fig. 14). However, morphologically *M. orantomantidis* differs from *M. orientalis* by producing zygospores and chlamydospores (Phookamsak *et al*. 2019).

**61. *Mucor radiatus*** H. Zhao, Y.C. Dai & X.Y. Liu, sp. nov., Fig. 77.

**Fig. 77.**
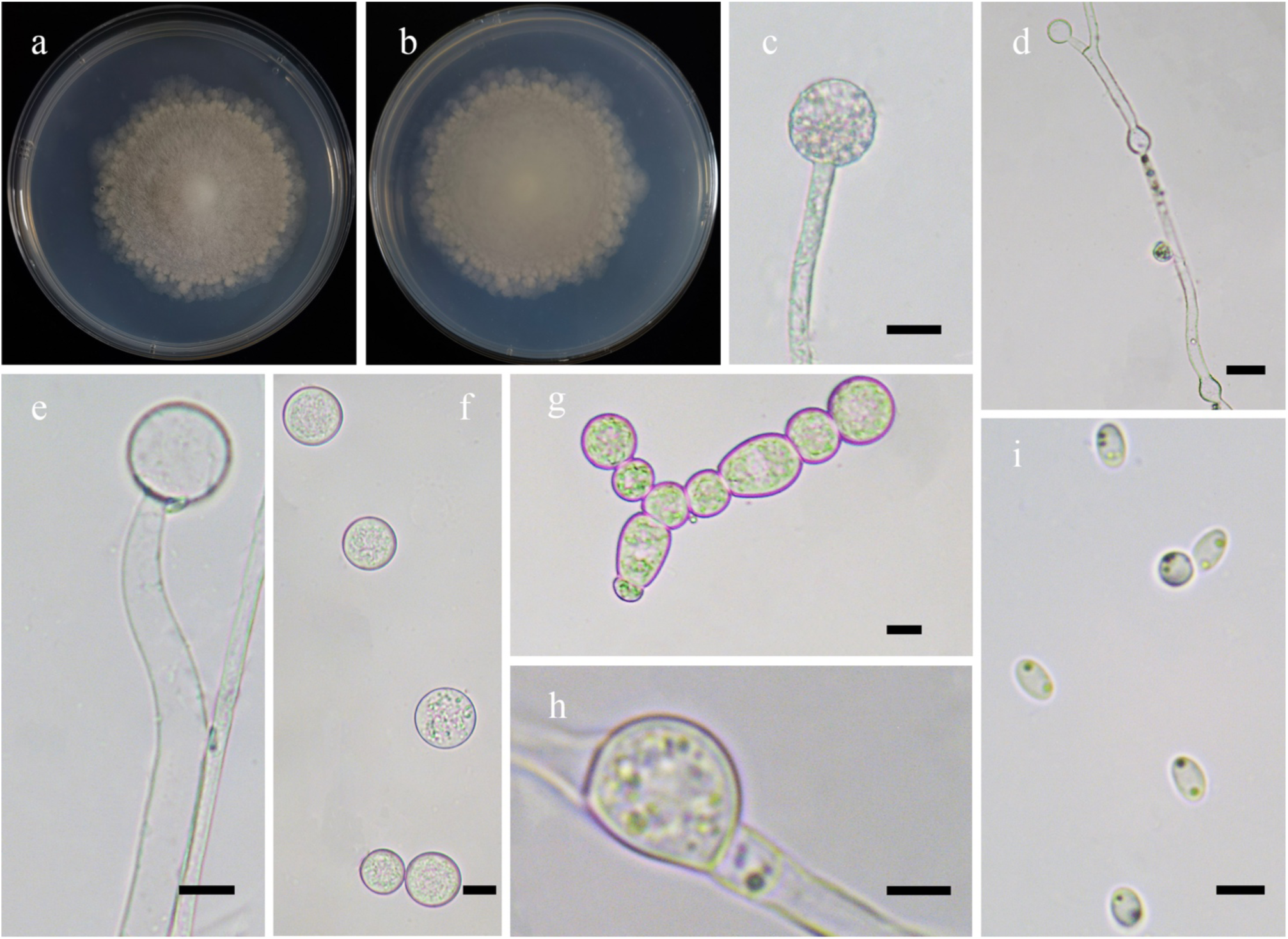
Morphologies of *Mucor radiatus* ex-holotype CGMCC 3.16149. **a, b.** Colonies on PDA (**a.** obverse, **b.** reverse); **c.** Sporangia; **d.** Aborted lateral sporangia with swellings in hyphae; **e.** Columellae; **f–h.** Chlamydospores; **i.** Sporangiospores. — Scale bars: **c, e–g.** 10 μm, **d.** 20 μm, **h, i.** 5 μm.

*Fungal Names*: FN570939.

*Etymology*: *radiatus* (Lat.) refers to the species having radiating colonies.

*Holotype*: HMAS 351551.

*Colonies* on PDA at 27 °C for 10 days, slow growing, reaching 60 mm in diameter, radiating, initially white, gradually becoming Light Brown, floccose, irregular at margin. *Hyphae* flourishing when old, branched, 2.5–9.0 μm in diameter. *Rhizoids* absent. *Stolons* absent. *Sporangiophores* arising from substrate or aerial hyphae, erect or curved, unbranched. *Fertile sporangia* always borne terminally on main axes, globose, smooth, 19.0–34.5 μm in diameter, walls deliquescent. *Aborted sporangia* mainly forming laterally or intercalary on sporangiophores, globose, monopodial, hyaline, thick-walled, 10.5–13.0 μm in diameter. *Apophyses* absent. *Collars* absent. *Columellae* forming between two aborted sporangia, subglobose to globose, hyaline, 12.5–26.5 μm long and 12.5–25.5 μm wide. *Sporangiospores* two types: mainly fusiform, with droplets, 3.5–5.5 μm long and 1.5–3.0 μm wide; and globose (rare), 3.0–5.0 μm in diameter. *Chlamydospores* abundantly produced on substrate hyphae and rarely on aerial hyphae, in chains, cylindrical, ellipsoid, ovoid or irregular when young, mostly globose, occasionally subglobose when old, 11.0–18.0 μm in diameter. *Zygospores* unknown.

*Materials examined*: China, Yunnan Province, Chuxiong, Mouding Country, 25°18’44″N, 113°12’ 40″E, from soil sample, 30 September 2021, Heng Zhao (holotype HMAS 351551, living ex-holotype culture CGMCC 3.16149, and living culture XY08167).

*GenBank accession numbers*: OL678209 and OL678210.

*Notes*: See notes of *Mucor abortisporangium*.

**62. *Mucor rhizosporus*** H. Zhao, Y.C. Dai & X.Y. Liu, sp. nov., Fig. 78.

**Fig. 78.**
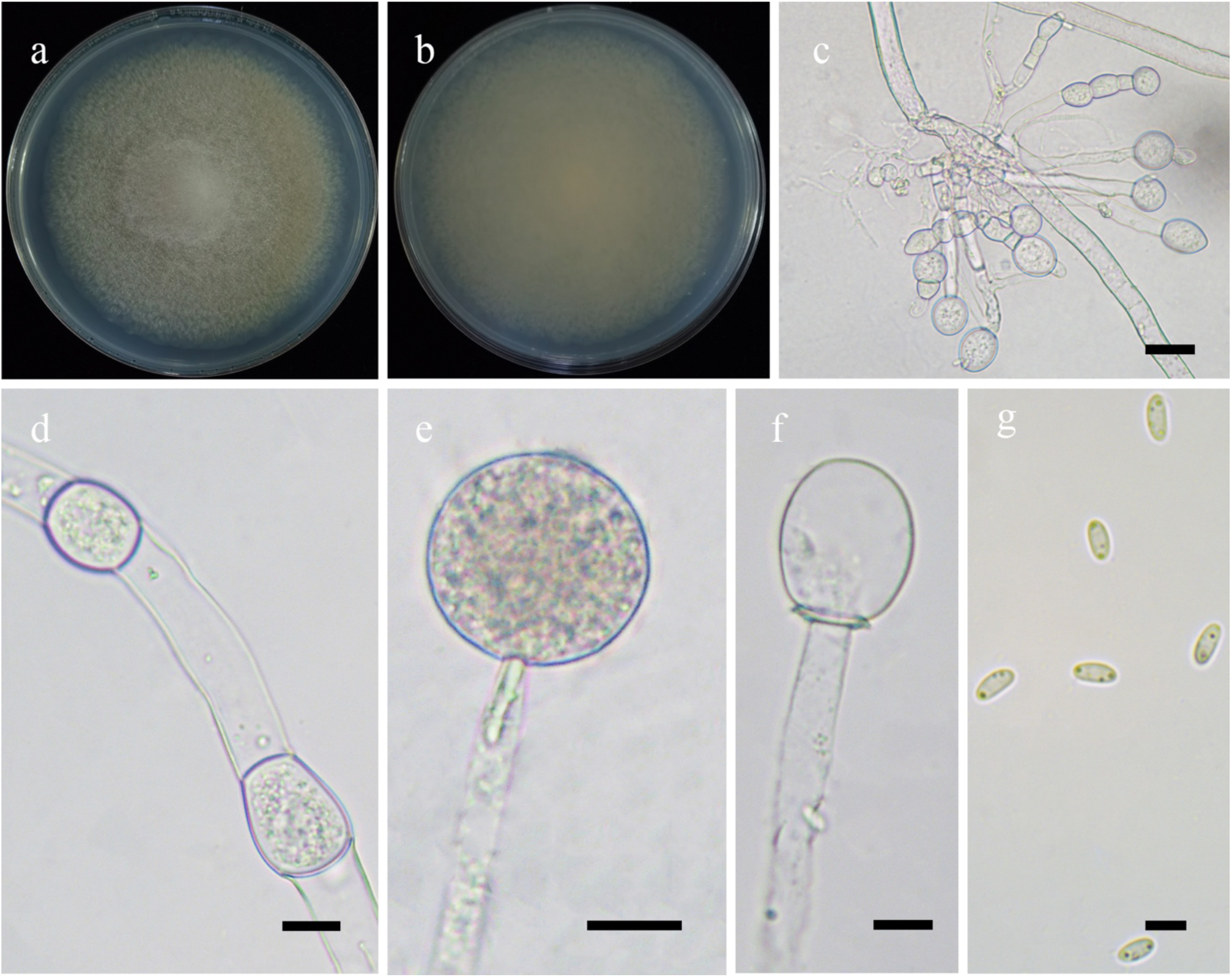
Morphologies of *Mucor rhizosporu*s ex-holotype CGMCC 3.16150. **a, b**. Colonies on PDA (**a.** obverse, **b.** reverse); **c.** Chlamydospores at rhizoids; **d.** Chlamydospores in aerial hyphae; **e.** Sporangia; **f.** Columellae; **g.** Sporangiospores. — Scale bars: **c–f.** 10 μm, **g.** 5 μm.

*Fungal Names*: FN570940.

*Etymology*: *rhizosporus* (Lat.) refers to the species producing chlamydospores in the rhizoids.

*Holotype*: HMAS 351552.

*Colonies* on PDA at 27 °C for 7 days, slow growing, reaching 80 mm in diameter, first white, gradually becoming Pale Yellow-Orange, floccose, irregular at margin. *Hyphae* flourishing, branched, aseptate when juvenile, septate with age, 2.5–11.5 μm in diameter. *Rhizoids* abundant, root-like, branched, swollen, always producing chlamydospores. *Stolons* present, abundant. *Sporangiophores* arising from substrate and aerial hyphae, erect, unbranched. *Sporangia* globose, smooth, 22.0–37.5 μm in diameter, walls deliquescent. *Apophyses* absent. *Collars* always absent. *Columellae* ovoid to oblong, hyaline, 13.5–31.5 μm long and 10.0–22.5 μm wide. *Sporangiospores* fusiform or ellipsoid, 5.0–6.0 μm long and 2.0–3.0 μm wide. *Chlamydospores* produced in aerial hyphae and rhizoids; abundant in rhizoids, in chains, variable in shape when young, globose or subglobose when old, in aerial hyphae, cylindrical, ellipsoid, ovoid, subglobose to globose or irregular, 10.0–19.5 μm long and 8.0–13.5 μm wide. *Zygospores* unknown.

*Material examined*: China, Guangxi Auto Region, Nanning, from soil sample, 29 August 2021, Heng Zhao (holotype HMAS 351552, living ex-holotype culture CGMCC 3.16150).

*GenBank accession number*: OL678211.

*Notes*: *Mucor rhizosporus* is closely related to *M. pernambucoensis* C.L. Lima *et al*. based on phylogenetic analysis of ITS rDNA sequences (Fig. 14). However, morphologically *M. pernambucoensis* differs from *M. rhizosporus* by larger sporangiospores (4.5–14.5 μm long and 2.5–5 μm wide vs. 5.0–6.0 μm long and 2.0–3.0 μm wide), sympodially branched sporangiophores, variable columellae, and neither rhizoids nor stolons (de Lima *et al*. 2018).

**63. *Mucor robustus*** H. Zhao, Y.C. Dai & X.Y. Liu, sp. nov., Fig. 79.

**Fig. 79.**
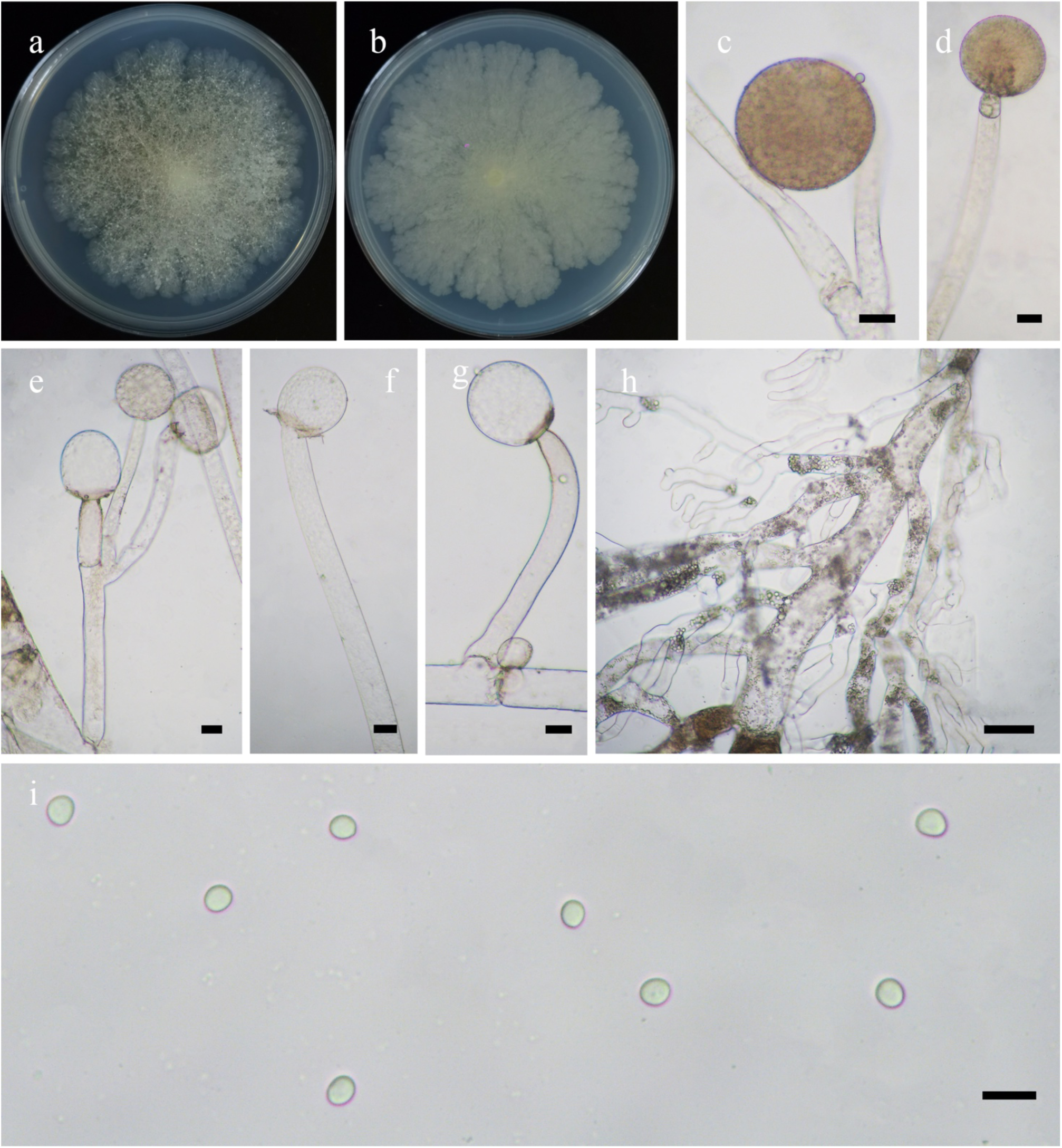
Morphologies of *Mucor robustus* ex-holotype CGMCC 3.16151. **a, b.** Colonies on PDA (**a.** obverse, **b.** reverse); **c, d.** Sporangia; **e–g.** Sporangiophores with columellae; **h.** Substrate hyphae; **i.** Sporangiospores. — Scale bars: **c–g.** 20 μm, **h.** 50 μm, **i.** 10 μm.

*Fungal Names*: FN570941.

*Etymology*: *robustus* (Lat.) refers to the species having robust hyphae.

*Holotype*: HMAS 351553.

*Colonies* on PDA at 27 °C for 7 days, reaching 80 mm in diameter, granulate, broadly lobed and zonate, initially white, gradually becoming Maize Yellow. *Hyphae* unbranched or simple branched, robust, hyaline, 14.0–77.0 μm in diameter, substrate hyphae abundant, root-like, branched. *Rhizoids* and *stolons* absent. *Sporangiophores* arising directly from substrate hyphae, erect to curved, unbranched, simple branched, or repeatedly unbranched, rarely with septa at the base on sporangiospores, generally constricted on the top, very long or short. *Sporangia* mostly globose, occasionally subglobose multi-spored, brown to black, 33.0–160.0 μm in diameter, walls deliquescent. *Columellae* hemispherical, conical, subglobose to globose, hyaline or subhyaline, rough, sometimes with pigment, 14.5–109.0 μm long and 16.0–105.5 μm wide. *Apophyses* absent. *Collar* usually absent, small if present. *Sporangiospores* ovoid to globose, 4.0–7.0 μm long and 3.5–6.0 μm wide. *Chlamydospores* absent. *Zygospores* unknown.

*Materials examined*: China, Xinjiang Auto Region, Altay, Burqin Country, 48°39’29″N, 87°2’9″E, from soil sample, 31 October 2021, Heng Zhao (holotype HMAS 351553, living ex-holotype culture CGMCC 3.16151, and living culture XY08976 and XY09024).

*GenBank accession numbers*: OL678212, OL678213 and OL678214.

*Notes*: *Mucor robustus* is closely related to *M. aligarensis* B.S. Mehrotra & B.R. Mehrotra, *M. flavus* (Mart.) Fr. and *M. rongii* F.R. Bai & C. Cheng based on phylogenetic analysis of ITS rDNA sequences (Fig. 14). However, morphologically *M. aligarensis* differs from *M. robustus* by larger sporangiospores (6.6–15.4 μm long and 5.5–11 μm wide vs. 4.0–7.0 μm long and 3.5–6.0 μm wide; Mehrotra & Mehrotra 1969). *M. flavus* differs from *M. robustus* by fusiform sporangiophores (https://www.mycobank.org). *M. rongii* differs from *M. robustus* by broadly ellipsoid to cylindrical or rather irregular sporangiospores (Bai *et al*. 2021).

**64. *Mucor sino-saturninus*** H. Zhao, Y.C. Dai & X.Y. Liu, sp. nov., Fig. 80.

**Fig. 80.**
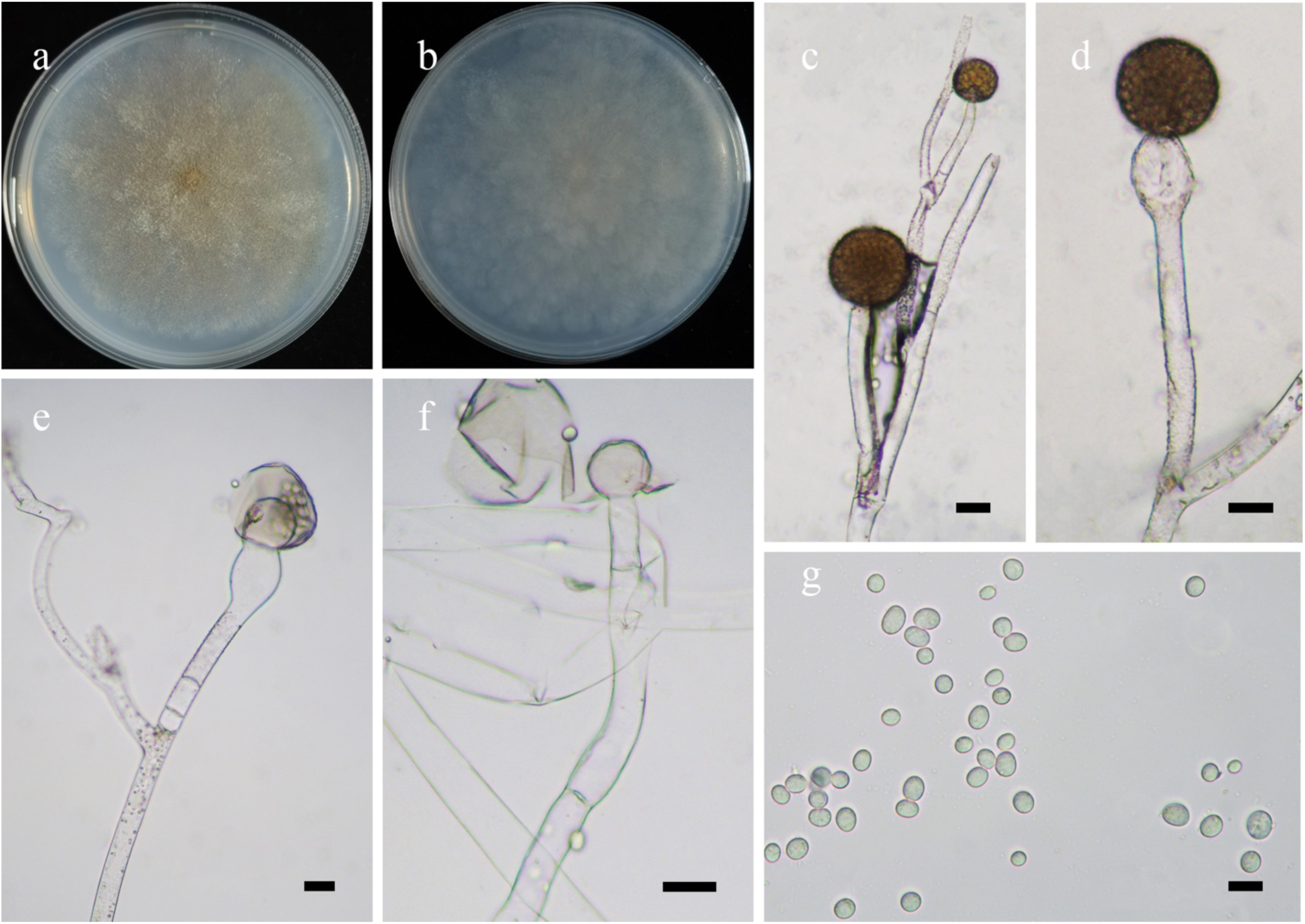
Morphologies of *Mucor sino-saturninus* ex-holotype CGMCC 3.16152. **a, b.** Colonies on PDA (**a.** obverse, **b.** reverse); **c.** Sporangiophores branched; **d.** Swellings below sporangia; **e, f.** Columellae with collars; **g.** Sporangiospores. — Scale bars: **c–f.** 20 μm, **g.** 10 μm.

*Fungal Names*: FN570942.

*Etymology*: *sino-saturninus* (Lat.) refers to the species being like *Mucor saturninus* but occurring in China.

*Holotype*: HMAS 351554.

*Colonies* on PDA at 27 °C for 7 days, slow growing, reaching 80 mm in diameter, broadly lobed, initially white, gradually becoming Ochraceous-Tawny to Dresden Brown, granulate. *Hyphae* unbranched or simple branched, hyaline, 7.5–43.5 μm in diameter. *Rhizoids* absent. *Stolons* absent. *Sporangiophores* arising directly from substrate or aerial hyphae, erect or bent, simple branched, sympodial, septate usually at the base and sometimes one to several below sporangia, generally constricted on the top, sometimes swollen below sporangia. *Sporangia* mostly globose, occasionally subglobose, brownish, with spines, 23.0–105.0 μm in diameter, walls deliquescent. *Apophyses* absent. *Collars* always present, distinct. *Columellae* subglobose to globose, hyaline, rough or smooth, 16.0–37.5 μm long and 14.5–37.0 μm wide. *Sporangiospores* ovoid to globose, subhyaline, 4.5–10.5 μm long and 3.0–7.0 μm wide. *Chlamydospores* absent. *Zygospores* unknown.

*Materials examined*: China, Yunnan Province, Diqing, Deqen Country, 98°46’15′′E, 28°27’38′′N, from soil sample, 19 October 2021, Heng Zhao (holotype HMAS 351554, living ex-holotype culture CGMCC 3.16152, and living culture XY08556).

*GenBank accession numbers*: OL678215 and OL678216.

*Notes*: *Mucor sino-saturninus* is closely related to *M. saturninus* Hagem based on phylogenetic analysis of ITS rDNA sequences (Fig. 14). However, morphologically *M. saturninus* differs from *M. sino-saturninus* by larger columellae (35–100 μm long and 25–90 μm wide vs. 16.0–37.5 μm long and 14.5–37.0 μm wide) and broadly ellipsoid sporangiospores (Hagem 1910).

**65. *Syncephalastrum breviphorum*** H. Zhao, Y.C. Dai & X.Y. Liu, sp. nov., Fig. 81.

**Fig. 81.**
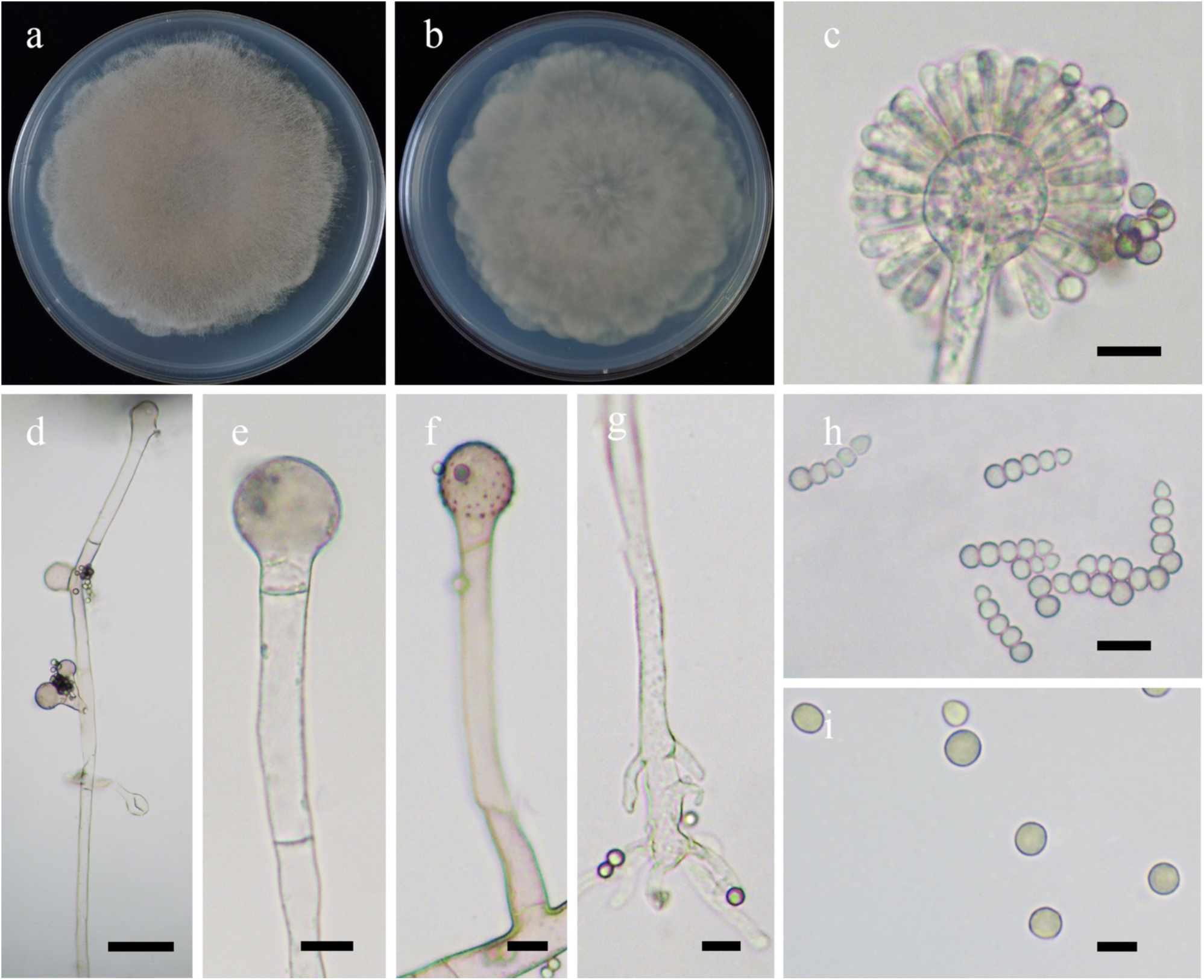
Morphologies of *Syncephalastrum breviphorum* ex-holotype CGMCC 3.16153. **a, b.** Colonies on PDA (**a.** obverse, **b.** reverse); **c.** Merosporangia; **d–f.** Sporangiophores with branched and vesicles; **g.** Rhizoids; **h, i.** Sporangiospores. — Scale bars: **c, e–h.** 10 μm, **d.** 20 μm, **i. 5** μm.

*Fungal Names*: FN570943.

*Etymology*: *breviphorum* (Lat.) refers to the species having short lateral sporangiophores.

*Holotype*: HMAS 351555.

*Colonies* on PDA at 27 °C for 6 days, reaching 90 mm in diameter, zonate, scaly, floccose, initially white, soon becoming Buckthorn Brown, reverse irregular at margin. *Hyphae* flourishing, branched, hyaline or slightly pigmented, aseptate at first, septate with age, 5.5–15.5 μm in diameter. *Rhizoids* root-like or finger-like, branched. *Stolons* present. *Sporangiophores* arising from aerial hyphae, erect, bent, even curved, simple or sympodially branched, abundant, and shorter when laterally branched. *Vesicles* globose to subglobose, subtended or irregular, rough, hyaline to subhyaline, sometimes with one septum to two septa, 12.5–21.5 μm in diameter, or 17.5–35.0 μm long and 19.0–29.0 μm wide. *Merosporangia* borne over entire surface of merosporangia, containing a single row of 3–5 sporangiospores, frequently five-spored, 10.6–16.5 μm long and 2.0– 4.0 μm wide. *Apophyses* absent. *Collars* absent. *Columellae* degenerated. *Sporangiospores* striated, variable shape, mainly globose, 3.0–5.0 μm in diameter, sometimes subglobose, ovoid, cylindrical or ellipsoid, 3.0–4.5 μm long and 2.5–3.5 μm wide. *Chlamydospores* absent. *Zygospores* unknown.

*Materials examined*: China, Fujian Province, Fuzhou, Fujian Academy of Agricultural Sciences, from paper sample, 1 October 2013, Xiao-Yong Liu (holotype HMAS 351555, living ex-holotype culture CGMCC 3.16153, and living culture XY06838).

*GenBank accession numbers*: OL678217 and OL678218.

*Notes*: *Syncephalastrum breviphorum* is closely related to *S. contaminatum* A.S. Urquhart & A. Idnurm based on phylogenetic analysis of ITS rDNA sequences (Fig. 15). However, morphologically *S. contaminatum* differs from *S. breviphorum* by globose sporangiospores and frequently four-spored merosporangia (Urquhart & Idnurm 2020).

**66. *Syncephalastrum elongatum*** H. Zhao, Y.C. Dai & X.Y. Liu, sp. nov., Fig. 82.

**Fig. 82.**
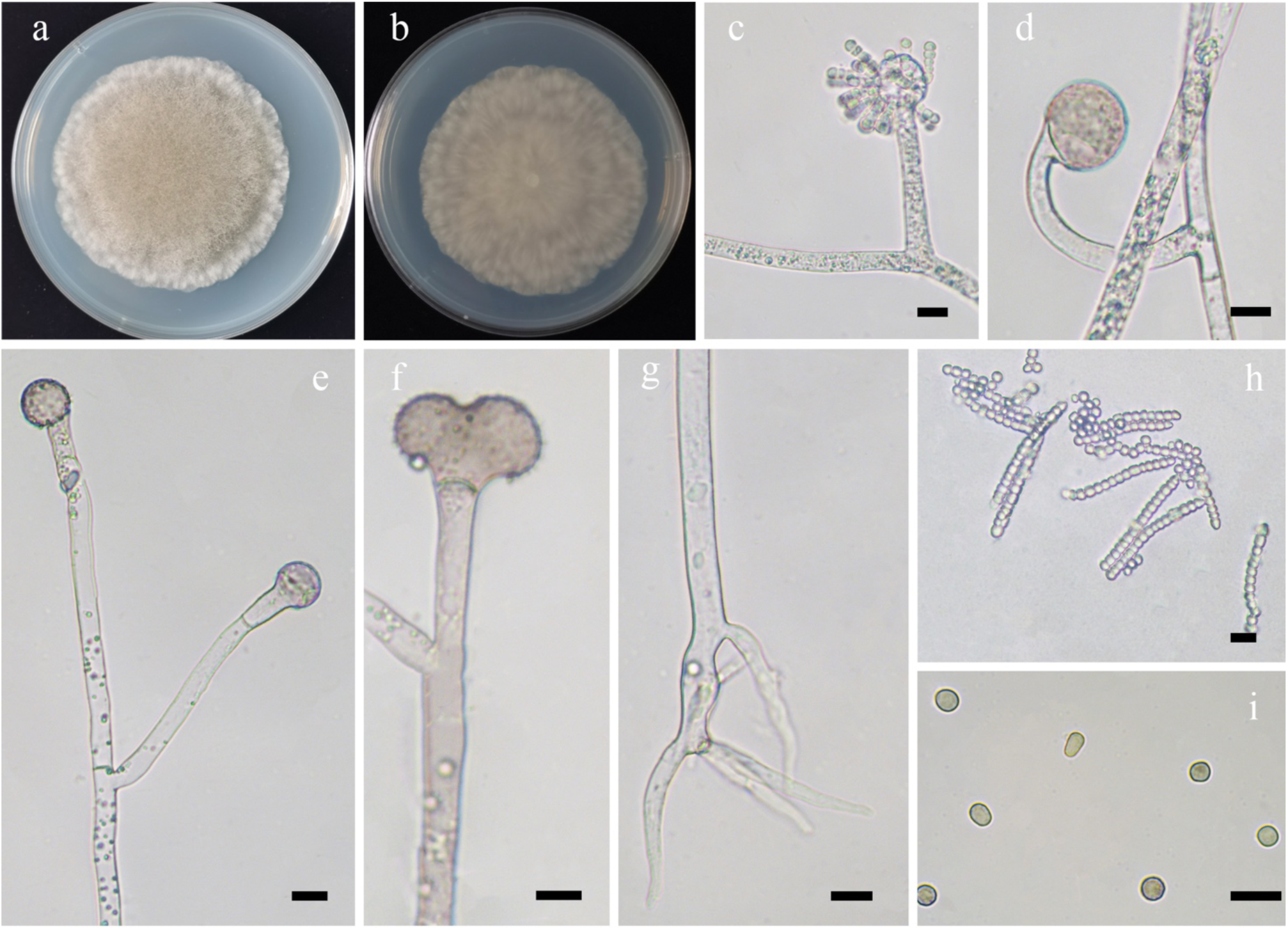
Morphologies of *Syncephalastrum elongatum* ex-holotype CGMCC 3.16154. **a, b.** Colonies on PDA (**a.** obverse, **b.** reverse); **c–f.** Sporangiophores with branched and vesicles; **g.** Rhizoids; **h, i.** Sporangiospores. — Scale bars: **c–i.** 10 μm.

*Fungal Names*: FN570944.

*Etymology*: *elongatum* (Lat.) refers to the species having long merosporangia.

*Holotype*: HMAS 351556.

*Colonies* PDA at 27 °C for 6 days, reaching 90 mm in diameter, scaly, floccose, granulate, initially white, soon becoming Clay Color, reverse irregular at margin. *Hyphae* flourishing, branched, aseptate at first, septate with age, hyaline or slightly pigmented, 3.5–24.0 μm in diameter. *Rhizoids* rare, root-like or finger-like. *Sporangiophores* arising from aerial hyphae, erect, bent, even curved, simple branched or unbranched, shorter and always curved when laterally branched, sometimes septate, 4.5–13.5 μm in wide. *Vesicles* subglobose, globose or subtended, rough, hyaline or Light Brown, 8.5–60.5 μm long and 10.0–45.0 μm wide. *Merosporangia* borne on entire surface of vesicles, containing a single row of 4–18 sporangiospores, frequently more than ten-spored, 11.5–51.5 μm long and 3.0–4.0 μm wide. *Apophyses* present. *Collars* absent. *Columellae* degenerated. *Sporangiospores* with striation, variable shape, subglobose, 3.0–5.5 μm in diameter, sometimes subglobose, ovoid, cylindrical or ellipsoid, 3.5–5.5 μm long and 2.5–3.5 μm wide. *Chlamydospores* absent. *Zygospores* unknown.

*Material examined*: China, Fujian Province, Fuzhou, Fujian Academy of Agricultural Sciences, from paper sample, 1 October 2013, Xiao-Yong Liu (holotype HMAS 351556, living ex-holotype culture CGMCC 3.16154).

*GenBank accession number*: OL678219.

*Notes*: *Syncephalastrum elongatum* is closely related to *S. verruculosum* P.C. Misra based on phylogenetic analysis of ITS rDNA sequences (Fig. 15). However, morphologically *S. verruculosum* differs from *S. elongatum* by merosporangia (13.0–23.0 μm long and 4.0–7.0 μm wide, containing a single row of 2–6 sporangiospores vs. 11.5–51.5 μm long and 3.0–4.0 μm wide, containing a single row of 4–18 sporangiospores), and the absence of rhizoids (Misra 1975).

**67. *Syncephalastrum simplex*** H. Zhao, Y.C. Dai & X.Y. Liu, sp. nov., Fig. 83.

**Fig. 83.**
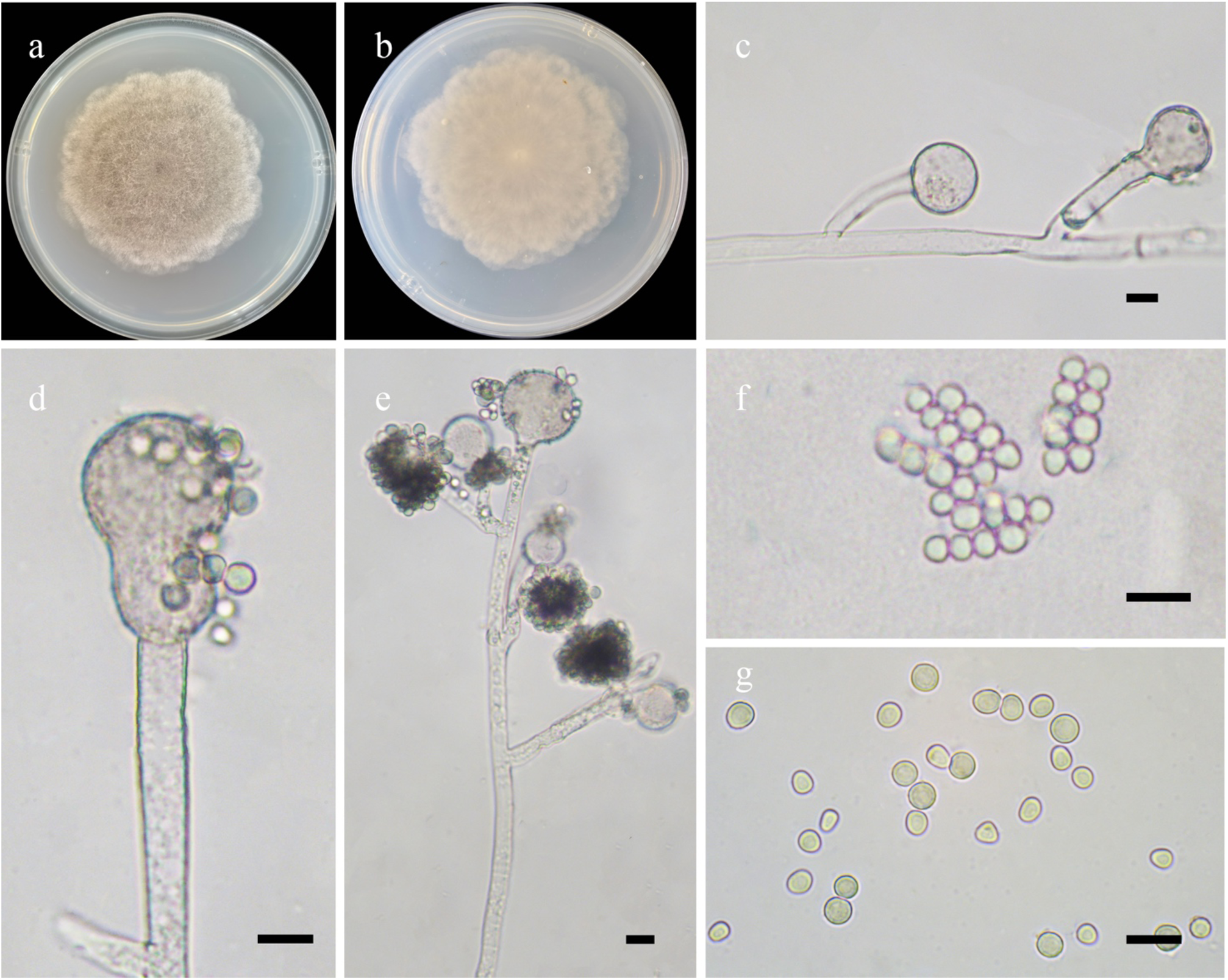
Morphologies of *Syncephalastrum simplex* ex-holotype CGMCC 3.16155. **a, b.** Colonies on PDA (**a.** obverse, **b.** reverse); **c, d.** Vesicles; **e.** Sporangiophores; **f, g.** Sporangiospores. — Scale bars: **c–g.** 10 μm.

*Fungal Names*: FN570945.

*Etymology*: *simplex* (Lat.) refers to the species having simple branching of the sporangiophores.

*Holotype*: HMAS 351557.

*Colonies* on PDA at 27 °C for 7 days, reaching 90 mm in diameter, lobed, floccose, granulate, initially white, soon becoming Clove Brown, reverse irregular at margin. *Hyphae* flourishing, branched, aseptate when juvenile, septate with age, 5.5–14.5 μm in diameter. *Rhizoids* absent. *Stolons* absent. *Sporangiophores* arising from aerial hyphae, erect, slightly bent, simple branched, or unbranched, 2.5–11.5 μm wide. *Vesicles* subglobose, globose or irregular, rough, hyaline to Light Brown, some with septa, 6.5–28.5 μm long and 7.5–28.0 μm wide. *Merosporangia* borne on entire surface of vesicles, containing a single row of 3–9 sporangiospores, mainly four- to five-spored. *Apophyses* present. *Collars* absent. *Columellae* degenerated. *Sporangiospores* with striation, variable shape, mainly globose, 3.5–6.5 μm in diameter, subglobose, ovoid or ellipsoid, 3.0–7.5 μm long and 2.5–5.5 μm wide. *Chlamydospores* absent. *Zygospores* unknown.

*Material examined*: China, Hubei Province, Shennongjia National Nature Reserve, from soil sample, 1 October 2013, Xiao-Yong Liu (holotype HMAS 351557, living ex-holotype culture CGMCC 3.16155).

*GenBank accession number*: OL678220.

*Notes*: *Syncephalastrum simplex* is closely related to *S. monosporum* R.Y. Zheng *et al*. based on phylogenetic analysis of ITS rDNA sequences (Fig. 15). However, morphologically *S. monosporum* differs from *S. simplex* by the absence of rhizoids or stolons (Zheng *et al*. 1988).

**68. *Syncephalastrum sympodiale*** H. Zhao, Y.C. Dai & X.Y. Liu, sp. nov., Fig. 84.

**Fig. 84.**
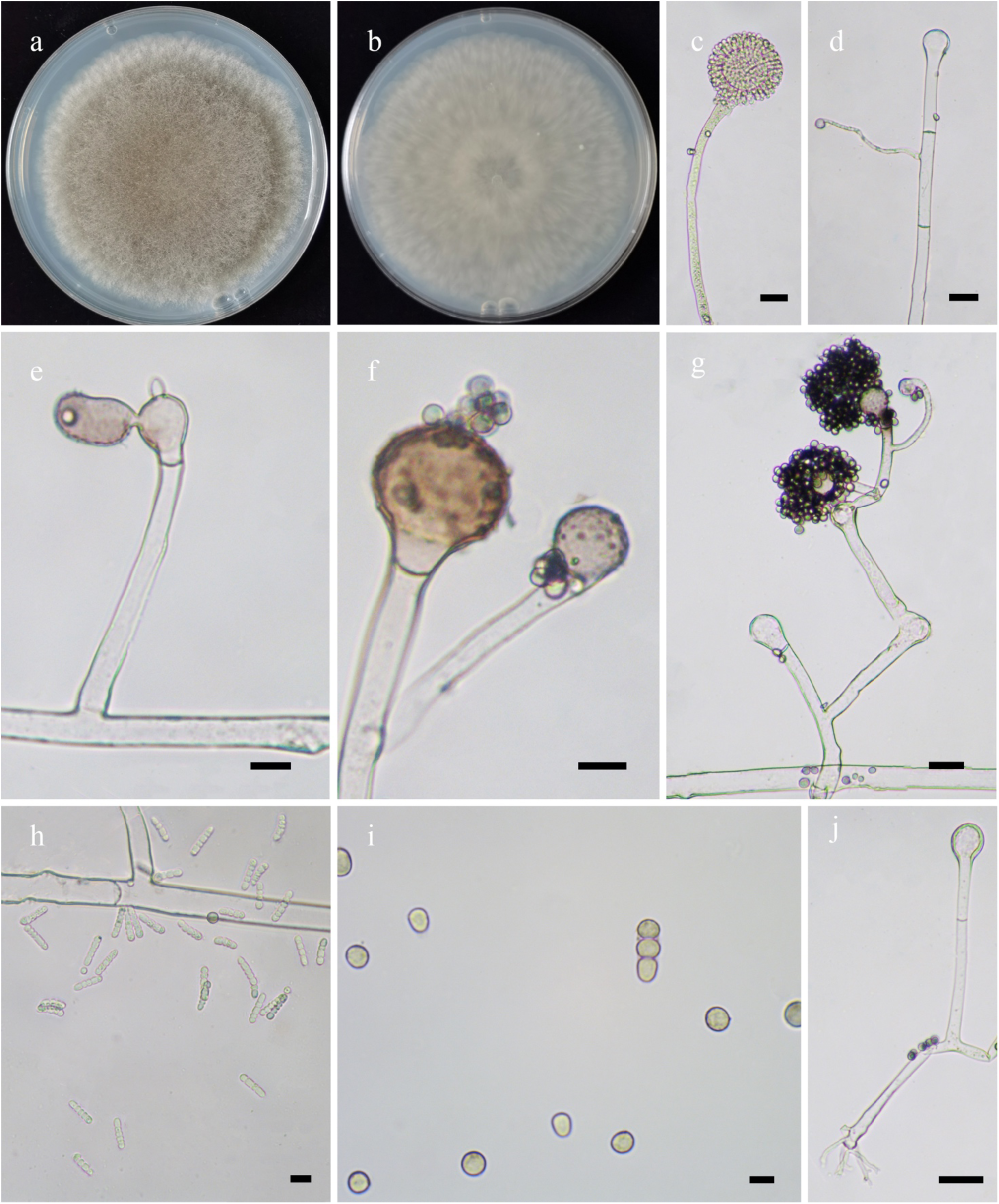
Morphologies of *Syncephalastrum sympodiale* ex-holotype CGMCC 3.16156. **a, b.** Colonies on PDA (**a.** obverse, **b.** reverse); **c.** Merosporangia; **d–g.** Sporangiophores with branches and vesicles; **h, i.** Sporangiospores; **j.** Rhizoids. — Scale bars: **c, g, j. 2**0 μm, **d–f, h.** 10 μm, **i.** 5 μm.

*Fungal Names*: FN570946.

*Etymology*: *sympodiale* (Lat.) refers to the species having sympodial branching of the sporangiophores.

*Holotype*: HMAS 351558.

*Colonies* on PDA at 27 °C for 7 days, reaching 80 mm in diameter, small scaly, floccose, granulate, initially white, gradually becoming Buckthorn Brown to Dresden Brown, reverse irregular at margin. *Hyphae* flourishing, branched, hyaline, aseptate at first, septate when old, 3.5– 16.5 μm in diameter. *Rhizoids* mostly root-like or finger-like, rarely unbranched. *Sporangiophores* arising from aerial hyphae, erect, bent, even curved, usually sympodial, unbranched, or simple branched, sometimes with one septum or two septa blow apophyses, shorter and always curved when laterally branched, 4.5–11.5 μm in wide. *Vesicles* mainly subglobose or globose, rarely subtended or irregular, rough, hyaline or Light Brown, 9.0–35.0 μm long and 8.5–30.5 μm wide. *Merosporangia* borne over entire surface of vesicles, containing a single row of 2–7 sporangiospores, frequently five- or six-spored, 6.0–19.5 μm long and 2.5–4.0 μm wide. *Apophyses* present. *Collars* absent. *Columellae* degenerated. *Sporangiospores* with striation, variable shape, mainly globose, 4.0–6.0 μm in diameter, sometimes subglobose, ovoid or ellipsoid, 4.0–6.0 μm long and 3.0–5.0 μm wide. *Chlamydospores* absent. *Zygospores* unknown.

*Materials examined*: China. Beijing, from soil sample, 1 October 2013, Xiao-Yong Liu (holotype HMAS 351558, living ex-holotype culture CGMCC 3.161536, and living culture XY06803). Hubei Province, Shennongjia National Nature Reserve, from mushroom sample, 1 October 2013, Xiao-Yong Liu (living culture XY06811).

*GenBank accession numbers*: OL678221, OL678222 and OL678223.

*Notes*: *Syncephalastrum sympodiale* is closely related to *S. elongatum* H. Zhao *et al*. and *S. verruculosum* P.C. Misra based on phylogenetic analysis of ITS rDNA sequences (Fig. 15). However, morphologically *S. verruculosum* differs from *S. sympodiale* by longer merosporangia (11.5–51.5 μm long vs. 6.0–19.5 μm long), the absence of sympodially branched sporangiophores and rhizoids. And *S. verruculosum* differs from *S. sympodiale* by larger vesicles (45–90 μm in diameter vs. 9.0–35.0 μm long and 8.5–30.5 μm wide), no sympodially branched sporangiophores, and the absence of rhizoids (Misra 1975).

**69. *Umbelopsis brunnea*** H. Zhao, Y.C. Dai & X.Y. Liu, sp. nov., Fig. 85.

**Fig. 85.**
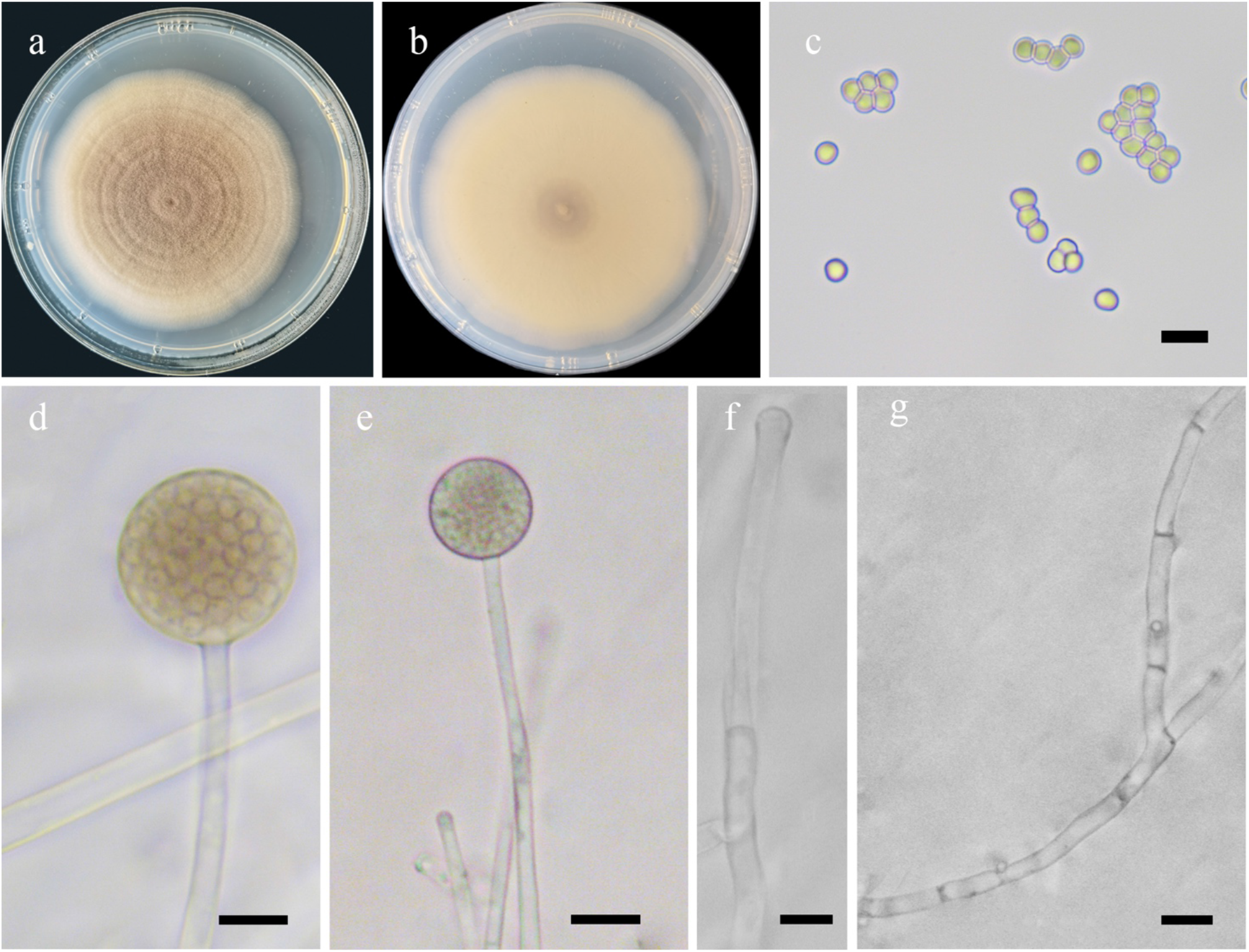
Morphologies of *Umbelopsis brunnea* ex-holotype CGMCC 3.16023. **a, b.** Colonies on PDA (**a,** obverse, **b,** reverse); **c.** Sporangiospores; **d, e.** Sporangia; **f.** Columellae; **g.** Hyphal branches showing irregular septate. — Scale bars: **c, d, f.** 5 μm, **e, g.** 10 μm.

*Fungal Names*: FN570832.

*Etymology*: *brunnea* (Lat.) refers to the species having brown colonies.

*Holotype*: HMAS 249876.

*Colonies* on PDA at 27 °C for 9 days, slow growing, reaching 70 mm in diameter, thin pancake-like, concentrically zoned, regular at margin, initially white, becoming Antique Brown to Amber Brown with age, reverse smooth. *Hyphae* branched, aseptate when juvenile, always irregularly septate with age, 2.0–5.0 μm in diameter. *Rhizoids* absent. *Stolons* absent. *Sporangia* globose, pigmented, smooth, 10.5–15.5 μm in diameter, walls deliquescent. *Apophyses* absent. *Collars* absent. *Columellae* hemispherical to cylindrical, smooth, 1.5–2.5 μm long and 2.0–2.5 μm wide. *Projections* absent. *Sporangiospores* subglobose, smooth, 2.0–3.0 μm in diameter. *Chlamydospores* absent. *Zygospores* absent.

*Materials examined*: China. Heilongjiang Province, Yichun, 48°14’11″N, 129°15’45″E, from soil sample, 23 Dec 2017, Mei-Lin Lv (holotype HMAS 249876, living ex-type culture CGMCC 3.16023). Inner Mongolia Auto Region, Hulun Buir, Genhe County, 48°14’11″N, 129°15’45″E, from soil sample, 23 Dec 2017, Mei-Lin Lv and Xiao Ju (living culture CGMCC 3.16024).

*GenBank accession numbers*: MW580596 and MW580597.

*Notes*: *Umbelopsis brunnea* is closely related to *U. globospora* H. Zhao *et al*. based on phylogenetic analysis of ITS rDNA sequences (Fig. 16). However, morphologically *U. globospora* differs from *U. brunnea* by Mouse Gray colonies and degenerated columellae.

**70. *Umbelopsis chlamydospora*** H. Zhao, Y.C. Dai & X.Y. Liu, sp. nov., Fig. 86.

**Fig. 86.**
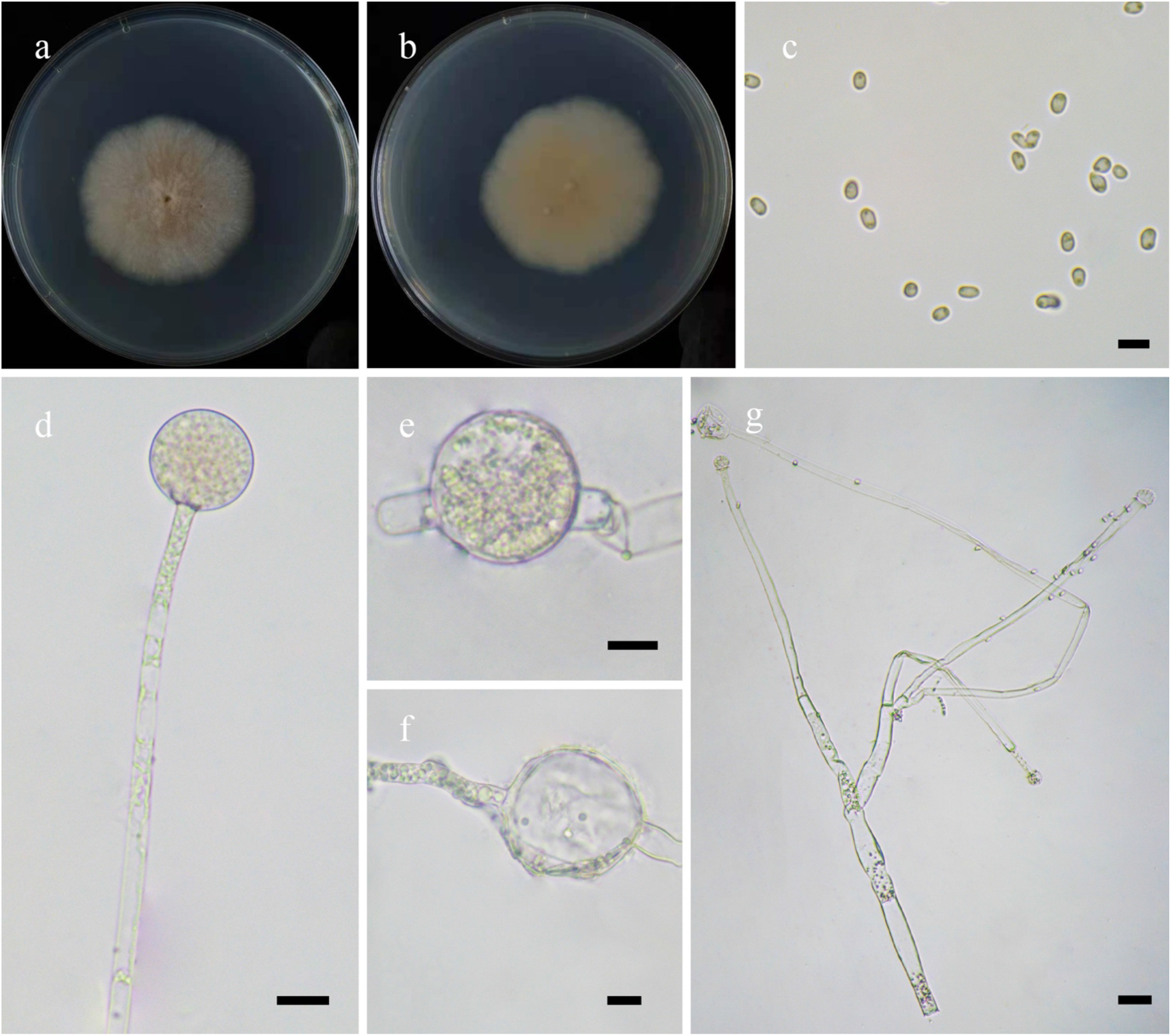
Morphologies of *Umbelopsis chlamydospora* ex-holotype CGMCC 3.16157. **a, b.** Colonies on PDA (**a.** obverse, **b.** reverse); **c.** Sporangiospores; **d.** Sporangia; **e, f.** Chlamydospores; **g.** Sporangiophores. — Scale bars: **c.** 5 μm, **d–e.** 10 μm, **g.** 20 μm.

*Fungal Names*: FN570947.

*Etymology*: *chlamydospora* (Lat.) refers to the species having chlamydospores.

*Holotype*: HMAS 351559.

*Colonies* on PDA at 25 °C for 9 days, slow growing, reaching 45 mm in diameter, white at first, gradually becoming Pale Salmon Color to Seashell Pink, granulate, flat, velvety. *Hyphae* branched, aseptate when juvenile, septate with age, sometimes with swellings, 1.0–4.5 μm in diameter. *Rhizoids* absent. *Stolons* absent. *Sporangiophores* arising from aerial or substrate hyphae, erect, sometimes bent, mainly unbranched and simple branched or sympodial, hyaline, with a septum below sporangium. *Sporangia* globose, pink, smooth, multi-spored, 10.0–16.5 μm in diameter, walls deliquescent. *Apophyses* absent. C*ollars* absent or present. *Columellae* subglobose to globose, smooth, 3.0–8.0 μm long and 3.5–10.0 μm wide. *Sporangiospores* ellipsoid, ovoid, 2.0– 4.0 μm long and 1.5–2.5 μm wide. *Chlamydospores* globose, 30.0–53.5 μm in diameter. *Zygospores* absent.

*Material examined*: China, Beijing, from soil sample, 10 October 2013, Xiao-Yong Liu (holotype HMAS 351559, living ex-holotype culture CGMCC 3.16157).

*GenBank accession number*: OL678224.

*Notes*: *Umbelopsis chlamydospora* is closely related to *U. heterosporus* C.A. de Souza *et al*. based on phylogenetic analysis of ITS rDNA sequences (Fig. 16). However, morphologically *U. heterosporus* differs from *U. chlamydospora* by a faster growth rate (75 mm in diameter for 7 days vs. 45 mm in diameter for 9 days) and variable shape and size of sporangiospores, including ellipsoid 4.5–9 μm long and 1.5–4 μm wide, cylindrical 4.5–8 μm long and 2.5–4 μm wide, reniform, 3.5–7.5 μm long and 2.5–4 μm wide, angular, 2.5–6 μm in diameter, and irregular, 5–11 μm long and 3.5–5 μm wide (Yuan *et al*. 2020).

**71. *Umbelopsis crustacea*** H. Zhao, Y.C. Dai & X.Y. Liu, sp. nov., Fig. 87.

**Fig. 87.**
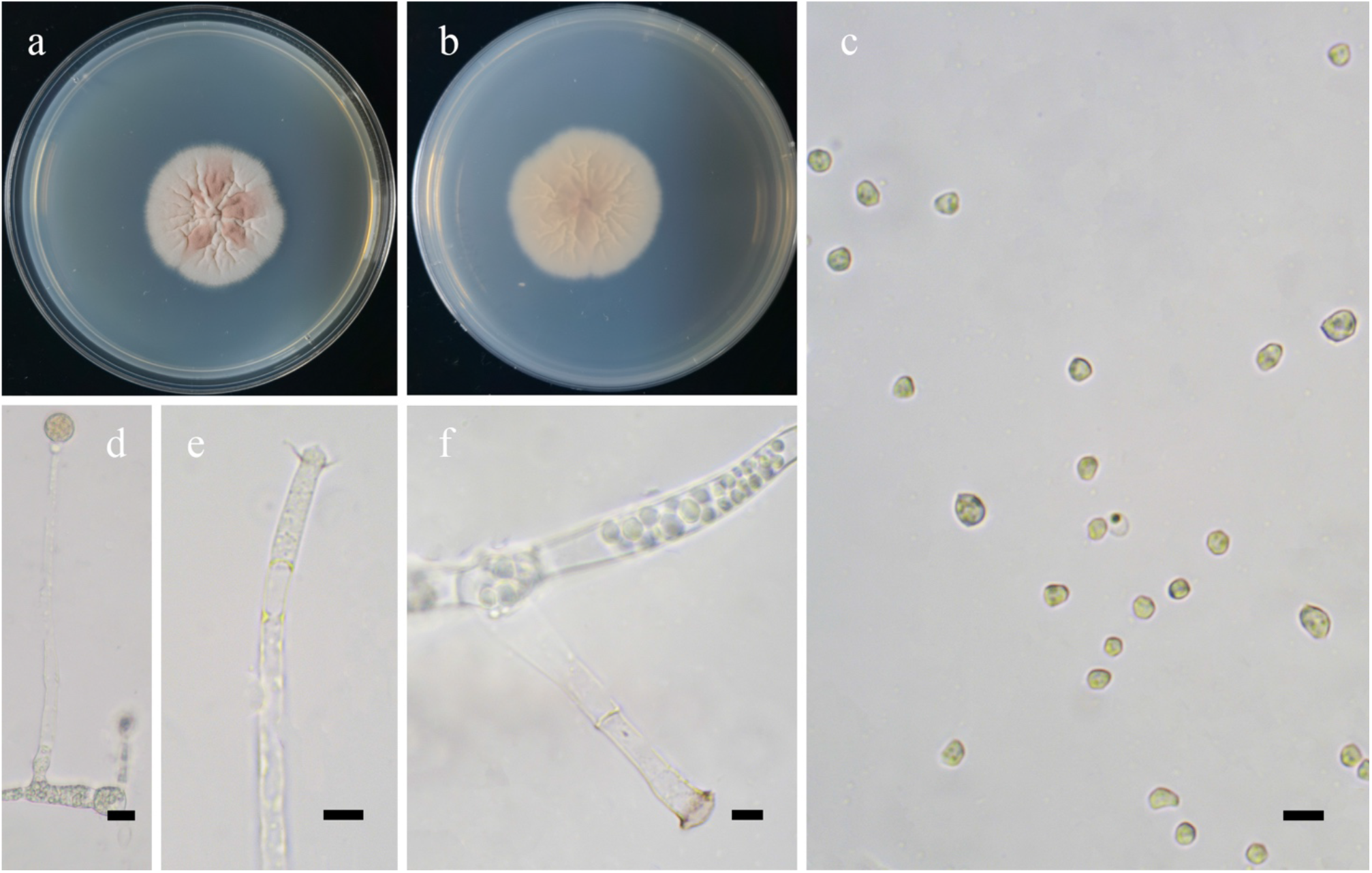
Morphologies of *Umbelopsis crustacea* ex-holotype CGMCC 3.16158. **a, b.** Colonies on PDA (**a.** obverse, **b.** reverse); **c.** Sporangiospores; **d.** Sporangia; **e.** Columellae with collars; **f.** Columellae with septa. — Scale bars: **c, e, f.** 5 μm, **d.** 10 μm.

*Fungal Names*: FN570948.

*Etymology*: *crustacea* (Lat.) refers to the species having a crusty texture to the colonies.

*Holotype*: HMAS 351560.

*Colonies* on PDA at 27°C for 12 days, slow growing, reaching 35 mm in diameter, less than 0.10 mm high, broadly lobed and concentrically zonate, irregular at margin, white at first, gradually becoming La France Pink to Peach Red, granulate, crusty. *Hyphae* simple branched, always swollen, aseptate when juvenile, frequently septate with age, 2.5–10.0 μm in diameter. *Rhizoids* absent. *Stolons* absent. *Sporangiophores* arising from substrate or aerial mycelia, erect, unbranched, simple branched or sympodial, hyaline, sometimes with one septum to several septa below sporangiophores, or aseptate. *Sporangia* globose, hyaline when young, red when old, smooth, multi-spored, 8.0–15.0 μm in diameter, walls deliquescent. *Apophyses* absent. *Collars* present or absent, distinct if present. *Columellae* hemispherical or degenerated, hyaline, subhyaline or pigmented, 1.5–2.5 μm long and 2.5–6.5 μm wide. *Sporangiospores* angular or irregular, subhyaline or hyaline, 2.0–5.0 μm long and 2.0–3.5 μm wide. *Chlamydospores* absent. *Zygospores* unknown.

*Materials examined*: China. Inner Mongolia Auto Region, Arxan, 119°41’16’’E, 47°18’26’’N, from soil sample, 29 January 2018, Yu-Chuan Bai (holotype HMAS 351560, living ex-holotype culture CGMCC 3.16158); Zhalantun,120°56’52′′E, 47°32’23′′N, from soil sample, 29 January 2018, Yu-Chuan Bai (living culture XY03542). Heilongjiang Province, Yichun, 129°23’58′′E, 48°48’21′′N, form soil sample, 28 March 2018, Yu-Chuan Bai (culture XY03944).

*GenBank accession numbers*: OL678225, OL678226 and OL678227.

*Notes*: *Umbelopsis crustacea* is closely related to *U. autotrophica* (E.H. Evans) W. Gams based on phylogenetic analysis of ITS rDNA sequences (Fig. 16). However, morphologically *U. autotrophica* differs from *U. crustacea* by globose sporangiospores (Evans 1971, Meyer & Gams 2003).

**72. *Umbelopsis globospora*** H. Zhao, Y.C. Dai & X.Y. Liu, sp. nov., Fig. 88.

**Fig. 88.**
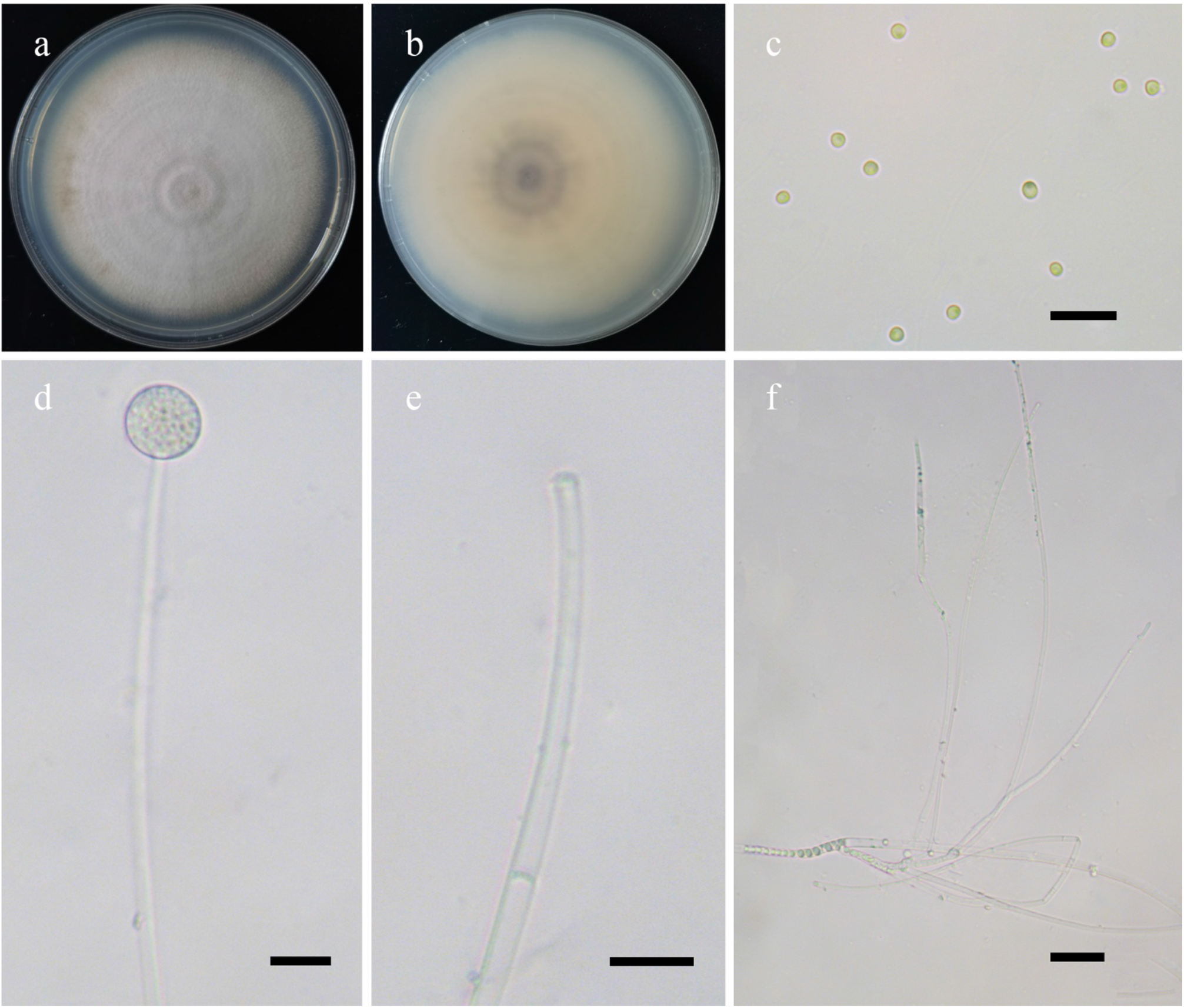
Morphologies of *Umbelopsis globospora* ex-holotype CGMCC 3.16159. **a, b.** Colonies on PDA (**a.** obverse, **b.** reverse); **c.** Sporangiospores; **d.** Sporangia; **e.** Degenerated columellae; **f.** Sporangiophores. — Scale bars: **c–e.** 10 μm, **f.** 5 μm.

*Fungal Names*: FN570949.

*Etymology*: *globospora* (Lat.) refers to the species having globose sporangiospores.

*Holotype*: HMAS 351561.

*Colonies* on PDA at 27°C for 12 days, slow growing, reaching 90 mm in diameter, less than 5 mm high, zonate, white at first, gradually becoming Mouse Gray, granulate, flat, velvety, regular at margin, forming sectors because of the whitish hyphae upon old sporulation layer. *Hyphae* branched, 1.5–5.0 μm in diameter. *Rhizoids* absent. *Stolons* absent. *Sporangiophores* arising from substrate mycelia, erect, sometimes bent, simple branched several times, hyaline, sometimes with one septum on branches of sporangiophores, swollen at the basal part, slightly constricted. *Sporangia* globose, smooth, hyaline when young, pigmented when old, smooth, multi-spored, 8.0– 13.0 μm in diameter, walls deliquescent. *Apophyses* absent. *Collars* absent. *Columellae* degenerated. *Sporangiospores* mostly globose, occasionally subglobose, subhyaline or hyaline, 1.5– 2.5 μm in diameter. *Chlamydospores* absent. *Zygospores* unknown.

*Material examined*: China, Zhejiang Province, Quzhou, Gutian Mountain Nature Reserve, from soil sample, 10 October 2020, Heng Zhao (holotype HMAS 351561, living ex-holotype culture CGMCC 3.16159).

*GenBank accession number*: OL678228.

*Notes*: See notes of *Umbelopsis brunnea*.

**73. *Umbelopsis gutianensis*** H. Zhao, Y.C. Dai & X.Y. Liu, sp. nov., Fig. 89.

**Fig. 89.**
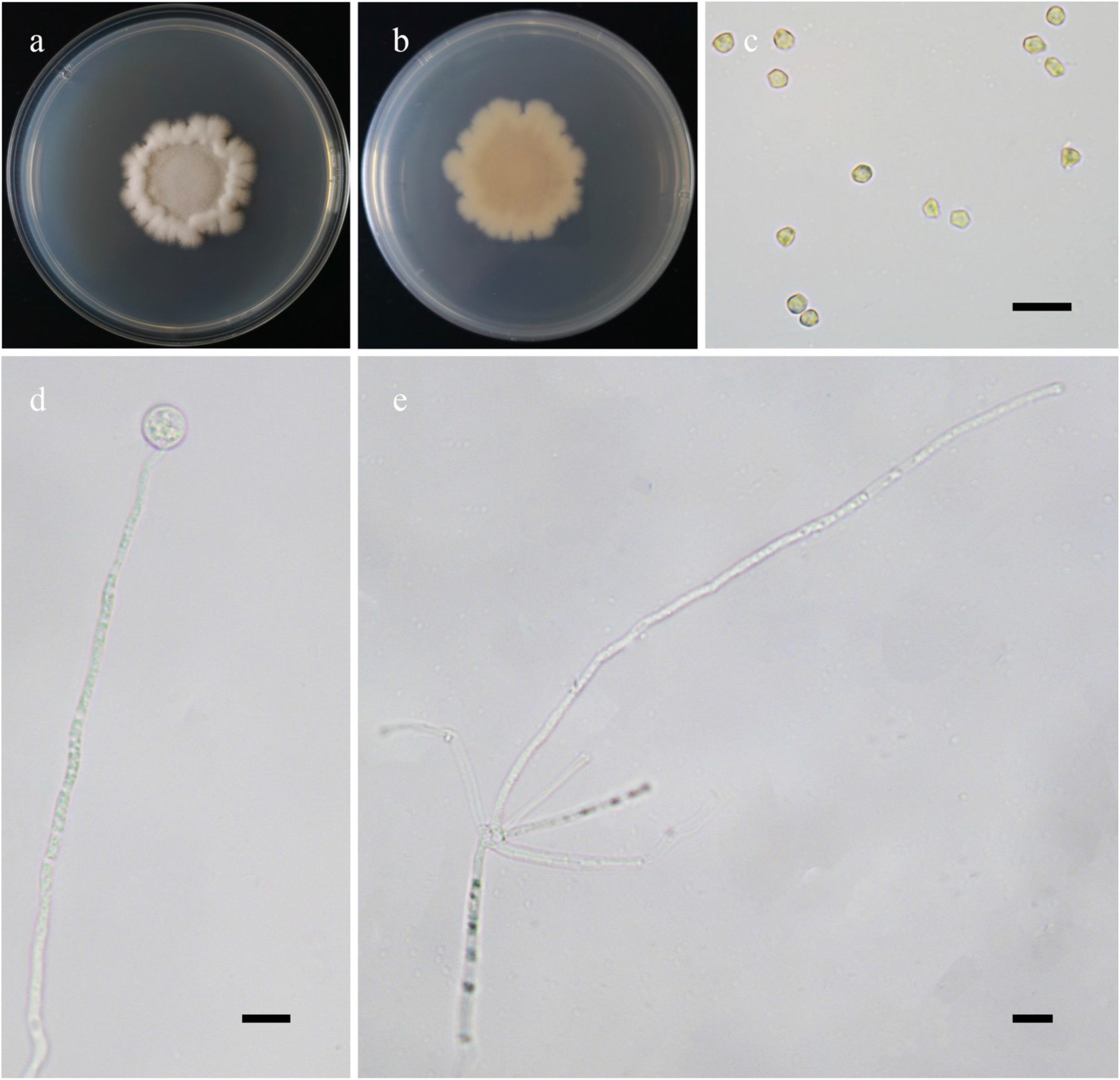
Morphologies of *Umbelopsis gutianensis* ex-holotype CGMCC 3.16160. **a, b.** Colonies on PDA (**a.** obverse, **b.** reverse); **c.** Sporangiospores; **d.** Sporangia; **e.** Sporangiophores. — Scale bars: **c–e.** 10 μm.

*Fungal Names*: FN570950.

*Etymology*: *gutianensis* (Lat.) refers to the Gutian Mountain Nature Reserve, Zhejiang Province, China, where the type was collected.

*Holotype*: HMAS 351562.

*Colonies* on PDA at 27°C for 12 days, slow growing, reaching 40 mm in diameter, broadly lobed and concentrically zonate, irregular at margin, white at first, afterwards becoming Brazil Red at the edge, velvety. *Hyphae* simple branched, aseptate when juvenile, septate with age, 2.5–6.5 μm in diameter. *Rhizoids* absent*. Stolons* absent. *Sporangiophores* arising from substrate mycelia, erect or bent, unbranched or simple branched, hyaline, always swollen on branches. *Sporangia* globose, hyaline when young, red when old, smooth, multi-spored, 8.0–12.5 μm in diameter, walls deliquescent. *Apophyses* absent. C*ollars* absent. *Columellae* degenerated. *Sporangiospores* angular, subhyaline or hyaline, 3.0–4.0 μm long and 2.0–3.0 μm wide. *Chlamydospores* absent. *Zygospores* unknown.

*Materials examined*: China, Zhejiang Province, Quzhou, Gutian Mountain Nature Reserve, from soil sample, 16 March 2021, Heng Zhao (holotype HMAS 351562, living ex-holotype culture CGMCC 3.16160, and living culture XY06604, XY06605).

*GenBank accession numbers*: OL678229, OL678230 and OL678231.

*Notes*: *Umbelopsis gutianensis* is closely related to *U. changbaiensis* Y.N. Wang *et al*. based on phylogenetic analysis of ITS rDNA sequences (Fig. 16). However, morphologically *U. changbaiensis* differs from *U. gutianensis* by depressed globose columellae, and by bearing collars (Wang *et al*. 2014).

**74. *Umbelopsis oblongispora*** H. Zhao, Y.C. Dai & X.Y. Liu, sp. nov., Fig. 90.

**Fig. 90.**
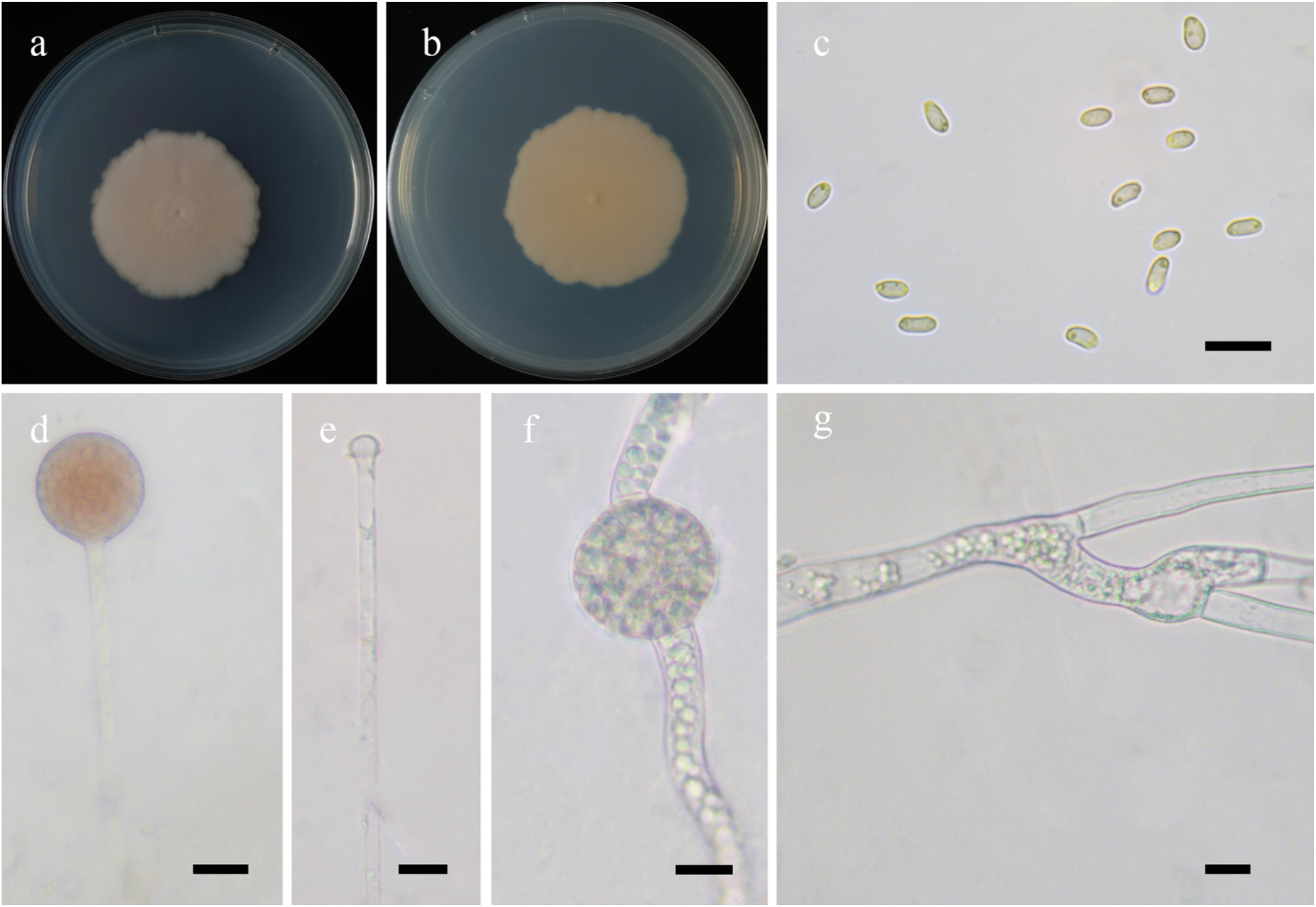
Morphologies of *Umbelopsis oblongispora* ex-holotype CGMCC 3.16161. **a, b.** Colonies on PDA (**a.** obverse, **b.** reverse); **c.** Sporangiospores; **d.** Sporangia; **e.** Columellae with collars; **f.** Chlamydospores; **g.** Sporangiophores. — Scale bars: **c–g.** 10 μm.

*Fungal Names*: FN570951.

*Etymology*: *oblongispora* (Lat.) refers to the species having oblong sporangiospores.

*Holotype*: HMAS 351563.

*Colonies* on PDA at 25 °C for 9 days, slow growing, reaching 45 mm in diameter, white at first, gradually becoming Pale Salmon Color to Seashell Pink, granulate, flat, velvety. *Hyphae* branched, aseptate when juvenile, septate with age, sometimes with swellings, 1.0–4.5 μm in diameter. *Rhizoids* absent. *Stolons* absent. *Sporangiophores* arising from aerial or substrate hyphae, erect, sometimes bent, mainly unbranched and simple branched or sympodial, hyaline, with a septum below sporangium. *Sporangia* globose, pink, smooth, multi-spored, 10.0–16.5 μm in diameter, walls deliquescent. *Apophyses* absent. C*ollars* absent or present. *Columellae* subglobose to globose, smooth, 3.0–8.0 μm long and 3.5–10.0 μm wide. *Sporangiospores* variable, oblong, ellipsoid, cylindrical, reniform, ovoid, or irregular, 2.0–4.0 μm long and 1.5–2.5 μm wide. *Chlamydospores* globose, 30.0–53.5 μm in diameter. *Zygospores* absent.

*Material examined*: China, Zhejiang Province, Quzhou, Gutian Mountain Nature Reserve, from soil sample, 16 March 2021, Heng Zhao (holotype HMAS 351563, living ex-holotype culture CGMCC 3.16161).

*GenBank accession number*: OL678232.

*Notes*: *Umbelopsis oblongispora* is closely related to *U. gibberispora* M. Sugiy. *et al*. based on phylogenetic analysis of ITS rDNA sequences (Fig. 16). However, morphologically *U. gibberispora* differs from *U. oblongispora* by two types of chlamydospores: subglobose to globose macrochlamydospores, and variably shaped microchlamydospores (Sugiyama *et al*. 2003).

**75. *Umbelopsis septata*** H. Zhao, Y.C. Dai & X.Y. Liu, sp. nov., Fig. 91.

**Fig. 91.**
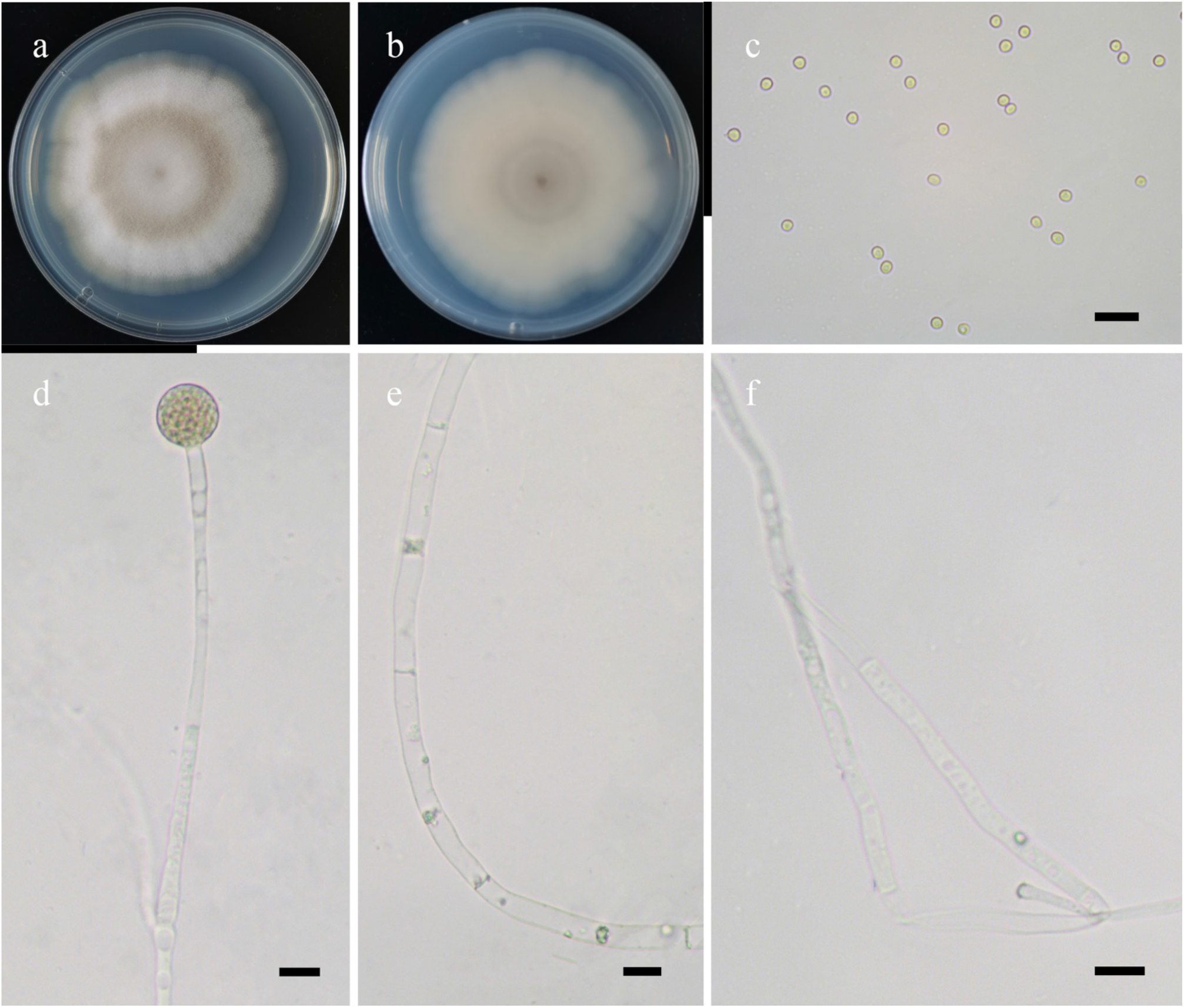
Morphologies of *Umbelopsis septata* ex-holotype CGMCC 3.16162. **a, b.** Colonies on PDA (**a.** obverse, **b.** reverse); **c.** Sporangiospores; **d.** Sporangia; **e.** Hyphae with septa; **f.** Sporangiophores. — Scale bars: **c–f.** 10 μm.

*Fungal Names*: FN570952.

*Etymology*: *septata* (Lat.) refers to the species having hyphae with abundant septa with age.

*Holotype*: HMAS 351564.

*Colonies* on PDA at 27°C for 12 days, slow growing, reaching 70 mm in diameter, less than 5 mm high, white at first, gradually becoming Smoke Gray, granulate, velvety, broadly lobed and concentrically zonate, irregular at margin, forming sectors because of the whitish hyphae upon old sporulation layer. *Hyphae* branched, sometimes swollen, aseptate when juvenile, frequently septate with age, 1.5–8.0 μm in diameter. *Rhizoids* absent*. Stolons* absent. *Sporangiophores* arising from substrate or aerial mycelia, erect, sometimes bent, hyaline, simple branched or unbranched, sometimes with a septum on branches of sporangiophores. *Sporangia* globose, hyaline when young, Light Brown when old, smooth, multi-spored, 11.5–16.5 μm in diameter, walls deliquescent. *Collars* absent. *Apophyses* absent. *Columellae* degenerated. *Sporangiospores* globose to subglobose, subhyaline or hyaline, 2.0–3.0 μm long and 1.5–3.0 μm. *Chlamydospores* absent. *Zygospores* unknown.

*Material examined*: China, Zhejiang Province, Quzhou, Gutian Mountain Nature Reserve, from soil sample, 16 March 2021, Heng Zhao (holotype HMAS 351564, living ex-holotype culture CGMCC 3.16162).

*GenBank accession number*: OL678233.

*Notes*: *Umbelopsis septata* is closely related to *U. globospora* H. Zhao *et al*. based on phylogenetic analysis of ITS rDNA sequences (Fig. 16). However, morphologically *U. globospora* differs from *U. septata* by a faster growth rate at the same condition (90 mm in diameter vs. 70 mm in diameter), and simple or several-time branched sporangiophores.

**76. *Umbelopsis tibetica*** H. Zhao, Y.C. Dai & X.Y. Liu, sp. nov., Fig. 92.

**Fig. 92.**
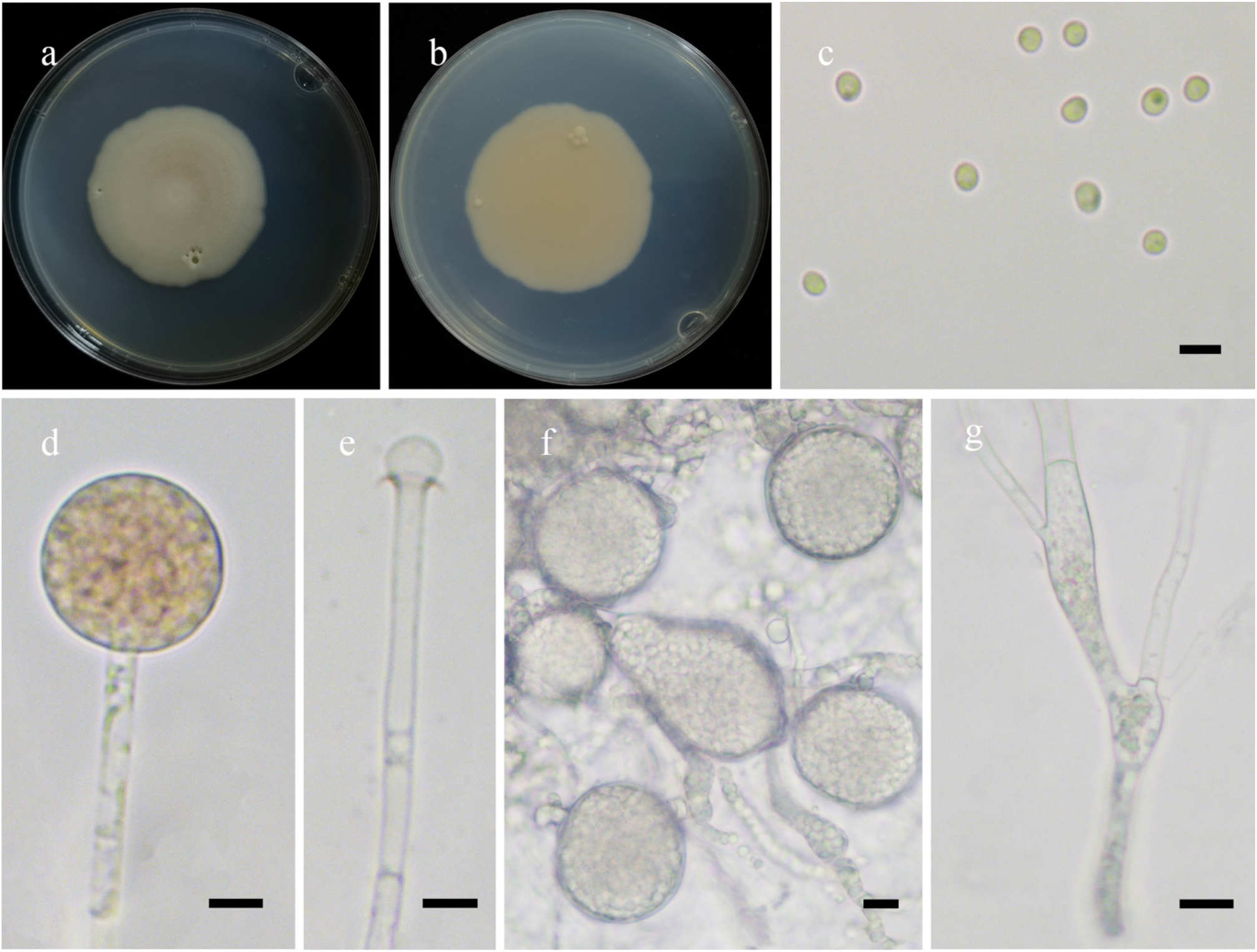
Morphologies of *Umbelopsis tibetica* ex-holotype CGMCC 3.16163. **a, b.** Colonies on PDA (**a.** obverse, **b.** reverse); **c.** Sporangiospores; **d.** Sporangia; **e.** Columellae with collars; **f.** Chlamydospores; **g.** Sporangiophores. — Scale bars: **c.** 5 μm, **d–h.** 10 μm.

*Fungal Names*: FN570953.

*Etymology*: *tibetica* (Lat.) refers to the Tibet Auto Region, China, where the type was collected.

*Holotype*: HMAS 351565.

*Colonies* on PDA at 27 °C for 12 days, slow growing, reaching 45 mm in diameter, thin pancake-like, concentrically zoned, regular at margin, white at first, gradually becoming light red. *Hyphae* abundant, simple branched, aseptate when juvenile, septate with age, 1.5–7.5 μm in diameter, swollen and containing droplets in substrate hyphae. *Rhizoids* absent. *Stolons* absent. *Sporangiophores* arising from substrate hyphae, erect, simple branched, hyaline, always swollen on branches. *Sporangia* globose, hyaline when young, brown when old, smooth, multi-spored, with one septum to several septa below sporangium, 13.0–18.5 μm in diameter, walls deliquescent. *Apophyses* absent. C*ollars* present, distinct. *Columellae* subglobose to ovoid, hyaline, smooth, 4.0– 7.0 μm long and 5.5–9.0 μm wide. *Sporangiospores* always ovoid to globose, rarely irregular, subhyaline or hyaline, 2.0–3.0 μm long and 1.5–2.5 μm wide. *Chlamydospores* in substrate hyphae, hyaline, abundant, irregular when young, globose when old, 26.5–47.5 μm in diameter. *Zygospores* unknow.

*Materials examined*: China, Tibet Auto Region, Linzhi, Bome County, from soil sample, 28 August 2021, Heng Zhao (holotype HMAS 35156, living ex-holotype culture CGMCC 3.16163, and living culture XY07881 and XY07882).

*GenBank accession numbers*: OL678234, OL678235 and OL678236.

*Notes*: *Umbelopsis tibetica* is closely related to *U. wigerinckiae* Sand.-Den. based on phylogenetic analysis of ITS rDNA sequences (Fig. 16). However, morphologically *U. wigerinckiae* differs from *U. tibetica* by umbellately branched sporangiospores, pink, coral to red sporangia, and smaller chlamydospores (5–10 μm in diameter vs. 26.5–47.5 μm in diameter; Crous *et al*. 2017).

### Species new to China

Along with these new species and new combinations above, we found the following 43 species belonging to Mucoromyceta for the first time in China (Table 1).

1. *Absidia caerulea* Bainier [as ‘cærulea’], *Bull. Soc. Bot. Fr.* 36: 184, 1889 *Materials examined*: China, Guangdong Province, Guangzhou, South China Agricultural University, from flower sample, 1 October 2013, Xiao-Yong Liu (living cultures XY00608 and XY00729). *GenBank accession numbers*: OL620081 and OL620082.
2. *Absidia koreana* Hyang B. Lee, Hye W. Lee & T.T. Nguyen, in Ariyawansa *et al*., *Fungal Diversity* 75: 248, 2015 *Materials examined*: China. Fujian Province, Kulangsu, from soil sample, 1 October 2013, Xiao-Yong Liu (living culture XY00816). Hubei Province, Shennongjia National Nature Reserve, from soil sample, 1 October 2013, Xiao-Yong Liu (living culture XY00596). *GenBank accession numbers*: OL620083 and OL620084.
3. *Absidia pararepens* Jurjević, M. Kolařík & Hubka, in Crous *et al*., *Persoonia* 44: 351, 2020 *Materials examined*: China. Beijing, from soil sample, 1 October 2013, Xiao-Yong Liu (living cultures XY00631 and XY00615). Tibet Auto Region, Naqu, from soil sample, 16 Nov 2020, Heng Zhao (Living culture XY05899). *GenBank accession numbers*: OL620085, OL620086 and OL620087.
4. *Actinomortierella ambigua* (B.S. Mehrotra) Vandepol & Bonito, in Vandepol *et al*., *Fungal Diversity*, 2020 *Material examined*: China, Yunnan Province, Dehong, 98°40’3′′E, 24°32’38′′N, from soil sample, 28 May 2021, Heng Zhao (living culture XY06923). *GenBank accession number*: OL620101.
5. *Backusella dispersa* (Hagem) Urquhart & Douch, in Urquhart *et al*., *Persoonia* 46: 16, 2020 *Material examined*: China, Xinjiang Auto Region, Ili, 43°47’4″N, 81°51’55″E, from soil sample, 28 October 2021, Heng Zhao (living culture XY08806). *GenBank accession number*: OL620088.
6. *Backusella locustae* Hyang B. Lee, S.H. Lee & T.T.T. Nguyen, in Wanasinghe *et al*., *Fungal Diversity* 89: 213, 2018 *Material examined*: China, Hunan Province, Changsha, 28°8’4″N, 112°57’19″E, from soil sample, 28 August 2021, Heng Zhao (living culture XY07940). *GenBank accession number*: OL620089.
7. *Backusella oblongielliptica* (H. Nagan., Hirahara & Seshita ex Pidopl. & Milko) Walther & de Hoog, *Persoonia* 30: 41, 2013 *Materials examined*: China. Yunnan Province, Dali, from soil sample, 28 October 2021, Heng Zhao (living culture XY08152). Zhejiang Province, Quzhou, Gutian Mountain Nature Reserve, from soil sample, 27 October 2021, Heng Zhao (living cultures XY08767 and XY08768). *GenBank accession numbers*: OL620090, OL620091 and OL620092.
8. *Backusella tuberculispora* (Schipper) Walther & de Hoog, *Persoonia* 30: 41, 2013 *Materials examined*: China, Yunnan Province, Lincang, Cangyuan County, from soil sample, 22 July 2021, Heng Zhao (living cultures XY07548 and XY7557). *GenBank accession numbers*: OL620093 and OL620094
9. *Cunninghamella antarctica* Caretta & Piont., *Sabouraudia* 15(1): 6, 1977 *Materials examined*: China. Anhui Province, Chuzhou, Quanjiao Country, 32°0’25″N, 117°55’8″E, from soil sample, 28 May 2021, Heng Zhao (living culture XY07003). Gansu Province, Lanzhou, from soil sample, 31 May 2021, Heng Zhao (living culture XY07050). *GenBank accession numbers*: OL620095 and OL620096.
10. *Dissophora globulifera* (O. Rostr.) Vandepol & Bonito, in Vandepol *et al*., *Fungal Diversity*, 104: 279, 2020 *Material examined*: China, Xinjiang Auto Region, Ili, Zhaosu County, 43°13’58″N, 81°10’45″E, from soil sample, 30 October 2021, Heng Zhao (living culture XY08870). *GenBank accession number*: OL620097.
11. *Gongronella namwonensis* Hyang B. Lee, A.L. Santiago & H.J. Lim, in Crous *et al*., *Persoonia* 44: 395, 2020 *Material examined*: China, Zhejiang Province, Hangzhou, from soil sample, 14 August 2021, Heng Zhao (living culture XY08131). *GenBank accession number*: OL620098.
12. *Isomucor trufemiae* J.I. Souza, Pires-Zottar. & Harakava, in Souza *et al*., *Mycologia* 104(1): 236, 2012 *Materials examined*: China, Yunnan Province, Lincang, Cangyuan Country, from soil sample, 22 July 2021, Heng Zhao (living cultures XY07543 and XY7554). *GenBank accession numbers*: OL620099 and OL620100.
13. *Mortierella bainieri* Costantin, *Bull. Soc. Mycol. Fr.* 4(3): 152, 1889 [1888] *Material examined*: China, Yunnan Province, Diqing, Pudacuo National Park, 27°49’40″N, 99°57’45″E, from soil sample, 28 October 2021, Heng Zhao (living culture XY08661). *GenBank accession number*: OL620102.
14. *Mortierella beljakovae* Milko, *Nov. sist. Niz. Rast.* 10: 83, 1973 *Material examined*: China, Heilongjiang Province, Mohe Country, 122°28’47′′E, 52°19’16′′N, from plant debris sample, 27 January 2018, Peng-Cheng Deng (living culture XY03433). *GenBank accession number*: OL620103.
15. *Mortierella chlamydospora* (Chesters) Plaäts-Nit., in Plaäts-Niterink *et al*., *Persoonia* 9(1): 91, 1976 *Materials examined*: China. Zhejiang Province, Wenzhou, Nanyandang Mountain, from soil sample, 22 July 2021, Jia-Jia Chen (living culture XY07448). Guangxi Auto Region, Guigang, from soil sample, 14 August 2021, Heng Zhao (living culture XY08109). *GenBank accession numbers*: OL620104 and OL620105.
16. *Mortierella cystojenkinii* W. Gams & Veenb.-Rijks, *Persoonia* 9(1): 137, 1976 *Materials examined*: China. Heilongjiang Province, Huma Country, 122°48’58′′E, 52°4’23′′N, from plant debris sample, 6 February 2018, Peng-Cheng Deng (living cultures XY03876 and XY03882). Inner Mongolia Auto Region, Zhalantun, 121°39’26′′E, 47°37’45′′N, from soil sample, 27 March 2018, Yu-Chuan Bai (living culture XY03898). *GenBank accession numbers*: OL620106, OL620107 and OL620108.
17. *Mortierella epicladia* W. Gams & Emden, *Persoonia* 9(1): 133, 1976 *Materials examined*: China, Yunnan Province, Dehong, Mangshi, 98°40’3′′E, 24°32’38′′N, from soil sample, 28 May 2021, Heng Zhao (living cultures XY06924 and XY06927). *GenBank accession numbers*: OL620109 and OL620110.
18. *Mortierella fluviae* Hyang B. Lee, K. Voigt & T.T.T. Nguyen, in Hyde *et al*., *Fungal Diversity* 80: 255, 2016 *Material examined*: China, Heilongjiang Province, Heihe, 127°6’54′′E, 48°4’37′′N, from soil sample, 4 April 2018, Ze Liu (living culture XY04370). *GenBank accession number*: OL620111.
19. *Mortierella lignicola* (G.W. Martin) W. Gams & R. Moreau, *Annls Sci. Univ. Besançon*, Sér. 2, Méd. Pharm. 3: 103, 1960 *Material examined*: China, Yunnan Province, Chuxiong, 25°18’53″N, 101°25’17″E, from soil sample, 28 September 2021, Heng Zhao (living culture XY08224). *GenBank accession number*: OL620112
20. *Mortierella longigemmata* Linnem., *Mucorales* (Lehre): 199, 1969 *Materials examined*: China, Yunnan Province. Diqing, Pudacuo National Park, 27°49’40″N, 99°57’45″E, from soil sample, 19 October 2021, Heng Zhao (living cultures XY08599 and XY08601). Yuxi, Ailao Mountain, from soil sample, 25 July 2021, Heng Zhao (living culture XY07567). *GenBank accession numbers*: OL620113, OL620114 and OL620115.
21. *Mortierella nantahalensis* C.Y. Chien, *Mycologia* 63(4): 826, 1971 *Materials examined*: China. Guangxi Auto Region, Fangchenggang, 21°44’46″N, 108°5’13″E, from soil sample, 22 July 2021, Heng Zhao (living culture XY07486), Hainan Province, Limu Mountain, from soil sample, 30 December 2021, Jia-Jia Chen (living culture XY06017). *GenBank accession numbers*: OL620116 and OL620117.
22. *Mortierella rishikesha* B.S. Mehrotra & B.R. Mehrotra, *Zentbl. Bakt. ParasitKde*, Abt. II 118: 184, 1964 [1963] *Material examined*: China, Beijing, 39°45’49″N, 115°36’34″E, from soil sample, 9 July 2021, Heng Zhao (living culture XY07315). *GenBank accession number*: OL620118.
23. *Mortierella schmuckeri* Linnem., *Arch. Mikrobiol.* 30: 263, 1958 *Materials examined*: China, Beijing, 39°52’11″N, 115°59’46″E, from soil sample, 12 July 2021, Heng Zhao (living culture XY07412); 40°27’37″N, 117°1’14″E, from soil sample, 13 July 2021, Heng Zhao (living culture XY07419). *GenBank accession numbers*: OL620119 and OL620120.
24. *Mortierella sclerotiella* Milko, *Nov. sist. Niz. Rast.*4: 160, 1967 *Material examined*: China, Inner Mongolia Auto Region, Arxan, 119°41’16′′E, 47°18’28′′N, from soil sample, 29 January 2018, Yu-Chuan Bai (living culture XY03528). *GenBank accession number*: OL620121.
25. *Mortierella turficola* Y. Ling, Rev. gén. Bot. 42: 743, 1930 *Material examined*: China, Heilongjiang Province, Mohe Country, 122°28’47′′E, 52°19’16′′N, from plant debris sample, 21 January 2018, Peng-Cheng Deng (living culture XY03346). *GenBank accession number*: OL620122.
26. *Mucor aligarensis* B.S. Mehrotra & B.R. Mehrotra, *Sydowia* 23(1‒6): 183, 1970 [1969] *Material examined*: China, Yunnan Province, Yuxi, Ailao Mountain, from soil sample, 25 July 2021, Heng Zhao (living culture XY07583). *GenBank accession number*: OL620123.
27. *Mucor atramentarius* L. Wagner & G. Walther, in Wagner *et al*., *Persoonia* 44: 87, 2019 *Material examined*: China, Tibet Auto Region, Ngari, Coqen County, from soil sample, 28 May 2021, Heng Zhao (living culture XY06751). *GenBank accession number*: OL620124.
28. *Mucor bainieri* B.S. Mehrotra & Baijal, *Aliso* 5(3): 237, 1963 *Material examined*: China, Yunnan Province, Dali, 26°24’45″N, 99°53’41″E, from soil sample, 28 October 2021, Heng Zhao (living culture XY08747). *GenBank accession number*: OL620125.
29. *Mucor fuscus* (Berk. & M.A. Curtis) Berl. & De Toni, in Berlese *et al*., *Syll. Fung*. (Abellini) 7(1): 204, 1888 *Materials examined*: China, Tibet Auto Region, Lhasa, Damxung County, from soil sample, 28 May 2021, Heng Zhao (living cultures XY06718 and XY06719). *GenBank accession numbers*: OL620126 and OL620127.
30. *Mucor fluvii* Hyang B. Lee, S.H. Lee & T.T.T. Nguyen, [as ‘fluvius’], in Wanasinghe *et al*., *Fungal Diversity* 89: 220, 2018 *Materials examined*: China, Zhejiang Province, Wenzhou, Nanyandang Mountain, from soil sample, 22 July 2021, Jia-Jia Chen (living cultures XY07446 and XY07447). *GenBank accession numbers*: OL620128 and OL620129.
31. *Mucor griseocyanus* Hagem, *Skr. VidenskSelsk. Christiania*, Kl. I, Math.-Natur.(no. 7): 28, 1908 *Materials examined*: China, Tibet Auto Region, Linzhi, Gongbujiangda County, from soil sample, 28 May 2021, Heng Zhao (living cultures XY06685 and XY06697). *GenBank accession numbers*: OL620130 and OL620131.
32. *Mucor lusitanicus* Bruderl., *Bull. Soc. Bot. Genève*, 2 sér. 8: 276, 1916 *Materials examined*: China. Beijing, 39°47’49″N, 115°34’1″E, from dung sample, 6 July 2021, Heng Zhao (living cultures XY07298 and XY07299). Shandong Province, Dongying, 37°45’23″N, 118°29’22″E, from soil sample, 30 December 2020, Heng Zhao (living cultures XY06042 and XY06074). *GenBank accession numbers*: OL620132, OL620133, OL620134 and OL620135.
33. *Mucor minutus* (Baijal & B.S. Mehrotra) Schipper, *Stud. Mycol.* 10: 24, 1975 *Materials examined*: China. Hubei Province, Tianmenshan, form soil sample, 22 July 2021, Heng Zhao (living culture XY07460). Yunnan Province, Dehong, Mangshi, 98°40’3′′E, 24°32’38′′N, from soil sample, 28 May 2021, Heng Zhao (living cultures XY06939 and XY06940). *GenBank accession numbers*: OL620136, OL620137 and OL620138.
34. *Mucor mucedo* Fresen., *Beitr*. *Mycol.* 1: 7, 1850 *Materials examined*: China, Beijing, 39°46’39″N, 115°28’57″E, from soil sample, 6 July 2021, Heng Zhao (living cultures XY07301 and XY07302). *GenBank accession numbers*: OL620139 and OL620140.
35. *Mucor nanus* Schipper & Samson, *Mycotaxon* 50: 477, 1994 *Materials examined*: China. Beijing, 39°30’19″N, 115°58’28″E, from soil sample, 6 July 2021, Heng Zhao (living culture XY07221). Hubei Province, Shiyan, Danjiangkou, 39°46’39″N, 111°8’16″E, from soil sample, 31 May 2021, Heng Zhao (living culture XY07037). *GenBank accession numbers*: OL620141 and OL620142.
36. *Mucor nidicola* A.A Madden, Stchigel, Guarro, Deanna A. Sutton & Starks, *Int. J. Syst. Evol. Microbiol.* 62(7): 1712, 2012 *Material examined*: China, Yunnan Province, Dali, 25°42’6″N, 100°9’19″E, from soil sample, 28 October 2021, Heng Zhao (living culture XY08161). *GenBank accession number*: OL620143.
37. *Mucor pseudocircinelloides* L. Wagner & G. Walther, in Wagner *et al*., *Persoonia* 44: 91, 2019 *Materials examined*: China, Shanxi Province, Shuozhou, Youyu Country, 10°2’40″N, 112°35’7″E, from soil sample, 1 August 2021, Heng Zhao (living cultures XY07713 and XY07714). *GenBank accession numbers*: OL620144 and OL620145.
38. *Mucor silvaticus* Hagem, *Skr. VidenskSelsk*. Christiania, Kl. I, Math.-Natur. (no. 7): 31, 1908 *Materials examined*: China, Inner Mongolia Auto Region, Hulun Buir, Genhe County, 122°15’48′′E, 52°8’22′′N, from plant debris sample, 4 April 2018, Peng-Cheng Deng (living cultures XY04328 and XY04329). *GenBank accession numbers*: OL620146 and OL620147.
39. *Mucor stercorarius* Hyang B. Lee, P.M. Kirk & T.T.T. Nguyen, in Tibpromma *et al*., *Fungal Diversity* 83: 110, 2017 *Material examined*: China, Anhui Province, Hefei, Lu’an, 116°0’28′′E, 31°45’27′′N, from soil sample, 28 May 2021, Heng Zhao (living culture XY07007). *GenBank accession number*: OL620148.
40. *Mucor variicolumellatus* L. Wagner & G. Walther, in Wagner *et al*., *Persoonia* 44: 92, 2019 *Materials examined*: China, Beijing, 40°29’46″N, 117°4’5″E, from rotten tree sample, 6 July 2021, Heng Zhao (living cultures XY07369 and XY07370). *GenBank accession numbers*: OL620149 and OL620150.
41. *Umbelopsis gibberispora* M. Sugiy., Tokum. & W. Gams, *Mycoscience* 44(3): 220, 2003 *Material examined*: China, Zhejiang Province, Quzhou, Gutian Mountain Nature Reserve, from soil sample, 28 May 2021, Heng Zhao (living culture XY06667). *GenBank accession number*: OL620151.
42. *Umbelopsis ovata* (H.Y. Yip) H.Y. Yip, *Trans. Br. Mycol. Soc.* 86(2): 334, 1986 *Materials examined*: China, Heilongjiang Province, Muhe Country, 122°38’47′′E, 52°40’1′′N, from plant debris sample, 17 April 2018, Peng-Cheng Deng (living cultures XY03175 and XY03176). *GenBank accession numbers*: OL620152 and OL620153.
43. *Umbelopsis sinsidoensis* Hyang B. Lee & T.T.T. Nguyen, in Wanasinghe *et al*., *Fungal Diversity* 89: 223, 2018 *Material examined*: China, Zhejiang Province, Quzhou, Gutian Mountain Nature Reserve, from soil sample, 25 January 2021, Heng Zhao (living culture XY06194). *GenBank accession number*: OL620154

## Discussion

Since Hibbett *et al*. (2007) abolished the phylum Zygomycota, the phylogeny of early diverging fungi has been undergoing drastic changes. Spatafora *et al*. (2016) proposed a phylum-level framework based on genomic data, accepting two phyla and six subordinate subphyla, Mucoromycota (Glomeromycotina, Mortierellomycotina and Mucoromycotina) and Zoopagomycota (Entomophthoromycotina, Kickxellomycatina and Zoopagomycotina). Depending on the phylogeny of 18S and 28S rDNA sequences, these phyla and subphyla were raised to subkingdoms (Mucoromyceta, Zoopagomyceta) and phyla (Entomophthoromycota, Glomeromycota, Kickxellomycota, Mortierellomycota, Mucoromycota and Zoopagomycota), respectively, along with a novel phylum Calcarisporiellomycota being erected (Tedersoo *et al*. 2018). In this paper, we sharpened our focus on the subkingdom Mucoromyceta, and looked further into its inner relationship by a time-scaled phylogenetic tree in concert with the monophyly concept. The results suggest that the divergence time of phyla in Mucoromyceta is earlier than 639 Mya, falling in the time interval revealed in other studies (Table 4; Gueidan *et al*. 2011, Chang *et al*. 2015, Tedersoo *et al*. 2018). Consequently, two new phyla are proposed (Fig. 2, 3, 4), namely Umbelopsidomycota and Endogonomycota. The former morphologically differs from Mucoromycota and Mortierellomycota by degenerated aerial hyphae and a large quantity of substrate-borne sporangiophores which produce a thin layer of colonies, and by abundant pigmented sporangia which cause reddish or ochraceous colonies, as well as a frequently occurring sectoring due to a possible genetic instability (Meyer & Gams 2003, Wang *et al*. 2013, 2014, 2015). Besides, it is physiologically different from its allied phylum Mortierellomycota by the inability to synthesize arachidonic acid (Zhao *et al*. 2021a). The later phylum Endogonomycota differs from all other phyla in the subkingdom Mucoromyceta by forming ectomycorrhizae with plants, by producing sexual spores (zygospores) in sporocarps rather than zygosporangia, and by producing asexual chlamydospores but not sporangiospores (Yao *et al*. 1996, Desirò *et al*. 2017, Yamamoto *et al*. 2020).

Some secondary ranks were validly introduced in the subkingdom Mucoromyceta, including subphyla, viz., Glomeromycotina, Mucoromycotina, Kickxellomycotina, Mortierellomycotina and Calcarisporiellomycotina (Hibbett *et al*. 2007, Hoffmann *et al*. 2011, Spatafora *et al*. 2016, Tedersoo *et al*. 2018), and subfamilies, viz., Absidioideae, Chaetocladioideae, Choanephoroideae, Cokeromycetoideae, Cunninghamelloideae, Dichotomocladioideae, Dicranophoroideae, Gilbertelloideae, Lichtheimioideae, Mycotyphoideae, Rhizomucoroideae, Kirkomycetoideae, Mucoroideae and Thamnidioideae (Voigt & Kirk 2015). In this taxonomic hierarchy of six phyla of the subkingdom Mucoromyceta, viz., Calcarisporiellomycota, Endogonomycota, Glomeromycota, Mortierellomycota, Mucoromycota and Umbelopsidomycota, we followed the principal ranks of taxa, without dealing with any secondary ranks.

In basal fungi, studies of divergence times have focused mainly on ranks above class (Chang *et al*. 2015, Tedersoo *et al*. 2018). For the first time, taking the subkingdom Mucoromyceta as a case, we established a minimum age range of earlier than 639 Mya, earlier than 585 Mya, 651–400 Mya and 570–107 Mya, for phyla, classes, orders, and families, respectively (Fig. 4, Table 4 and 5). The results suggest that Mucoromyceta contains six phyla (two new phyla, Endogonomycota and Umbelopsidomycota), eight classes, 15 orders (five orders, Claroideoglomerales, Cunninghamellales, Lentamycetales, Phycomycetales and Syncephalastrales), 41 families (six new families, Circinellaceae, Gongronellaceae, Protomycocladaceae, Rhizomucoraceae, Syzygitaceae and Thermomucoraceae) and 121 genera. In details, 1) Calcarisporiellomycota comprises one class, one order, one family and two genera; 2) Endogonomycota includes one class, one order, two families and seven genera; 3) Glomeromycota diverges into three classes, six orders, 16 families and 41 genera; 4) Mortierellomycota accommodates one class, one family and 14 genera; 5) Mucoromycota contains one classes, five orders, 19 families and 55 genera, including five new orders, six new families and one revived family; 6) Umbelopsidomycota consists of one class, one order, two families and two genera. In addition, most families of Mucoromycota, such as Backusellaceae, Lentamycetaceae, Phycomycetaceae, Pilobolaceae, Radiomycetaceae, Rhizopodaceae and Saksenaeaceae were phylogenetically and morphologically well-supported, by other authors (von Arx 1982, Benny *et al*. 2001, O’Donnell *et al*. 2001, Voigt & Wöstemeyer 2001, Hoffmann *et al*. 2013). This updated classification is vigorously supported by molecular dating analyses, except for the Choanephoraceae and Mycotyphaceae which are yet unique in traditional morphological characteristics (Benny *et al*. 1985, Benny 1991, Hoffmann *et al*. 2013).

In this study, we accepted the Glomeromycota as the phylum rank (Schüßler *et al*. 2001, Tedersoo *et al*. 2018), rather than the subphylum erected by Spatafora *et al*. (2016). And Glomeromycota divided into three classes, five orders and 14 families by Oehl *et al*. (2011a, b, c). In this study, the phylogenetic analyses and divergence time analyses suggest that its class, order and family ranks are well supported, while the family Claroideoglomeraceae is raised up to a newly order Claroideoglomerales in the Glomeromycetes. Moreover, the divergence time of higher taxa of Glomeromycota provided in our study (Fig. 4 and Table 4), class rank: mean stem divergence time earlier than 585 Ma, order rank: mean stem divergence time earlier than 400 Ma and family rank: mean stem divergence time during 107–539 Ma.

In the past decades, a great step has been made for the phylogenetic relationship within Mucorales and Mortierellales based on morphology, multi-loci and genomic phylogeny (Gams 1976, 1977, Zheng *et al*. 2007, Hoffmann *et al*. 2009a, 2013, Wagner *et al*. 2013, Walther *et al*. 2013, Vandepol *et al*. 2020). However, the results of the MCC tree (Fig. 4) suggest the genus *Mucor* is polyphyletic. For instance, *Mucor abundans*, *M. azygosporus*, *M. indicus*, *M. luteus*, *M. mucedo* and *M. silvaticus* are located in different clades, causing difficulties in defining the boundaries of the genus. This implies that it is necessary to resolve the polyphyletic genus into different genera including new monophyletic genera (Wanger *et al*. 2013). A perfect example was to re-circumscribe the genera of *Absidia*, *Lentamyces* and *Lichtheimia* based on a combination of physiological, phylogenetic and morphological characters (Hoffmann *et al*. 2007, Hoffmann and Voigt 2009, Hoffmann 2010). Another example is that of Vandepol *et al*. (2020), where 14 genera were accepted in Mortierellaceae by multiple genes recalled from genomics with some new genera being established to resolve the polyphyly. Although the ITS rDNA was recommended as the fungal DNA barcode (Schoch *et al*. 2012, Nilsson *et al*. 2014), it evolved too fast (White *et al*. 1990) to be aligned across families and higher taxa. Therefore, we strongly suggest that multiple markers, i.e., physiological, phylogenetic, phylogenomic, morphological data and divergence time, should be considered comprehensively as much as possible when dealing with taxa at the generic level for basal or early diverging fungi.

Other eight genera with an uncertain taxonomic position in the Catalogue of Life or Index Fungorum are classified into definite families in this study. In detail, *Ambomucor* is assigned to the family Mucoraceae, *Densospora* is assigned to the family Densosporaceae, *Echinochlamydosporium* is assigned to the family Calcarisporiellaceae, *Intraornatospora* and *Paradentiscutata* are assigned to the family Gigasporaceae, *Modicella* is assigned to the family Mortierellaceae, and a new family Syzygitaceae was proposed to accommodate the genera *Sporodiniella* and *Syzygites*.

Currently, it is a common requirement to delineate fungal species using genealogical concordance and phylogenetic species recognition (GCPSR; Taylor *et al*. 2000, Hibbett & Taylor 2013). Species in major genera such as *Absidia* (Hurdeal *et al*. 2021, Zong *et al*. 2021), *Apophysomyces* (Alvarez *et al*. 2010), *Backusella* (Urquhart *et al*. 2021, Nguyen *et al*. 2021), *Cunninghamella* (Zheng & Chen 2001) and *Rhizopus* (Zheng *et al*. 2007, Liu *et al*. 2008) were characterized by unique morphological features and molecular data, making it easy to add new species. However, early morphological studies left a number of species complexes and cryptic species, especially in the species-rich genera like *Mortierella* and *Mucor* (Petkovits *et al*. 2011, Hoffmann *et al*. 2013, Wagner *et al*. 2013, 2020). This problem hinders studies on the diversity of basal fungal species. An important progress has been made in resolving the *Mucor circinelloides* complex based on five genes, phenotypic features, mating behaviors and maximum growth temperatures (Schipper 1976, Wagner *et al*. 2020). In brief, we are looking forward to better solving the species identification and definition in the subkingdom Mucoromyceta, providing a solid foundation for further studies.

Although studies on Chinese Mucoromyceta began relatively late, there are currently 308 species recorded (Fig. 1 and Table S1), accounting for nearly 40 % of the 792 species recorded worldwide (https://nmdc.cn/fungarium/fungi/chinadirectories, November 12, 2021). However, 46 type strains originated from China only (less than 6 %, Table 6), far less than those in the USA (126) and close to those in India (51), Brazil (46), and Germany (44). In mainland China, Prof. Zheng’s team has long been devoted to the systematic study on Mucoromyceta, particularly *Absidia* (Zhao *et al*. 2021b, 2022a, b, Zong *et al*. 2021), *Actinomucor* (Zheng & Liu 2005), *Ambomucor* (Zheng & Liu 2014, Liu & Zheng 2015), *Backusella* (Zheng *et al*. 2013), *Blakeslea* (Zheng & Chen 1986), *Circinella* (Zheng *et al*. 2017), *Cunninghamella* (Liu *et al*. 2001, Zheng & Chen 2001, Zhao *et al*. 2021b,), *Mortierella* (Chen 1992a, b), *Mucor* (Pei 2000a, b, Zheng 2002), *Pilaira* (Liu *et al*. 2012, Zheng & Liu 2009), *Pilobolus* (Hu *et al*. 1989), *Pirella* (Liu 2004), *Rhizomucor* (Zheng *et al*. 2009), *Rhizopus* (Zheng *et al*. 2007, Liu *et al*. 2008), *Syncephalastrum* (Zheng *et al*. 1988), and *Umbelopsis* (Wang *et al*. 2013, 2014, 2015). In Taiwan of China, Prof. Ho’s team also contributed a large number of species in Mucoromyceta, including *Absidia* (Ho *et al*. 2004, Hsu *et al*. 2009, Hsu & Ho 2010), *Blakeslea* (Ho & Chen 2003), *Circinella* (Ho 1995a), *Chaetocladium* (Ho *et al*. 2008), *Gongronella* (Ho & Chen 1990), *Lichtheimia* (Yang *et al*. 2012), *Rhizopus* (Ho 1988, 1995b, Ho & Chen 1994), and *Thamnidium* (Ho 2002). In the current study, we further explored the Mucoromyceta in China and found 73 species new to science and 43 species new to China. Our results increase the diversity of basal fungi in China and the world as well. To sum up with these taxa proposed herein, a total of six phyla, eight classes, 15 orders, 41 families, 121 genera an 868 species belonging to Mucoromyceta in the world are outlined. Specifically, six phyla, seven classes, 13 orders, 32 families, 56 genera and 426 species are recorded in China. We believe more taxa in Mucoromyceta will be described from China when sampling from different ecosystems, as has occurred in the Ascomycota and Basidiomycota (Wu *et al*. 2020, 2022, Zheng *et al*. 2020, Dai *et al*. 2021, Wang *et al*. 2021b, Liu *et al*. 2022).

**Table 5.** A comparison of estimated fossil age and inferred divergence time of the five phyla in the subkingdom Mucoromyceta in different studies. * New taxa proposed in the present study are in bold.

| Phyla | Inferred mean of stem age (Mya) | References |
| --- | --- | --- |
| Calcarisporiellomycota | **748** | **This study** |
|  | 626 | (Tedersoo *et al.* 2018) |
| **Endogonomycota\*** | **708** | **This study** |
| Glomeromycota | **810** | **This study** |
|  | 610 | (Redecker *et al.* 2000) |
|  | 590 | (Berbee & Taylor 2001) |
|  | 725 | (Berney & Pawlowski 2006) |
|  | 720 | (Lucking *et al.* 2009) |
|  | 510 | (Gueidan *et al.* 2011) |
|  | 520 | (Gaya *et al.* 2015) |
|  | 511 | (Chang *et al.* 2015) |
|  | 642 | (Tedersoo *et al.* 2018) |
| Mortierellomycota | **777** | **This study** |
|  | 727 | (Tedersoo *et al.* 2018) |
|  | 511 | (Chang *et al.* 2015) |
| Mucoromycota | **639** | **This study** |
|  | 610 | (Berbee & Taylor 2001) |
|  | 510 | (Gueidan *et al.* 2011) |
|  | 520 | (Gaya *et al.* 2015) |
|  | 578 | (Chang *et al.* 2015) |
|  | 626 | (Tedersoo *et al.* 2018) |
| Umbelopsidomycota* | **694** | **This study** |

**Table 6.**
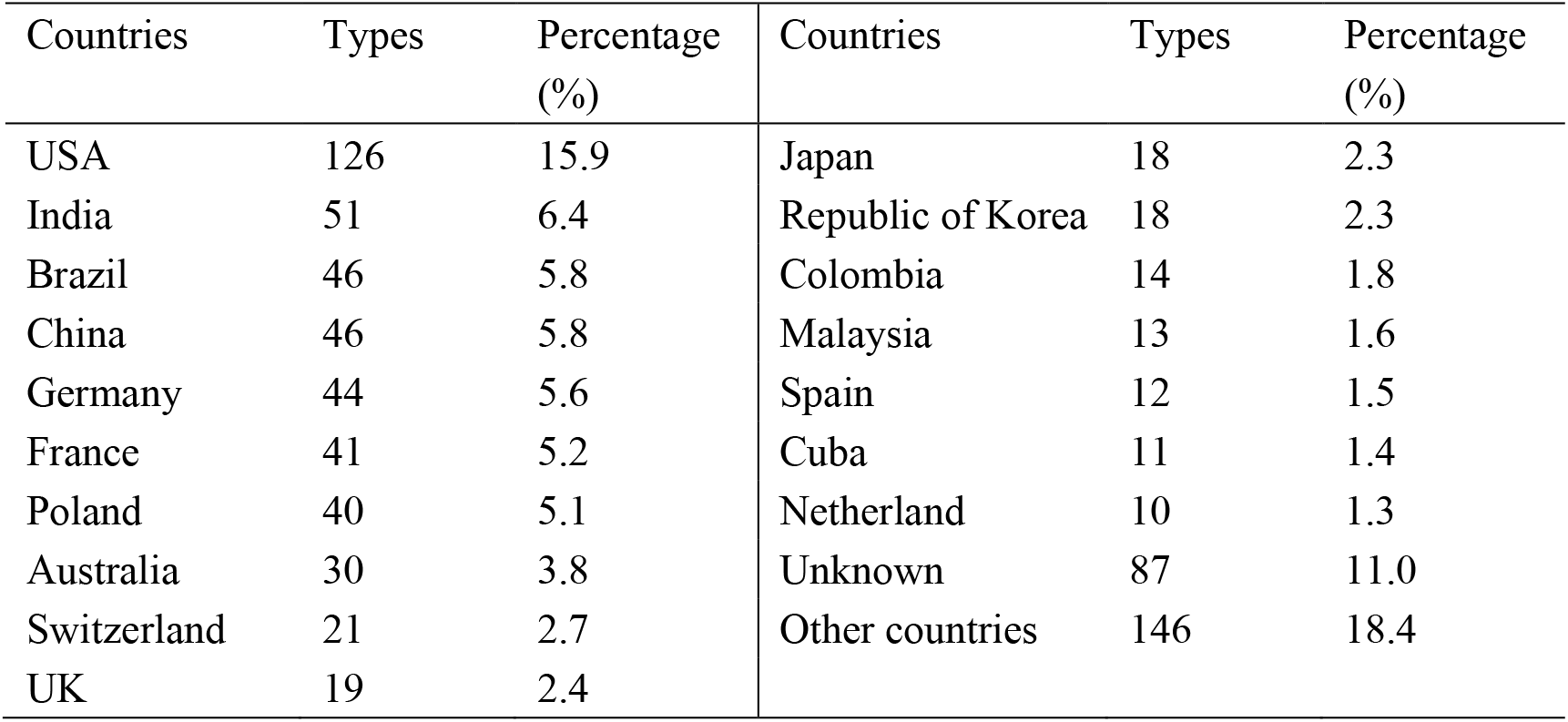
The origin of taxonomic types in Mucoromyceta

| Countries | Types | Percentage (%) | Countries | Types | Percentage (%) |
| --- | --- | --- | --- | --- | --- |
| USA | 126 | 15.9 | Japan | 18 | 2.3 |
| India | 51 | 6.4 | Republic of Korea | 18 | 2.3 |
| Brazil | 46 | 5.8 | Colombia | 14 | 1.8 |
| China | 46 | 5.8 | Malaysia | 13 | 1.6 |
| Germany | 44 | 5.6 | Spain | 12 | 1.5 |
| France | 41 | 5.2 | Cuba | 11 | 1.4 |
| Poland | 40 | 5.1 | Netherland | 10 | 1.3 |
| Australia | 30 | 3.8 | Unknown | 87 | 11.0 |
| Switzerland | 21 | 2.7 | Other countries | 146 | 18.4 |
| UK | 19 | 2.4 |  |  |  |

## Acknowledgments

We thank the following scholars for their help with sampling and culture isolation: Yu-Chuan Bai and Peng-Cheng Deng (North Minzu University), Meng Zhou, Hong-Ming Zhou, Shun Liu, Xiao-Wu Man, Chang-Ge Song and Tai-Min Xu (Beijing Forestry University), Ya-Ning Wang, Jing Yang, Mao-Qiang He, Yue Zhang, Jie Li, Xiao-Ling Liu, Gui-Qing Chen, Feng-Yan Bai, Xue-Wei Wang (Prof.), Xue-Wei Wang (Ph.D. student), Xin-Yu Zhu and Long Wang (Institute of Microbiology, Chinese Academy of Chinese), Xiao Ju (Nanjing Agricultural University), Zhi-Kang Zhang (Ludong University), Ze Liu, Xiang Sun and Qi-Ming Wang (Hebei University), Zhong-Dong Yu (Northwest A&F University), Yan-Yan Long (Guangxi Academy of Agricultural Sciences), Nian-Kai Zeng (Hainan Medical University), Ze-Fen Yu and Min Qiao (Yunnan University), Pei-Wu Cui (Hunan University of Chinese Medicine), Jun-Feng Liang (Chinese Academy of Forestry), Yan-Feng Hang (Guizhou University), Wang-Qiu Deng (Guangdong Institute of Microbiology), Pu Liu, Li-Ying Ren (Jilin Agricultural University), Biao Xu (Tarim University), Rafael F. Castañeda-Ruiz (Academia de Ciencias de Cuba, Cuba), Kerstin Voigt (Friedrich Schiller University Jena, Germany), Manfred Binder (Clark University, USA), and Timothy Y. James (Michigan University, USA), Nicolas Corradi and Mathu Malar C (University of Ottawa, USA), and Francis Michel Martin (Head of Department at French National Institute for Agriculture, Food, and Environment, France).

## Author contributions

Heng Zhao is responsible for sample collection, culture isolations, identifications and descriptions, data analyses, drafting and editing the manuscript; Yu-Cheng Dai and Xiao-Yong Liu for funding acquisition, conceiving this study, and improving the manuscript.

## Data availability

All sequences have been deposited at GenBank database. All the trees have been deposited at TreeBase (Added when revised).

## Compliance with ethical standards

Conflict of interest: All authors declare no conflict of interest.

Consent to participate: All authors agreed to participate in this research.

Consent for publication: All authors read and approved submitted manuscript.

---

**Table S1.** Species publication years, types origin, and record time in China.

