## Supplementary material for "Outline and divergence time of subkingdom Mucoromyceta: two new phyla, five new orders, six new families and seventy-three new species": Table S1

**Table S1.** Species publication years, types origin, and record time in China

| Name Authors | Published year | Type locality | Distributed in China | First recorded in China |
| --- | --- | --- | --- | --- |
| *Absidia anomala* Hesselt. & J.J. Ellis | 1964 | Cuba |  |  |
| *Absidia bonitoensis* C.L. Lima, D.X. Lima, Hyang B. Lee & A.L. Santiago | 2021 | Brazil |  |  |
| *Absidia caatingaensis* D.X. Lima & A.L. Santiago | 2015 | Brazil |  |  |
| *Absidia caerulea* Bainier | 1889 | France |  |  |
| *Absidia californica* J.J. Ellis & Hesselt. | 1965 | USA |  |  |
| *Absidia clavata* B.S. Mehrotra & Nand | 1967 | India |  |  |
| *Absidia cuneospora* G.F. Orr & Plunkett | 1959 | USA |  |  |
| *Absidia cylindrospora* Hagem | 1908 |  | √ | 2004 |
| *Absidia dubia* Bainier | 1882 |  |  |  |
| *Absidia egyptiaca* R. Sartory, J. Mey. & Tawfic | 1939 | Egypt |  |  |
| *Absidia fassatiae* Váňová | 1971 | Czech Republic |  |  |
| *Absidia glauca* Hagem | 1908 |  | √ | 2000 |
| *Absidia heterospora* Y. Ling | 1930 |  | √ | 2010 |
| *Absidia idahoensis* Hesselt., M.K. Mahoney & S.W. Peterson | 1990 | USA | √ | 2018 |
| *Absidia inflata* J.H. Mirza, S.M. Khan, S. Begum & Shagufta | 1979 | Pakistan |  |  |
| *Absidia jindoensis* Hyang B. Lee & T.T.T. Nguyen | 2018 | Republic of Korea |  |  |
| *Absidia koreana* Hyang B. Lee, Hye W. Lee & T.T. Nguyen | 2015 | Republic of Korea |  |  |
| *Absidia macrospora* Váňová | 1968 | Czech Republic |  |  |
| *Absidia multispora* T.R.L. Cordeiro, D.X Lima, Hyang B. Lee & A.L. Santiago | 2020 | Brazil |  |  |
| *Absidia narayanae* Subrahm. | 1990 | India |  |  |
| *Absidia panacisoli* T. Yuan Zhang, Ying Yu, He Zhu, S.Z. Yang, T.M. Yang, Meng Y. Zhang & Yi X. Zhang | 2018 | China |  |  |
| *Absidia pararepens* Jurjević, M. Kolařík & Hubka | 2020 |  |  |  |
| *Absidia pseudocylindrospora* Hesselt. & J.J. Ellis | 1962 | Tanzania | √ | 2010 |
| *Absidia psychrophilia* Hesselt. & J.J. Ellis | 1964 | Canada |  |  |
| *Absidia reflexa* Tiegh. | 1878 | France |  |  |
| *Absidia repens* Tiegh. | 1878 | France | √ | 2000 |
| *Absidia saloaensis* T.R.L. Cordeiro, D.X Lima, Hyang B. Lee & A.L. Santiago | 2020 | Brazil |  |  |
| *Absidia spinosa* Lendn. | 1907 | Switzerland | √ | 2004 |
| *Absidia stercoraria* Hyang B. Lee, H.S. Lee & T.T.T. Nguyen | 2016 | Republic of Korea |  |  |
| *Absidia terrestris* Rosas de Paz, Dania García, Guarro, Cano & Stchigel | 2018 | Mexico |  |  |
| *Absidia ushtrina* S.C. Agarwal | 1975 | India |  |  |
| *Acaulospora alpina* Oehl, Sýkorová & Sieverd. | 2006 | Switzerland |  |  |
| *Acaulospora aspera* Corazon-Guivin, Oehl & G.A. Silva | 2019 | Peru |  |  |
| *Acaulospora baetica* Palenz., Oehl, Azcón-Aguilar & G.A. Silva | 2015 | Spain |  |  |
| *Acaulospora bireticulata* F.M. Rothwell & Trappe | 1979 | USA | √ | 1990 |
| *Acaulospora brasiliensis* (B.T. Goto, L.C. Maia & Oehl) C. Walker, M. Krüger & A. Schüßler | 2011 | Brazil |  |  |
| *Acaulospora capsicula* Błaszk. | 1990 | Poland | √ | 2007 |
| *Acaulospora cavernata* Błaszk. | 1989 | Poland | √ | 2000 |
| *Acaulospora colliculosa* Kaonongbua, J.B. Morton & Bever | 2010 | USA |  |  |
| *Acaulospora colombiana* (Spain & N.C. Schenck) Kaonongbua, J.B. Morton & Bever | 2010 | Colombia | √ | 2018 |
| *Acaulospora colossica* P.A. Schultz, Bever & J.B. Morton | 1999 | USA | √ | 2009 |
| *Acaulospora delicata* C. Walker, C.M. Pfeiff. & Bloss | 1986 | USA | √ | 2003 |
| *Acaulospora denticulata* Sieverd. & S. Toro | 1987 | Colombia | √ | 1995 |
| *Acaulospora dilatata* J.B. Morton | 1986 | USA | √ | 1998 |
| *Acaulospora elegans* Trappe & Gerd. | 1974 | USA | √ | 1990 |
| *Acaulospora endographis* B.T. Goto | 2013 | Brazil |  |  |
| *Acaulospora entreriana* M.S. Velázquez & Cabello | 2008 | Argentina |  |  |
| *Acaulospora excavata* Ingleby & C. Walker | 1994 | Ivory Coast | √ | 2001 |
| *Acaulospora foveata* Trappe & Janos | 1982 | Mexico | √ | 1998 |
| *Acaulospora fragilissima* D. Redecker, Crossay & Cilia | 2018 | France |  |  |
| *Acaulospora gedanensis* Błaszk. | 1988 | Poland | √ | 2006 |
| *Acaulospora herrerae* E. Furrazola, B.T. Goto, G.A. Silva, Sieverd. & Oehl | 2012 | Cuba |  |  |
| *Acaulospora ignota* Błaszk., Góralska, Chwat & B.T. Goto | 2015 | Brazil |  |  |
| *Acaulospora koskei* Błaszk. | 1995 | Poland | √ | 2005 |
| *Acaulospora lacunosa* J.B. Morton | 1986 | USA | √ | 2000 |
| *Acaulospora laevis* Gerd. & Trappe | 1974 | USA | √ | 1990 |
| *Acaulospora longula* Spain & N.C. Schenck | 1984 | Colombia | √ | 1992 |
| *Acaulospora mellea* Spain & N.C. Schenck | 1984 | Colombia | √ | 1992 |
| *Acaulospora minuta* Oehl, Tchabi, Hountondji, Palenz., I.C. Sánchez & G.A. Silva | 2011 |  |  |  |
| *Acaulospora morrowiae* Spain & N.C. Schenck | 1984 | Colombia | √ | 2018 |
| *Acaulospora nivalis* Oehl, Palenz., I.C. Sánchez, G.A. Silva & Sieverd. | 2012 | Switzerland |  |  |
| *Acaulospora papillosa* C.M.R. Pereira & Oehl | 2016 | Brazil |  |  |
| *Acaulospora paulinae* Błaszk. | 1988 | Poland | √ | 2008 |
| *Acaulospora polonica* Błaszk. | 1988 | Poland | √ | 2001 |
| *Acaulospora punctata* Oehl, Palenz., Sánchez-Castro, G.A. Silva, C. Castillo & Sieverd. | 2011 | Switzerland |  |  |
| *Acaulospora pustulata* Palenz., Oehl, Azcón-Aguilar & G.A. Silva | 2013 | Spain |  |  |
| *Acaulospora reducta* Oehl, B.T. Goto & C.M.R. Pereira | 2016 | Brazil |  |  |
| *Acaulospora rehmii* Sieverd. & S. Toro | 1987 | Colombia | √ | 2003 |
| *Acaulospora rugosa* J.B. Morton | 1986 | USA | √ | 2001 |
| *Acaulospora saccata* D. Redecker, Crossay & Cilia | 2018 | France |  |  |
| *Acaulospora scrobiculata* Trappe | 1977 | Mexico | √ | 1992 |
| *Acaulospora sieverdingii* Oehl, Sýkorová & Błaszk. | 2011 |  |  |  |
| *Acaulospora soloidea* Vaing. & B.F. Rodrigues | 2011 | India |  |  |
| *Acaulospora spinosa* C. Walker & Trappe | 1981 | USA | √ | 1990 |
| *Acaulospora spinosissima* Oehl, Palenz., I.C. Sánchez, Tchabi, Hount. & G.A. Silva | 2019 | Benin |  |  |
| *Acaulospora spinulifera* Oehl, V.M. Santos, J.S. Pontes & G.A. Silva | 2017 | Brazil |  |  |
| *Acaulospora splendida* Sieverd., Chaverri & I. Rojas | 1988 | Costa Rica | √ | 2009 |
| *Acaulospora sporocarpia* S.M. Berch | 1985 | USA |  |  |
| *Acaulospora taiwania* H.T. Hu | 1988 | China |  |  |
| *Acaulospora terricola* Swarupa, Kunwar & Manohar. | 2003 | India |  |  |
| *Acaulospora thomii* Błaszk. | 1988 | Poland |  |  |
| *Acaulospora tortuosa* Palenz., Oehl, Azcón-Aguilar & G.A. Silva | 2013 | Spain |  |  |
| *Acaulospora tsugae* T.C. Lin & Oehl | 2019 | China |  |  |
| *Acaulospora tuberculata* Janos & Trappe | 1982 | Panama | √ | 1997 |
| *Acaulospora verna* Błaszk. | 2012 |  |  |  |
| *Acaulospora viridis* Palenz., Oehl, Azcón-Aguilar & G.A. Silva | 2014 | Spain |  |  |
| *Acaulospora walkeri* Kramad. & Hedger | 1990 | Indonesia |  |  |
| *Actinomucor elegans* (Eidam) C.R. Benj. & Hesselt. | 1957 |  | √ | 1973 |
| *Ambispora appendicula* (Spain, Sieverd. & N.C. Schenck) C. Walker | 2008 | Colombia | √ | 2018 |
| *Ambispora callosa* (Sieverd.) C. Walker, Vestberg & A. Schüßler | 2007 | Democratic Republic of the Congo | √ | 2017 |
| *Ambispora fecundispora* (N.C. Schenck & G.S. Sm.) C. Walker | 2008 | USA | √ | 2018 |
| *Ambispora fennica* C. Walker, Vestberg & A. Schüßler | 2007 | Finland |  |  |
| *Ambispora gerdemannii* (S.L. Rose, B.A. Daniels & Trappe) C. Walker, Vestberg & A. Schüßler | 2007 | USA | √ | 2014 |
| *Ambispora granatensis* Palenz., N. Ferrol & Oehl | 2011 | Spain |  |  |
| *Ambispora jimgerdemannii* (Spain, Oehl & Sieverd.) C. Walker | 2008 |  | √ | 2018 |
| *Ambispora leptoticha* (N.C. Schenck & G.S. Sm.) C. Walker, Vestberg & A. Schüßler | 2007 | USA | √ | 2017 |
| *Ambispora nicolsonii* (C. Walker, L.E. Reed & F.E. Sanders) Oehl, G.A. Silva, B.T. Goto & Sieverd. | 2011 | Great Britain |  |  |
| *Ambispora reticulata* Oehl & Sieverd. | 2012 | Switzerland |  |  |
| *Ambomucor clavatus* R.Y. Zheng & X.Y. Liu | 2014 | China | √ | 2013 |
| *Ambomucor ovalisporus* X.Y. Liu & R.Y. Zheng | 2015 | China | √ | 2015 |
| *Ambomucor seriatoinflatus* X.Y. Liu & R.Y. Zheng | 2014 | China | √ | 2013 |
| *Apophysomyces elegans* P.C. Misra, K.J. Srivast. & Lata | 1979 | India | √ | 1987 |
| *Apophysomyces mexicanus* A. Bonifaz, Cano, Stchigel & Guarro | 2014 | Mexico |  |  |
| *Apophysomyces ossiformis* E. Álvarez, Stchigel, Cano, Deanna A. Sutton & Guarro | 2010 | USA |  |  |
| *Apophysomyces thailandensis* Khuna, Suwannar. & Lumyong | 2019 | Thailand |  |  |
| *Apophysomyces trapeziformis* E. Álvarez, Stchigel, Cano, Deanna A. Sutton & Guarro | 2010 | USA |  |  |
| *Apophysomyces variabilis* E. Álvarez, Stchigel, Cano, Deanna A. Sutton & Guarro | 2010 | Netherlands |  |  |
| *Aquamortierella elegans* Embree & Indoh | 1967 | New Zealand |  |  |
| *Archaeospora ecuadoriana* A. Schüßler & C. Walker | 2019 | Ecuador |  |  |
| *Archaeospora europaea* Oehl, Palenz., Sánchez-Castro, V.M. Santos & G.A. Silva | 2019 | Switzerland |  |  |
| *Archaeospora myriocarpa* (Spain, Sieverd. & N.C. Schenck) Oehl, G.A. Silva, B.T. Goto & Sieverd. | 2011 | Colombia |  |  |
| *Archaeospora schenckii* (Sieverd. & S. Toro) C. Walker & A. Schüßler | 2010 | Colombia | √ | 2018 |
| *Archaeospora spainiae* (Oehl, Palenz., Sánchez-Castro & G.A. Silva) A. Schüßler & C. Walker | 2019 | Switzerland |  |  |
| *Archaeospora trappei* (R.N. Ames & Linderman) J.B. Morton & D. Redecker | 2001 | USA |  |  |
| *Archaeospora undulata* (Sieverd.) Sieverd., G.A. Silva, B.T. Goto & Oehl | 2011 | Democratic Republic of the Congo | √ | 2018 |
| *Backusella azygospora* T.R.L. Cordeiro, Hyang B. Lee & A.L. Santiago | 2019 | Brazil |  |  |
| *Backusella chlamydospora* Hyang B. Lee & T.T.T. Nguyen | 2021 | Republic of Korea |  |  |
| *Backusella circina* J.J. Ellis & Hesselt. | 1969 | USA | √ | 2000 |
| *Backusella constricta* D.X. Lima, C.A.F. de Souza & A.L. Santiago | 2016 | Brazil |  |  |
| *Backusella gigacellularis* J.I. Souza, Pires-Zottar. & Harakava | 2014 | Brazil |  |  |
| *Backusella grandis* (Schipper & Samson) Walther & de Hoog | 2013 | India |  |  |
| *Backusella granulispora* L.S. Loh & Kuthub. | 2001 | Malaysia |  |  |
| *Backusella indica* (Baijal & B.S. Mehrotra) Walther & de Hoog | 2013 | India |  |  |
| *Backusella koreana* Hyang B. Lee, J.S. Kim & T.T.T. Nguyen | 2021 | Republic of Korea |  |  |
| *Backusella lamprospora* (Lendn.) Benny & R.K. Benj. | 1975 | France | √ | 2018 |
| *Backusella locustae* Hyang B. Lee, S.H. Lee & T.T.T. Nguyen | 2018 | Republic of Korea |  |  |
| *Backusella oblongielliptica* (H. Nagan., Hirahara & Seshita ex Pidopl. & Milko) Walther & de Hoog | 2013 | Japan |  |  |
| *Backusella oblongispora* (Naumov) Walther & de Hoog | 2013 | former Soviet Union |  |  |
| *Backusella recurva* (E.E. Butler) Walther & de Hoog | 2013 | USA |  |  |
| *Backusella thermophila* Hyang B. Lee, A.L. Santiago, P.M. Kirk, K. Voigt & T.T.T. Nguyen | 2021 | Republic of Korea |  |  |
| *Backusella tuberculispora* (Schipper) Walther & de Hoog | 2013 | India |  |  |
| *Backusella variabilis* (A.K. Sarbhoy) Walther & de Hoog | 2013 | India |  |  |
| *Benjaminiella multispora* Benny, Samson & M.C. Sriniv. | 1985 | India |  |  |
| *Benjaminiella poitrasii* (R.K. Benj.) Arx | 1981 | USA |  |  |
| *Benjaminiella youngii* P.M. Kirk | 1989 | Spain |  |  |
| *Bifiguratus adelaidae* Torr.-Cruz & Porras-Alfaro | 2017 | USA |  |  |
| *Blakeslea monospora* B.S. Mehrotra & Baijal | 1968 | India | √ | 2003 |
| *Blakeslea trispora* Thaxt. | 1914 | USA | √ | 1973 |
| *Bulbospora minima* Oehl, Marinho, B.T. Goto & G.A. Silva | 2014 | Brazil |  |  |
| *Calcarisporiella thermophila* (H.C. Evans) de Hoog | 1974 | Great Britain |  |  |
| *Cetraspora armeniaca* (Błaszk.) Oehl, F.A. Souza & Sieverd. | 2009 | Poland | √ | 2014 |
| *Cetraspora auronigra* Oehl, L.L. Lima, Kozovits, Magna & G.A. Silva | 2014 | Brazil |  |  |
| *Cetraspora gilmorei* (Trappe & Gerd.) Oehl, F.A. Souza & Sieverd. | 2009 | USA | √ | 2014 |
| *Cetraspora helvetica* Oehl, Jansa, F.A. Souza & G.A. Silva | 2011 | Switzerland | √ | 2014 |
| *Cetraspora pellucida* (T.H. Nicolson & N.C. Schenck) Oehl, F.A. Souza & Sieverd. | 2009 | USA | √ | 2014 |
| *Chaetocladium brefeldii* Tiegh. & G. Le Monn. | 1873 | France | √ | 2008 |
| *Chaetocladium jonesiae* (Berk. & Broome) Fresen. | 1863 | Great Britain |  |  |
| *Chlamydoabsidia padenii* Hesselt. & J.J. Ellis | 1966 | USA |  |  |
| *Choanephora cucurbitarum* (Berk. & Ravenel) Thaxt. | 1903 | USA | √ | 1963 |
| *Choanephora infundibulifera* (Curr.) D.D. Cunn. | 1891 | India | √ | 1964 |
| *Circinella angarensis* (Schostak.) Zycha | 1935 |  | √ | 2017 |
| *Circinella lacrymispora* Aramb. & Cabello | 1996 | Argentina |  |  |
| *Circinella minor* Lendn. | 1905 | Switzerland |  |  |
| *Circinella mucoroides* Saito | 1907 |  | √ | 2017 |
| *Circinella muscae* (Sorokīn) Berl. & De Toni | 1888 |  | √ | 1995 |
| *Circinella nodulosa* R.Y. Zheng, X.Y. Liu & Y.N. Wang | 2017 | China | √ | 2017 |
| *Circinella ramosa* R.Y. Zheng, X.Y. Liu & Y.N. Wang | 2017 | China | √ | 2017 |
| *Circinella simplex* Tiegh. | 1875 | France | √ | 2017 |
| *Circinella umbellata* Tiegh. & G. Le Monn. | 1873 | France | √ | 1973 |
| *Claroideoglomus candidum* (E. Furrazola, Kaonongbua & Bever) Oehl, G.A. Silva & Sieverd. | 2011 | USA |  |  |
| *Claroideoglomus claroideum* (N.C. Schenck & G.S. Sm.) C. Walker & A. Schüßler | 2010 | USA | √ | 2017 |
| *Claroideoglomus drummondii* (Błaszk. & Renker) C. Walker & A. Schüßler | 2010 | Poland |  |  |
| *Claroideoglomus etunicatum* (W.N. Becker & Gerd.) C. Walker & A. Schüßler | 2010 | USA | √ | 2015 |
| *Claroideoglomus hanlinii* Błaszk., Chwat & Góralska | 2015 | Cuba |  |  |
| *Claroideoglomus lamellosum* (Dalpé, Koske & Tews) C. Walker & A. Schüßler | 2010 | Canada | √ | 2018 |
| *Claroideoglomus luteum* (L.J. Kenn., J.C. Stutz & J.B. Morton) C. Walker & A. Schüßler | 2010 | Canada | √ | 2018 |
| *Claroideoglomus walkeri* (Błaszk. & Renker) C. Walker & A. Schüßler | 2010 | Poland |  |  |
| *Cokeromyces recurvatus* Poitras | 1950 | USA |  |  |
| *Corymbiglomus corymbiforme* Błaszk. & Chwat | 2012 |  |  |  |
| *Corymbiglomus pacificum* Oehl, J. Medina, P. Cornejo, Sánchez-Castro, G.A. Silva & Palenz. | 2014 | Chile |  |  |
| *Cunninghamella bigelovii* Z.H. Xin, Y.Hui Zhao & Hui Wang | 2015 | China |  |  |
| *Cunninghamella binariae* R.Y. Zheng | 2001 |  | √ | 2001 |
| *Cunninghamella blakesleeana* Lendn. | 1927 |  | √ | 1973 |
| *Cunninghamella candida* Yosh. Yamam. | 1929 |  |  |  |
| *Cunninghamella clavata* R.Y. Zheng & G.Q. Chen | 1998 | China | √ | 1998 |
| *Cunninghamella echinulata* (Thaxt.) Thaxt. ex Blakeslee | 1905 | USA | √ | 1940 |
| *Cunninghamella elegans* Lendn. | 1907 | Switzerland | √ | 1972 |
| *Cunninghamella gigacellularis* A.L. Santiago, C.L. Lima & C.A.F. de Souza | 2016 | Brazil |  |  |
| *Cunninghamella homothallica* Komin. & Tubaki | 1952 | Japan | √ | 2001 |
| *Cunninghamella intermedia* K.B. Deshp. & Mantri | 1966 | India | √ | 2001 |
| *Cunninghamella multiverticillata* R.Y. Zheng & G.Q. Chen | 2001 | China | √ | 2001 |
| *Cunninghamella phaeospora* Boedijn | 1959 | Indonesia | √ | 2001 |
| *Cunninghamella polymorpha* Pišpek | 1929 |  | √ | 1992 |
| *Cunninghamella septata* R.Y. Zheng | 2001 | Belgium | √ | 2001 |
| *Cunninghamella vesiculosa* P.C. Misra | 1966 | India | √ | 2001 |
| *Densospora nanospora* McGee | 1996 | Australia |  |  |
| *Densospora nuda* McGee | 1996 | Australia |  |  |
| *Densospora solicarpa* McGee | 1996 | Australia |  |  |
| *Densospora tubiformis* (P.A. Tandy) McGee | 1996 | Australia | √ | 2018 |
| *Dentiscutata colliculosa* B.T. Goto & Oehl | 2010 | Brazil | √ | 2014 |
| *Dentiscutata erythropus* (Koske & C. Walker) C. Walker & D. Redecker | 2013 | USA | √ | 2017 |
| *Dentiscutata heterogama* (T.H. Nicolson & Gerd.) Sieverd., F.A. Souza & Oehl | 2009 | USA | √ | 2014 |
| *Dentiscutata nigerita* Khade | 2010 | India | √ | 2014 |
| *Dentiscutata nigra* (J.F. Redhead) Sieverd., F.A. Souza & Oehl | 2009 | USA | √ | 2014 |
| *Dentiscutata reticulata* (Koske, D.D. Mill. & C. Walker) Sieverd., F.A. Souza & Oehl | 2009 | USA | √ | 2014 |
| *Dentiscutata savannicola* (R.A. Herrera & Ferrer) C. Walker & A. Schüßler | 2014 | Cuba |  |  |
| *Desertispora omanana* (Symanczik, Błaszk. & Al-Yahya'ei) Symanczik, Błaszk., Kozłowska & Al-Yahya’ei | 2018 | Oman |  |  |
| *Dichotomocladium elegans* Benny & R.K. Benj. | 1975 | USA |  |  |
| *Dichotomocladium floridanum* Benny & R.K. Benj. | 1993 | USA |  |  |
| *Dichotomocladium hesseltinei* (B.S. Mehrotra & A.K. Sarbhoy) Benny & R.K. Benj. | 1975 | India |  |  |
| *Dichotomocladium robustum* Benny & R.K. Benj. | 1975 | USA |  |  |
| *Dichotomocladium sphaerosporum* R.K. Benj. & Benny | 1993 | Pakistan |  |  |
| *Dicranophora fulva* J. Schröt. | 1886 | Germany |  |  |
| *Dissophora decumbens* Thaxt. | 1914 | USA |  |  |
| *Dissophora nadsonii* Filippov | 1932 | Russia |  |  |
| *Dissophora ornata* (W. Gams) W. Gams | 1989 | Colombia |  |  |
| *Diversispora arenaria* (Błaszk., Tadych & Madej) Oehl, G.A. Silva & Sieverd. | 2011 |  |  |  |
| *Diversispora aurantia* (Błaszk., Blanke, Renker & Buscot) C. Walker & A. Schüßler | 2010 | Poland | √ | 2016 |
| *Diversispora celata* C. Walker, Gamper & A. Schüßler | 2009 | Switzerland | √ | 2016 |
| *Diversispora clara* Oehl, B. Estrada, G.A. Silva & Palenz. | 2011 | Spain |  |  |
| *Diversispora eburnea* (L.J. Kenn., J.C. Stutz & J.B. Morton) C. Walker & A. Schüßler | 2010 | USA | √ | 2018 |
| *Diversispora epigaea* (B.A. Daniels & Trappe) C. Walker & A. Schüßler | 2010 | USA | √ | 2016 |
| *Diversispora gibbosa* (Błaszk.) Błaszk. & Kovács | 2011 | Poland |  |  |
| *Diversispora insculpta* (Błaszk.) Oehl, G.A. Silva & Sieverd. | 2011 | Poland |  |  |
| *Diversispora jakucsiae* Błaszk., T.K. Balázs & Kovács | 2015 | Poland |  |  |
| *Diversispora peloponnesiaca* Błaszk., B.T. Goto, Orfanoudakis & Niezgoda | 2019 | Poland |  |  |
| *Diversispora peridiata* Błaszk., Chwat, Kovács & Góralska | 2015 | Poland |  |  |
| *Diversispora sabulosa* Błaszk. & Kozłowska | 2018 | Lithuania |  |  |
| *Diversispora slowinskiensis* Błaszk., Chwat, Kovács & Góralska | 2015 | Poland |  |  |
| *Diversispora sporocarpia* Chachuła, Mleczko, Zubek, Niezgoda, A. Kozłowska, Jobim, B.T. Goto & Błaszk. | 2019 | Poland |  |  |
| *Diversispora spurca* (C.M. Pfeiff., C. Walker & Bloss) C. Walker & A. Schüßler | 2004 | USA | √ | 2007 |
| *Diversispora tenera* (P.A. Tandy) Oehl, G.A. Silva & Sieverd. | 2011 | Australia | √ | 2018 |
| *Diversispora trimurales* (Koske & Halvorson) C. Walker & A. Schüßler | 2010 | USA | √ | 2018 |
| *Diversispora varaderana* Błaszk., Chwat, Kovács & Góralska | 2015 | Cuba |  |  |
| *Dominikia achra* (Błaszk., D. Redecker, Koegel, Schützek, Oehl & Kovács) Błaszk., Chwat & Kovács | 2014 | Poland |  |  |
| *Dominikia aurea* (Oehl & Sieverd.) Błaszk., Chwat, G.A. Silva & Oehl | 2014 | Switzerland | √ | 2018 |
| *Dominikia bernensis* Oehl, Palenz., Sánchez-Castro, N.M.F. Sousa & G.A. Silva | 2014 | Switzerland |  |  |
| *Dominikia compressa* (Sieverd., Oehl, Palenz., Sánchez-Castro & G.A. Silva) Oehl, Palenz., Sánchez-Castro & G.A. Silva | 2014 | Switzerland |  |  |
| *Dominikia difficilevidera* Błaszk., Góralska & Chwat | 2015 | Poland |  |  |
| *Dominikia distichi* Błaszk., Chwat & Kovács | 2014 | South Africa |  |  |
| *Dominikia duoreactiva* Błaszk., Góralska & Chwat | 2015 | Egypt |  |  |
| *Dominikia iranica* (Błaszk., Kovács & Balázs) Błaszk., Chwat & Kovács | 2014 | Iran |  |  |
| *Dominikia lithuanica* Błaszk., Chwat & Góralska | 2016 | Lithuania |  |  |
| *Dominikia minuta* (Błaszk., Tadych & Madej) Błaszk., Chwat & Kovács | 2014 | Poland |  |  |
| *Echinochlamydosporium variabile* X.Z. Jiang, H.Y. Yu, M.C. Xiang, X.Y. Liu & Xing Z. Liu | 2011 | China | √ | 2011 |
| *Ellisomyces anomalus* (Hesselt. & P. Anderson) Benny & R.K. Benj. | 1975 | USA |  |  |
| *Endogone acrogena* Gerd., Trappe & Hosford | 1974 | USA | √ | 1984 |
| *Endogone aggregata* P.A. Tandy | 1975 | Australia | √ | 1984 |
| *Endogone alba* (Petch) Gerd. & Trappe | 1974 |  | √ | 1984 |
| *Endogone aurantiaca* Błaszk. | 1997 | Poland |  |  |
| *Endogone botryocarpa* Koh. Yamam., Degawa & A. Yamada | 2020 | Japan |  |  |
| *Endogone carolinensis* (Y.J. Yao) Desirò, M.E. Sm., Bonito, Bidartondo & Trappe | 2017 | USA |  |  |
| *Endogone corticioides* Koh. Yamam., Degawa & A. Yamada | 2017 | Japan | √ | 2017 |
| *Endogone crassa* P.A. Tandy | 1975 | Australia | √ | 1984 |
| *Endogone incrassata* Thaxt. | 1922 | USA & Canada | √ | 1984 |
| *Endogone irregularis* Szem. | 1965 | Hungary |  |  |
| *Endogone lanata* Harkn. | 1899 | USA |  |  |
| *Endogone ludwigii* Bucholtz | 1912 | Germany |  |  |
| *Endogone maritima* Błaszk., Tadych & Madej | 1998 | Poland |  |  |
| *Endogone minutissima* Beeli | 1924 |  |  |  |
| *Endogone multiplex* Thaxt. | 1922 | USA |  |  |
| *Endogone occidentalis* Kanouse | 1936 | USA |  |  |
| *Endogone oregonensis* Gerd. & Trappe | 1974 | USA | √ | 1984 |
| *Endogone pegleri* Y.J. Yao | 1995 | Australia |  |  |
| *Endogone pisiformis* Link | 1809 | Germany | √ | 1984 |
| *Endogone pseudopisiformis* Y.J. Yao | 1995 | USA |  |  |
| *Endogone reticulata* P.A. Tandy | 1975 | Australia | √ | 1984 |
| *Endogone rosea* Zeller | 1941 | USA |  |  |
| *Endogone sphagnophila* G.F. Atk. | 1918 |  |  |  |
| *Endogone stratosa* Trappe, Gerd. & Fogel | 1974 | USA | √ | 1984 |
| *Endogone tjibodensis* Boedijn | 1935 | Indonesia |  |  |
| *Endogone torrendii* Bres. | 1920 | 'Lusitania' |  |  |
| *Endogone tuberculosa* Lloyd | 1918 | Australia | √ | 1984 |
| *Endogone verrucosa* Gerd. & Trappe | 1974 | USA | √ | 1984 |
| *Endogone xylogena* (Sacc.) J. Schröt. | 1887 |  |  |  |
| *Entrophospora hexagoni* Rhatwal & Gandhe | 2009 | India |  |  |
| *Entrophospora infrequens* (I.R. Hall) R.N. Ames & R.W. Schneid. | 1979 | New Zealand | √ | 1990 |
| *Fennellomyces gigacellularis* J.H. Mirza, S.M. Khan, S. Begum & Shagufta | 1979 | Pakistan |  |  |
| *Fennellomyces heterothallicus* P.C. Misra, N.N. Gupta & Lata | 1979 | India |  |  |
| *Fennellomyces linderi* (Hesselt. & Fennell) Benny & R.K. Benj. | 1975 | USA |  |  |
| *Fennellomyces verticillatus* J.H. Mirza, S.M. Khan, S. Begum & Shagufta | 1979 | Pakistan |  |  |
| *Funneliformis badius* (Oehl, D. Redecker & Sieverd.) C. Walker & A. Schüßler | 2010 | Germany |  |  |
| *Funneliformis caledonius* (T.H. Nicolson & Gerd.) C. Walker & A. Schüßler | 2010 | Great Britain |  |  |
| *Funneliformis constrictus* (Trappe) C. Walker & A. Schüßler | 2010 | Mexico |  |  |
| *Funneliformis coronatus* (Giovann.) C. Walker & A. Schüßler | 2010 | Italy |  |  |
| *Funneliformis dimorphicus* (Boyetchko & J.P. Tewari) Oehl, G.A. Silva & Sieverd. | 2011 | Canada | √ | 2018 |
| *Funneliformis fragilistratus* (Skou & I. Jakobsen) C. Walker & A. Schüßler | 2010 | Denmark |  |  |
| *Funneliformis geosporus* (T.H. Nicolson & Gerd.) C. Walker & A. Schüßler | 2010 | Great Britain |  |  |
| *Funneliformis halonatus* (S.L. Rose & Trappe) Oehl, G.A. Silva & Sieverd. | 2011 | Great Britain | √ | 2018 |
| *Funneliformis mosseae* (T.H. Nicolson & Gerd.) C. Walker & A. Schüßler | 2010 | Great Britain | √ | 2015 |
| *Funneliformis verruculosus* (Błaszk.) C. Walker & A. Schüßler | 2010 | Poland |  |  |
| *Funneliglomus sanmartinense* Corazon-Guivin, G.A. Silva & Oehl | 2019 | Peru |  |  |
| *Gamsiella multidivaricata* (R.K. Benj.) Benny & M. Blackw. | 2004 | USA |  |  |
| *Geosiphon pyriformis* (Kütz.) F. Wettst. | 1915 | Germany |  |  |
| *Gigaspora albida* N.C. Schenck & G.S. Sm. | 1982 | USA | √ | 2010 |
| *Gigaspora alboaurantiaca* W.N. Chou | 1991 | China | √ | 2014 |
| *Gigaspora candida* Bhattacharjee, Mukerji, J.P. Tewari & Skoropad | 1982 | India | √ | 2014 |
| *Gigaspora decipiens* I.R. Hall & L.K. Abbott | 1984 | Australia | √ | 2002 |
| *Gigaspora gigantea* (T.H. Nicolson & Gerd.) Gerd. & Trappe | 1974 | USA | √ | 1984 |
| *Gigaspora lazzarii* Montecchi, Ruini & G. Gross | 1996 | Italy |  |  |
| *Gigaspora margarita* W.N. Becker & I.R. Hall | 1976 | USA | √ | 1984 |
| *Gigaspora ramisporophora* Spain, Sieverd. & N.C. Schenck | 1989 | Brazil | √ | 2012 |
| *Gigaspora rosea* T.H. Nicolson & N.C. Schenck | 1979 | USA | √ | 1984 |
| *Gigaspora tuberculata* Neeraj, Mukerji, B.C. Sharma & A.K. Varma | 1993 | India | √ | 2009 |
| *Gilbertella persicaria* (E.D. Eddy) Hesselt. | 1960 | USA | √ | 1964 |
| *Glomus ambisporum* G.S. Sm. & N.C. Schenck | 1985 | USA | √ | 1991 |
| *Glomus arborense* McGee | 1986 | Australia | √ | 1991 |
| *Glomus atrouva* McGee & Pattinson | 2002 | Australia |  |  |
| *Glomus australe* (Berk.) S.M. Berch | 1983 | Australia | √ | 1991 |
| *Glomus avelingiae* R.C. Sinclair | 2000 | South Africa |  |  |
| *Glomus bagyarajii* V.S. Mehrotra | 1997 |  |  |  |
| *Glomus boreale* (Thaxt.) Trappe & Gerd. | 1974 | Canada | √ | 1984 |
| *Glomus botryoides* F.M. Rothwell & Victor | 1984 | USA | √ | 1991 |
| *Glomus brohultii* R.A. Herrera, Ferrer & Sieverd. | 2003 | Cuba |  |  |
| *Glomus caesaris* Sieverd. & Oehl | 2002 | Germany |  |  |
| *Glomus canadense* (Thaxt.) Trappe & Gerd. | 1974 | Canada | √ | 1991 |
| *Glomus canum* McGee | 2002 | Australia |  |  |
| *Glomus cerebriforme* McGee | 1986 | Australia | √ | 1991 |
| *Glomus citricola* D.Z. Tang & M. Zang | 1984 | China | √ | 2011 |
| *Glomus clavisporum* (Trappe) R.T. Almeida & N.C. Schenck | 1990 | Mexico | √ | 2018 |
| *Glomus convolutum* Gerd. & Trappe | 1974 | USA | √ | 1991 |
| *Glomus coremioides* (Berk. & Broome) D. Redecker & J.B. Morton | 2000 | Sri Lanka | √ | 2001 |
| *Glomus corymbiforme* Błaszk. | 1995 | Poland |  |  |
| *Glomus crenatum* E. Furrazola, Ferrer, R.A. Herrera & B.T. Goto | 2011 | Cuba |  |  |
| *Glomus cuneatum* McGee & A. Cooper | 2002 | Australia |  |  |
| *Glomus delhiense* Mukerji, Bhattacharjee & J.P. Tewari | 1983 | India | √ | 1991 |
| *Glomus dolichosporum* M.Q. Zhang & You S. Wang | 1997 | China | √ | 1997 |
| *Glomus flavisporum* (M. Lange & E.M. Lund) Trappe & Gerd. | 1974 | Denmark | √ | 1991 |
| *Glomus formosanum* C.G. Wu & Z.C. Chen | 1986 | China | √ | 1991 |
| *Glomus fragile* (Berk. & Broome) Trappe & Gerd. | 1974 | Sri Lanka | √ | 2018 |
| *Glomus fuegianum* (Speg.) Trappe & Gerd. | 1974 | Argentina | √ | 1991 |
| *Glomus globiferum* Koske & C. Walker | 1986 | USA | √ | 1991 |
| *Glomus glomerulatum* Sieverd. | 1987 | Colombia | √ | 1991 |
| *Glomus goaensis* Khade | 2009 | India |  |  |
| *Glomus herrerae* Torres-Arias, E. Furrazola & B.T. Goto | 2019 | Cuba |  |  |
| *Glomus heterosporum* G.S. Sm. & N.C. Schenck | 1985 | USA | √ | 1991 |
| *Glomus hoi* S.M. Berch & Trappe | 1985 | USA | √ | 1991 |
| *Glomus hyderabadensis* Swarupa, Kunwar, G.S. Prasad & Manohar. | 2004 | India | √ | 2006 |
| *Glomus indicum* Błaszk., Wubet & Harikumar | 2010 | Kerala and Eritrea |  |  |
| *Glomus kerguelense* Dalpé & Strullu | 2002 | Kerguelen |  |  |
| *Glomus liquidambaris* (C.G. Wu & Z.C. Chen) R.T. Almeida & N.C. Schenck ex Y.J. Yao | 1995 | China | √ | 2003 |
| *Glomus macrocarpum* Tul. & C. Tul. | 1844 | France | √ | 1990 |
| *Glomus magnicaule* I.R. Hall | 1977 | New Zealand | √ | 2003 |
| *Glomus majewskii* Błaszk. | 2010 | Denmark |  |  |
| *Glomus melanosporum* Gerd. & Trappe | 1974 | USA | √ | 1991 |
| *Glomus microcarpum* Tul. & C. Tul. | 1844 | France | √ | 1990 |
| *Glomus monosporum* Gerd. & Trappe | 1974 | USA | √ | 1991 |
| *Glomus mortonii* Bentiv. & Hetrick | 1991 | USA | √ | 2012 |
| *Glomus multicaule* Gerd. & B.K. Bakshi | 1976 | India | √ | 1991 |
| *Glomus multiforum* Tadych & Błaszk. | 1997 | Poland | √ | 2006 |
| *Glomus multisubstensum* Mukerji, Bhattacharjee & J.P. Tewari | 1983 | India | √ | 1991 |
| *Glomus mume* B.P. Cai, Jun Y. Chen, Q.X. Zhang & L.D. Guo | 2013 | China |  |  |
| *Glomus nanolumen* Koske & Gemma | 1990 | USA | √ | 2008 |
| *Glomus pallidum* I.R. Hall | 1977 | New Zealand | √ | 1990 |
| *Glomus pansihalos* S.M. Berch & Koske | 1986 | USA | √ | 1991 |
| *Glomus pellucidum* McGee & Pattinson | 2002 | Australia |  |  |
| *Glomus przelewicense* Błaszk. | 1988 | Poland |  |  |
| *Glomus pubescens* (Sacc. & Ellis) Trappe & Gerd. | 1974 | USA | √ | 1984 |
| *Glomus pustulatum* Koske, Friese, C. Walker & Dalpé | 1986 | USA | √ | 1991 |
| *Glomus radiatum* (Thaxt.) Trappe & Gerd. | 1974 | USA | √ | 1991 |
| *Glomus reticulatum* Bhattacharjee & Mukerji | 1980 | India | √ | 1991 |
| *Glomus rubiforme* (Gerd. & Trappe) R.T. Almeida & N.C. Schenck | 1990 | USA | √ | 2000 |
| *Glomus segmentatum* Trappe, Spooner & Ivory | 1979 | Belize | √ | 1991 |
| *Glomus sinuosum* (Gerd. & B.K. Bakshi) R.T. Almeida & N.C. Schenck | 1990 | India | √ | 2001 |
| *Glomus spinosum* H.T. Hu | 2002 | China | √ | 2002 |
| *Glomus spinuliferum* Sieverd. & Oehl | 2003 | Germany | √ | 2009 |
| *Glomus taiwanense* (C.G. Wu & Z.C. Chen) R.T. Almeida & N.C. Schenck ex Y.J. Yao | 1995 | China | √ | 2001 |
| *Glomus tenebrosum* (Thaxt.) S.M. Berch | 1983 | Canada | √ | 1991 |
| *Glomus tetrastratosum* Błaszk., Chwat & Góralska | 2015 | Poland |  |  |
| *Glomus trufemii* B.T. Goto, G.A. Silva & Oehl | 2012 | Brazil |  |  |
| *Glomus versiforme* (P. Karst.) S.M. Berch | 1983 | Finland | √ | 1991 |
| *Glomus warcupii* McGee | 1986 | Australia | √ | 1991 |
| *Gongronella brasiliensis* C.A.F. de Souza, D.X. Lima & A.L. Santiago | 2017 | Brazil |  |  |
| *Gongronella butleri* (Lendn.) Peyronel & Dal Vesco | 1955 | Malaysia | √ | 1990 |
| *Gongronella guangdongensis* F. Liu, T.T. Liu & L. Cai | 2015 | China |  |  |
| *Gongronella koreana* Hyang B. Lee & T.T.T. Nguyen | 2015 | Republic of Korea |  |  |
| *Gongronella lacrispora* Hesselt. & J.J. Ellis | 1962 | USA |  |  |
| *Gongronella namwonensis* Hyang B. Lee, A.L. Santiago & H.J. Lim | 2020 | Republic of Korea |  |  |
| *Gongronella orasabula* Hyang B. Lee, K. Voigt, P.M. Kirk & T.T.T. Nguyen | 2016 | Republic of Korea |  |  |
| *Gongronella pedratalhadensis* L.W.S. Freitas, H.B. Lee & A.L. Santiago | 2020 | Brazil |  |  |
| *Gongronella sichuanensis* Z.Y. Zhang, Y.F. Han, W.H. Chen & Z.Q. Liang | 2019 | China |  |  |
| *Halteromyces radiatus* Shipton & Schipper | 1975 | Australia |  |  |
| *Helicostylum elegans* Corda | 1842 | Czech Republic |  |  |
| *Helicostylum pulchrum* (Preuss) Pidopl. & Milko | 1971 |  |  |  |
| *Hesseltinella vesiculosa* H.P. Upadhyay | 1970 | Brazil |  |  |
| *Hyphomucor assamensis* (B.S. Mehrotra & B.R. Mehrotra) Schipper & Lunn | 1986 | India |  |  |
| *Innospora majewskii* (Blaszk. & Kovács) Błaszk., Kovács, Chwat & Kozłowska | 2017 | Poland |  |  |
| *Intraornatospora intraornata* (B.T. Goto & Oehl) B.T. Goto, Oehl & G.A. Silva | 2012 | Brazil |  |  |
| *Isomucor trufemiae* J.I. Souza, Pires-Zottar. & Harakava | 2012 | Brazil |  |  |
| *Jimgerdemannia ambigua* Koh. Yamam., Degawa & A. Yamada | 2020 | Japan |  |  |
| *Jimgerdemannia flammicorona* (Trappe & Gerd.) Trappe, Desirò, M.E. Sm., Bonito & Bidartondo | 2017 | Italy |  |  |
| *Jimgerdemannia lactiflua* (Berk. & Broome) Trappe, Desirò, M.E. Sm., Bonito & Bidartondo | 2017 | Great Britain |  |  |
| *Kamienskia bistrata* (Błaszk., D. Redecker, Koegel, Symanczik, Oehl & Kovács) Błaszk., Chwat & Kovács | 2014 | Poland |  |  |
| *Kirkiana ramosa* L.S. Loh, Kuthub. & Nawawi | 2001 | Malaysia |  |  |
| *Kirkomyces cordensis* (B.S. Mehrotra & B.R. Mehrotra) Benny | 1996 | India |  |  |
| *Kuklospora spinosa* B.P. Cai, Jun Y. Chen, Q.X. Zhang & L.D. Guo | 2013 | China | √ | 2013 |
| *Lentamyces indicus* S.D. Patil & J.C. Kale ex Kerst. Hoffm. & K. Voigt | 2009 |  |  |  |
| *Lentamyces minutissimus* (Faurel & Schotter) Kerst. Hoffm. & K. Voigt | 2009 | Congo |  |  |
| *Lentamyces parricidus* (Renner & Muskat ex Hesselt. & J.J. Ellis) Kerst. Hoffm. & K. Voigt | 2008 | Tunisia |  |  |
| *Lentamyces zychae* (Hesselt. & J.J. Ellis) Kerst. Hoffm. & K. Voigt | 2008 | Germany |  |  |
| *Lichtheimia blakesleeana* (Lendn.) Kerst. Hoffm., Walther & K. Voigt | 2009 |  |  |  |
| *Lichtheimia brasiliensis* A.L. Santiago, D.X. Lima & R.J.V. Oliveira | 2013 | Brazil |  |  |
| *Lichtheimia corymbifera* (Cohn) Vuill. | 1903 | Poland | √ | 2012 |
| *Lichtheimia hyalospora* (Saito) Kerst. Hoffm., Walther & K. Voigt | 2009 |  | √ | 2012 |
| *Lichtheimia ornata* (A.K. Sarbhoy) Alastr.-Izq. & Walther | 2010 | India | √ | 2012 |
| *Lichtheimia ramosa* (Zopf) Vuill. | 1903 |  | √ | 2012 |
| *Lichtheimia sphaerocystis* Alastr.-Izq. & Walther | 2010 | India |  |  |
| *Lobosporangium transversale* (Malloch) M. Blackw. & Benny | 2004 | USA |  |  |
| *Microdominikia litorea* (Błaszk. & Kozłowska) Oehl, Corazon-Guivin & G.A. Silva | 2019 | Greece |  |  |
| *Microkamienskia divaricata* (Błaszk., Chwat & Góralska) Corazon-Guivin, G.A. Silva & Oehl | 2019 | South Africa |  |  |
| *Microkamienskia perpusilla* (Blaszk. & Kovács) Corazon-Guivin, G.A. Silva & Oehl | 2019 | Poland |  |  |
| *Microkamienskia peruviana* Corazon-Guivin, G.A. Silva & Oehl | 2019 | Peru |  |  |
| *Modicella malleola* (Harkn.) Gerd. & Trappe | 1974 | USA |  |  |
| *Modicella reniformis* (Bres.) Gerd. & Trappe | 1974 | Brazil |  |  |
| *Mortierella acrotona* W. Gams | 1976 | India |  |  |
| *Mortierella alliacea* Linnem. | 1953 | Germany |  |  |
| *Mortierella alpina* Peyronel | 1913 | Italy | √ | 1992 |
| *Mortierella ambigua* B.S. Mehrotra | 1963 | India |  |  |
| *Mortierella amoeboidea* W. Gams | 1976 | Germany |  |  |
| *Mortierella angusta* (Linnem.) Linnem. | 1969 | Germany |  |  |
| *Mortierella antarctica* Linnem. | 1969 | Antarctica |  |  |
| *Mortierella apiculata* Marchal | 1891 | Belgium |  |  |
| *Mortierella arcuata* E. Wolf | 1954 | Germany |  |  |
| *Mortierella armillariicola* W. Gams | 1976 | Netherlands |  |  |
| *Mortierella baccata* E. Wolf | 1954 | Germany |  |  |
| *Mortierella bainieri* Costantin | 1889 | France |  |  |
| *Mortierella basiparvispora* W. Gams & Grinb. | 1976 | Chile |  |  |
| *Mortierella beljakovae* Milko | 1973 | Ukraine |  |  |
| *Mortierella biramosa* Tiegh. | 1875 | France |  |  |
| *Mortierella bisporalis* (Thaxt.) Björl. | 1936 | USA | √ | 2018 |
| *Mortierella calciphila* Wrzosek | 2016 | Poland |  |  |
| *Mortierella camargensis* W. Gams & R. Moreau | 1960 |  |  |  |
| *Mortierella candelabrum* Tiegh. & G. Le Monn. | 1873 | France |  |  |
| *Mortierella capitata* Marchal | 1891 | Belgium |  |  |
| *Mortierella cephalosporina* Chalab. | 1965 | Ukraine |  |  |
| *Mortierella chienii* P.M. Kirk | 2012 | USA |  |  |
| *Mortierella chlamydospora* (Chesters) Plaäts-Nit. | 1976 | Great Britain |  |  |
| *Mortierella claussenii* Linnem. | 1958 | Europe |  |  |
| *Mortierella clonocystis* W. Gams | 1976 | Spain |  |  |
| *Mortierella cogitans* Degawa | 1998 | Japan |  |  |
| *Mortierella cystojenkinii* W. Gams & Veenb.-Rijks | 1976 | Netherlands |  |  |
| *Mortierella dichotoma* Linnem. ex W. Gams | 1977 | Germany |  |  |
| *Mortierella echinosphaera* Plaäts-Nit. | 1976 | Netherlands |  |  |
| *Mortierella echinula* Linnem. | 1953 |  |  |  |
| *Mortierella echinulata* Harz | 1871 | Germany |  |  |
| *Mortierella elasson* Sideris & G.E. Paxton | 1929 | USA |  |  |
| *Mortierella elongata* Linnem. | 1941 | Germany | √ | 1992 |
| *Mortierella elongatula* W. Gams & Domsch | 1976 | Germany |  |  |
| *Mortierella epicladia* W. Gams & Emden | 1976 | Spain |  |  |
| *Mortierella epigama* W. Gams & Domsch | 1972 | Germany |  |  |
| *Mortierella exigua* Linnem. | 1941 | Germany | √ | 2018 |
| *Mortierella fimbriata* S.H. Ou | 1940 | China | √ | 1940 |
| *Mortierella fimbricystis* W. Gams | 1976 | Argentina |  |  |
| *Mortierella fluviae* Hyang B. Lee, K. Voigt & T.T.T. Nguyen | 2016 | Republic of Korea |  |  |
| *Mortierella formicae* Siedlecki | 2017 | Poland |  |  |
| *Mortierella formosana* S.F. Wei, H.M. Ho & K. Voigt | 2015 | China | √ | 2015 |
| *Mortierella gamsii* Milko | 1974 |  | √ | 1986 |
| *Mortierella gemmifera* M. Ellis | 1941 | Great Britain | √ | 1992 |
| *Mortierella globalpina* W. Gams & Veenb.-Rijks | 1976 | Netherlands | √ | 1992 |
| *Mortierella globulifera* O. Rostr. | 1916 |  |  |  |
| *Mortierella hepiali* Q.T. Chen & B. Liu | 1986 | China | √ | 1986 |
| *Mortierella histoplasmatoides* W. Gams | 1991 | USA |  |  |
| *Mortierella horticola* Linnem. | 1941 | Germany | √ | 1992 |
| *Mortierella humicola* Oudem. | 1902 |  |  |  |
| *Mortierella humilis* Linnem. ex W. Gams | 1977 | Germany | √ | 1992 |
| *Mortierella humilissima* Pišpek | 1929 |  |  |  |
| *Mortierella hyalina* (Harz) W. Gams | 1970 |  | √ | 1992 |
| *Mortierella hypsicladia* Degawa & W. Gams | 2004 | Japan |  |  |
| *Mortierella indohii* C.Y. Chien | 1974 | USA | √ | 1992 |
| *Mortierella insignis* Linnem. | 1941 | Germany |  |  |
| *Mortierella jenkinii* (A.L. Sm.) Naumov | 1935 | Great Britain | √ | 1992 |
| *Mortierella kuhlmanii* W. Gams | 1976 | USA |  |  |
| *Mortierella lignicola* (G.W. Martin) W. Gams & R. Moreau | 1960 | Colombia |  |  |
| *Mortierella longigemmata* Linnem. | 1969 | Germany |  |  |
| *Mortierella macrocystis* W. Gams | 1961 |  |  |  |
| *Mortierella macrocystopsis* W. Gams & Carreiro | 1989 | USA |  |  |
| *Mortierella mairei* Vuill. | 1918 | France |  |  |
| *Mortierella mehrotraensis* Baijal | 1968 | India |  |  |
| *Mortierella microspora* E. Wolf | 1954 | Germany |  |  |
| *Mortierella microzygospora* Degawa | 1998 | Japan |  |  |
| *Mortierella minutissima* Tiegh. | 1878 | France | √ | 2018 |
| *Mortierella mundensis* Linnem. | 1941 | Germany |  |  |
| *Mortierella mutabilis* Linnem. | 1941 | Germany | √ | 1992 |
| *Mortierella nantahalensis* C.Y. Chien | 1971 | USA |  |  |
| *Mortierella niveovelutina* Cif. & Ashford | 1929 |  |  |  |
| *Mortierella oligospora* Björl. | 1936 |  |  |  |
| *Mortierella ovalispora* Chalab. | 1965 | Ukraine |  |  |
| *Mortierella paraensis* Pfenning & W. Gams | 1993 | Brazil |  |  |
| *Mortierella parazychae* W. Gams | 1976 | Netherlands |  |  |
| *Mortierella parvispora* Linnem. | 1941 | Germany | √ | 1992 |
| *Mortierella pilulifera* Tiegh. | 1875 | France |  |  |
| *Mortierella pisiformis* H.M. Ho, S.F. Wei & K. Voigt | 2015 | China | √ | 2015 |
| *Mortierella plectoconfusa* E. Wolf | 1954 | Germany |  |  |
| *Mortierella polycephala* Coem. | 1863 | Belgium | √ | 1992 |
| *Mortierella polygonia* W. Gams & Veenb.-Rijks | 1976 | Netherlands |  |  |
| *Mortierella pseudozygospora* W. Gams & Carreiro | 1989 | USA |  |  |
| *Mortierella pulchella* Linnem. | 1941 | Germany |  |  |
| *Mortierella pusilla* Oudem. | 1902 |  | √ | 2011 |
| *Mortierella pygmaea* Chalab. | 1965 | Ukraine |  |  |
| *Mortierella repens* A.L. Sm. | 1898 | Great Britain |  |  |
| *Mortierella reticulata* Tiegh. & G. Le Monn. | 1873 | France | √ | 1992 |
| *Mortierella rhizogena* Dasz. | 1912 | Switzerland |  |  |
| *Mortierella rishikesha* B.S. Mehrotra & B.R. Mehrotra | 1964 | India |  |  |
| *Mortierella rostafinskii* Bref. | 1881 |  |  |  |
| *Mortierella sarnyensis* Milko | 1973 | Ukraine |  |  |
| *Mortierella schmuckeri* Linnem. | 1958 | Mexico |  |  |
| *Mortierella sclerotiella* Milko | 1967 | Ukraine |  |  |
| *Mortierella selenospora* W. Gams | 1976 | Netherlands |  |  |
| *Mortierella signyensis* K. Voigt, P.M. Kirk & Bridge | 2012 | Antarctica |  |  |
| *Mortierella simplex* Tiegh. & G. Le Monn. | 1873 | France |  |  |
| *Mortierella strangulata* Tiegh. | 1875 | France |  |  |
| *Mortierella striospora* K.B. Deshp. & Mantri | 1963 | India |  |  |
| *Mortierella stylospora* Dixon-Stew. | 1932 | Great Britain | √ | 2011 |
| *Mortierella subtilissima* Oudem. | 1902 |  |  |  |
| *Mortierella sugadairana* Y. Takash., Degawa & K. Narisawa | 2018 | Japan |  |  |
| *Mortierella thereuopodae* Degawa | 2014 | Japan |  |  |
| *Mortierella tirolensis* Linnem. | 1969 | Italy |  |  |
| *Mortierella traversoana* Peyronel | 1913 |  |  |  |
| *Mortierella tsukubaensis* Ts. Watan. | 2001 | Japan |  |  |
| *Mortierella tuberosa* Tiegh. | 1875 | France |  |  |
| *Mortierella turficola* Y. Ling | 1930 |  |  |  |
| *Mortierella verrucosa* Linnem. | 1953 | Germany | √ | 1992 |
| *Mortierella verticillata* Linnem. | 1941 | Germany | √ | 1992 |
| *Mortierella wolfii* B.S. Mehrotra & Baijal | 1963 | India |  |  |
| *Mortierella wuyishanensis* F.J. Chen | 1992 | China | √ | 1992 |
| *Mortierella zonata* Linnem. ex W. Gams | 1977 | Germany |  |  |
| *Mortierella zychae* Linnem. | 1941 | Germany | √ | 1992 |
| *Mucor abundans* Povah | 1917 |  | √ | 1972 |
| *Mucor adventitius* Oudem. | 1902 |  | √ | 1973 |
| *Mucor albus* Schrank | 1789 | Germany |  |  |
| *Mucor aligarensis* B.S. Mehrotra & B.R. Mehrotra | 1970 | India |  |  |
| *Mucor amethystinus* L. Wagner & G. Walther | 2019 | Armenia |  |  |
| *Mucor amphibiorum* Schipper | 1978 | Australia |  |  |
| *Mucor ardhlaengiktus* B.S. Mehrotra & B.M. Mehrotra | 1979 | India |  |  |
| *Mucor atramentarius* L. Wagner & G. Walther | 2019 | 'Earth' |  |  |
| *Mucor azygosporus* R.K. Benj. | 1963 | USA |  |  |
| *Mucor bacilliformis* Hesselt. | 1954 | USA |  |  |
| *Mucor bondarzevii* S.A. Kulik | 1960 | Russia |  |  |
| *Mucor caatinguensis* A.L. Santiago, C.A.F. de Souza & D.X. Lima | 2016 | Brazil |  |  |
| *Mucor caninus* Pers. | 1796 |  |  |  |
| *Mucor cheongyangensis* Hyang B. Lee & T.T.T. Nguyen | 2020 | Republic of Korea |  |  |
| *Mucor chuxiongensis* C.Y. Chai, W.J. Liu, H. Chen & F.L. Hui | 2019 | China |  |  |
| *Mucor circinatus* D.X. Lima, G. Walther & A.L. Santiago | 2017 | Brazil |  |  |
| *Mucor circinelloides* Tiegh. | 1875 | France | √ | 1973 |
| *Mucor corticola* Hagem | 1910 |  | √ | 2018 |
| *Mucor ctenidius* (Durrell & M. Fleming) Walther & de Hoog | 2013 | USA |  |  |
| *Mucor delicatus* L.S. Loh & Kuthub. | 2001 | Malaysia |  |  |
| *Mucor durus* Walther & de Hoog | 2013 | Great Britain | √ | 2018 |
| *Mucor ellipsoideus* E. Álvarez, Cano, Stchigel, Deanna A. Sutton & Guarro | 2011 | USA |  |  |
| *Mucor endophyticus* (R.Y. Zheng & H. Jiang) J. Pawłowska & Walther | 2013 | China | √ | 2018 |
| *Mucor erectus* Bainier | 1884 | France | √ | 2011 |
| *Mucor eugeniae* L.S. Loh & Nawawi | 2001 |  |  |  |
| *Mucor eugenieae* L.S. Loh & Nawawi | 2001 | Malaysia |  |  |
| *Mucor exponens* (Burgeff) Walther & de Hoog | 2013 |  |  |  |
| *Mucor falcatus* Schipper | 1967 | Germany | √ | 1973 |
| *Mucor flavus* Bainier | 1903 | France | √ | 2009 |
| *Mucor fluvii* Hyang B. Lee, S.H. Lee & T.T.T. Nguyen | 2018 | Republic of Korea |  |  |
| *Mucor fragilis* Bainier | 1884 | France | √ | 1972 |
| *Mucor fuscus* Bainier | 1903 | France |  |  |
| *Mucor fusiformis* Walther & de Hoog | 2013 | Finland |  |  |
| *Mucor genevensis* Lendn. | 1907 | Switzerland | √ | 1974 |
| *Mucor gigasporus* G.Q. Chen & R.Y. Zheng | 1987 | China | √ | 1986 |
| *Mucor griseocyanus* Hagem | 1908 |  |  |  |
| *Mucor guilliermondii* Nadson & Filippov | 1925 |  | √ | 2011 |
| *Mucor hachijyoensis* Ts. Watan. | 1994 | Japan |  |  |
| *Mucor heterogamus* Vuill. | 1887 | France | √ | 2018 |
| *Mucor hiemalis* Wehmer | 1903 |  | √ | 1972 |
| *Mucor inaequisporus* Dade | 1937 | Ghana |  |  |
| *Mucor indicus* Lendn. | 1930 | India | √ | 2007 |
| *Mucor irregularis* Stchigel, Cano, Guarro & E. Álvarez | 2011 | China | √ | 2013 |
| *Mucor japonicus* (Komin.) Walther & de Hoog | 2013 |  |  |  |
| *Mucor koreanus* Hyang B. Lee, S.J. Jeon & T.T. Nguyen | 2016 | Republic of Korea |  |  |
| *Mucor lahorensis* J.H. Mirza, S.M. Khan, S. Begum & Shagufta | 1979 | Pakistan |  |  |
| *Mucor lausannensis* Lendn. | 1907 | Switzerland | √ | 2005 |
| *Mucor laxorrhizus* Y. Ling | 1930 |  |  |  |
| *Mucor lilianae* Voglmayr & Clémençon | 2015 | USA |  |  |
| *Mucor lusitanicus* Bruderl. | 1916 |  |  |  |
| *Mucor luteus* Linnem. ex Wrzosek | 2010 | Germany | √ | 1974 |
| *Mucor megalocarpus* Walther & de Hoog | 2013 | France |  |  |
| *Mucor meguroensis* Ts. Watan. | 1994 | Japan |  |  |
| *Mucor merdicola* C.A.F. de Souza & A.L. Santiago | 2016 | Brazil |  |  |
| *Mucor minutus* (Baijal & B.S. Mehrotra) Schipper | 1975 | India |  |  |
| *Mucor moelleri* (Vuill.) Lendn. | 1908 | Germany | √ | 2018 |
| *Mucor mucedo* Fresen. | 1850 |  | √ | 1972 |
| *Mucor multiplex* (R.Y. Zheng) Walther & de Hoog | 2013 | China |  |  |
| *Mucor nanus* Schipper & Samson | 1994 | Sweden |  |  |
| *Mucor nidicola* A.A. Madden, Stchigel, Guarro, Deanna A. Sutton & Starks | 2012 | USA |  |  |
| *Mucor odoratus* Treschew | 1940 | Denmark |  |  |
| *Mucor orantomantidis* Hyang B. Lee, P.M. Kirk & T.T.T. Nguyen | 2019 | Republic of Korea |  |  |
| *Mucor pakistanicus* J.H. Mirza, S.M. Khan, S. Begum & Shagufta | 1979 | Pakistan |  |  |
| *Mucor parviseptatus* Walther & de Hoog | 2013 | Australia | √ | 2005 |
| *Mucor pernambucoensis* C.L. Lima, D.X. Lima & A.L. Santiago | 2018 | Brazil |  |  |
| *Mucor piriformis* A. Fisch. | 1892 |  | √ | 1973 |
| *Mucor plasmaticus* Tiegh. | 1875 | France |  |  |
| *Mucor plumbeus* Bonord. | 1864 |  | √ | 1987 |
| *Mucor prayagensis* B.S. Mehrotra & Nand ex Schipper | 1978 | India |  |  |
| *Mucor pseudocircinelloides* L. Wagner & G. Walther | 2019 | South Africa |  |  |
| *Mucor pseudolusitanicus* L. Wagner & G. Walther | 2019 | Germany |  |  |
| *Mucor psychrophilus* Milko | 1971 | Russia |  |  |
| *Mucor punjabensis* J.H. Mirza, S.M. Khan, S. Begum & Shagufta | 1979 | Pakistan |  |  |
| *Mucor racemosus* Fresen. | 1850 |  | √ | 1972 |
| *Mucor ramosissimus* Samouts. | 1927 | Russia | √ | 2011 |
| *Mucor ravidus* L.S. Loh & Nawawi | 2001 | Malaysia |  |  |
| *Mucor renisporus* K. Jacobs & Botha | 2008 | South Africa |  |  |
| *Mucor rudolphii* Voglmayr & Clémençon | 2015 | Switzerland |  |  |
| *Mucor saprophilus* Novot. | 1950 | former Soviet Union |  |  |
| *Mucor sarawakensis* L.S. Loh & Nawawi | 2001 | Malaysia |  |  |
| *Mucor saturninus* Hagem | 1910 | Norway | √ | 2011 |
| *Mucor septatum* C.A.F. de Souza, T.R.L. Cordeiro & A.L. Santiago | 2018 | Brazil |  |  |
| *Mucor silvaticus* Hagem | 1908 |  |  |  |
| *Mucor sinensis* Milko & Beliakova | 1971 | former Soviet Union |  |  |
| *Mucor souzae* C.A. de Souza, D.X. Lima & A.L. Santiago | 2018 | Brazil |  |  |
| *Mucor stercorarius* Hyang B. Lee, P.M. Kirk, K. Voigt & T.T.T. Nguyen | 2017 | Republic of Korea |  |  |
| *Mucor stercoreus* (Tode) Link | 1824 |  |  |  |
| *Mucor strictus* Hagem | 1908 |  | √ | 2011 |
| *Mucor subtilissimus* Oudem. | 1898 |  | √ | 1972 |
| *Mucor succosus* Berk. | 1841 | Great Britain |  |  |
| *Mucor suhagiensis* M.D. Mehrotra | 1964 | India |  |  |
| *Mucor sympodialis* L.S. Loh & Kuthub. | 2001 | Malaysia |  |  |
| *Mucor tanatus* L.S. Loh & Nawawi | 2001 | Malaysia |  |  |
| *Mucor thermohyalospora* Subrahm. | 1983 | India |  |  |
| *Mucor troglophilus* Zalar | 1997 | Slovenia |  |  |
| *Mucor ucrainicus* Milko | 1971 | Ukraine |  |  |
| *Mucor variicolumellatus* L. Wagner & G. Walther | 2019 | Germany |  |  |
| *Mucor variosporus* Schipper | 1978 |  | √ | 2018 |
| *Mucor verticillatus* L.S. Loh & Kuthub. | 2001 | Malaysia |  |  |
| *Mucor zonatus* Milko | 1967 | Germany | √ | 2011 |
| *Mycocladus verticillatus* Beauverie | 1900 | France |  |  |
| *Mycotypha guadalcanalensis* Matsush. | 1971 | Solomon Islands |  |  |
| *Mycotypha indica* P.M. Kirk & Benny | 1985 | India |  |  |
| *Mycotypha microspora* Fenner | 1932 |  | √ | 2011 |
| *Nanoglomus plukenetiae* Corazon-Guivin, G.A. Silva & Oehl | 2019 | Peru |  |  |
| *Nawawiella apophysa* L.S. Loh & Kuthub. | 2001 | Malaysia |  |  |
| *Oehlia diaphana* (J.B. Morton & C. Walker) Błaszk., Kozłowska, Niezgoda, B.T. Goto & Dalpé | 2018 | USA |  |  |
| *Otospora bareae* Palenz., N. Ferrol & Oehl | 2008 | Spain |  |  |
| *Pacispora chimonobambusae* (C.G. Wu & Y.S. Liu) Sieverd. & Oehl ex C. Walker, Vestberg & A. Schüßler | 2007 | China | √ | 2018 |
| *Pacispora coralloidea* Sieverd. & Oehl | 2004 | Switzerland |  |  |
| *Pacispora franciscana* Sieverd. & Oehl | 2004 | Italy |  |  |
| *Pacispora patagonica* (Novas & Fracchia) C. Walker, Vestberg & A. Schüßler | 2007 | USA |  |  |
| *Pacispora robigina* Sieverd. & Oehl | 2004 | Switzerland | √ | 2008 |
| *Pacispora scintillans* (S.L. Rose & Trappe) Sieverd. & Oehl ex C. Walker, Vestberg & A. Schüßler | 2007 | USA | √ | 2017 |
| *Paradentiscutata bahiana* Oehl, Magna, B.T. Goto & G.A. Silva | 2012 | Brazil |  |  |
| *Paradentiscutata maritima* B.T. Goto, D.K. Silva, Oehl & G.A. Silva | 2012 | Brazil |  |  |
| *Paraglomus albidum* (C. Walker & L.H. Rhodes) Oehl, G.A. Silva & Sieverd. | 2011 | USA | √ | 2018 |
| *Paraglomus bolivianum* (Sieverd. & Oehl) Oehl & G.A. Silva | 2013 | Bolivia |  |  |
| *Paraglomus brasilianum* (Spain & J. Miranda) J.B. Morton & D. Redecker | 2001 | Brazil | √ | 2009 |
| *Paraglomus laccatum* (Błaszk.) Renker, Błaszk. & Buscot | 2007 | Poland |  |  |
| *Paraglomus lacteum* (S.L. Rose & Trappe) Oehl, G.A. Silva & Sieverd. | 2011 | USA | √ | 2018 |
| *Paraglomus occultum* (C. Walker) J.B. Morton & D. Redecker | 2001 | USA | √ | 2002 |
| *Paraglomus pernambucanum* Oehl, C.M. Mello, Magna & G.A. Silva | 2013 | Brazil | √ | 2016 |
| *Paraglomus turpe* Oehl, V.M. Santos & Palenz. | 2016 | Switzerland |  |  |
| *Parasitella parasitica* (Bainier) Syd. | 1903 |  |  |  |
| *Peridiospora reticulata* C.G. Wu & Suh J. Lin | 1997 | China | √ | 1997 |
| *Peridiospora tatachia* C.G. Wu & Suh J. Lin | 1997 | China | √ | 1997 |
| *Pervetustus simplex* Błaszk., Chwat, Kozłowska, Crossay, Symanczik & Al-Yahya'ei | 2017 | Poland |  |  |
| *Phascolomyces articulosus* Boedijn ex Benny & R.K. Benj. | 1976 | Indonesia |  |  |
| *Phycomyces blakesleeanus* Burgeff | 1925 |  | √ | 1973 |
| *Phycomyces microsporus* Tiegh. | 1875 | France |  |  |
| *Phycomyces nitens* (C. Agardh) Kunze | 1823 | Finland | √ | 1973 |
| *Pilaira anomala* (Ces.) J. Schröt. | 1886 |  | √ | 2009 |
| *Pilaira australis* Urquhart, Coulon & Idnurm | 2017 | Australia |  |  |
| *Pilaira caucasica* Milko | 1970 | Armenia | √ | 2018 |
| *Pilaira dimidiata* Grove | 1884 | Great Britain |  |  |
| *Pilaira moreaui* Y. Ling | 1926 |  |  |  |
| *Pilaira praeampla* R.Y. Zheng & X.Y. Liu | 2009 | China | √ | 2009 |
| *Pilaira subangularis* R.Y. Zheng & X.Y. Liu | 2009 | China | √ | 2009 |
| *Pilobolus crystallinus* (F.H. Wigg.) Tode | 1784 | Germany | √ | 1973 |
| *Pilobolus kleinii* Tiegh. | 1878 | France | √ | 1972 |
| *Pilobolus lentiger* Corda | 1837 | Czech Republic |  |  |
| *Pilobolus longipes* Tiegh. | 1878 | France | √ | 2005 |
| *Pilobolus minutus* Speg. | 1880 | Argentina |  |  |
| *Pilobolus oedipus* Mont. | 1826 |  |  |  |
| *Pilobolus roridus* (Bolton) Pers. | 1801 |  | √ | 2005 |
| *Pilobolus umbonatus* Buller | 1934 |  | √ | 1940 |
| *Pirella circinans* Bainier | 1882 |  | √ | 2004 |
| *Pirella naumovii* (Milko) Benny & Schipper | 1992 | Armenia |  |  |
| *Poitrasia circinans* (H. Nagan. & N. Kawak.) P.M. Kirk | 1984 | Japan | √ | 2017 |
| *Protomycocladus faisalabadensis* (J.H. Mirza, S.M. Khan, S. Begum & Shagufta) Schipper & Samson | 1994 | Pakistan |  |  |
| *Racocetra beninensis* Oehl, Tchabi & Lawouin | 2009 | Benin | √ | 2014 |
| *Racocetra crispa* F.A. Souza, I.R. Silva, M.B.B. de B. Barreto, B.T. Goto & Oehl | 2018 | Brazil |  |  |
| *Racocetra tropicana* Oehl, B.T. Goto & G.A. Silva | 2011 | Brazil | √ | 2014 |
| *Racocetra undulata* T.C. Lin & C.H. Yen | 2011 | China | √ | 2011 |
| *Radiomyces embreei* R.K. Benj. | 1960 | USA |  |  |
| *Radiomyces mexicanus* Benny & R.K. Benj. | 1992 | Mexico |  |  |
| *Radiomyces spectabilis* Embree | 1959 | USA |  |  |
| *Redeckera fulva* (Berk. & Broome) C. Walker & A. Schüßler | 2010 | Sri Lanka |  |  |
| *Redeckera megalocarpa* (D. Redecker) C. Walker & A. Schüßler | 2010 | France |  |  |
| *Redeckera pulvinata* (Henn.) C. Walker & A. Schüßler | 2010 | Venezuela |  |  |
| *Rhizomucor chlamydosporus* R.Y. Zheng, X.Y. Liu & R.Y. Li | 2009 | China | √ | 2007 |
| *Rhizomucor miehei* (Cooney & R. Emers.) Schipper | 1978 | USA | √ | 1990 |
| *Rhizomucor pakistanicus* M. Qureshi & J.H. Mirza | 1979 | Pakistan |  |  |
| *Rhizomucor pusillus* (Lindt) Schipper | 1978 |  | √ | 1990 |
| *Rhizomucor regularior* (R.Y. Zheng & G.Q. Chen) R.Y. Zheng, X.Y. Liu & R.Y. Li | 2009 | China | √ | 2008 |
| *Rhizomucor tauricus* (Milko & Schkur.) Schipper | 1978 | former Soviet Union |  |  |
| *Rhizophagus aggregatus* (N.C. Schenck & G.S. Sm.) C. Walker | 2016 | USA | √ | 2018 |
| *Rhizophagus antarcticus* (Cabello) C. Walker | 2016 | Antarctica |  |  |
| *Rhizophagus arabicus* Błaszk., Symanczik & Al-Yahya'ei | 2014 | Oman |  |  |
| *Rhizophagus clarus* (T.H. Nicolson & N.C. Schenck) C. Walker & A. Schüßler | 2010 | USA | √ | 2018 |
| *Rhizophagus custos* (C. Cano & Dalpé) C. Walker & A. Schüßler | 2010 | Spain |  |  |
| *Rhizophagus fasciculatus* (Thaxt.) C. Walker & A. Schüßler | 2010 | Canada | √ | 2018 |
| *Rhizophagus intraradices* (N.C. Schenck & G.S. Sm.) C. Walker & A. Schüßler | 2010 | USA | √ | 2017 |
| *Rhizophagus invermaius* (I.R. Hall) C. Walker | 2016 | New Zealand | √ | 2018 |
| *Rhizophagus irregularis* (Błaszk., Wubet, Renker & Buscot) C. Walker & A. Schüßler | 2010 | Poland | √ | 2017 |
| *Rhizophagus litchii* S. Pandey & A.P. Misra | 1971 | China |  |  |
| *Rhizophagus manihotis* (R.H. Howeler, Sieverd. & N.C. Schenck) C. Walker & A. Schüßler | 2010 | Colombia | √ | 2018 |
| *Rhizophagus marattiacearum* (C. West) E.J. Butler | 1939 | New Zealand & Australia |  |  |
| *Rhizophagus melanus* (Sudová, Sýkorová & Oehl) C. Walker | 2016 | Norway |  |  |
| *Rhizophagus natalensis* Błaszk., Chwat & B.T. Goto | 2015 | Brazil |  |  |
| *Rhizophagus prolifer* (Dalpé & Declerck) C. Walker & A. Schüßler | 2010 | France |  |  |
| *Rhizophagus theae* (Zimm.) E.J. Butler | 1939 |  |  |  |
| *Rhizopodopsis javensis* Boedijn | 1959 | Indonesia |  |  |
| *Rhizopus americanus* (Hesselt. & J.J. Ellis) R.Y. Zheng, G.Q. Chen & X.Y. Liu | 2000 | USA | √ | 2007 |
| *Rhizopus arrhizus* A. Fisch. | 1892 |  | √ | 1973 |
| *Rhizopus caespitosus* Schipper & Samson | 1994 | India |  |  |
| *Rhizopus homothallicus* Hesselt. & J.J. Ellis | 1962 | Guatemala |  |  |
| *Rhizopus koreanus* Hyang B. Lee & T.T.T. Nguyen | 2016 | Republic of Korea |  |  |
| *Rhizopus lyococcus* (Ehrenb.) G.Y. Liou, F.L. Lee, G.F. Yuan & Stalpers | 2007 |  | √ | 2018 |
| *Rhizopus megasporus* Boedijn | 1959 | Indonesia |  |  |
| *Rhizopus microsporus* Tiegh. | 1875 | France | √ | 2005 |
| *Rhizopus niveus* M. Yamaz. | 1919 | Japan | √ | 2007 |
| *Rhizopus schipperae* Weitzman, McGough, Rinaldi & Della-Latta | 1996 | USA |  |  |
| *Rhizopus sexualis* (G. Sm.) Callen | 1940 | Great Britain | √ | 1991 |
| *Rhizopus stolonifer* (Ehrenb.) Vuill. | 1902 |  | √ | 1973 |
| *Sacculospora baltica* (Błaszk., Madej & Tadych) Oehl, Palenz., Sánchez-Castro, B.T. Goto, G.A. Silva & Sieverd. | 2011 | Poland |  |  |
| *Sacculospora felinovii* A. Willis, Błaszk., T. Prabhu, Chwat, Góralska, Sashidhar, P. Harris, J. D'Souza, Vaing. & Adholeya | 2016 | India |  |  |
| *Saksenaea erythrospora* E. Álvarez, Cano, Stchigel & Guarro | 2010 | USA |  |  |
| *Saksenaea loutrophoriformis* Deanna A. Sutton, Stchigel, Chander, Guarro & Cano | 2017 | USA |  |  |
| *Saksenaea oblongispora* E. Álvarez, Stchigel, Cano & Guarro | 2010 | Brazil |  |  |
| *Saksenaea trapezispora* Deanna A. Sutton, Stchigel, Wiederh., Guarro & Cano | 2016 | USA |  |  |
| *Saksenaea vasiformis* S.B. Saksena | 1953 | India | √ | 2012 |
| *Sclerocarpum amazonicum* Jobim, Błaszk., Niezgoda, A. Kozłowska & B.T. Goto | 2019 | Brazil |  |  |
| *Sclerogone eucalypti* Warcup | 1990 | Australia |  |  |
| *Scutellospora alborosea* (Ferrer & R.A. Herrera) C. Walker & F.E. Sanders | 1986 | Cuba |  |  |
| *Scutellospora alterata* Oehl, J.S. Pontes, Palenz., I.C. Sánchez & G.A. Silva | 2013 | Brazil |  |  |
| *Scutellospora arenicola* Koske & Halvorson | 1990 | USA | √ | 2014 |
| *Scutellospora aurigloba* (I.R. Hall) C. Walker & F.E. Sanders | 1986 | New Zealand | √ | 1990 |
| *Scutellospora biornata* Spain, Sieverd. & S. Toro | 1989 | Colombia |  |  |
| *Scutellospora calospora* (T.H. Nicolson & Gerd.) C. Walker & F.E. Sanders | 1986 | Great Britain | √ | 1990 |
| *Scutellospora castanea* C. Walker | 1993 | France | √ | 2012 |
| *Scutellospora coralloidea* (Trappe, Gerd. & I. Ho) C. Walker & F.E. Sanders | 1986 | USA | √ | 1997 |
| *Scutellospora crenulata* R.A. Herrera, Cuenca & C. Walker | 2001 | Venezuela | √ | 2008 |
| *Scutellospora dipapillosa* (C. Walker & Koske) C. Walker & F.E. Sanders | 1986 | USA | √ | 2009 |
| *Scutellospora dipurpurescens* J.B. Morton & Koske | 1988 | USA | √ | 2006 |
| *Scutellospora gregaria* (N.C. Schenck & T.H. Nicolson) C. Walker & F.E. Sanders | 1986 | USA | √ | 1998 |
| *Scutellospora hawaiiensis* Koske & Gemma | 1995 | USA |  |  |
| *Scutellospora minuta* (Ferrer & R.A. Herrera) C. Walker & F.E. Sanders | 1986 | Cuba |  |  |
| *Scutellospora nodosa* Błaszk. | 1991 | Poland |  |  |
| *Scutellospora ovalis* D. Redecker, Crossay & Cilia | 2018 | France |  |  |
| *Scutellospora pernambucana* Oehl, D.K. Silva, N. Freitas & L.C. Maia | 2009 | Brazil |  |  |
| *Scutellospora persica* (Koske & C. Walker) C. Walker & F.E. Sanders | 1986 | USA | √ | 2002 |
| *Scutellospora projecturata* Kramad. & C. Walker | 2000 | Indonesia |  |  |
| *Scutellospora rubra* Stürmer & J.B. Morton | 1999 | Brazil | √ | 2012 |
| *Scutellospora scutata* C. Walker & Dieder. | 1989 | Brazil |  |  |
| *Scutellospora spinosissima* C. Walker & Cuenca | 1998 | Venezuela |  |  |
| *Scutellospora striata* Cuenca & R.A. Herrera | 2008 | Venezuela |  |  |
| *Scutellospora tepuiensis* E. Furrazola & Cuenca | 2017 | Venezuela |  |  |
| *Scutellospora tricalypta* (R.A. Herrera & Ferrer) C. Walker & F.E. Sanders | 1986 | Cuba | √ | 2014 |
| *Scutellospora trirubiginopa* X.L. Pan & G.Yun Zhang | 1997 | China | √ | 1997 |
| *Scutellospora verrucosa* (Koske & C. Walker) C. Walker & F.E. Sanders | 1986 | USA | √ | 2003 |
| *Scutellospora weresubiae* Koske & C. Walker | 1986 | USA |  |  |
| *Sieverdingia tortuosa* (N.C. Schenck & G.S. Sm.) Błaszk., Niezgoda & B.T. Goto | 2019 | USA |  |  |
| *Spinellus arvernensis* L. Ling | 1930 |  |  |  |
| *Spinellus chalybeus* (Dozy & Molk.) Vuill. | 1904 |  |  |  |
| *Spinellus fusiger* (Link) Tiegh. | 1875 |  | √ | 2008 |
| *Spinellus gigasporus* Cooke & Massee | 1889 | Australia |  |  |
| *Spinellus sphaerosporus* Tiegh. | 1875 | France |  |  |
| *Sporodiniella umbellata* Boedijn | 1959 | Indonisia | √ | 1997 |
| *Syncephalastrum alma-ataense* Novobr. | 1972 | Kazakhstan |  |  |
| *Syncephalastrum contaminatum* A.S. Urquhart & A. Idnurm | 2020 |  |  |  |
| *Syncephalastrum monosporum* R.Y. Zheng, G.Q. Chen & F.M. Hu | 1988 | China | √ | 1988 |
| *Syncephalastrum racemosum* Cohn ex J. Schröt. | 1886 | Poland | √ | 1973 |
| *Syzygites megalocarpus* Ehrenb. | 1818 | Germany |  |  |
| *Thamnidium elegans* Link | 1809 | Germany | √ | 1973 |
| *Thamnostylum lucknowense* (J.N. Rai, J.P. Tewari & Mukerji) Arx & H.P. Upadhyay | 1970 | India |  |  |
| *Thamnostylum nigricans* (Tiegh.) Benny & R.K. Benj. | 1975 | France |  |  |
| *Thamnostylum piriforme* (Bainier) Arx & H.P. Upadhyay | 1970 |  | √ | 2018 |
| *Thamnostylum repens* (Tiegh.) H.P. Upadhyay | 1973 | France |  |  |
| *Thermomucor indicae-seudaticae* Subrahm., B.S. Mehrotra & Thirum. | 1977 | India |  |  |
| *Tortumyces cameronensis* L.S. Loh | 2001 | Malaysia |  |  |
| *Tortumyces fimicola* L.S. Loh | 2001 | Malaysia |  |  |
| *Tricispora nevadensis* (Palenz., N. Ferrol, Azcón-Aguilar & Oehl) Oehl, Palenz., G.A. Silva & Sieverd. | 2011 | Spain |  |  |
| *Umbelopsis angularis* W. Gams & M. Sugiy. | 2003 | Netherlands | √ | 2013 |
| *Umbelopsis autotrophica* (E.H. Evans) W. Gams | 2003 | Great Britain |  |  |
| *Umbelopsis changbaiensis* Y.N. Wang, X.Y. Liu & R.Y. Zheng | 2014 | China | √ | 2014 |
| *Umbelopsis dimorpha* Mahoney & W. Gams | 2004 | New Zealand | √ | 2013 |
| *Umbelopsis fusiformis* H.Y. Yip | 1986 | Australia |  |  |
| *Umbelopsis gibberispora* M. Sugiy., Tokum. & W. Gams | 2003 | Japan |  |  |
| *Umbelopsis isabellina* (Oudem.) W. Gams | 2003 |  | √ | 2010 |
| *Umbelopsis longicollis* (Dixon-Stew.) Y.N. Wang, X.Y. Liu & R.Y. Zheng | 2015 |  |  |  |
| *Umbelopsis nana* (Linnem.) Arx | 1982 | Germany | √ | 2013 |
| *Umbelopsis ovata* (H.Y. Yip) H.Y. Yip | 1986 | Australia |  |  |
| *Umbelopsis ramanniana* (Möller) W. Gams | 2003 |  | √ | 2011 |
| *Umbelopsis sinsidoensis* Hyang B. Lee & T.T.T. Nguyen | 2018 | Republic of Korea |  |  |
| *Umbelopsis swartii* H.Y. Yip | 1986 | Australia |  |  |
| *Umbelopsis versiformis* Amos & H.L. Barnett | 1966 | USA | √ | 2002 |
| *Umbelopsis vinacea* (Dixon-Stew.) Arx | 1982 | Australia | √ | 2011 |
| *Umbelopsis westeae* H.Y. Yip | 1986 | Australia |  |  |
| *Umbelopsis wiegerinckiae* Sand.-Den. | 2017 | Netherlands |  |  |
| *Utharomyces epallocaulus* Boedijn ex P.M. Kirk & Benny | 1980 | China |  |  |
| *Vinositunica ingens* Koh. Yamam., Degawa & A. Yamada | 2020 | Japan |  |  |
| *Vinositunica radiata* Koh. Yamam., Degawa & A. Yamada | 2020 | Japan |  |  |
| *Zychaea mexicana* Benny & R.K. Benj. | 1975 | Mexico |  |  |
